# BCL-XL blockage in TNBC models confers vulnerability to inhibition of specific cell cycle regulators

**DOI:** 10.1101/2021.03.16.435600

**Authors:** Olivier Castellanet, Fahmida Ahmad, Yaron Vinik, Gordon B. Mills, Bianca Habermann, Jean-Paul Borg, Sima Lev, Fabienne Lamballe, Flavio Maina

## Abstract

Cell cycle regulators are frequently altered in Triple-Negative Breast Cancer (TNBC). Emerging agents targeting these signals offer the possibility to design new combinatorial therapies. However, preclinical models that recapitulate TNBC primary resistance and heterogeneity are essential to evaluate the potency of these combined treatments.

**Methods:** Bioinformatic processing of human breast cancer datasets was used to analyse correlations between expression levels of cell cycle regulators and patient survival outcome. The *MMTV-R26*^*Met*^ mouse model of TNBC resistance and heterogeneity was employed to analyse expression and targeting vulnerability of cell cycle regulators in the presence of BCL-XL blockage. Robustness of outcomes and selectivity was further explored using a panel of human breast cancer cells. Alterations of protein expression, phosphorylation, and/or cellular localisation were analysed by western blots, reverse phase protein array, and immunocytochemistry. Bioinformatics was performed to highlight drug’s mechanisms of action.

**Results:** We report that high expression levels of BCL-XL and specific cell cycle regulators correlate with poor survival outcomes of TNBC patients. Blockage of BCL-XL confers vulnerability to drugs targeting CDK1/2/4, but not FOXM1, CDK4/6, Aurora A and Aurora B, to all *MMTV-R26*^*Met*^ and human TNBC cell lines tested. Mechanistically, we show that, co-targeting of BCL-XL and CDK1/2/4 synergistically inhibited cell growth by combinatorial depletion of survival and RTK/AKT signals, and concomitantly restoring FOXO3a tumour suppression actions. This was accompanied by an accumulation of DNA damage and consequently apoptosis.

**Conclusions:** Our studies illustrate the possibility to exploit the vulnerability of TNBC cells to CDK1/2/4 inhibition by targeting BCL-XL. Moreover, they underline that specificity matters in targeting cell cycle regulators for combinatorial anticancer therapies.

## Introduction

Targeting of the anti-apoptotic protein BCL-XL together with anti-mitotic agents has been proposed as an efficient therapeutic strategy for different human cancers, including triple-negative breast cancer (TNBC), a particular aggressive subtype of breast cancer [1]. TNBC is defined by the lack of oestrogen (ER) and progesterone (PR) receptors and the absence of amplification/overexpression of the HER2 receptor tyrosine kinase (RTK) [2]. The disease accounts for ∼15% of all breast cancer types and is characterised by an extraordinary molecular heterogeneity. Current therapeutic options for targeted treatment are limited due to primary or acquired resistance, and major efforts are devoted to search for molecular alterations and predict vulnerabilities for effective targeted therapy [3]. However, the high heterogeneity of TNBC challenges the identification of generic targets, and highlights the requirement for precision therapy and preclinical models that recapitulate TNBC heterogeneity.

We have recently reported the generation of a rather unique mouse model in which a subtle increase in the wild-type MET RTK expression levels in the mammary gland (*MMTV-R26*^*Met*^ mice) leads to spontaneous tumour formation faithfully recapitulating TNBC features [4]. These include histological, molecular and signalling heterogeneity, as well as primary resistance to chemotherapy and approved targeted therapies [5, 6]. We further exploited the *MMTV-R26*^*Met*^ TNBC model to identify a highly effective therapeutic protocol based on the combined inhibition of the anti-apoptotic molecule BCL-XL and of the cell cycle checkpoint regulator WEE1 [4]. Cell cycle proteins are frequently overexpressed and/or overactivated in several cancer types including TNBC. For example, loss of RB or p16^INK4^ is frequent across TNBC subtypes [7, 8]. Alterations in cyclin D and E, CDK4/6, and CDK2 in TNBC have also been reported [9]. Furthermore, mutations in p53, a master cell cycle regulator of the G1/S checkpoint, are present in a large proportion of TNBC, thus leaving cells to mainly rely on the G2/M checkpoint to maintain DNA integrity [10, 11]. Therefore, targeting cell cycle regulators provide a clear rationale for designing anti-cancer therapy [10]. The vulnerability of TNBC to inhibitors of cell cycle regulators has been demonstrated by several studies using different drugs targeting distinct signalling components. For example, the CDK4/6 inhibitor Palbociclib, effectively used in ER-positive breast cancer, has recently been explored in combination with chemotherapy in RB-positive TNBC cell models [12-14]. Additionally, it has been reported that a combinatorial treatment based on CDK4/6 plus BET inhibitors leads to cell division errors and death, although leading to the emergence of heterogeneous mechanisms of resistance [15].

In the present study, we employed the *MMTV-R26*^*Met*^ model system that recapitulates TNBC heterogeneity and primary resistance to explore the effect of inhibiting specific cell cycle regulators while blocking BCL-XL function. Our results show that targeting specific cell cycle components is important and uncover a detrimental effect on TNBC cells of CDK1/2/4 plus BCL-XL inhibition. We provide evidence that this combinatorial targeting significantly reduces the levels of RTK and AKT signalling besides perturbing cell cycle and DNA repair.

## Methods

### Mouse cell lines

*MMTV-R26*^*Met*^ MGT cell lines were derived from independent *MMTV-R26*^*Met*^ tumours and established as previously described [4]. Cells were cultured in complete DMEM/F12 (Dulbecco’s modified Eagle’s media/F12, 1/1, ThermoFisher Scientific) medium (DMEM/F12, supplemented with 10% foetal bovine serum (FBS, ThermoFisher Scientific), penicillin-streptomycin (P/S, 100 U/ml/0.1 mg/ml, ThermoFisher Scientific), glutamine (2mM, ThermoFisher Scientific), glucose (0.25%, Sigma), insulin (10 µg/ml, Sigma), transferrin (10µg/ml, Sigma), sodium selenite (5ng/ml, Sigma), hydrocortisone (0.5 µg/ml, Sigma), EGF (20ng/ml, Roche), and HGF (10 ng/ml, Peprotech), at 37C° in a 5% CO_2_ atmosphere. PCR-based assays were performed on all cell lines to verify that they were free of *Mycoplasma* contamination. Mouse embryonic fibroblasts were kindly provided by P. Perrin, either untreated or treated with Mitomycin C (Sigma).

### Human cell lines

All human TNBC (MDA-MB-231, MDA-MB-468, SUM-159, Hs578t, HCC-1937 and BT-549) and non-TNBC (MCF-7, SKBR-3, and BT-474) cell lines used in this study were obtained from the American Type Culture Collection (ATCC) without further authentication and were tested by PCR-based assay to verify that they were free of *Mycoplasma* contamination. These human breast cancer cells were grown in DMEM/F12 medium supplemented with P/S, glutamine (2mM), sodium pyruvate (1mM, ThermoFisher Scientific), non-essential amino acids (ThermoFisher Scientific), and insulin (10µg/ml).

### Drugs

Drug concentration and sources are reported in Table S9. Calculation of the Synergy maps and Bliss score has been performed using online SynergyFinder tool v1.0 [16] using “Viability” parameter as readout and “Bliss Method” with correction activated. The CompuSyn software v1.0 using the Chou-Talalay equation was used to measure synergy or additive effects of drug combinations. Combination index CI<1 indicates synergism, CI<0.5 indicates strong synergism, CI=1 means additive effect and CI>1 stands for antagonism.

### Cell viability assay

Cell viability assay was performed on *MMTV-R26*^*Met*^ MGT and human breast cancer cells as previously described [4]. Briefly, cells were plated in 96-well plates, and treated after 24hrs with either single or combined drugs at the indicated concentrations. Cell viability was determined 48hrs later using the Cell Titer Glo Luminescent Assay (Promega). Data are mean values of at least three independent experiments done in triplicates.

### Cell cycle analysis by flow cytometry

*MMTV-R26*^*Met*^ cells were treated for 12hrs with vehicle, A1155463 (A11, 1µM), R547 (3µM), alone or in combination. After trypsinisation, cell suspension was then processed as previously described [4]. Three independent experiments were performed.

### Immunocytochemistry

Protocols used were as described in [4]. Percentage of TritonX-100 and antibodies used for immunofluorescence staining are detailed in Table S8. Quantification of nucleus versus cytoplasmic staining intensity was determined using Image J Intensity Ratio Nuclei Cytoplasm Tool, RRID:SCR_018573.

### Western blotting

Protein extracts were prepared, and western blot analysis was performed as previously described [4]. The antibodies used are reported in Table S8. ACTIN and Ponceau staining were used as loading controls. Ponceau stainings are shown in supplementary information (non-edited gels). For densitometric analysis, the intensity of each protein band was measured using the ImageJ software (National Institutes of Health, USA).

### Reverse phase protein array (RPPA)

Protein lysates of *MMTV-R26*^*Met*^ MGT cells treated or not with A1155463 (1µM), R547 (3µM), or the drug combination A1155463 + R547 (1µM, 3µM) for 12hrs, were prepared according to the MD Anderson Cancer Center platform instructions. RPPA of cells treated with Adavosertib (3µM), or A1155463 + Adavosertib (1µM, 3µM) was previously reported [4], and used in this study to compare signalling changes occurring with the different drug combinations. Samples were screened with 426 antibodies to identify signalling changes in protein expression and phosphorylation levels.

### RNA-seq

Total RNA from dissected *MMTV-R26*^*Met*^ tumours (n=4) and control mammary gland tissues (n=3) was processed for transcriptome analysis. RNA integrity was assessed using the Agilent RNA 6000 Pico kit and Agilent 2100 Bioanalyzer (Agilent Technologies, Santa Clara, California) according to the manufacturer’s instructions. Sequencing was performed as previously described [17].

### Bioinformatic analysis

Analysis of publicly available microarray data: Kaplan-Meier curve reporting the probability (in percent) of the overall survival of human TNBC patients according to BCL-XL and/or specific cell cycle regulator levels. The NCBI dataset used was the GSE31519 Affymetrix Human Genome U133A Array. The database includes 580 TNBC patients phenotyped by immunohistochemistry.

Raw RNA-seq reads were mapped against the latest release (mm10) of the mouse genome using STAR aligner [18]. We used featureCounts [19] to quantify mapped reads. Differential expression analysis was performed with the RStudio (RStudio, Inc., Boston, MA) software using the DESeq2 package [20]. For data shown in Figure 3C and S2A, a cut-off p-value < 0.05 was applied. The Genseset enrichment analysis (GSEA) software [21, 22] was used on the WikiPathways database to highlight enriched pathways in the *MMTV-R26*^*Met*^ tumours versus controls. Ranking of the enriched pathways was then performed using the Enrichr software. RPPA analyses of drug perturbation effects were done using biological duplicates. Expression levels of proteins were Log2 transformed before analysis. GSEA was employed to determine the statistically significant enriched pathways between cells treated with either A11+R547 or A11+Adavosertib. Differential gene expressions were obtained from the RPPA data outcomes. GSEA v4.1.0 was obtained from the Broad Institute (http://software.broadinstitute.org/gsea/index.jsp). The collection of annotated gene sets of REACTOME was obtained from the GSEA website (MSigDB, http://www.gsea-msigdb.org/gsea/msigdb/index.jsp).

### Statistical analysis

The probability of overall survival rates was calculated using the Kaplan-Meier method. P values were computed using the Logrank (Mantel Cox) test. P values are indicated in figures. P > 0.05 was considered as non-significant (ns). * P<0.05; ** P<0.01; *** P <0.001. For RPPA studies, analysis of fold-change proteins and p-values to determine significantly differentially expressed proteins were done by the Limma package in R. Data are presented as the mean ± standard error of the mean (s.e.m.), according to sample distributions. For two sided comparisons, unpaired Student’s t test was used for data showing normal distributions, and Wilcoxon test was used in other situations. For multiple comparisons, we used ANOVA test followed by Tukey test. All statistical analyses were performed using the GraphPad Prism and software.

## Results

### High expression levels of BCL-XL and of selective cell cycle regulators correlate with poor survival outcomes of TNBC patients

Recent studies using breast cancer tissue-specific microarray databases (GSE42568, GSE45827, and GSE54002) have reported a strong enrichment in cell cycle pathway genes [23], which we also illustrate in Figure 1A. Common up-and down-regulated genes from these three microarray datasets were highly enriched in cell cycle regulation pathways from KEGG and WikiPathways (Figure 1B, Table S1). Analysis of the Pan-Cancer Atlas cohort of the TCGA database revealed that about 50% of breast cancer patients have altered cell cycle genes, with breast cancer ranking second among all cancer types analysed for the percentage of amplification events (Figure 1C). Among the four breast cancer sub- types, HER2^+^ and TNBC patients show the highest frequency of alteration in cell cycle regulators (Figure 1D). Gene amplification is the major type of alterations found in breast cancer patients (Figure 1D). We next used a TNBC microarray database (GSE31519; 580 patients) to analyse the relevance of cell cycle gene levels in patient outcomes. We found that TNBC patients with high levels of *BCL-XL* or specific cell cycle regulators such as *CDK1, CDK6* and **WEE1**, exhibit a shorter overall survival rate (Figure 2A and B, Table S2). Interestingly, patients that concurrently highly express the anti-apoptotic factor *BCL-XL* and the cell cycle modulators *CDK1, CDK6*, or *WEE1*, have poor clinical outcome, thus highlighting the detrimental effect of their concomitant high expression levels (Figure 2C, Table S2). Intriguingly, high levels of other cell cycle modulators such as *CDK2, CDK4, FOXM1, Aurora A*, or *Aurora B*, either alone or together with *BCL-XL* levels, is not associated with altered patient survival (Figure 2B and C, Table S2). The apparent significant difference between the survival curves in the combination sets is mainly due to *BCL-XL* levels (Figure 2C, Table S2). Furthermore, patients with high levels of *BCL-XL, CDK1*, and *WEE1* exhibit more aggressive TNBC tumours (classified as grade 3; Figure 2D). This was also observed taking into consideration co-expression of high versus low *BCL-XL* with *CDK1* or *WEE1* (Figure 2E). This significant increase in aggressiveness was not observed in patients with high levels of *CDK6* (Figure 2D and E). These results may suggest a greater implication of specific cell cycle regulators, particularly when overexpressed with BCL-XL, on survival of TNBC patients.

**Figure 1.**
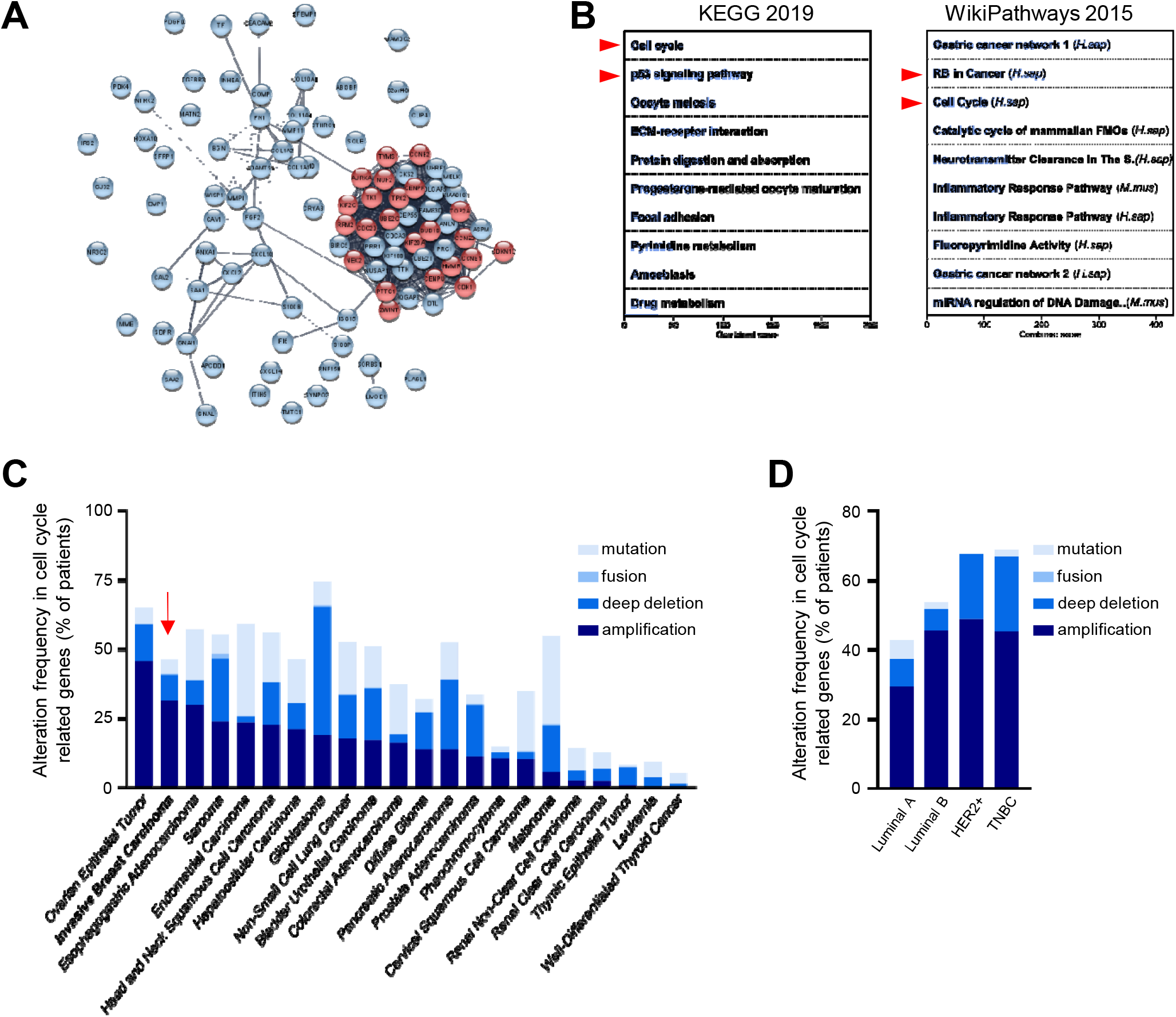
Cell cycle regulators are enriched and altered in breast cancer. **(A-B)** Human breast tumours are enriched in cell cycle regulators. **(A)** Analysis from three breast cancer databases (GSE42568, GSE45827 and GSE54002) delineate 97 common differential expressed genes, as reported in [23]. In the STRING network, proteins are represented by the nodes, and interaction between proteins by edges. Out of the 97 genes, 23 are involved in cell cycle regulation according to the Reactome database (red circles; see Table S1). See Figure S1A for high magnification. **(B)** Enrichment pathway analyses of the 97 differentially expressed genes using the Enrichr software, according to KEGG 2019 and WikiPathways 2015 databases, ordered according to the combined score. The 10-top ranked enriched pathways are shown and highlight signals involved in cell cycle regulation (red arrowhead). **(C-D)** Analysis of the Pan-Cancer Atlas cohort (TCGA database) of cell cycle-related gene alterations. Only patients with mutations and copy number variations data were selected for this analysis. Information regarding cell cycle-related gene alterations have been extracted with the use of cBioPortal for Cancer Genomics. The histograms report frequency (% of patients) of cell cycle-related gene alterations in different cancer types **(C)** and in the four breast cancer sub-types **(D)**. Alterations include mutation, gene fusion, deep deletion, and amplification. Note that breast cancer (indicated by a red arrow) ranks second among all cancer types analysed, sorted by amplification events.

**Figure 2.**
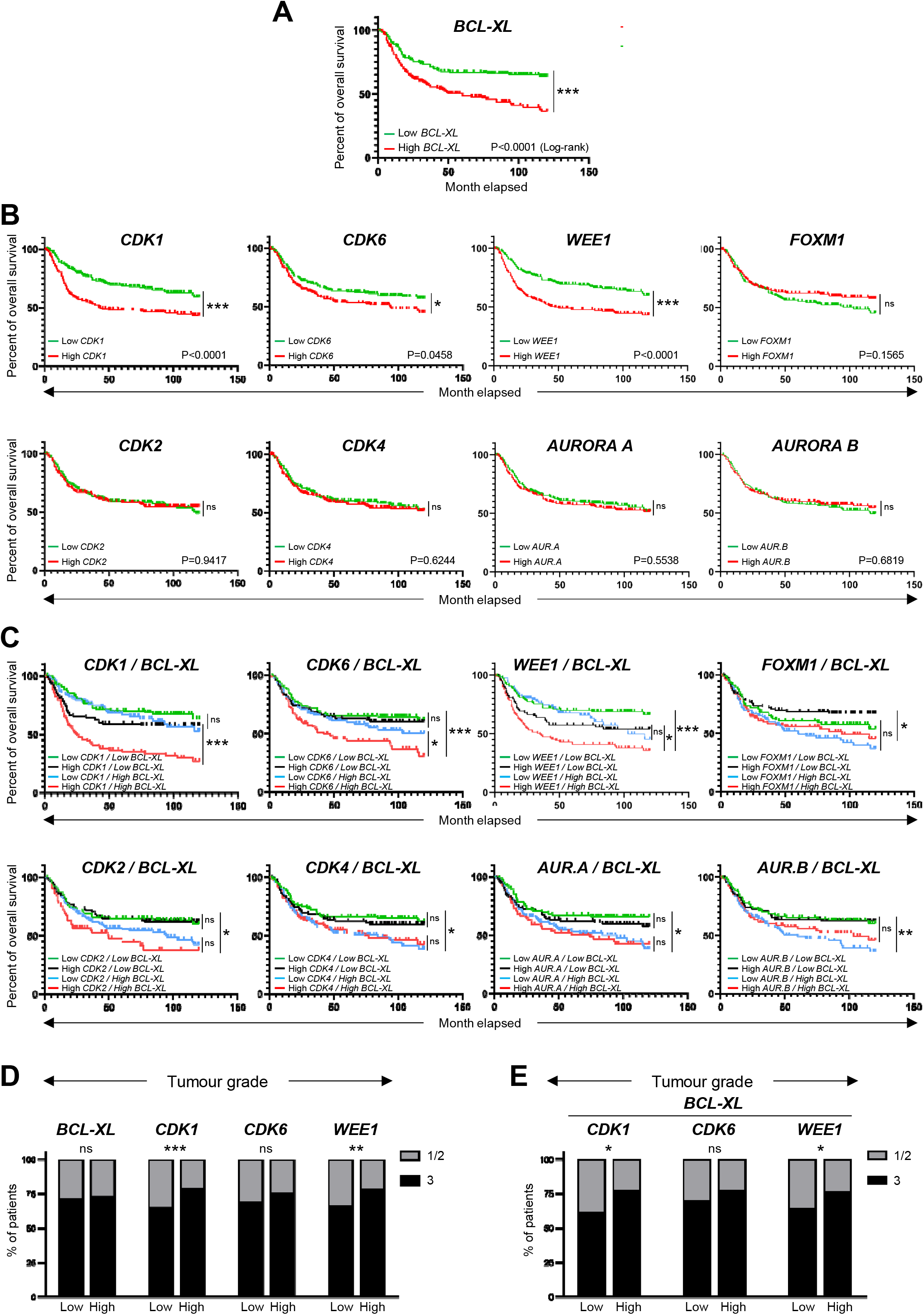
High expression levels of *BCL-XL, CDK1, CDK6*, and *WEE1* correlate with lower survival rate of TNBC patients. **(A-C)** Analysis of publicly available microarray data from 580 TNBC patients (GSE31519 Affymetrix Human Genome U133A Array). Kaplan-Meier curves reporting the probability of the overall survival of human TNBC patients according to the expression levels of *BCL-XL* **(A)**, specific cell cycle regulators (*CDK1, CDK2, CDK4, CDK6, WEE1, FOXM1, Aurora A*, and *Aurora B*) alone **(B)** or in combination with *BCL-XL* **(C)**. The median of each gene expression levels was used as a threshold to segregate high versus low expressers. P values were computed using the Logrank (Mantel Cox) test. **(D-E)** Tumour grades reported in patients with high or low levels of either single **(D)** or combined **(E)** indicated genes. Tumour grades were defined according to standard pathological scores (1-3; 3 corresponds to the most aggressive grade). ns: not significant; * P<0.05; ** P<0.01; ***p<0.001.

### TNBC cells are vulnerable to inhibition of selective cell cycle regulators following BCL-XL blockage

The above findings, together with our recent studies uncovering the vulnerability of TNBC cells to combined WEE1 and BCL-XL targeting [4], drove us to explore the sensitivity of TNBC cells to inhibition of specific cell cycle regulators in combination with BCL-XL targeting. We addressed this issue using the *MMTV-R26*^*Met*^ TNBC model system, employing RNA-seq analysis, proteomic profiling, and cell viability assay as illustrated in Figure 3A. By Gene Set Enrichment Analysis (GSEA) of RNA-seq outcomes comparing *MMTV-R26*^*Met*^ tumours (n=4) to control mammary gland tissues (*MMTV-Cre*; n=3), we found a striking enrichment in genes related to cell cycle regulation (Figure 3B), and to DNA replication and DNA damage pathways (Figure S2A, B, and Table S3). Specifically, we found significantly high mRNA levels of *FoxM1* (a member of the Forkhead superfamily of transcription factors regulating a plethora of genes throughout the cell cycle to control DNA replication, mitosis, and cell proliferation), *Aurora A* and *B* (cell cycle regulated kinases involved in microtubule formation and/or stabilization at the spindle pole during chromosome segregation), and Cyclin-dependent kinases *Cdk1, Cdk2*, and *Cdk4* (which control progression through the cell cycle in concert with their cyclin regulatory subunits). Specific cyclins such as *Cyclin A2, B1, B2*, and *E1* were also upregulated in *MMTV-R26*^*Met*^ tumours (see Table S4). Instead, comparable levels between tumours and normal mammary glands were found for *Cdk3, Cdk5*, and *Cdk6* (Figure 3C).

**Figure 3.**
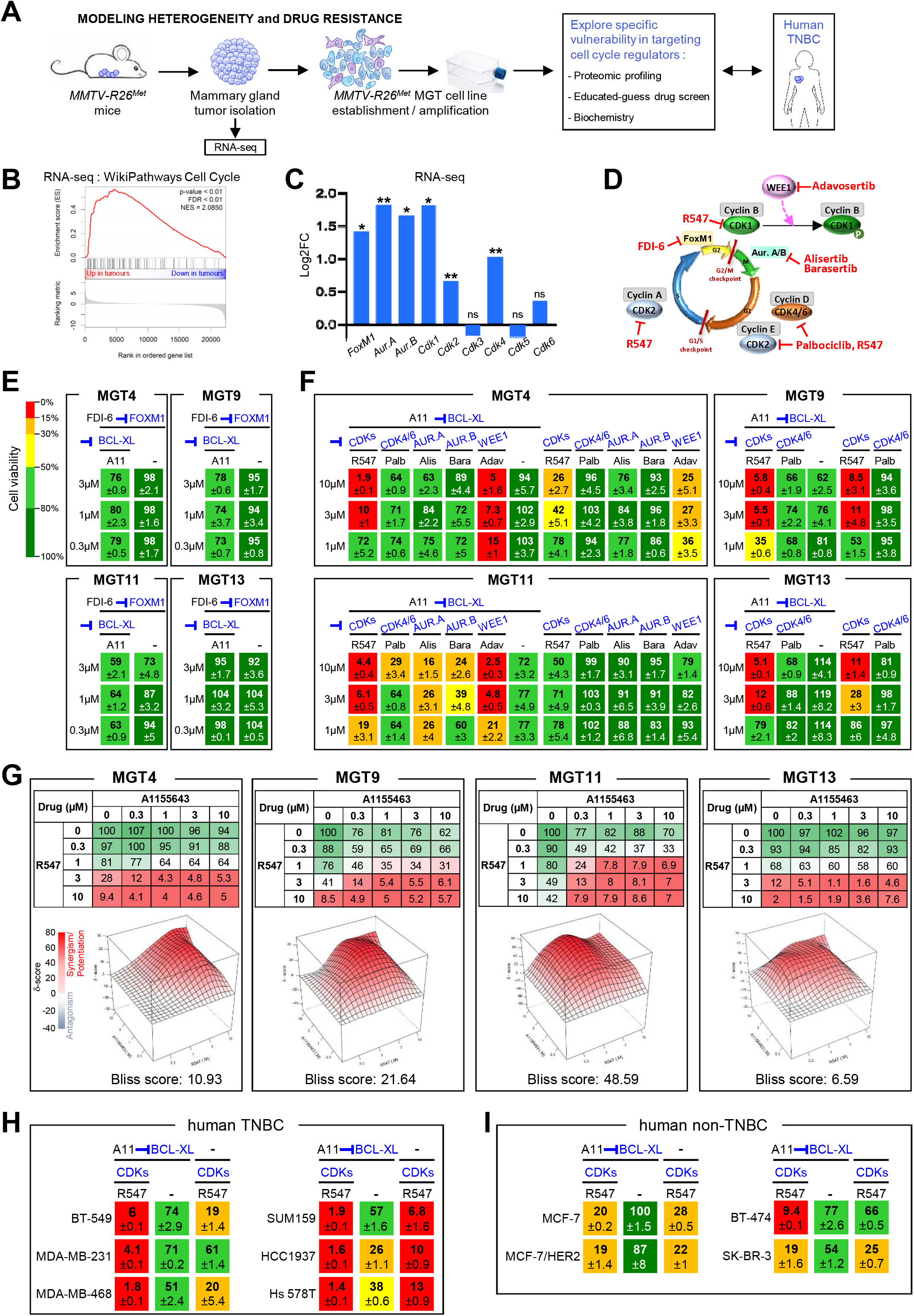
Combined inhibition of BCL-XL and CDK1/2/4 is deleterious for *MMTV-R26*^*Met*^ and human TNBC cell lines. **(A)** Scheme illustrating the strategy used in this study to explore the vulnerability of TNBC cells to the inhibition of specific cell cycle regulators in combination with BCL-XL targeting. RNA-seq studies were performed on *MMTV-R26*^*Met*^ tumours. Proteomic profiling, “educated-guess” drug screens and biochemical analyses were carried out on cell lines established from the *MMTV-R26*^*Met*^ TNBC mouse model treated with drugs targeting specific cell cycle regulators together with BCL-XL. **(B)** GSEA enrichment performed, using the WikiPathways database, with cell cycle geneset on *MMTV-R26*^*Met*^ tumours versus controls. The barcode plot indicates the position of a member of this geneset in the ranked list. Red and blue colours represent up-or downregulated genes in the *MMTV-R26*^*Met*^ tumours versus the controls, respectively. NES: normalized enrichment score; FDR: false discovery rate. **(C)** Histogram depicting upregulation of *FoxM1, Aurora A, Aurora B, Cdk1, Cdk2*, and *Cdk4* transcript levels (as Log_2_FC; from RNA-seq data) in *MMTV-R26*^*Met*^ tumours versus control tissues. Note that *Cdk3, Cdk5*, and *Cdk6* mRNA levels were similar to controls. **(D)** Scheme illustrating key cell cycle regulators and their inhibitors used in this study. Their position within the cell cycle corresponds to the phase in which they act. **(E)** Effects of FOXM1 inhibition (FDI-6, 0.3-3µM) combined with BCL-XL targeting (A11: A1155463, 0.3µM) were analysed on the viability of the four tumorigenic *MMTV-R26*^*Met*^ MGT cell lines. Numbers indicate percentage of cell viability in the presence of drugs, compared to untreated cells (used as control). Percentages are reported using a green (high)-to-red (low) colour code (the scale depicted on the left is used as a reference in all studies). **(F)** Cell viability of the four *MMTV-R26*^*Met*^ cell lines exposed to A1155463 (A11; targeting BCL-XL) alone or in combination with specific cell cycle regulators: R547 (targeting CDK1/2/4), Palbociclib (Palb; CDK4/6 inhibitor), Alisertib (Alis; targeting Aurora A), Barasertib (Bara; targeting Aurora B). Adavosertib (Adav; targeting WEE1) was used as control. **(G)** Top panel: Detailed matrix analysis of the MGT4, MGT9, MGT11 and MGT13 cell viability in response to A1155463 alone or in combination with R547. Bottom panel: Loewe plots and Bliss scores highlight synergism of the two drugs. **(H-I)** Cell viability of a panel of human TNBC **(H)** and human non-TNBC **(I)** cell lines when exposed to A1155463 (1µM) and R547 (3µM). In all figures, cell viability is presented as percentage of untreated cells (used as control). Values are expressed as means ± s.e.m. At least three independent experiments were performed. For RNAseq studies, statistical analyses were performed using “R”. Adjusted p-values are reported. ns: not significant; * P<0.05; ** P<0.01. Aur.A: Aurora A; Aur.B: Aurora B; CDK: Cyclin-dependent kinase.

In light of these results, we evaluated cell viability of mammary gland tumour (MGT) cell lines (MGT4, 9, 11, and 13) derived from the *MMTV-R26*^*Met*^ model [4] when treated with inhibitors of various cell cycle regulators, acting during different phases, as illustrated in Figure 3D. Inhibition of either FOXM1 (with FDI-6), CDK4/6 (with Palbociclib), Aurora A or B (with Alisertib or Barasertib, respectively) together with BCL-XL blockage (with A1155463) did not alter the viability of the *MMTV-R26*^*Met*^ TNBC cells we tested (Figure 3D-F). We observed only a partial response in MGT11 cells following combined inhibition of BCL-XL with CDK4/6, Aurora A or Aurora B (Figure 3F). In contrast, combined targeting of CDK1/2/4 (with R547) and BCL-XL (with A1155463) was highly deleterious for all *MMTV-R26*^*Met*^ TNBC cells we tested (Figure 3D and F). Concomitant inhibition of CDK1/2/4 and BCL-XL was synergistic for 3 out of 4 *MMTV-R26*^*Met*^ MGT cell lines, as shown by the Bliss score and the Chou-Talalay combination index score calculation (Figure 3G and S2C). These results were corroborated by the deleterious effect of CDK1/2/4 and BCL-XL co-targeting we observed in all six human TNBC cell lines tested (Figure 3H). In contrast, we found only a modest effect on human non-TNBC cells (with the exception of the BT-474 cells), similar to the non-tumorigenic *MMTV-R26*^*Met*^ MGT2 cells (Figure 3I and S2D). These findings illustrate the vulnerability of TNBC cells to the inhibition of specific cell cycle regulators when BCL-XL is targeted.

### Combined BCL-XL and CDK1/2/4 inhibition interferes with cell cycle and survival signals, and triggers apoptosis

We next examined, in the context of BCL-XL blockage (by A1155463), the molecular and biological consequences of CDK1/2/4 targeting (by R547) [24], and assessed specificity in alterations compared to WEE1 inhibition (by Adavosertib). We first assessed the status of cell cycle regulators and observed a drastic downregulation of both RB protein expression and phosphorylation levels following CDK1/2/4 targeting (R547), in contrast to unchanged levels following WEE1 inhibition (Adavosertib) (Figure 4A, B, and S3). This is consistent with RB being a direct target of CDK2 and CDK4/6 [25]. FOXM1 phosphorylation levels were also severely reduced following CDK1/2/4 inhibition (Figure 4A, B, and S3). This is again consistent with FOXM1 being a direct target of CDK1 and CDK2 [26]. In contrast, WEE1 led to a slight increase in phospho-FOXM1 concurrent with an increase in CDK1 activity, as illustrated by loss of CDK1 phosphorylation (Figure 4A, B, and S3). Changes in p53 protein and phosphorylation levels were more pronounced following WEE1 inhibition than with CDK1/2/4 inhibition (Figure 4A and S3), although varying in relation to the MGT p53 status that we reported previously [4]. These findings were corroborated by a semi-quantitative reverse phase protein array (RPPA) proteomic profiling, a high-throughput antibody-based technique to analyse protein activities in signalling networks (Figure S2E). Interestingly, we found a specific downregulation of Cyclin D3 levels following CDK1/2/4 inhibition (Figure 4C), correlating with downregulation of PP1 (Figure 4C), a phosphatase that stabilizes Cyclin D3 by keeping it in a dephosphorylated state [27]. No significant changes were observed in Cyclin B1, Cyclin D1, and Cyclin E1 levels (Figure S2F). Collectively, these findings illustrate that CDK1/2/4 inhibition has drastic consequences on cell cycle regulators we tested, not significantly exacerbated by BCL-XL targeting.

**Figure 4.**
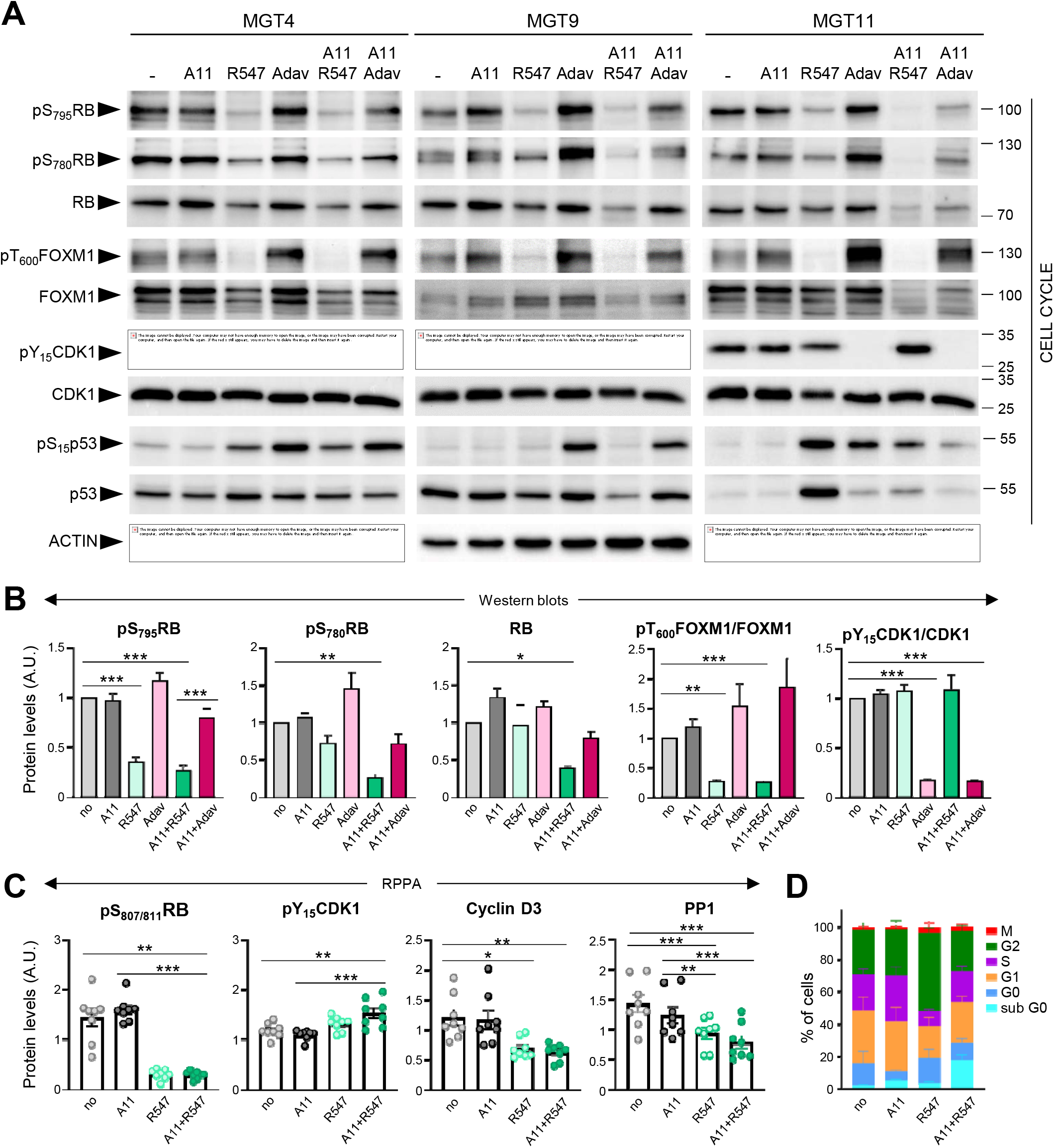
Inhibition of BCL-XL together with CDK1/2/4 perturbs cell cycle regulators. **(A)** Western blots performed on MGT4, MGT9, and MGT11 cells treated for 12hrs with A1155463 (A11: 1µM; BCL-XL inhibitor), R547 (3µM; CDK1/2/4 inhibitor), Adavosertib (Adav: 3µM; WEE1 inhibitor), alone or in combination. Adavosertib treatment was used for comparative studies. ACTIN, together with Ponceau (see the non-edited gels in Supplementary data), was used as loading control in all studies. Molecular weight markers are indicated on the right. **(B)** Protein levels estimated by densitometric analysis (using Image J) of western blots. Mean values obtained with the three MGT cell lines are shown as fold of control (not treated cells). Statistical analyses were done using ANOVA followed by Tukey test. **(C)** Changes in the expression/phosphorylation levels of the reported proteins in all four tumorigenic *MMTV-R26*^*Met*^ cell lines (MGT4, MGT9, MGT11, MGT13), either untreated or treated with the indicated drugs, based on the RPPA analysis (Table S5). **(D)** Histogram reporting the percentage of cells in each phase of the cell cycle when treated with the indicated drugs (for 12hrs), compared to non-treated cells (no). Distribution of cells in the different cell cycle phases was determined by flow cytometry using PI and Ki67 staining. Three independent experiments were done. Statistical analyses were performed by one-way (B-C) or two-way (D) ANOVA followed by Tukey test, and are reported in Table S7. * P<0.05; ** P<0.01; ***P<0.001. A.U.: arbitrary units.

We then analysed the cell cycle effects of R547 alone or in combination with BCL-XL targeting (with A1155463), by following the distribution of cells in the cycle phases through flow cytometry. In the presence of R547, we observed a decrease in the percentage of cells in S phase accompanied by an accumulation in G2 (Figure 4D and S2G). These results indicate a cell cycle blockage at G1-S and G2-M transitions, as previously reported [24]. Combined A1155463+R547 treatment led to an accumulation of cells in G1, consequently reducing the percentage of cells in G2 compared to R547 monotherapy, in agreement with previously reported actions of BCL-XL on cell cycle regulation [28, 29].

We next explored the consequences of BCL-XL+CDK1/2/4 inhibition on regulators of cell survival and DNA damage and compared them with those linked to the BCL-XL+WEE1 targeting that we previously reported [4]. We found a drastic decrease in anti-apoptotic XIAP and MCL1 protein levels in cells treated with BCL-XL+CDK1/2/4 targeting, accompanied by increased cleavage of Caspase3, Caspase 7, and PARP, as shown by western blot (Figure 5A, B, and S3) and RPPA (Figure 5C) analyses. Moreover, the combined treatments led to a high extent of DNA damage in cells, as revealed by the levels of γH2AX (a histone variant considered as a double-strand break sensor), by western blot (Figure 5A) and immuno-cytochemistry (Figure 5D). The high levels of DNA damage were accompanied by a downregulation of protein and/or phosphorylation levels of both ATM and ATR in cells treated with BCL-XL+CDK1/2/4 inhibitors, in contrast to unchanged, or a slight upregulation following BCL-XL+WEE1 blockage (Figure 5A and B), indicating a deficiency in the DNA damage detection mechanism induced by the A1155463+R547 drugs. Furthermore, this combined BCL-XL+CDK1/2/4 treatment did not affect the percentage of phospho-S_10_-Histone H3 (pH3)-positive cells, and did not trigger any mitotic catastrophe events in comparison to those observed in BCL-XL+WEE1 treated cells (Figure 5E and F). This agrees with the blockage of CDK1 by R547, therefore promoting cell cycle exit, in contrast to premature entry into mitosis observed with BCL-XL+WEE1 blockage (Figure 5F), as we showed in [4]. Together, these results illustrate that CDK1/2/4 inhibition together with BCL-XL targeting is as detrimental as BCL-XL+WEE1 blockage for TNBC cells, with similar perturbations of cell survival regulators, while having distinct effects on cell cycle and DNA repair components.

**Figure 5.**
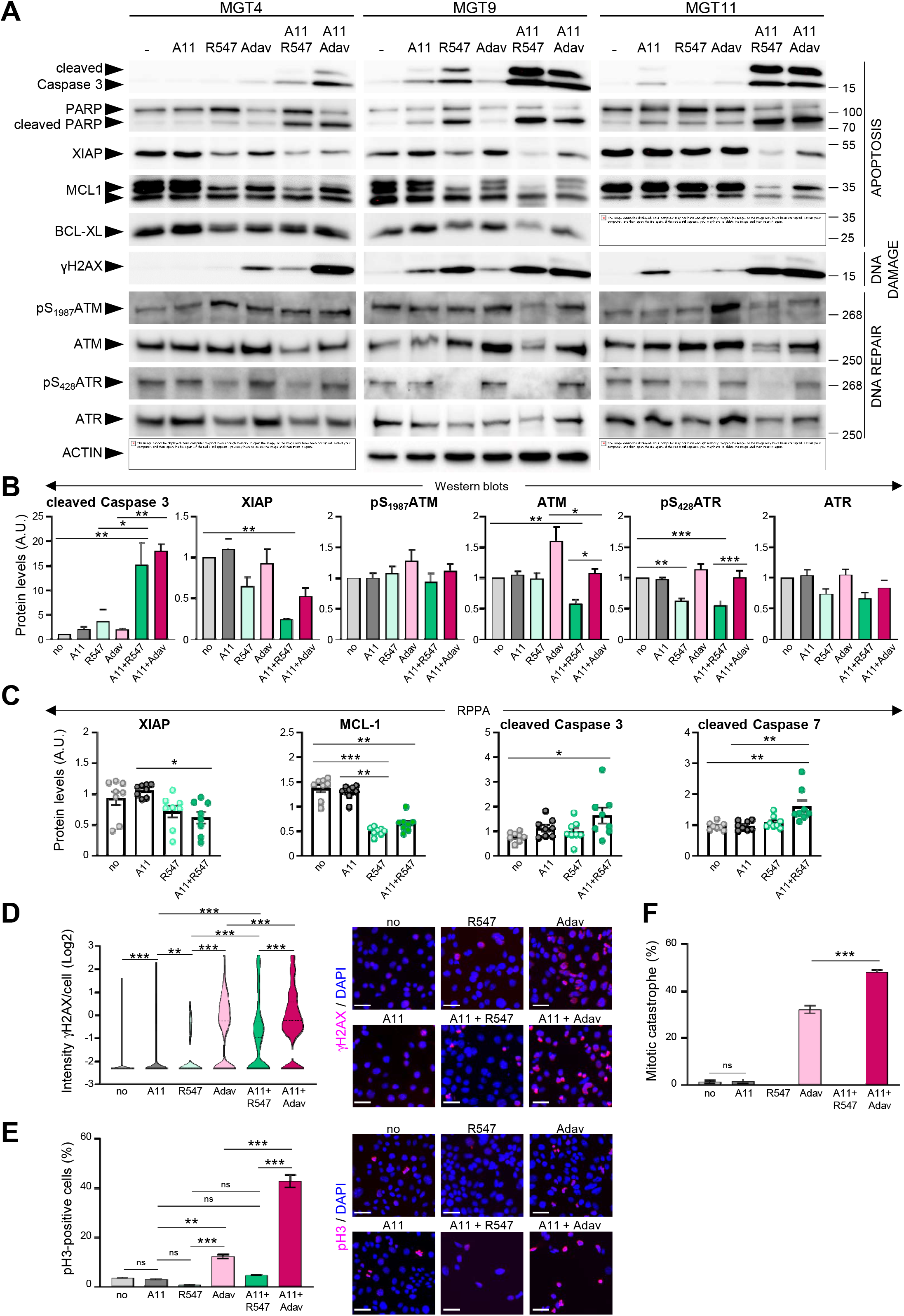
Co-targeting of BCL-XL and CDK1/2/4 induces DNA damage, interferes with survival signals, and triggers apoptosis. **(A)** *MMTV-R26*^*Met*^ cells were treated for 12hrs with either A1155463 (A11: 1µM; BCL-XL inhibitor), R547 (3µM; CDK1/2/4 inhibitor) or Adavosertib (Adav: 3 µM; WEE1 inhibitor), alone or in combination, then subjected to western blot analysis. Note that the ACTIN panels are the same as in Figure 4 as western blots were performed simultaneously. **(B)** Densitometric analysis (using Image J) of western blots depicting protein levels. Mean values obtained with the three MGT cell lines are shown as fold of control (untreated cells). **(C)** Graphs depicting changes in levels of anti-apoptotic proteins (XIAP and MCL-1) as well as apoptosis markers (cleaved-Caspase3 and cleaved-Caspase 7) in the four tumorigenic *MMTV-R26*^*Met*^ cell lines following treatment with the indicated drugs, based on the RPPA analysis (Table S6). **(D-F)** DNA damage and mitotic catastrophe analysis. MGT11 cells were treated or not with A1155463 (0.3µM), R547 (3µM), Adavosertib (3µM), or in combination. **(D)** Immunostaining with anti-γH2AX antibodies (to assess DNA damage) was performed after 12hrs of treatment. The violin plot depicts the number of cells according to their γH2AX staining intensity. Representative images of γH2AX immunostaining (red) are shown on the right. **(E)** Cells treated for 16hrs with the indicated drugs were immunostained with anti-pH3 antibodies. The graph reports the percentage of cells in mitosis (pH3-positive cells) versus the total number of cells. Representative images of pH3 immunostaining (red) are shown on the right. **(F)** Histogram reporting the number of mitotic catastrophe, (revealed by anti-pH3/α-Tubulin (microtubules) immunostaining) in treated cells. Mitotic catastrophe was analysed in cells in metaphase and anaphase among the pH3-positive cells. In contrast to WEE1 targeting (with Adavosertib), as previously reported [4], R547 does not induce mitotic catastrophe. In all experiments, DAPI was used to counterstain the nuclear DNA. Three independent experiments were performed. For multiple comparisons, statistical significance was assessed by One-way ANOVA followed by Tukey test. ns: not significant; *P<0.05; ** P<0.01; *** P<0.001. Scale bar: 50µm.

### Combined BCL-XL and CDK1/2/4 inhibition leads to downregulation of RTK and AKT signalling

To obtain further insights on signalling changes occurring in cells treated with BCL-XL+CDK1/2/4 inhibition and associated with their death, we bioinformatically explored RPPA outcomes on expression and/or phosphorylation levels of 426 proteins (Figure S2E and Table S5). First, we compared alterations occurring with single (BCL-XL or CDK1/2/4) versus combined BCL-XL+CDK1/2/4 targeting. We found that combined BCL-XL+CDK1/2/4 inhibition caused 177 alterations: 73 of which were already present following CDK1/2/4 inhibition (Figure 6A). Only 4 alterations were found following BCL-XL targeting (Figure 6A). Interestingly, protein-protein interaction network analysis, using the STRING tool, highlighted three main enriched pathways: RTK signalling, AKT signalling, and cell cycle/DNA damage (Figure 6B and S4). Furthermore, we found that CDK1/2/4 targeting leads to alteration of RTK signalling (Figure 6C: violet), AKT signalling (orange), cell cycle regulators (green), and components of DNA damage/repair and apoptosis (yellow). Interestingly, the combined treatment exacerbated perturbation of AKT signalling and DNA damage/apoptosis effectors (Figure 6C).

**Figure 6.**
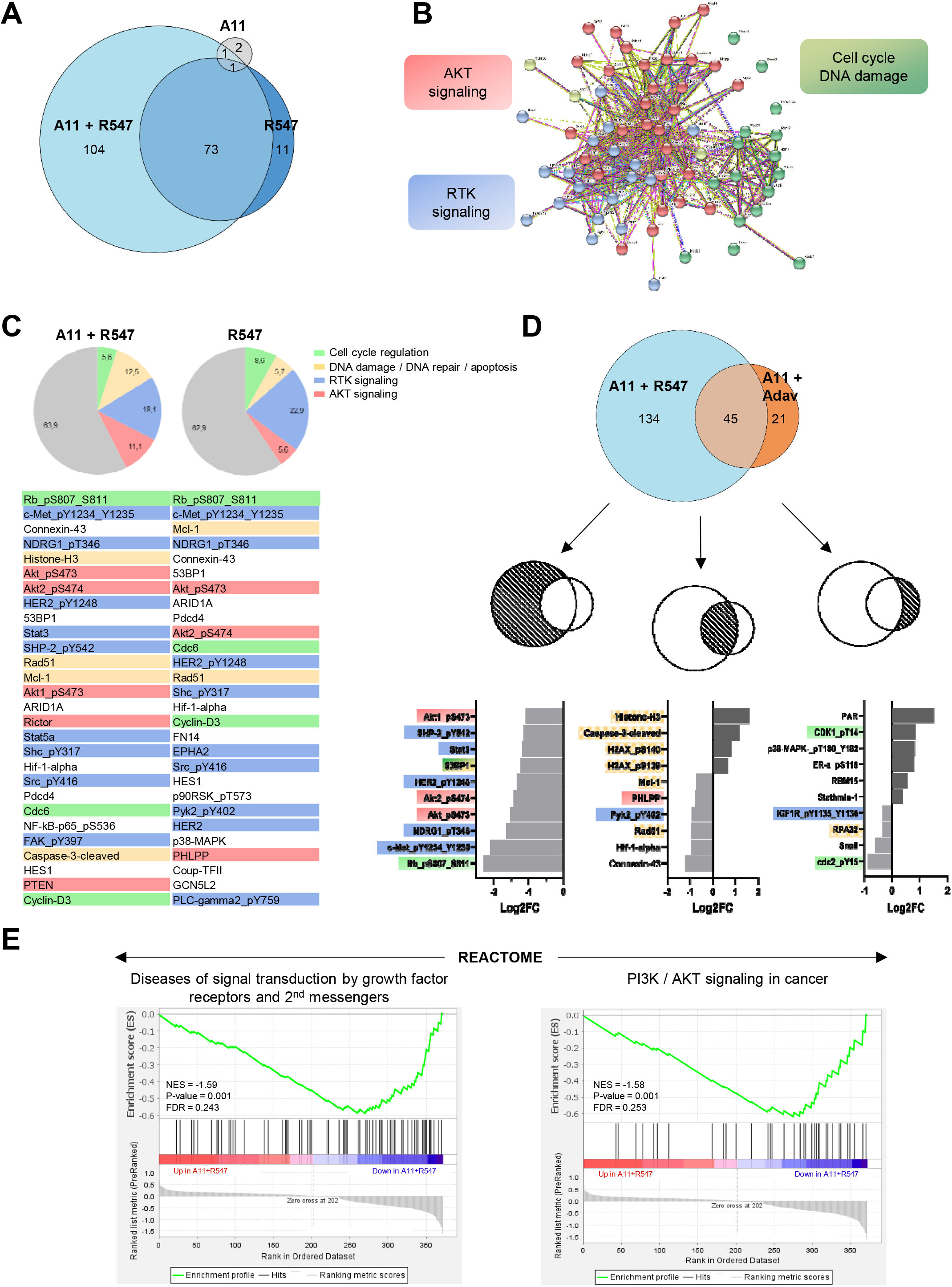
Proteomic analyses highlight downregulation of RTK and AKT signalling in *MMTV-R26*^*Met*^ MGT cells when subjected to combined inhibition of BCL-XL and CDK1/2/4. **(A)** Euler diagram showing the number of dysregulated signals in all *MMTV-R26*^*Met*^ cells treated with the indicated drugs. The diagram was obtained by the Eulerr package in R (https://cran.r-project.org/package=eulerr). The area of each circle is proportional to the number of dysregulated signals. A cut-off of p-value<0.05 was applied. **(B)** Projection of the dysregulated genes in cells treated with A1155463+R547 onto the STRING protein-protein interaction network highlights 3 main clusters using the kmeans clustering method. Clustering was based on protein interaction. The cut-off value was predefined as p-value < 0.05 and fold change <-0.5 or >0.5. See Figure S4 for high magnification. **(C)** Diagrams representing the proportional distribution of signals in all *MMTV-R26*^*Met*^ cells treated with R547 alone or in combination with A1155463. Percentages are indicated. The cut-off value was predefined as p-value < 0.05. The 30-top ranked dysregulated signals are listed. **(D)** Euler diagram depicting the number of dysregulated signals in cells treated with either A11+R547 or A11+Adavosertib. A cut-off of p-value <0.05 was applied. The 10-top ranked dysregulated signals among either the specific (A11+R547: 134; A11+Adav: 21) or the common (A11+R547/A11+Adav: 45) signals are shown. For panels **C** and **D**: among signals, we highlighted RTK signalling (blue), AKT signalling (red), proteins involved in DNA damage/repair/apoptosis (yellow), and those in cell cycle regulation (green). **(E)** GSEA enrichment plots performed, using the REACTOME database, on all signals from A11+R547 versus A11+Adav data. Note that gene-sets related to RTK signalling and second messengers, as well as PI3K/AKT signalling, are in the top 10 down-regulated Reactome pathways in BCL-XL+CDK1/2/4-treated *MMTV-R26*^*Met*^ cells. The barcode plot indicates the position of a member of a gene set in the ranked list of proteins. Red and blue colours represent proteins up-or downregulated in A11+R547 versus A11+Adav, respectively. NES: normalized enrichment score; FDR: false discovery rate.

We then compared signalling changes linked to BCL-XL+CDK1/2/4 inhibition with those occurring in cells following BCL-XL+WEE1 blockage. The top ranked signalling alterations observed specifically upon BCL-XL+CDK1/2/4 inhibition were mainly signals involved in RTK and/or AKT pathways (e.g. downregulation of: phospho-MET, phospho-HER2, phospho-AKT, phospho-NDRG1, phospho-SHP2, STAT3), while BCL-XL+WEE1 targeting mainly impacted regulators of cell cycle and DNA repair (Figure 6D). DNA damage/repair/apoptosis deregulation was seen with both combinatorial settings (Figure 6D).

To further examine the mechanisms underlying the effects of these drug combinations, we performed a GSEA using all signals from the RPPA outcomes from cells treated with BCL-XL+CDK1/2/4 versus BCL-XL+WEE1 inhibitors. Using the Reactome database, we found that gene-sets related to RTK signalling and second messengers, as well as PI3K/AKT signalling, were among the most significant deregulated signals in *MMTV-R26*^*Met*^ MGT cells treated with BCL-XL+CDK1/2/4 inhibitors (Figure 6E). This approach delineated a possible mechanism underlying combined BCL-XL plus CDK1/2/4 targeting, showing a pronounced alteration of RTK and AKT signalling.

Interestingly, it has recently been shown that CDK1, together with Aurora kinase, ensures RTK storage by suppressing endosomal degradation and recycling pathways [30]. We therefore further investigated the striking correlation between BCL-XL+CDK1/2/4 inhibition and the downregulation of RTK and ATK signalling. Western blot results revealed a consistent downregulation of expression and/or phosphorylation levels of several proteins, including MET, GAB1, ERKs, and AKT in cells with BCL-XL+CDK1/2/4 inhibition, although with slightly different intensities among the *MMTV-R26*^*Met*^ MGT cells (Figure 7A and S3). These downregulations were further confirmed by performing kinetic studies (Figure 7C). Additionally, RPPA analysis showed downregulation of protein expression and/or phosphorylation levels of AKT, RICTOR, and FOXO3a following BCL-XL+CDK1/2/4 inhibition (Figure 7B).

**Figure 7.**
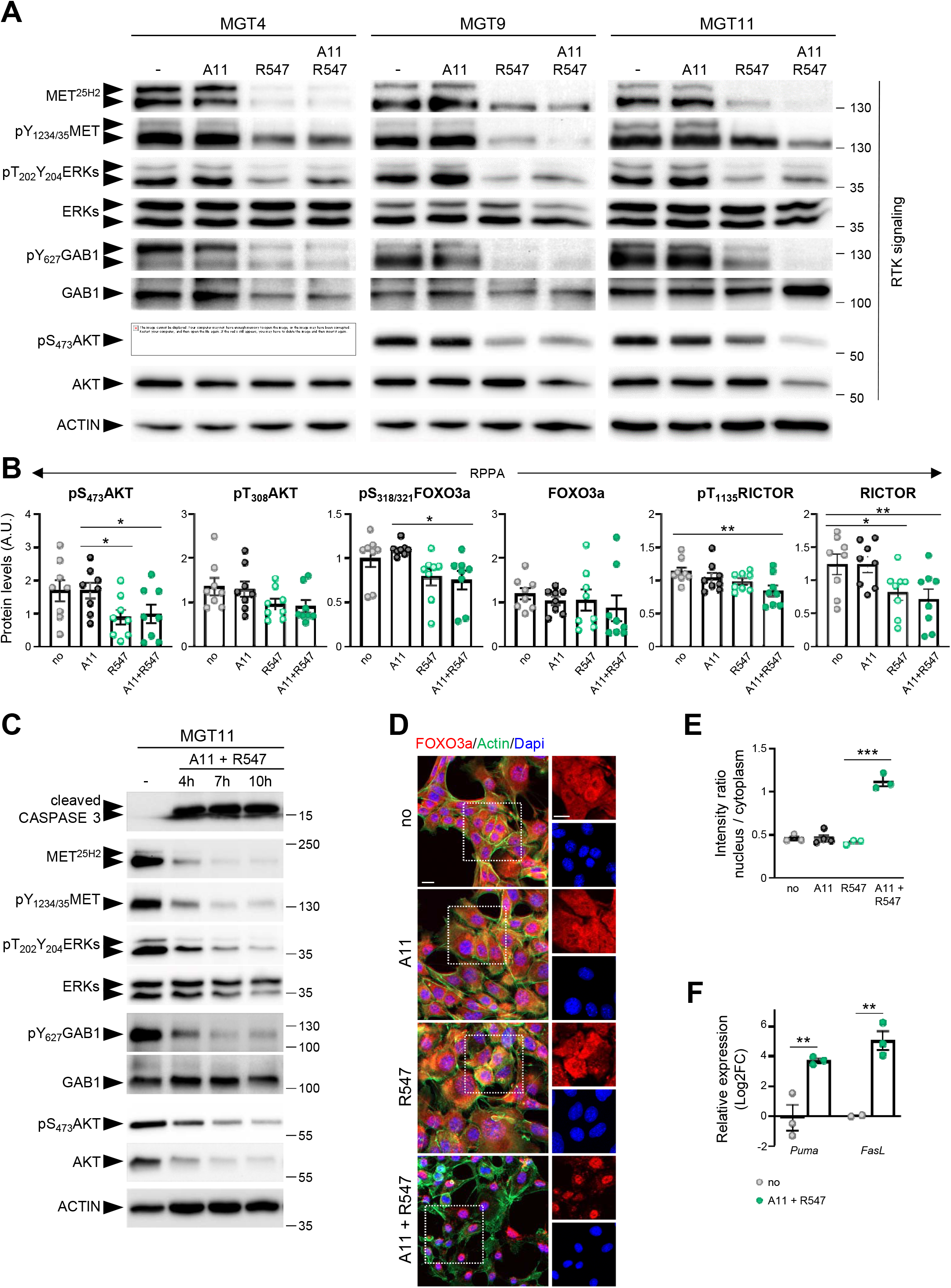
Combined BCL-XL and CDK1/2/4 inhibition leads to RTK/AKT signalling downregulation and FOXO3a nuclear retention with activation of apoptosis associated genes. **(A)** Western blots performed on *MMTV-R26*^*Met*^ MGT cells after a 12hr-treatment with the indicated drugs. **(B)** Histograms reporting downregulation of expression and/or phosphorylation levels of AKT, FOXO3a, and RICTOR following BCL-XL and CDK1/2/4 inhibition, based on the RPPA outcomes. **(C)** Kinetic analysis of changes in expression and/or phosphorylation levels of the indicated RTK signalling components after combined targeting of BCL-XL and CDK1/2/4. **(D)** MGT11 cells treated with the indicated drugs were immunostained with anti-FOXO3a (red) and phalloidin (to detect F-actin, green). Note the nuclear retention of the FOXO3a protein when cells were treated with BCL-XL+CDK1/2/4 inhibitors (A11+R547). DAPI (blue) was used to counterstain the nuclear DNA. The right panels depict split red and blue channels of the indicated areas. **(E)** Graph depicting the intensity ratio of nuclear versus cytoplasmic FOXO3a in cells when exposed to the indicated drugs. **(F)** RT-qPCR analysis of FOXO3a target genes in cells treated with A11+R547. Statistical significance was assessed by One-way ANOVA followed by Tukey test. *P<0.05; ** P<0.01; *** P<0.001. Scale bar: 20 µm.

The correlation between reduced phosphorylation of AKT and FOXO3a was particularly interesting, considering that AKT phosphorylates FOXO3a leading to its translocation from nucleus to cytoplasm, thus preventing its tumour suppressor function [31]. We therefore explored this aspect by performing immuno-cytochemistry to follow FOXO3a localisation in untreated and treated cells. We found a striking FOXO3a nuclear localization in cells treated with BCL-XL+CDK1/2/4 inhibition compared with cells either untreated or treated with single drugs (Figure 7D and E). This FOXO3a nuclear retention was accompanied by a transcriptional expression of *PUMA* and *FasL*, two FOXO3a target genes involved in apoptosis (Figure 7F). Together, these findings show that combined BCL-XL and CDK1/2/4 targeting, while perturbing cell cycle regulators, leads to a combinatorial depletion of survival and RTK/AKT signals, restoring FOXO3a tumour suppression actions.

### Discussion

In this study, we provide evidence that combinatorial targeting of BCL-XL with CDK1/2/4 could be an efficient therapeutic approach for TNBC. This drug combination was potent across different TNBC subtypes, as demonstrated by its high efficacy in different human TNBC cell lines and in the heterogenous cell lines generated from distinct *MMTV-R26*^*Met*^ tumours. Such combination appears to be less effective on non-TNBC cells, based on the cell lines used in this study.

The alteration of cell cycle regulators in several types of cancer [10], including breast cancer, and the possibility to modulate their function, has fostered the interest to design treatment options targeting them, with some of them already being exploited in clinical trials [10]. The relevance of alterations of cell cycle regulators in breast cancer is strengthened by several -omics analyses and bioinformatic processing. For example, a recent study revisited breast cancer microarray datasets uncovering that differentially expressed genes are mainly enriched in cell cycle regulators [23]. Specifically, CDK4/6 oncogenic activation has been reported in luminal breast cancer, constituting one of the main tumorigenic drivers. In contrast, dysregulations associated with TNBC formation include *c-Myc* activation, *p53* mutations, *PTEN*-loss, and *CDKs/Cyclins* overactivation/overexpression [10, 32]. In this study, we have shown that TNBC patients with high levels of *CDK1, CDK6*, and *WEE1*, are characterised by a worse prognosis compared to those with low expression levels. Interestingly, we have also shown that poor TNBC prognosis is further exacerbated by high expression levels of BCL-XL.

The identification of vulnerabilities of cancer cells is particularly challenging for those types of cancer, like TNBC, highly heterogeneous and lacking major drivers to which cells are addicted to [33]. To uncover TNBC signalling vulnerabilities, relevant model systems mimicking disease characteristics are essential. We have recently reported the uniqueness of the *MMTV-R26*^*Met*^ model, which recapitulates several features of TNBC including primary therapy resistance and marker intertumoral heterogeneity. This model has previously allowed us to uncover the potency of WEE1 targeting while lowering survival inputs through BCL-XL inhibition [4]. The present study further confirms the vulnerability of TNBC cells to targeting cell cycle regulators in the context of BCL-XL blockage.

In this regard, an intriguing aspect emerging from the present studies is that in the presence of BCL-XL, the specific cell cycle regulator targeted determines the effect on TNBC cell viability. Specifically, TNBC cells are sensitive to combined inhibition of CDK1/2/4 by R547, but not of FOXM1, CDK4/6, Aurora A and B. FOXM1 is a transcription factor overexpressed in most solid tumours, such as breast, liver, prostate, colon, and pancreas [34]. This member of the Forkhead family regulates the expression of a large set of G2/M specific genes, and aberrant FOXM1 upregulation has been shown to be a key driver in cancer progression [26]. Our RNA-seq analysis showed an up-regulation of *CDK1, CDK2, CDK4, FoxM1, Aurora A/B*, but not of *CDK3, CDK5, CDK6*, genes in *MMTV-R26*^*Met*^ tumours versus normal mammary glands, as reported in TNBC patients [35, 36]. Nevertheless, their inhibition together with BCL-XL targeting had no effect on the viability of the four *MMTV-R26*^*Met*^ TNBC cell lines we examined. This apparent contradiction to previous studies [37] is likely explained by the fact that the effectiveness of FOXM1 or CDK4/6 targeting is conditioned by the set of molecular alterations present in cancer cells. Indeed, it has been shown that Palbociclib (a potent CDK4/6 inhibitor currently used in the clinic) in RB-proficient TNBC cells effectively potentiates subsequent treatments with chemotherapeutic agents like paclitaxel or cisplatin, although eliciting antagonist effects when simultaneously used [14, 38]. Furthermore, it has been shown that FoxM1 and Aurora A targeting overcomes TNBC paclitaxel resistance [39]. Our RPPA and biochemical studies showed that RB is hyper-phosphorylated in *MMTV-R26*^*Met*^ TNBC cells [4], although levels may not be sufficient to confer the sensitivity of cells to Palbociclib treatment. Alternatively, it is possible that the effects of Palbociclib in TNBC cells is conditioned by the drug used in combination. Here, we show that Palbociclib does not synergize with BCL-XL inhibition. The R547 drug we used in these studies is a potent ATP-competitive inhibitor of CDK1/2/4 [24]. The strong combinatorial effects of BCL-XL and CDK1/2/4 blockage we highlighted here point to this combination as a possible effective option to surpass, at least in part, the molecular heterogeneity of TNBC (e.g. the RB levels and its phosphorylation status). Future studies are needed to clarify whether BCL-XL+CDK1/2/4 inhibition would also minimize heterogeneous mechanisms of resistance otherwise occurring by targeting CDK4/6 with chemotherapeutics previously reported [40]. It would also be important to assess the net contribution of each CDK among the CDK1/2/4 inhibited by R547 while blocking BCL-XL. However, this would require rather laborious experimental setting using single and combined conditional shRNA sequences targeting each individual CDK, as agents blocking specifically individual CDKs are not available. In relation to ongoing clinical trials using drugs blocking cell cycle regulators as targeted anticancer therapies, our studies underline the importance of carefully evaluating the best signals to target to achieve optimal response while minimizing side effects in patients.

Beside regulating cell cycle, new functions of CDK1/2/4 have recently emerged. It has been shown that CDK1/2 targets are hyperphosphorylated in basal-like breast cancer, generating a genome integrity vulnerability [41]. Additionally, CDK1 phosphorylates several proteins involved in epigenetic regulation, such as the H3K79 methyltransferase Dot1l, responsible for placing activating marks on gene bodies [42]. Furthermore, it has been shown that CDK1 promotes storage of RTKs by suppressing endosomal degradation and recycling pathways [30]. The latter function may be particularly relevant for the effects we observed in TNBC cells following BCL-XL plus CDK1/2/4 targeting as most of the altered signals belong to RTK and AKT signalling, beside cell cycle regulators. In view of the findings reported in [30], it is tempting to speculate that inhibition of CDK1 perturbs RTK trafficking and storage, reducing the recycling pool due to increased processing and degradation. Lowering RTK and AKT signalling is likely detrimental for the cells, particularly in a context of reduced stress support pathway associated with BCL-XL inhibition. This also leads to nuclear retention, and therefore transcriptional activity, of FOXO3A, a tumour suppressor regulating expression of a plethora of genes involved in multiple biological processes. Thus, the effectiveness on TNBC cells of a combinatorial targeting of BCL-XL plus CDK1/2/4 inhibition likely resides on the acquisition of DNA damage (illustrated by an accumulation of γH2AX) associated with cell cycle progression perturbation, an unbalance of survival/apoptotic signals (reduced XIAP-MCL1 and increased FASL-PUMA levels), with a concomitant depletion of RTK and AKT inputs. Future studies will define whether there is also a perturbation of epigenetic marks on gene bodies, as a consequence of CDK1/2 targeting [42]. This would be particularly relevant also in view of how high levels of oncogene sets are ensured by gene body hypermethylation, as we reported in liver cancer [17].

## Acknowledgments

We thank: all members of our labs for helpful discussions and comments; R. Dono, F. Helmbacher, V. Géli for valuable feedback on the study; C. Sequera for help in processing data related to the overall survival of TNBC patients according to levels of cell cycle regulators; A.L. Bailly for performing cell cycle analysis reported in Figure 4D and Figure S2G; S. Richelme for assistance as lab manager; F. Daian for computational work to calculate the Synergy maps and Bliss scores; P. Perrin for providing us with mouse embryonic fibroblasts; the animal house platform for excellent help with mouse husbandry.

## Author contributions

O.C.: performed most of the biochemical experiments, cell viability assays, and computational work to analyse RNA-seq and RPPA outcomes; largely contributed to data analysis and interpretation.

F.A.: performed the majority of the biochemical studies and cell viability assays, prepared samples for RPPA; data analysis and interpretation.

Y.V.: performed computational work to analyse RNA-seq and RPPA outcomes, provided inputs on studies and on the manuscript.

G.B.M.: contributed to RPPA studies; provided input on the manuscript.

B.H.: processed RNA-seq outcomes and generated raw data; provided input on the manuscript.

J.P.B.: supervised the FACS sorting analysis; provided input on the manuscript.

S.L.: supervised RPPA analyses performed by Y.V.; provided input on studies and on the manuscript.

F.L.: designed and supervised studies, contributed to experimental work, data analysis, and interpretation; contributed to write the manuscript.

F.M.: designed and supervised the study, contributed to experimental work, analysed, and interpreted data, ensured financial support, and wrote the manuscript.

