## Supplementary Information, Figures and Tables for "BCL-XL blockage in TNBC models confers vulnerability to inhibition of specific cell cycle regulators"

[1] Aix Marseille Univ, CNRS, Developmental Biology Institute of Marseille (IBDM), Turing Center for Living Systems, Parc Scientifique de Luminy, Marseille (France). [2] Weizmann Institute of Science, Department of Molecular Cell Biology, Rehovot (Israel). [3] Knight Cancer Institute, Portland, OR 97201, USA. [4] Aix Marseille Univ, Centre de Recherche en Cancérologie de Marseille (CRCM), Equipe labellisée Ligue 'Cell polarity, cell signaling and cancer', Inserm, CNRS, Institut Paoli-Calmettes, Marseille (France). [5] Institut Universitaire de France (IUF).

[6] shared first authors

**Keywords:** cancer mouse model, triple-negative breast cancer, signalling reprogramming, drug resistance, cell cycle regulators, BCL-XL, CDKs, MET

**Supplementary Figure Legends**

**Figure S1. Human breast tumours are enriched in cell cycle regulators.** High magnification of the STRING network reported in Figure 1A.

**Figure S2. *MMTV-R26$^{Met}$* TNBC cells are vulnerable to combined targeting of BCL-XL and CDK1/2/4.** **(A)** RNA-seq data from *MMTV-R26$^{Met}$* tumours (n=4) versus control mammary gland tissues (n=3). Enrichment pathway analysis, using WikiPathways 2015, ordered according to the combined score. The 10-top ranked enriched pathways are shown and highlight enrichment in cell cycle/DNA replication (red arrowhead) and DNA damage (green arrowhead) regulatory pathways. The cut-off applied to identify deregulated genes was p-value <0.05. **(B)** GSEA enrichment performed, using the WikiPathways database with the DNA damage response geneset, on *MMTV-R26$^{Met}$* tumours versus controls. The barcode plot indicates the position of a member of this geneset in the ranked list. Red and blue colours represent up- or downregulated genes in the *MMTV-R26$^{Met}$* tumours versus the controls, respectively. NES: normalized enrichment score; FDR: false discovery rate. **(C)** Cell viability outcomes of combined drug effects (see detailed matrix, Figure 3G) were used by the Compusyn software to simulate combination index (Y-axis) for each affected fraction (X-axis, from 0 to 1). Black dots correspond to tested concentrations. Based on the combination index scores, co-targeting of BCL-XL (with A1155463) and CDK1/2/4 (with R547) resulted in synergistic interactions for MGT4, MGT9 and MGT11 cells. **(D)** Cell viability assay performed on the non-tumorigenic *MMTV-R26$^{Met}$* MGT2 cell line as well as on mitomycin C-treated mouse embryonic fibroblasts (MEFs). **(E)** Heatmap reporting expression or phosphorylation levels of proteins in the four tumorigenic *MMTV-R26$^{Met}$* cell lines as determined by RPPA. **(F)** Comparable protein levels of Cyclin B1, D1, and E1 in the four tumorigenic *MMTV-R26$^{Met}$* cell lines untreated or treated with the indicated drugs (based on the RPPA analysis; Table S5). **(G)** Representative graphs showing cell cycle analysis as measured by flow cytometry using PI and Ki67 staining of MGT11 cells untreated (no) or treated with the indicated drugs: A1155463 (A11, 1µM); R547 (3µM).

**Figure S3. Signalling perturbation in *MMTV-R26$^{Met}$* TNBC cell lines by blocking BCL-XL and CDK1/2/4.** MGT13 cells were untreated (-) or treated for 12hrs with either A1155463 (A11: 1µM), R547 (3µM) or Adavosertib (Adav: 3 µM), alone or in combination, then subjected to western blot analysis. Molecular weight markers are indicated on the right. ACTIN and Ponceau (see also not edited gels) were used as loading controls.

**Figure S4. Combined targeting of BCL-XL and CDK1/2/4 in *MMTV-R26$^{Met}$* MGT cells leads to downregulation of RTK and AKT signalling as well as signals involved in cell cycle regulation and DNA damage/repair.** High magnification of the STRING network reported in Figure 6B.

**Supplementary Table Legends**

**Table S1 - Enrichr analysis on Breast cancer patients (GSE42568, GSE45827 and GSE54002) (Figure 1B)**.

**Table S2 - Microarray analysis of GSE31519 (TNBC patients) (Figure 2A-C)**.

**Table S3 - Enrichr analysis on *MMTV-R26$^{Met}$* tumours (Figure 3B and C)**.

**Table S4 – RNA-seq data cell cycle regulators in *MMTVR26$^{Met}$* tumours versus controls (Figure 3B and C)**.

**Table S5 - All deregulated signals in treated MGT cells**.

**Table S6 - Antibodies used for RPPA analysis of MMTV-R26Met treated cells**.

**Table S7 – Statistics of cell cycle distribution of MGT11 treated cells (Figure 4D)**.

**Table S8 - List of antibodies used in this study**.

**Table S9 - List of drugs used in this study.**

**Table S10 - List of oligonucleotides (Figure 7F).**

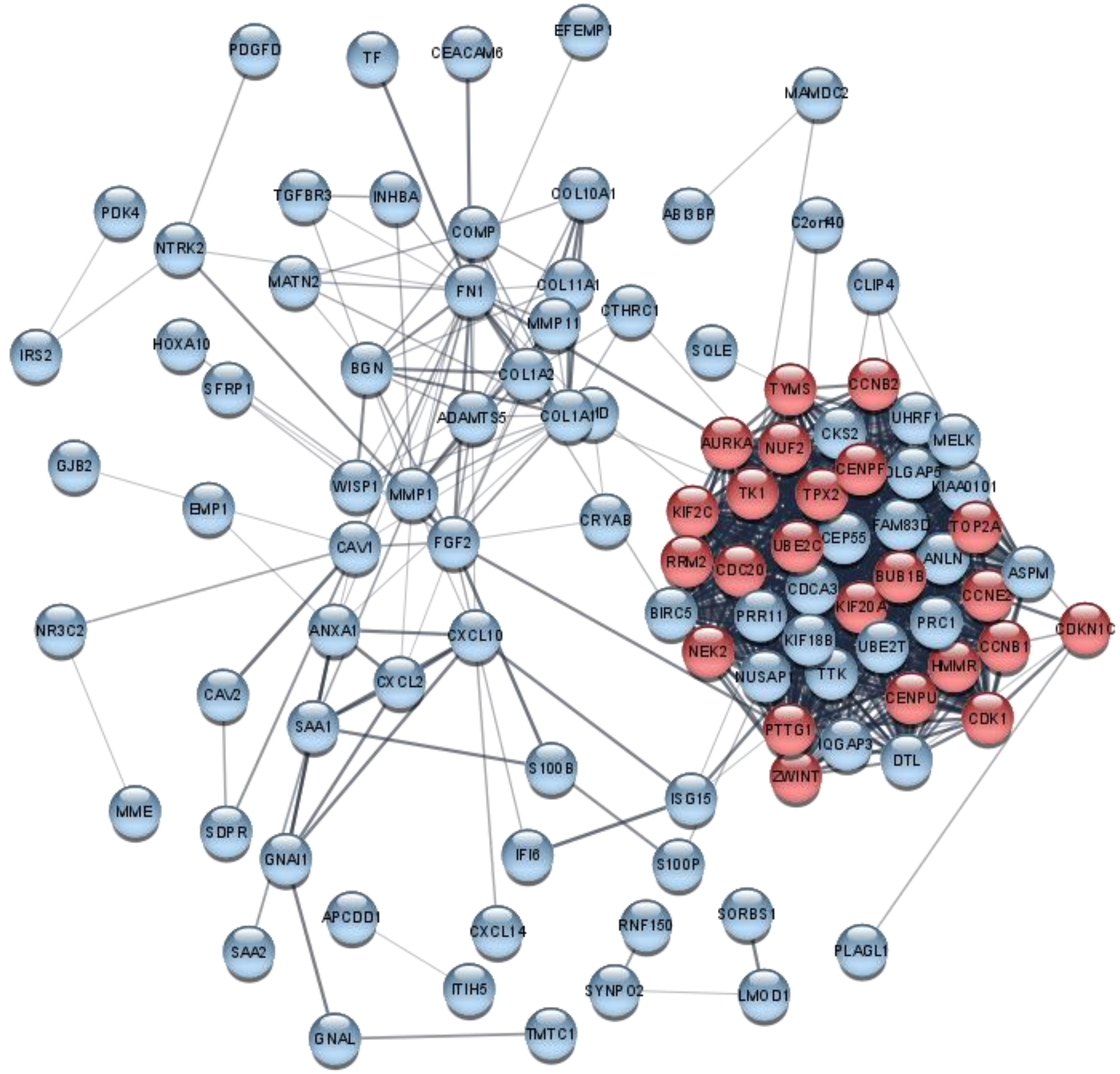
PDGFD
TF
CEACAM6
EFEMP1
MAMDC2
TGFBR3
INHBA
COL10A1
ABI3BP
C2orf40
PDK4
NTRK2
COMP
MATN2
FN1
COL11A1
CLIP4
IRS2
HOXA10
MMP11
CTHRC1
SQLE
BGN
COL1A2
TYMS
CCNB2
SFRP1
ADAMTS5
COL1A1
UHRF1
CKS2
MELK
AURKA
NUF2
GJB2
CENPF
KIAA0101
WISP1
MMP1
TK1
TPX2
EMP1
KIF2C
CRYAB
FAM83D
TOP2A
CAV1
FGF2
UBE2C
CEP55
RRM2
CDC20
ANLN
ASPM
BUB1B
CDCA3
CCNE2
BIRC5
KIF20A
NR3C2
ANXA1
CXCL10
PRR11
PRC1
CDKN1C
KIF18B
CCNB1
NEK2
UBE2T
CXCL2
NUSAP1
TTK
HMMR
CAV2
CENPU
SAA1
PTTG1
CDK1
IQGAP3
DTL
S100B
ZWINT
ISG15
MME
SDPR
IFI6
S100P
GNAI1
APCDD1
SORBS1
RNF150
CXCL14
PLAGL1
SAA2
ITIH5
SYNPO2
LMOD1
GNAL
TMTC1

**Figure S1**

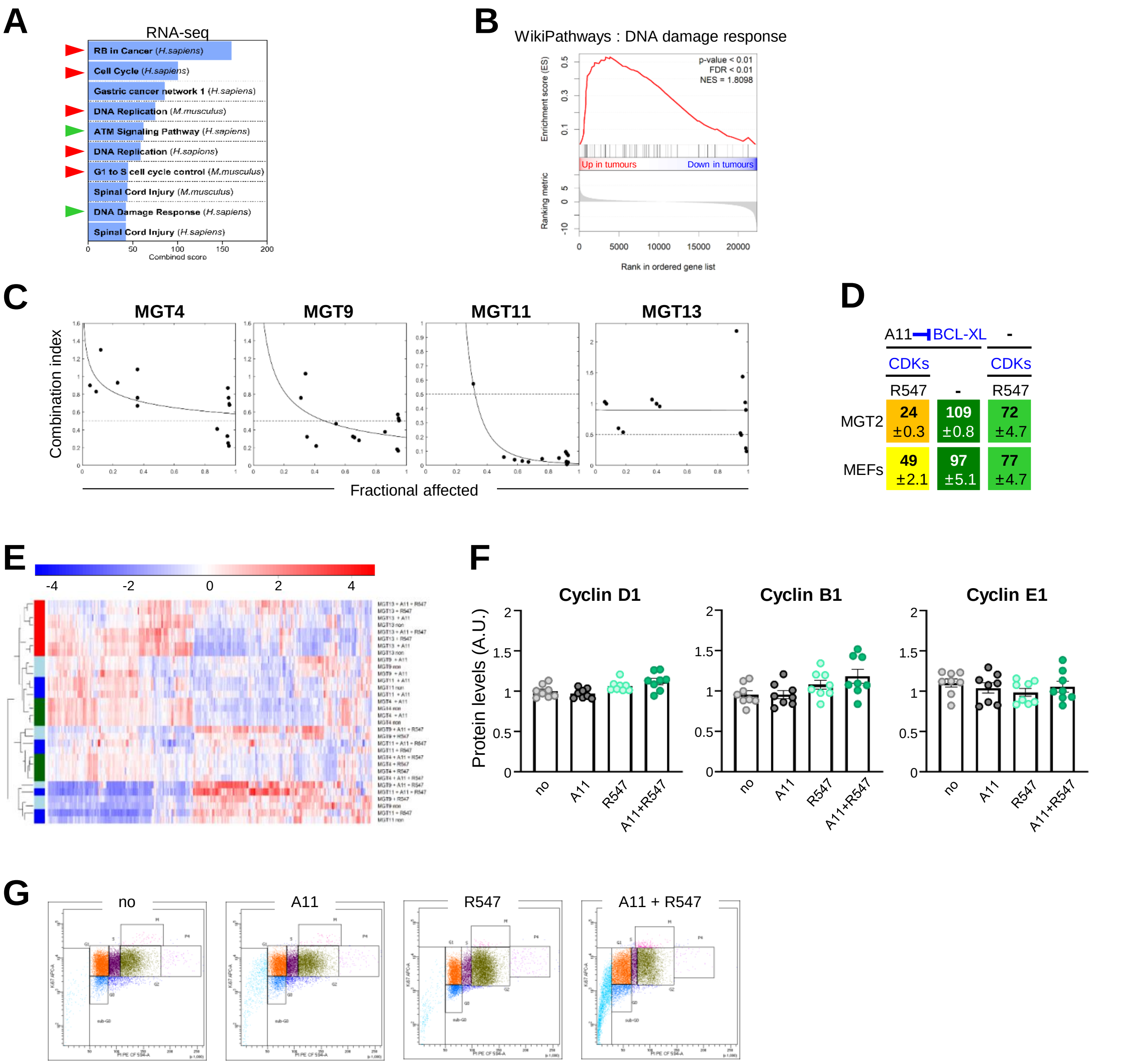
A
RNA-seq
RB in Cancer (H.sapiens)
Cell Cycle (H.sapiens)
Gastric cancer network 1 (H.sapiens)
DNA Replication (M.musculus)
ATM Signaling Pathway (H.sapiens)
DNA Replication (H.sapiens)
G1 to S cell cycle control (M.musculus)
Spinal Cord Injury (M.musculus)
DNA Damage Response (H.sapiens)
Spinal Cord Injury (H.sapiens)
Combined score
B
WikiPathways : DNA damage response
p-value < 0.01
FDR < 0.01
NES = 1.8098
Enrichment score (ES)
Up in tumours
Down in tumours
Ranking metric
Rank in ordered gene list
C
MGT4
MGT9
MGT11
MGT13
Combination index
Fractional affected
D
A11 BCL-XL
-
CDKs
CDKs
R547
-
R547
MGT2
24 ±0.3
109 ±0.8
72 ±4.7
MEFs
49 ±2.1
97 ±5.1
77 ±4.7
E
F
Cyclin D1
Cyclin B1
Cyclin E1
Protein levels (A.U.)
no
A11
R547
A11+R547
G
no
A11
R547
A11 + R547

Figure S2

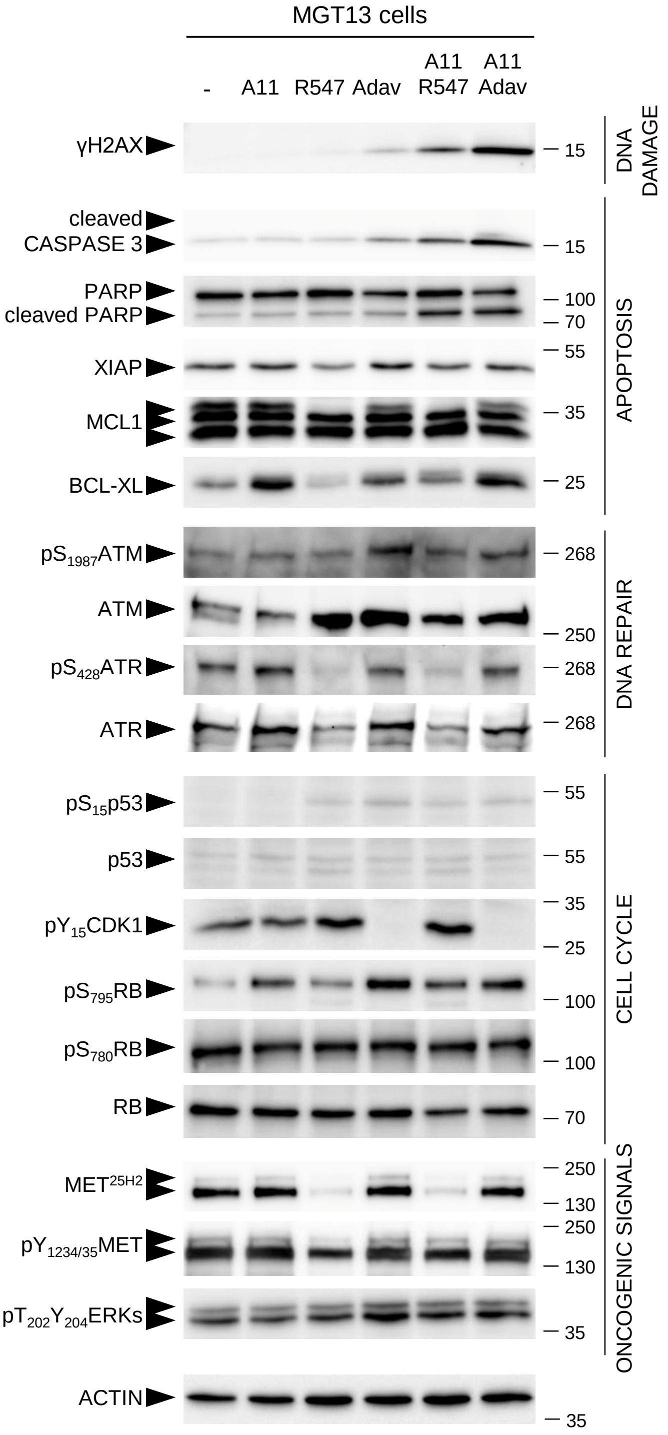

MGT13 cells
-
A11
R547
Adav
A11 R547
A11 Adav
γH2AX
cleaved CASPASE 3
PARP
cleaved PARP
XIAP
MCL1
BCL-XL
pS1987ATM
ATM
pS428ATR
ATR
pS15p53
p53
pY15CDK1
pS795RB
pS780RB
RB
MET25H2
pY1234/35MET
pT202Y204ERKs
ACTIN
15
15
100
70
55
35
25
268
250
268
268
55
55
35
25
100
100
70
250
130
250
130
35
35
DNA DAMAGE
APOPTOSIS
DNA REPAIR
CELL CYCLE
ONCOGENIC SIGNALS

**Figure S3**

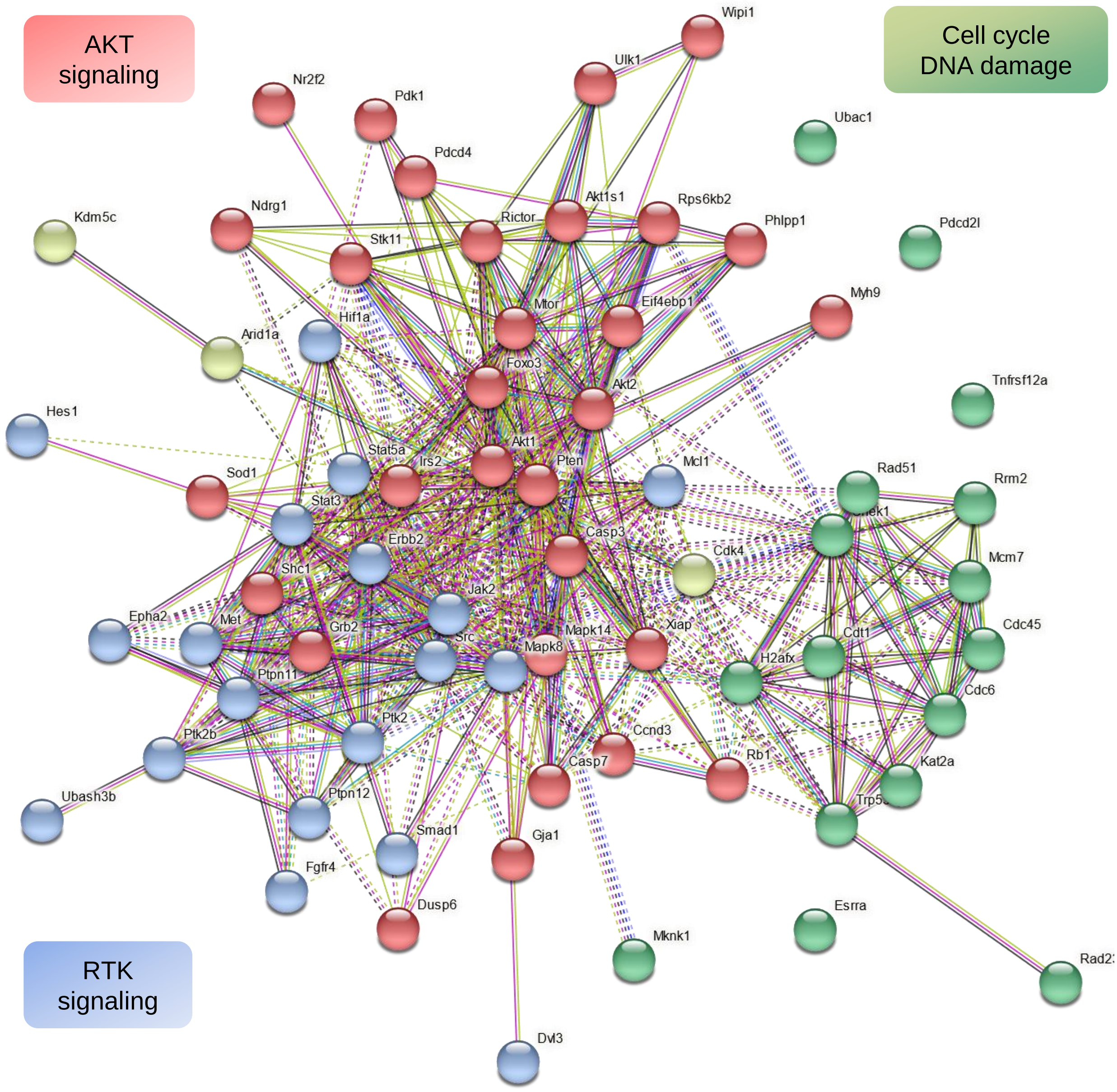
AKT
signaling
Cell cycle
DNA damage
RTK
signaling
Wipi1
Ulk1
Nr2f2
Pdk1
Ubac1
Pdcd4
Akt1s1
Rps6kb2
Phlpp1
Pdcd2l
Kdm5c
Ndrg1
Rictor
Stk11
Myh9
Mtor
Eif4ebp1
Hif1a
Arid1a
Foxo3
Akt2
Tnfrsf12a
Hes1
Akt1
Stat5a
Irs2
Pten
Mcl1
Rad51
Rrm2
Sod1
Stat3
Erbb2
Casp3
Cdk4
Mcm7
Shc1
Jak2
Epha2
Met
Grb2
Src
Mapk14
Xiap
Cdt1
Cdc45
Mapk8
H2afx
Ptpn11
Cdc6
Ptk2
Ptk2b
Ccnd3
Casp7
Rb1
Kat2a
Ubash3b
Ptpn12
Smad1
Gja1
Fgfr4
Dusp6
Esrra
Mknk1
Dvl3

**Figure S4**

Table S1. KEGG_2019_Human

| Term | Overlap | P-value | Adjusted P-va | Old P-value | Old Adjusted | Odds Ratio | Combined Sco | Genes |
|---|---|---|---|---|---|---|---|---|
| Cell cycle | 9/124 | 7,34E-08 | 1,01E-05 | 0 | 0 | 13,5665904 | 222,867345 | CDC20;CDKN1C;CCNB2;CCNB1;PTTG1;CCNE2;CDK1;BUB1B;TTK |
| p53 signaling | 5/72 | 8,16E-05 | 0,00372669 | 0 | 0 | 12,5284594 | 117,937959 | CCNB2;CCNB1;RRM2;CCNE2;CDK1 |
| Oocyte meios | 7/125 | 1,23E-05 | 8,40E-04 | 0 | 0 | 10,1046902 | 114,274837 | CDC20;CCNB2;CCNB1;PTTG1;CCNE2;CDK1;AURKA |
| ECM-receptor | 5/82 | 1,52E-04 | 0,00519038 | 0 | 0 | 10,8958838 | 95,8253101 | COMP;COL1A1;COL1A2;FN1;HMMR |
| Protein digest | 5/90 | 2,35E-04 | 0,0053598 | 0 | 0 | 9,8664008 | 82,4539977 | COL1A1;COL1A2;MME;COL11A1;COL10A1 |
| Progesterone- | 5/99 | 3,66E-04 | 0,00715502 | 0 | 0 | 8,91768842 | 70,5746889 | CCNB2;CCNB1;CDK1;GNAI1;AURKA |
| Focal adhesion | 7/199 | 2,33E-04 | 0,0053598 | 0 | 0 | 6,18691631 | 51,7468483 | COMP;COL1A1;COL1A2;CAV2;PDGFD;CAV1;FN1 |
| Pyrimidine me | 3/57 | 0,00521504 | 0,06495098 | 0 | 0 | 9,17731481 | 48,2378767 | RRM2;TK1;TYMS |
| Amoebiasis | 4/96 | 0,00292675 | 0,04455164 | 0 | 0 | 7,22871757 | 42,1713458 | COL1A1;COL1A2;GNAL;FN1 |
| Drug metabol | 4/108 | 0,00446298 | 0,06114286 | 0 | 0 | 6,3907563 | 34,5863775 | RRM2;MAOA;FMO2;TK1 |
| Bacterial inva | 3/74 | 0,01071473 | 0,11213358 | 0 | 0 | 6,97394366 | 31,634756 | CAV2;CAV1;FN1 |
| PI3K-Akt signa | 8/354 | 0,00157007 | 0,02688739 | 0 | 0 | 3,92681578 | 25,3540258 | COMP;COL1A1;NTRK2;COL1A2;CCNE2;PDGFD;FN1;FGF2 |
| Relaxin signali | 4/130 | 0,00853441 | 0,0974345 | 0 | 0 | 5,26904095 | 25,0998624 | COL1A1;COL1A2;MMP1;GNAI1 |
| Gap junction | 3/88 | 0,01705122 | 0,13846223 | 0 | 0 | 5,82117647 | 23,7011154 | PDGFD;CDK1;GNAI1 |
| Nitrogen meta | 1/17 | 0,09959855 | 0,3591282 | 0 | 0 | 10,1746926 | 23,4690239 | CA3 |
| Phenylalanine | 1/17 | 0,09959855 | 0,3591282 | 0 | 0 | 10,1746926 | 23,4690239 | MAOA |
| Cocaine addic | 2/49 | 0,03656234 | 0,21222213 | 0 | 0 | 6,97379989 | 23,0744662 | MAOA;GNAI1 |
| Mineral absor | 2/51 | 0,03932841 | 0,21222213 | 0 | 0 | 6,68848035 | 21,6426382 | TF;MT1M |
| IL-17 signaling | 3/93 | 0,01972463 | 0,14222498 | 0 | 0 | 5,49638889 | 21,578202 | CXCL10;MMP1;CXCL2 |
| Small cell lung | 3/93 | 0,01972463 | 0,14222498 | 0 | 0 | 5,49638889 | 21,578202 | CCNE2;CKS2;FN1 |
| Steroid biosyr | 1/19 | 0,11064906 | 0,37451509 | 0 | 0 | 9,04326047 | 19,907759 | SQLE |
| Regulation of | 2/55 | 0,04508917 | 0,21222213 | 0 | 0 | 6,18244191 | 19,1600873 | IRS2;GNAI1 |
| AGE-RAGE sig | 3/100 | 0,0238342 | 0,16326427 | 0 | 0 | 5,09793814 | 19,0491277 | COL1A1;COL1A2;FN1 |
| One carbon p | 1/20 | 0,11612376 | 0,37451509 | 0 | 0 | 8,56686799 | 18,4453131 | TYMS |
| Human T-cell | 5/219 | 0,01145891 | 0,11213358 | 0 | 0 | 3,89335498 | 17,3993568 | CDC20;CCNB2;PTTG1;CCNE2;BUB1B |
| Cellular senes | 4/160 | 0,01718145 | 0,13846223 | 0 | 0 | 4,24929972 | 17,2688364 | CCNB2;CCNB1;CCNE2;CDK1 |
| Viral myocard | 2/59 | 0,05113982 | 0,21894236 | 0 | 0 | 5,74742642 | 17,0882011 | CAV1;DMD |
| Histidine meta | 1/23 | 0,13234809 | 0,37451509 | 0 | 0 | 7,39754098 | 14,9601933 | MAOA |
| Renin-angiote | 1/23 | 0,13234809 | 0,37451509 | 0 | 0 | 7,39754098 | 14,9601933 | MME |
| Platelet activa | 3/124 | 0,04115729 | 0,21222213 | 0 | 0 | 4,08181818 | 13,0224458 | COL1A1;COL1A2;GNAI1 |
| RIG-I-like rece | 2/70 | 0,06913496 | 0,28701483 | 0 | 0 | 4,81502188 | 12,8642688 | CXCL10;ISG15 |
| Human papillo | 6/330 | 0,01639699 | 0,13846223 | 0 | 0 | 3,09480848 | 12,7216971 | COMP;COL1A1;COL1A2;CCNE2;FN1;ISG15 |
| Chemokine sig | 4/190 | 0,02991776 | 0,19517778 | 0 | 0 | 3,55850727 | 12,48788 | CXCL10;CXCL14;CXCL2;GNAI1 |

| | | | | | | | | |
|---|---|---|---|---|---|---|---|---|
| Melanoma | 2/72 | 0,07260184 | 0,2925427 | 0 | 0 | 4,67697757 | 12,2666132 | PDGFD;FGF2 |
| Dopaminergic | 3/131 | 0,04713409 | 0,21222213 | 0 | 0 | 3,85722656 | 11,782897 | GNAL;MAOA;GNAI1 |
| PPAR signaling | 2/74 | 0,07612391 | 0,29797073 | 0 | 0 | 4,54660239 | 11,7092874 | MMP1;SORBS1 |
| FoxO signaling | 3/132 | 0,04802107 | 0,21222213 | 0 | 0 | 3,82713178 | 11,619614 | CCNB2;CCNB1;IRS2 |
| Proteoglycans | 4/201 | 0,03568244 | 0,21222213 | 0 | 0 | 3,35793201 | 11,1923116 | CAV2;CAV1;FN1;FGF2 |
| Human immu | 4/212 | 0,04204774 | 0,21222213 | 0 | 0 | 3,17857143 | 10,0727331 | CCNB2;CCNB1;CDK1;GNAI1 |
| Regulation of | 4/214 | 0,0432698 | 0,21222213 | 0 | 0 | 3,14797919 | 9,88559987 | PDGFD;FN1;FGF2;IQGAP3 |
| Tyrosine meta | 1/36 | 0,19930744 | 0,49153762 | 0 | 0 | 4,64683841 | 7,49491679 | MAOA |
| Prostate canc | 2/97 | 0,12002643 | 0,37451509 | 0 | 0 | 3,44184428 | 7,29685903 | CCNE2;PDGFD |
| Aldosterone-r | 1/37 | 0,20424058 | 0,49153762 | 0 | 0 | 4,51753188 | 7,1759037 | NR3C2 |
| Pathways in ca | 7/530 | 0,04545612 | 0,21222213 | 0 | 0 | 2,23310477 | 6,90254422 | CCNE2;MMP1;FN1;CKS2;BIRC5;FGF2;GNAI1 |
| Longevity regu | 2/102 | 0,13025611 | 0,37451509 | 0 | 0 | 3,26892562 | 6,66289654 | IRS2;CRYAB |
| Chagas diseas | 2/103 | 0,132326 | 0,37451509 | 0 | 0 | 3,23639637 | 6,54556865 | GNAL;GNAI1 |
| Alcoholism | 3/180 | 0,0996122 | 0,3591282 | 0 | 0 | 2,78248588 | 6,41772194 | NTRK2;MAOA;GNAI1 |
| Ferroptosis | 1/40 | 0,21885981 | 0,49153762 | 0 | 0 | 4,16939891 | 6,33466728 | TF |
| Glycine, serine | 1/40 | 0,21885981 | 0,49153762 | 0 | 0 | 4,16939891 | 6,33466728 | MAOA |
| Bladder cance | 1/41 | 0,22367345 | 0,49424617 | 0 | 0 | 4,06495902 | 6,08755302 | MMP1 |
| Tryptophan m | 1/42 | 0,22845766 | 0,49680474 | 0 | 0 | 3,96561375 | 5,85484959 | MAOA |
| TNF signaling | 2/110 | 0,14701818 | 0,40282981 | 0 | 0 | 3,02555862 | 5,80059803 | CXCL10;CXCL2 |
| Serotonergic s | 2/113 | 0,15341456 | 0,41211362 | 0 | 0 | 2,94334003 | 5,51761901 | MAOA;GNAI1 |
| Viral carcinoge | 3/201 | 0,12705258 | 0,37451509 | 0 | 0 | 2,48472222 | 5,12636531 | CDC20;CCNE2;CDK1 |
| Epstein-Barr v | 3/201 | 0,12705258 | 0,37451509 | 0 | 0 | 2,48472222 | 5,12636531 | CXCL10;CCNE2;ISG15 |
| Cytokine-cyto | 4/294 | 0,10797968 | 0,37451509 | 0 | 0 | 2,27029846 | 5,05325813 | CXCL10;INHBA;CXCL14;CXCL2 |
| Type II diabete | 1/46 | 0,24730381 | 0,51334276 | 0 | 0 | 3,61238616 | 5,04700089 | IRS2 |
| Rap1 signaling | 3/206 | 0,13395065 | 0,37451509 | 0 | 0 | 2,4229064 | 4,87072954 | PDGFD;FGF2;GNAI1 |
| Arginine and p | 1/49 | 0,26113811 | 0,52610294 | 0 | 0 | 3,38609973 | 4,5465359 | MAOA |
| Malaria | 1/49 | 0,26113811 | 0,52610294 | 0 | 0 | 3,38609973 | 4,5465359 | COMP |
| Fanconi anem | 1/54 | 0,28363701 | 0,53969819 | 0 | 0 | 3,06588308 | 3,86319665 | UBE2T |
| Ubiquitin mec | 2/137 | 0,20619262 | 0,49153762 | 0 | 0 | 2,41714111 | 3,81653161 | CDC20;UBE2C |
| Insulin signali | 2/137 | 0,20619262 | 0,49153762 | 0 | 0 | 2,41714111 | 3,81653161 | IRS2;SORBS1 |
| Ras signaling p | 3/232 | 0,1717482 | 0,45249046 | 0 | 0 | 2,14497817 | 3,77886337 | NTRK2;PDGFD;FGF2 |
| Legionellosis | 1/55 | 0,28805456 | 0,54059554 | 0 | 0 | 3,00895568 | 3,74496241 | CXCL2 |
| Fluid shear str | 2/139 | 0,21068578 | 0,49153762 | 0 | 0 | 2,38161308 | 3,70909437 | CAV2;CAV1 |
| Signaling path | 2/139 | 0,21068578 | 0,49153762 | 0 | 0 | 2,38161308 | 3,70909437 | INHBA;FGF2 |
| Glutathione m | 1/56 | 0,29244509 | 0,54141861 | 0 | 0 | 2,95409836 | 3,63199999 | RRM2 |

| | | | | | | | | |
|---|---|---|---|---|---|---|---|---|
| Parkinson dise | 2/142 | 0,21744393 | 0,49153762 | 0 | 0 | 2,33022432 | 3,55548945 | GNAL;GNAI1 |
| Gastric cancer | 2/149 | 0,23328333 | 0,49937214 | 0 | 0 | 2,21847417 | 3,22899256 | CCNE2;FGF2 |
| Long-term de | 1/60 | 0,30974027 | 0,56579222 | 0 | 0 | 2,7532648 | 3,22688465 | GNAI1 |
| Cushing syndr | 2/155 | 0,24691631 | 0,51334276 | 0 | 0 | 2,13082699 | 2,98040008 | CCNE2;GNAI1 |
| Cytosolic DNA | 1/63 | 0,32243586 | 0,57368458 | 0 | 0 | 2,61964569 | 2,96504871 | CXCL10 |
| Hepatitis B | 2/163 | 0,26513575 | 0,52610294 | 0 | 0 | 2,02412607 | 2,68705434 | CCNE2;BIRC5 |
| Amphetamine | 1/68 | 0,3430825 | 0,59447095 | 0 | 0 | 2,42353805 | 2,59266304 | MAOA |
| cGMP-PKG sig | 2/166 | 0,27197183 | 0,52610294 | 0 | 0 | 1,98679702 | 2,58692252 | IRS2;GNAI1 |
| Adipocytokine | 1/69 | 0,34713632 | 0,59447095 | 0 | 0 | 2,38777724 | 2,5263584 | IRS2 |
| Renin secretic | 1/69 | 0,34713632 | 0,59447095 | 0 | 0 | 2,38777724 | 2,5263584 | GNAI1 |
| Adherens junc | 1/72 | 0,35914949 | 0,60004245 | 0 | 0 | 2,28653891 | 2,3414537 | SORBS1 |
| Arrhythmoger | 1/72 | 0,35914949 | 0,60004245 | 0 | 0 | 2,28653891 | 2,3414537 | DMD |
| MAPK signalir | 3/295 | 0,27265189 | 0,52610294 | 0 | 0 | 1,67679795 | 2,17909857 | NTRK2;PDGFD;FGF2 |
| Gastric acid se | 1/75 | 0,37094338 | 0,61132439 | 0 | 0 | 2,19350908 | 2,17531576 | GNAI1 |
| Pertussis | 1/76 | 0,37482664 | 0,61132439 | 0 | 0 | 2,16415301 | 2,12366531 | GNAI1 |
| Kaposi sarcon | 2/186 | 0,31740264 | 0,57216003 | 0 | 0 | 1,7690442 | 2,03012707 | FGF2;CXCL2 |
| Hypertrophic | 1/85 | 0,40872305 | 0,64077486 | 0 | 0 | 1,93140125 | 1,7280585 | DMD |
| Colorectal car | 1/86 | 0,41237491 | 0,64077486 | 0 | 0 | 1,90858245 | 1,69066501 | BIRC5 |
| Salmonella inf | 1/86 | 0,41237491 | 0,64077486 | 0 | 0 | 1,90858245 | 1,69066501 | CXCL2 |
| GABAergic syr | 1/89 | 0,42319682 | 0,64077486 | 0 | 0 | 1,8432377 | 1,58503312 | GNAI1 |
| TGF-beta sign | 1/90 | 0,42676001 | 0,64077486 | 0 | 0 | 1,82243507 | 1,55186445 | INHBA |
| Dilated cardio | 1/91 | 0,43030136 | 0,64077486 | 0 | 0 | 1,80209472 | 1,51965145 | DMD |
| Morphine adc | 1/91 | 0,43030136 | 0,64077486 | 0 | 0 | 1,80209472 | 1,51965145 | GNAI1 |
| Rheumatoid a | 1/91 | 0,43030136 | 0,64077486 | 0 | 0 | 1,80209472 | 1,51965145 | MMP1 |
| NF-kappa B si | 1/95 | 0,44425109 | 0,64285224 | 0 | 0 | 1,72506104 | 1,39965477 | CXCL2 |
| Circadian entr | 1/97 | 0,45109836 | 0,64285224 | 0 | 0 | 1,6889515 | 1,3445234 | GNAI1 |
| Hematopoieti | 1/97 | 0,45109836 | 0,64285224 | 0 | 0 | 1,6889515 | 1,3445234 | MME |
| Choline metal | 1/99 | 0,45786195 | 0,64285224 | 0 | 0 | 1,65431582 | 1,29233095 | PDGFD |
| HIF-1 signaling | 1/100 | 0,46121268 | 0,64285224 | 0 | 0 | 1,63752277 | 1,26727233 | TF |
| Melanogenesi | 1/101 | 0,46454286 | 0,64285224 | 0 | 0 | 1,62106557 | 1,24287332 | GNAI1 |
| Toll-like recep | 1/104 | 0,47441141 | 0,64994363 | 0 | 0 | 1,57361133 | 1,17341109 | CXCL10 |
| Parathyroid h | 1/106 | 0,48089 | 0,65229634 | 0 | 0 | 1,54348165 | 1,13000872 | GNAI1 |
| Endocytosis | 2/244 | 0,44390972 | 0,64285224 | 0 | 0 | 1,34109692 | 1,08915051 | CAV2;CAV1 |
| Insulin resista | 1/108 | 0,48728938 | 0,65400926 | 0 | 0 | 1,51447832 | 1,08875411 | IRS2 |
| Cholinergic sy | 1/112 | 0,49985432 | 0,65400926 | 0 | 0 | 1,45960715 | 1,01214793 | GNAI1 |

| | | | | | | | | |
|---|---|---|---|---|---|---|---|---|
| Leukocyte tra | 1/112 | 0,49985432 | 0,65400926 | 0 | 0 | 1,45960715 | 1,01214793 | GNAI1 |
| Toxoplasmosi | 1/113 | 0,50294753 | 0,65400926 | 0 | 0 | 1,44650176 | 0,99413645 | GNAI1 |
| Glutamatergi | 1/114 | 0,50602176 | 0,65400926 | 0 | 0 | 1,43362832 | 0,97655264 | GNAI1 |
| Sphingolipid s | 1/119 | 0,52111231 | 0,65869989 | 0 | 0 | 1,37253404 | 0,89460353 | GNAI1 |
| Neurotrophin | 1/119 | 0,52111231 | 0,65869989 | 0 | 0 | 1,37253404 | 0,89460353 | NTRK2 |
| AMPK signalir | 1/120 | 0,5240751 | 0,65869989 | 0 | 0 | 1,36093126 | 0,8793253 | IRS2 |
| Autophagy | 1/128 | 0,547132 | 0,67425753 | 0 | 0 | 1,27468698 | 0,76871935 | IRS2 |
| Purine metab | 1/129 | 0,54993507 | 0,67425753 | 0 | 0 | 1,26466445 | 0,75621251 | RRM2 |
| Vascular smo | 1/132 | 0,55824144 | 0,67630873 | 0 | 0 | 1,23551495 | 0,7202604 | PPP1R14A |
| Apelin signalir | 1/137 | 0,5717488 | 0,67630873 | 0 | 0 | 1,18979026 | 0,66515884 | GNAI1 |
| Estrogen signa | 1/137 | 0,5717488 | 0,67630873 | 0 | 0 | 1,18979026 | 0,66515884 | GNAI1 |
| Measles | 1/138 | 0,57440071 | 0,67630873 | 0 | 0 | 1,18104583 | 0,65480492 | CCNE2 |
| MicroRNAs in | 2/299 | 0,55121784 | 0,67425753 | 0 | 0 | 1,08968473 | 0,64904369 | CCNE2;IRS2 |
| Apoptosis | 1/143 | 0,58741786 | 0,67630873 | 0 | 0 | 1,13917109 | 0,6060605 | BIRC5 |
| Adrenergic sig | 1/145 | 0,59251345 | 0,67630873 | 0 | 0 | 1,12323543 | 0,58788088 | GNAI1 |
| Breast cancer | 1/147 | 0,59754661 | 0,67630873 | 0 | 0 | 1,10773636 | 0,57039892 | FGF2 |
| Phospholipase | 1/148 | 0,60004002 | 0,67630873 | 0 | 0 | 1,10014498 | 0,56190887 | PDGFD |
| Retrograde er | 1/148 | 0,60004002 | 0,67630873 | 0 | 0 | 1,10014498 | 0,56190887 | GNAI1 |
| Non-alcoholic | 1/149 | 0,6025181 | 0,67630873 | 0 | 0 | 1,09265618 | 0,55358068 | IRS2 |
| Phagosome | 1/152 | 0,60986134 | 0,67630873 | 0 | 0 | 1,07078493 | 0,52952848 | COMP |
| Oxytocin signa | 1/153 | 0,61227906 | 0,67630873 | 0 | 0 | 1,06368637 | 0,52180955 | GNAI1 |
| Hepatitis C | 1/155 | 0,61707001 | 0,67630873 | 0 | 0 | 1,04976581 | 0,50679838 | CXCL10 |
| Wnt signaling | 1/158 | 0,62414654 | 0,67830319 | 0 | 0 | 1,02954996 | 0,48529907 | SFRP1 |
| Hippo signalin | 1/160 | 0,62879201 | 0,67830319 | 0 | 0 | 1,01649655 | 0,4716084 | BIRC5 |
| Protein proce | 1/165 | 0,640158 | 0,68516911 | 0 | 0 | 0,9852559 | 0,43946379 | CRYAB |
| Alzheimer dis | 1/171 | 0,65334236 | 0,68852234 | 0 | 0 | 0,95019286 | 0,40445339 | MME |
| Influenza A | 1/171 | 0,65334236 | 0,68852234 | 0 | 0 | 0,95019286 | 0,40445339 | CXCL10 |
| NOD-like rece | 1/178 | 0,66811958 | 0,69872047 | 0 | 0 | 0,91229045 | 0,3679159 | CXCL2 |
| Axon guidanc | 1/181 | 0,67425978 | 0,69979992 | 0 | 0 | 0,896949 | 0,35352331 | GNAI1 |
| Transcriptiona | 1/186 | 0,68424404 | 0,7035586 | 0 | 0 | 0,8724856 | 0,3310565 | HOXA10 |
| Calcium signal | 1/188 | 0,68815221 | 0,7035586 | 0 | 0 | 0,86306654 | 0,32256701 | GNAL |
| cAMP signalin | 1/212 | 0,73146969 | 0,74230627 | 0 | 0 | 0,7639655 | 0,23889163 | GNAI1 |
| Human cytom | 1/225 | 0,75238358 | 0,75791581 | 0 | 0 | 0,71915252 | 0,20460537 | GNAI1 |
| Olfactory tran | 1/444 | 0,93733125 | 0,93733125 | 0 | 0 | 0,35958258 | 0,02327166 | GNAL |

Table S1. WikiPathways_2015

| Term | Overlap | P-value | Adjusted P-va | Old P-value | Old Adjusted | Odds Ratio | Combined Sco | Genes |
|---|---|---|---|---|---|---|---|---|
| Gastric cancer | 6/43 | 2,42E-07 | 1,77E-05 | 0 | 0 | 27,4982675 | 418,898028 | TOP2A;TPX2;CENPF;UBE2C;S100P;AURKA |
| RB in Cancer(H | 9/87 | 3,21E-09 | 4,68E-07 | 0 | 0 | 20,0394737 | 391,924624 | TOP2A;CCNB2;ANLN;CCNB1;RRM2;CCNE2;CDK1;TTK;TYMS |
| Cell Cycle(Hon | 7/104 | 3,63E-06 | 1,77E-04 | 0 | 0 | 12,3053679 | 154,130002 | CDC20;CCNB2;CCNB1;PTTG1;CCNE2;CDK1;BUB1B |
| Catalytic cycle | 1/5 | 0,03037691 | 0,13044201 | 0 | 0 | 40,7233607 | 142,290378 | FMO2 |
| Neurotransmi | 1/5 | 0,03037691 | 0,13044201 | 0 | 0 | 40,7233607 | 142,290378 | MAOA |
| Inflammatory | 3/30 | 8,16E-04 | 0,01010409 | 0 | 0 | 18,3796296 | 130,693445 | COL1A1;COL1A2;FN1 |
| Inflammatory | 3/31 | 9,00E-04 | 0,01010409 | 0 | 0 | 17,7223214 | 124,295002 | COL1A1;COL1A2;FN1 |
| Fluoropyrimid | 3/35 | 0,00128675 | 0,01252437 | 0 | 0 | 15,5039063 | 103,188341 | RRM2;TK1;TYMS |
| Gastric cancer | 3/35 | 0,00128675 | 0,01252437 | 0 | 0 | 15,5039063 | 103,188341 | TOP2A;UBE2C;UBE2T |
| miRNA regulat | 4/63 | 6,13E-04 | 0,01010409 | 0 | 0 | 11,2906993 | 83,5172474 | CCNB2;CCNB1;CCNE2;CDK1 |
| Focal Adhesio | 8/186 | 1,99E-05 | 7,27E-04 | 0 | 0 | 7,698681 | 83,3253619 | COMP;COL1A1;COL1A2;CAV2;PDGFD;CAV1;COL11A1;FN1 |
| Hedgehog Sign | 2/22 | 0,0079962 | 0,05306568 | 0 | 0 | 16,4107438 | 79,2440187 | CCNB1;CDK1 |
| Integrated Par | 8/195 | 2,80E-05 | 8,17E-04 | 0 | 0 | 7,32480818 | 76,7939415 | TOP2A;ANXA1;GPRC5A;PTTG1;BUB1B;INHBA;TYMS;FGF2 |
| Endochondral | 4/67 | 7,74E-04 | 0,01010409 | 0 | 0 | 10,5716953 | 75,7331046 | CDKN1C;ADAMTS5;COL10A1;FGF2 |
| G1 to S cell cy | 4/67 | 7,74E-04 | 0,01010409 | 0 | 0 | 10,5716953 | 75,7331046 | CDKN1C;CCNB1;CCNE2;CDK1 |
| DNA Damage | 4/69 | 8,65E-04 | 0,01010409 | 0 | 0 | 10,2453782 | 72,259229 | CCNB2;CCNB1;CCNE2;CDK1 |
| Differentiation | 3/47 | 0,00302157 | 0,02594994 | 0 | 0 | 11,26875 | 65,3810548 | TF;INHBA;FGF2 |
| Senescence ar | 5/108 | 5,45E-04 | 0,01010409 | 0 | 0 | 8,13477045 | 61,1261816 | COL1A1;FN1;COL10A1;INHBA;CXCL14 |
| Focal Adhesio | 7/183 | 1,39E-04 | 0,0033881 | 0 | 0 | 6,75484914 | 59,9785461 | COL1A1;COL1A2;CAV2;PDGFD;CAV1;COL11A1;FN1 |
| Type II interfe | 3/50 | 0,00360385 | 0,02923119 | 0 | 0 | 10,5478723 | 59,3397317 | CXCL10;IFI6;ISG15 |
| Nanoparticle-r | 2/28 | 0,01277452 | 0,08109044 | 0 | 0 | 12,6198347 | 55,026299 | COL1A1;FN1 |
| Matrix Metallo | 2/29 | 0,01366612 | 0,08313558 | 0 | 0 | 12,1518212 | 52,1657675 | MMP11;MMP1 |
| Osteoblast(Mu | 1/10 | 0,05983844 | 0,2184103 | 0 | 0 | 18,0947177 | 50,9566617 | COL1A1 |
| Matrix Metallo | 2/30 | 0,01458397 | 0,0851704 | 0 | 0 | 11,7172373 | 49,5385124 | MMP11;MMP1 |
| Endochondral | 3/59 | 0,00574285 | 0,03992646 | 0 | 0 | 8,84866071 | 45,6573209 | ADAMTS5;COL10A1;FGF2 |
| Serotonin Tran | 1/11 | 0,06562331 | 0,22255402 | 0 | 0 | 16,2844262 | 44,3559154 | MAOA |
| Spinal Cord In | 4/96 | 0,00292675 | 0,02594994 | 0 | 0 | 7,22871757 | 42,1713458 | CXCL10;ANXA1;CDK1;CXCL2 |
| Integrated Car | 2/35 | 0,01955477 | 0,10574061 | 0 | 0 | 9,93939394 | 39,1069034 | MMP1;CDK1 |
| Iron Homeost | 1/12 | 0,07137288 | 0,22255402 | 0 | 0 | 14,8032787 | 39,0782475 | TF |
| Iron metabolis | 1/12 | 0,07137288 | 0,22255402 | 0 | 0 | 14,8032787 | 39,0782475 | TF |
| miRNA Regula | 4/105 | 0,00403731 | 0,03102352 | 0 | 0 | 6,58157917 | 36,278832 | CCNB2;CCNB1;CCNE2;CDK1 |
| Quercetin and | 1/13 | 0,07708735 | 0,22255402 | 0 | 0 | 13,5689891 | 34,7748227 | MMP1 |
| Dopamine me | 1/13 | 0,07708735 | 0,22255402 | 0 | 0 | 13,5689891 | 34,7748227 | MAOA |
| ATM Signaling | 2/40 | 0,02513048 | 0,12230166 | 0 | 0 | 8,62940409 | 31,7879108 | CCNB1;CDK1 |
| Spinal Cord In | 4/114 | 0,00540304 | 0,03944222 | 0 | 0 | 6,04033613 | 31,5353437 | CXCL10;ANXA1;CDK1;CXCL2 |
| Osteoblast Sig | 1/14 | 0,08276695 | 0,22255402 | 0 | 0 | 12,5245902 | 31,2078529 | COL1A1 |

| | | | | | | | | |
|---|---|---|---|---|---|---|---|---|
| Biogenic Amin | 1/14 | 0,08276695 | 0,22255402 | 0 | 0 | 12,5245902 | 31,2078529 | MAOA |
| Biogenic Amin | 1/15 | 0,08841187 | 0,22255402 | 0 | 0 | 11,6293911 | 28,209984 | MAOA |
| Cholesterol Bi | 1/15 | 0,08841187 | 0,22255402 | 0 | 0 | 11,6293911 | 28,209984 | SQLE |
| Cholesterol Bi | 1/15 | 0,08841187 | 0,22255402 | 0 | 0 | 11,6293911 | 28,209984 | SQLE |
| GPCRs, Class C | 1/15 | 0,08841187 | 0,22255402 | 0 | 0 | 11,6293911 | 28,209984 | GPRC5A |
| Leptin Insulin | 1/15 | 0,08841187 | 0,22255402 | 0 | 0 | 11,6293911 | 28,209984 | IRS2 |
| Regulation of | 2/44 | 0,02999915 | 0,13044201 | 0 | 0 | 7,8059819 | 27,3723489 | CDK1;KIF2C |
| Type II interfe | 2/44 | 0,02999915 | 0,13044201 | 0 | 0 | 7,8059819 | 27,3723489 | CXCL10;ISG15 |
| GPCRs, Class C | 1/16 | 0,09402234 | 0,22878769 | 0 | 0 | 10,8535519 | 25,6602159 | GPRC5A |
| Neurotransmi | 1/16 | 0,09402234 | 0,22878769 | 0 | 0 | 10,8535519 | 25,6602159 | MAOA |
| ACE Inhibitor I | 1/17 | 0,09959855 | 0,23838342 | 0 | 0 | 10,1746926 | 23,4690239 | NR3C2 |
| TGF Beta Signa | 2/52 | 0,04074045 | 0,16640625 | 0 | 0 | 6,55438017 | 20,9775147 | TGFBR3;INHBA |
| Cardiac Proge | 2/53 | 0,04217145 | 0,16640625 | 0 | 0 | 6,42553881 | 20,3433322 | INHBA;FGF2 |
| IL-4 Signaling I | 2/53 | 0,04217145 | 0,16640625 | 0 | 0 | 6,42553881 | 20,3433322 | BIRC5;IRS2 |
| Integrin-media | 3/97 | 0,02202049 | 0,11482114 | 0 | 0 | 5,26143617 | 20,0764923 | CAV2;CAV1;SORBS1 |
| Nucleotide Me | 1/19 | 0,11064906 | 0,25241816 | 0 | 0 | 9,04326047 | 19,907759 | RRM2 |
| TGF Beta Signa | 2/55 | 0,04508917 | 0,17323734 | 0 | 0 | 6,18244191 | 19,1600873 | TGFBR3;INHBA |
| Integrin-media | 3/100 | 0,0238342 | 0,11999287 | 0 | 0 | 5,09793814 | 19,0491277 | CAV2;CAV1;SORBS1 |
| Nucleotide Me | 1/20 | 0,11612376 | 0,26083182 | 0 | 0 | 8,56686799 | 18,4453131 | RRM2 |
| Nicotine Activ | 1/21 | 0,12156503 | 0,2649029 | 0 | 0 | 8,13811475 | 17,1494977 | GNAI1 |
| G1 to S cell cy | 2/60 | 0,05269574 | 0,19727122 | 0 | 0 | 5,64804788 | 16,6234515 | CCNE2;CDK1 |
| Chemokine sig | 4/165 | 0,01900837 | 0,10574061 | 0 | 0 | 4,11628999 | 16,3123458 | CXCL10;CXCL14;CXCL2;GNAI1 |
| Oxidative Stre | 1/22 | 0,12697307 | 0,26866766 | 0 | 0 | 7,75019516 | 15,9946996 | MAOA |
| Signal Transdu | 1/22 | 0,12697307 | 0,26866766 | 0 | 0 | 7,75019516 | 15,9946996 | GNAI1 |
| Angiogenesis( | 1/23 | 0,13234809 | 0,27215242 | 0 | 0 | 7,39754098 | 14,9601933 | FGF2 |
| Cytokines and | 1/23 | 0,13234809 | 0,27215242 | 0 | 0 | 7,39754098 | 14,9601933 | CXCL2 |
| Signal Transdu | 1/24 | 0,13769029 | 0,27341557 | 0 | 0 | 7,07555239 | 14,0290403 | GNAI1 |
| IL-9 Signaling I | 1/24 | 0,13769029 | 0,27341557 | 0 | 0 | 7,07555239 | 14,0290403 | IRS2 |
| PPAR signaling | 2/68 | 0,06572515 | 0,22255402 | 0 | 0 | 4,96143251 | 13,5063766 | MMP1;SORBS1 |
| Cytokines and | 1/25 | 0,14299986 | 0,27837305 | 0 | 0 | 6,78039617 | 13,1872716 | CXCL2 |
| Primary Focal | 2/70 | 0,06913496 | 0,22255402 | 0 | 0 | 4,81502188 | 12,8642688 | CDKN1C;MME |
| EPO Receptor | 1/26 | 0,14827699 | 0,28114859 | 0 | 0 | 6,50885246 | 12,423272 | IRS2 |
| EPO Receptor | 1/26 | 0,14827699 | 0,28114859 | 0 | 0 | 6,50885246 | 12,423272 | IRS2 |
| One Carbon M | 1/28 | 0,15873477 | 0,29335793 | 0 | 0 | 6,02610808 | 11,0911759 | TYMS |
| Extracellular v | 1/28 | 0,15873477 | 0,29335793 | 0 | 0 | 6,02610808 | 11,0911759 | TGFBR3 |
| Apoptosis Mo | 2/80 | 0,08700295 | 0,22255402 | 0 | 0 | 4,19559229 | 10,2448527 | BIRC5;HN1 |
| One Carbon M | 1/30 | 0,16906518 | 0,29739176 | 0 | 0 | 5,60994912 | 9,97152177 | TYMS |
| Prostaglandin | 1/30 | 0,16906518 | 0,29739176 | 0 | 0 | 5,60994912 | 9,97152177 | ANXA1 |

| | | | | | | | | |
|---|---|---|---|---|---|---|---|---|
| Dopminergic I | 1/30 | 0,16906518 | 0,29739176 | 0 | 0 | 5,60994912 | 9,97152177 | CDKN1C |
| TCA Cycle(Mu | 1/30 | 0,16906518 | 0,29739176 | 0 | 0 | 5,60994912 | 9,97152177 | PDK4 |
| Prostaglandin | 1/31 | 0,17418309 | 0,30084348 | 0 | 0 | 5,4226776 | 9,47693309 | ANXA1 |
| Integrated Bre | 3/151 | 0,06639735 | 0,22255402 | 0 | 0 | 3,33260135 | 9,03834202 | ANXA1;MMP1;AURKA |
| Oxidative Stre | 1/32 | 0,17926975 | 0,30084348 | 0 | 0 | 5,2474881 | 9,01971654 | MAOA |
| Statin Pathwa | 1/32 | 0,17926975 | 0,30084348 | 0 | 0 | 5,2474881 | 9,01971654 | SQLE |
| Trans-sulfurat | 1/32 | 0,17926975 | 0,30084348 | 0 | 0 | 5,2474881 | 9,01971654 | TYMS |
| G Protein Sign | 2/88 | 0,10217292 | 0,24060075 | 0 | 0 | 3,80376706 | 8,67672967 | GNAL;GNAI1 |
| Alpha 6 Beta 4 | 1/33 | 0,18432532 | 0,30237636 | 0 | 0 | 5,08324795 | 8,59604194 | IRS2 |
| Amino Acid m | 2/92 | 0,11001275 | 0,25241816 | 0 | 0 | 3,63397612 | 8,02076309 | MAOA;PDK4 |
| SIDS Susceptib | 3/161 | 0,0771817 | 0,22255402 | 0 | 0 | 3,12009494 | 7,99241283 | NTRK2;TF;MAOA |
| EGF/EGFR Sign | 3/161 | 0,0771817 | 0,22255402 | 0 | 0 | 3,12009494 | 7,99241283 | CAV2;CAV1;AURKA |
| Endothelin Pa | 1/35 | 0,19434398 | 0,31526913 | 0 | 0 | 4,78375121 | 7,83638527 | GNAI1 |
| G Protein Sign | 2/96 | 0,11800568 | 0,26104288 | 0 | 0 | 3,47863548 | 7,43392223 | GNAL;GNAI1 |
| Striated Musc | 1/38 | 0,20914356 | 0,33067346 | 0 | 0 | 4,39521489 | 6,8773438 | DMD |
| Parkinsons Dis | 1/40 | 0,21885981 | 0,33067346 | 0 | 0 | 4,16939891 | 6,33466728 | CCNE2 |
| Interleukin-11 | 1/40 | 0,21885981 | 0,33067346 | 0 | 0 | 4,16939891 | 6,33466728 | BIRC5 |
| Wnt Signaling | 2/106 | 0,13858049 | 0,27341557 | 0 | 0 | 3,14256198 | 6,21065764 | SFRP1;CDK1 |
| Striated Musc | 1/41 | 0,22367345 | 0,33067346 | 0 | 0 | 4,06495902 | 6,08755302 | DMD |
| Splicing factor | 1/42 | 0,22845766 | 0,33067346 | 0 | 0 | 3,96561375 | 5,85484959 | CAV2 |
| Aryl Hydrocarl | 1/43 | 0,23321262 | 0,33067346 | 0 | 0 | 3,87099922 | 5,63541889 | PLAGL1 |
| Hair Follicle D | 1/43 | 0,23321262 | 0,33067346 | 0 | 0 | 3,87099922 | 5,63541889 | INHBA |
| IL-7 Signaling I | 1/43 | 0,23321262 | 0,33067346 | 0 | 0 | 3,87099922 | 5,63541889 | IRS2 |
| One carbon m | 1/44 | 0,23793852 | 0,33402907 | 0 | 0 | 3,78078536 | 5,42823602 | TYMS |
| Apoptosis-rela | 1/52 | 0,27472018 | 0,37838817 | 0 | 0 | 3,18643523 | 4,11688147 | BIRC5 |
| Vitamin B12 M | 1/54 | 0,28363701 | 0,38613899 | 0 | 0 | 3,06588308 | 3,86319665 | SAA2 |
| Cardiac Hyper | 1/56 | 0,29244509 | 0,39171544 | 0 | 0 | 2,95409836 | 3,63199999 | FGF2 |
| BDNF signalin | 2/142 | 0,21744393 | 0,33067346 | 0 | 0 | 2,33022432 | 3,55548945 | NTRK2;IRS2 |
| Calcium Regul | 2/143 | 0,21970111 | 0,33067346 | 0 | 0 | 2,31358068 | 3,506202 | GJB2;GNAI1 |
| Regulation of | 2/144 | 0,22196034 | 0,33067346 | 0 | 0 | 2,29717146 | 3,45783246 | FN1;FGF2 |
| SIDS Susceptib | 1/59 | 0,30545619 | 0,40542367 | 0 | 0 | 2,8008762 | 3,32169607 | MAOA |
| Regulation of | 2/148 | 0,23101551 | 0,33067346 | 0 | 0 | 2,23378241 | 3,27309535 | FN1;FGF2 |
| Calcium Regul | 2/149 | 0,23328333 | 0,33067346 | 0 | 0 | 2,21847417 | 3,22899256 | GJB2;GNAI1 |
| Interferon typ | 1/61 | 0,31399813 | 0,41300655 | 0 | 0 | 2,70724044 | 3,13598135 | IRS2 |
| miR-targeted | 4/362 | 0,18393964 | 0,30237636 | 0 | 0 | 1,83268391 | 3,10300445 | COL1A1;CDKN1C;TYMS;IQGAP3 |
| Alpha6-Beta4 | 1/64 | 0,32661604 | 0,42199949 | 0 | 0 | 2,57793391 | 2,88463066 | IRS2 |
| SREBP signalli | 1/64 | 0,32661604 | 0,42199949 | 0 | 0 | 2,57793391 | 2,88463066 | SQLE |
| Oncostatin M | 1/65 | 0,33077064 | 0,42361854 | 0 | 0 | 2,53752561 | 2,8073409 | MMP1 |

| | | | | | | | | |
|---|---|---|---|---|---|---|---|---|
| Insulin Signali | 2/163 | 0,26513575 | 0,36866494 | 0 | 0 | 2,02412607 | 2,68705434 | IRS2;SORBS1 |
| TSH signaling | 1/67 | 0,33900372 | 0,43038733 | 0 | 0 | 2,46038251 | 2,66150454 | GNAI1 |
| AMPK Signalir | 1/69 | 0,34713632 | 0,43691295 | 0 | 0 | 2,38777724 | 2,5263584 | CCNB1 |
| Folate Metabo | 1/70 | 0,35116532 | 0,4382063 | 0 | 0 | 2,35305298 | 2,46246561 | SAA2 |
| EGFR1 Signali | 2/172 | 0,28563706 | 0,38613899 | 0 | 0 | 1,9160914 | 2,40092633 | CAV2;CAV1 |
| Alzheimers Di: | 1/73 | 0,36310496 | 0,44926546 | 0 | 0 | 2,25466758 | 2,28412108 | MME |
| Arrhythmoger | 1/74 | 0,3670362 | 0,45031333 | 0 | 0 | 2,22366944 | 2,22877228 | DMD |
| Alzheimers Di: | 1/79 | 0,3863343 | 0,46615544 | 0 | 0 | 2,08060109 | 1,97876027 | MME |
| Apoptosis(Mu | 1/79 | 0,3863343 | 0,46615544 | 0 | 0 | 2,08060109 | 1,97876027 | BIRC5 |
| MicroRNAs in | 1/80 | 0,39012331 | 0,46686888 | 0 | 0 | 2,05416061 | 1,93356581 | FGF2 |
| Apoptosis(Ho | 1/86 | 0,41237491 | 0,48948567 | 0 | 0 | 1,90858245 | 1,69066501 | BIRC5 |
| Selenium Micı | 1/88 | 0,41961166 | 0,4940589 | 0 | 0 | 1,86451856 | 1,61919566 | SAA2 |
| Androgen rece | 1/90 | 0,42676001 | 0,49449969 | 0 | 0 | 1,82243507 | 1,55186445 | CAV1 |
| Corticotropin- | 1/90 | 0,42676001 | 0,49449969 | 0 | 0 | 1,82243507 | 1,55186445 | GNAI1 |
| IL-3 Signaling | 1/97 | 0,45109836 | 0,51858552 | 0 | 0 | 1,6889515 | 1,3445234 | BIRC5 |
| Neural Crest D | 1/101 | 0,46454286 | 0,5298692 | 0 | 0 | 1,62106557 | 1,24287332 | FGF2 |
| Toll-like recep | 1/104 | 0,47441141 | 0,53693075 | 0 | 0 | 1,57361133 | 1,17341109 | CXCL10 |
| MicroRNAs in | 1/109 | 0,49045966 | 0,55082393 | 0 | 0 | 1,50037948 | 1,0688887 | FGF2 |
| Iron uptake ar | 1/115 | 0,50907714 | 0,56642886 | 0 | 0 | 1,42098073 | 0,95938328 | TF |
| ESC Pluripoter | 1/116 | 0,51211376 | 0,56642886 | 0 | 0 | 1,4085531 | 0,94261568 | FGF2 |
| Adipogenesis | 1/129 | 0,54993507 | 0,59918298 | 0 | 0 | 1,26466445 | 0,75621251 | IRS2 |
| Metapathway | 1/129 | 0,54993507 | 0,59918298 | 0 | 0 | 1,26466445 | 0,75621251 | FMO2 |
| Adipogenesis( | 1/134 | 0,56369436 | 0,60526179 | 0 | 0 | 1,21681252 | 0,69752937 | IRS2 |
| PodNet: prote | 2/306 | 0,5638055 | 0,60526179 | 0 | 0 | 1,0642127 | 0,60984277 | CXCL10;BIRC5 |
| Regulation of | 1/146 | 0,59503778 | 0,63412786 | 0 | 0 | 1,11543245 | 0,57905486 | CXCL10 |
| Insulin Signali | 1/153 | 0,61227906 | 0,6477735 | 0 | 0 | 1,06368637 | 0,52180955 | SORBS1 |
| Purine metabo | 1/158 | 0,62414654 | 0,65557838 | 0 | 0 | 1,02954996 | 0,48529907 | RRM2 |
| miR-targeted | 1/166 | 0,64238943 | 0,66992041 | 0 | 0 | 0,97923497 | 0,43337078 | TYMS |
| miR-targeted | 1/172 | 0,65549268 | 0,67873711 | 0 | 0 | 0,94458825 | 0,39896398 | FGF2 |
| TNF-alpha NF- | 1/179 | 0,67017896 | 0,68840509 | 0 | 0 | 0,90711917 | 0,36303861 | CAV1 |
| Metapathway | 1/181 | 0,67425978 | 0,68840509 | 0 | 0 | 0,896949 | 0,35352331 | FMO2 |
| Non-odorant ( | 1/256 | 0,79596052 | 0,80701553 | 0 | 0 | 0,63072967 | 0,1439361 | GPRC5A |
| PluriNetWork | 1/284 | 0,82873423 | 0,83444964 | 0 | 0 | 0,56751434 | 0,10661084 | UHRF1 |
| mRNA proces: | 1/398 | 0,91625787 | 0,91625787 | 0 | 0 | 0,4021968 | 0,0351751 | RBMS3 |

Table S2. GSE31519.

| SampleName | datas | Array_type | GSM_accessic | Biops | nodal | age | tumo | grade | event | event | adjuv | 206665_s_at Bcl | MEDIAN = -0,004 | 204252_at CDK2 | MEDIAN = 0,004 | 203213_at Cdk1 | MEDIAN = 0,006 | 202246_s_at Cdk | MEDIAN = 0,008 | 207143_at CDK6 | MEDIAN = -0,003 | 202580_x_at Fox | MEDIAN= 0,0040 | 204092_s_at Aur | MEDIAN= 0,0044 | 209464_at AurkB | MEDIAN = 0,001 | 212533_at Wee1 | MEDIAN = 0,005129 |
|---|---|---|---|---|---|---|---|---|---|---|---|---|---|---|---|---|---|---|---|---|---|---|---|---|---|---|---|---|---|
| TNBC_001 | 1 | GPL96 | GSM782523 | 1 | 0 | 59 | 2 | 3 | 93 | 0 | NO | -0,002958 | | 0,003285 | | 0,00418 | | 0,008441 | | -0,000767 | | 0,004304 | | 0,006657 | | 0,001442 | | 0,004491 | |
| TNBC_002 | 1 | GPL96 | GSM782524 | 1 | 0 | 65 | 2 | 12 | 37 | 0 | NO | -0,002184 | | 0,003536 | | 0,006012 | | 0,010858 | | -0,00565 | | 0,000883 | | 0,004826 | | 0,000528 | | 0,002893 | |
| TNBC_003 | 1 | GPL96 | GSM782525 | 1 | 1 | 66 | 2 | 12 | 31 | 0 | NO | -0,011604 | | 0,004552 | | 0,005921 | | 0,008522 | | -0,003047 | | 0,00161 | | 0,00362 | | -0,00409 | | 0,006735 | |
| TNBC_004 | 1 | GPL96 | GSM782526 | 1 | 0 | | | | | | NO | -0,013867 | | 0,004121 | | 0,007375 | | 0,006574 | | -0,000547 | | 0,004008 | | 0,00033 | | 0,000694 | | 0,006116 | |
| TNBC_005 | 1 | GPL96 | GSM782527 | 1 | 1 | | | | | | NO | -0,005577 | | 0,004448 | | 0,004856 | | 0,009406 | | -0,003133 | | 0,001876 | | 0,001655 | | 0,00091 | | 0,005595 | |
| TNBC_006 | 1 | GPL96 | GSM782528 | 1 | 0 | 60 | 2 | 12 | 120 | 0 | NO | -0,011456 | | 0,004132 | | 0,000932 | | 0,008469 | | -0,008924 | | 0,003279 | | 0,002474 | | -0,001456 | | 0,007706 | |
| TNBC_007 | 1 | GPL96 | GSM782529 | 1 | 0 | 57 | 2 | 3 | 25 | 1 | NO | -0,010638 | | 0,004279 | | 0,003992 | | 0,008222 | | -0,002437 | | 0,004311 | | 0,003622 | | 0,000774 | | 0,007266 | |
| TNBC_008 | 2 | GPL96 | GSM782530 | 1 | 1 | 46 | 1 | 12 | 23 | 1 | YES | -0,005653 | | 0,006488 | | 0,002608 | | 0,013323 | | -0,00193 | | -0,003385 | | 0,002261 | | -0,004651 | | 0,002943 | |
| TNBC_009 | 2 | GPL96 | GSM782531 | 1 | 0 | 55 | 2 | 3 | 21 | 1 | YES | 0,00091 | | 0,004673 | | 0,004147 | | 0,009802 | | -0,000725 | | 0,00514 | | 0,003852 | | 0,00001 | | 0,001386 | |
| TNBC_010 | 2 | GPL96 | GSM782532 | 1 | 0 | 61 | 2 | 3 | 45 | 0 | YES | -0,004565 | | 0,003617 | | 0,002205 | | 0,007898 | | -0,004851 | | 0,001676 | | 0,001742 | | -0,001946 | | 0,003999 | |
| TNBC_011 | 2 | GPL96 | GSM782533 | 1 | 0 | 29 | 1 | 3 | 45 | 1 | YES | -0,006712 | | 0,004987 | | 0,007575 | | 0,009869 | | -0,00761 | | 0,002873 | | 0,003671 | | -0,001149 | | 0,003628 | |
| TNBC_012 | 2 | GPL96 | GSM782534 | 1 | | 69 | 1 | 3 | 52 | 0 | YES | -0,00074 | | 0,005474 | | 0,003782 | | 0,008851 | | -0,002667 | | 0,007149 | | 0,006029 | | 0,002048 | | 0,003067 | |
| TNBC_013 | 2 | GPL96 | GSM782535 | 1 | 1 | 47 | 2 | 3 | 16 | 1 | YES | -0,000904 | | 0,004052 | | 0,003298 | | 0,008554 | | -0,008195 | | 0,000577 | | 0,003768 | | 0,0005 | | 0,000241 | |
| TNBC_014 | 2 | GPL96 | GSM782536 | 1 | 0 | 51 | 1 | 12 | 70 | 0 | YES | -0,00406 | | 0,004113 | | 0,00553 | | 0,009481 | | -0,003865 | | 0,003475 | | 0,000114 | | 0,000858 | | 0,002797 | |
| TNBC_015 | 2 | GPL96 | GSM782537 | 1 | 0 | 51 | 2 | 12 | 77 | 0 | YES | 0,000635 | | 0,001768 | | 0,002073 | | 0,007535 | | -0,001199 | | 0,000497 | | 0,002408 | | -0,000974 | | 0,002325 | |
| TNBC_016 | 2 | GPL96 | GSM782538 | 1 | 0 | 57 | 2 | 3 | 47 | 0 | YES | -0,00809 | | 0,005426 | | 0,007126 | | 0,010951 | | -0,003125 | | 0,006831 | | 0,004625 | | 0,001995 | | 0,007268 | |
| TNBC_017 | 2 | GPL96 | GSM782539 | 1 | 1 | 80 | 2 | 3 | 11 | 1 | YES | -0,000585 | | 0,003954 | | 0,002814 | | 0,008472 | | -0,004793 | | 0,005081 | | -0,000293 | | 0,00247 | | 0,004354 | |
| TNBC_018 | 2 | GPL96 | GSM782540 | 1 | 0 | 48 | 2 | 3 | 76 | 0 | YES | -0,002578 | | 0,005595 | | 0,003848 | | 0,010732 | | -0,003981 | | 0,006581 | | 0,003435 | | 0,000609 | | 0,006098 | |
| TNBC_019 | 2 | GPL96 | GSM782541 | 1 | 0 | 53 | 2 | 3 | 69 | 0 | YES | -0,000529 | | 0,005678 | | 0,005096 | | 0,011099 | | -0,002094 | | 0,001432 | | 0,002026 | | -0,000092 | | 0,005187 | |
| TNBC_020 | 2 | GPL96 | GSM782542 | 1 | 0 | 53 | 1 | 3 | 28 | 0 | YES | -0,000726 | | 0,006516 | | 0,005703 | | 0,011776 | | -0,002961 | | 0,010893 | | 0,002887 | | 0,000743 | | 0,004308 | |
| TNBC_021 | 2 | GPL96 | GSM782543 | 1 | 0 | 67 | 2 | 12 | 34 | 0 | YES | -0,000486 | | 0,003464 | | -0,003234 | | 0,006512 | | -0,005703 | | -0,006835 | | 0,001113 | | -0,005604 | | 0,002696 | |
| TNBC_022 | 2 | GPL96 | GSM782544 | 1 | 0 | 56 | 2 | 3 | 10 | 1 | YES | -0,001126 | | 0,006658 | | 0,004444 | | 0,012656 | | -0,000172 | | 0,00679 | | 0,001872 | | 0,005661 | | 0,00452 | |
| TNBC_023 | 2 | GPL96 | GSM782545 | 1 | 1 | 53 | 2 | 3 | 29 | 0 | YES | 0,000768 | | 0,004916 | | 0,00696 | | 0,009014 | | 0,001547 | | 0,009019 | | 0,003199 | | 0,00342 | | 0,005419 | |
| TNBC_024 | 2 | GPL96 | GSM782546 | 1 | 0 | 63 | 1 | 3 | 36 | 0 | YES | -0,001015 | | 0,002052 | | -0,004994 | | 0,010749 | | -0,011838 | | -0,005597 | | 0,002378 | | -0,003245 | | 0,000581 | |
| TNBC_025 | 2 | GPL96 | GSM782547 | 1 | 0 | 47 | 2 | 3 | 31 | 0 | YES | -0,000265 | | 0,004724 | | 0,004215 | | 0,008398 | | -0,008395 | | 0,008263 | | 0,006453 | | 0,003136 | | 0,003372 | |
| TNBC_026 | 2 | GPL96 | GSM782548 | 1 | 0 | 45 | 1 | 3 | 36 | 0 | YES | -0,002184 | | 0,004784 | | 0,005162 | | 0,010091 | | -0,003442 | | 0,005687 | | 0,003399 | | 0,002833 | | 0,001624 | |
| TNBC_027 | 2 | GPL96 | GSM782549 | 1 | 0 | 42 | 2 | 3 | 6 | 1 | YES | -0,002787 | | 0,005446 | | 0,009752 | | 0,009427 | | -0,001479 | | 0,008935 | | 0,004591 | | 0,00552 | | 0,005911 | |
| TNBC_028 | 2 | GPL96 | GSM782550 | 1 | 0 | 51 | 1 | 3 | 33 | 0 | YES | -0,000543 | | 0,004677 | | 0,003572 | | 0,010773 | | -0,004153 | | 0,004828 | | 0,003636 | | 0,002191 | | 0,003005 | |
| TNBC_029 | 2 | GPL96 | GSM782551 | 1 | 0 | 54 | 2 | 3 | 10 | 1 | YES | -0,000808 | | 0,004457 | | 0,001781 | | 0,010468 | | -0,00635 | | -0,000426 | | 0,006136 | | -0,000359 | | 0,005349 | |
| TNBC_030 | 2 | GPL96 | GSM782552 | 1 | 0 | 40 | 1 | 3 | 27 | 0 | YES | -0,001569 | | 0,004811 | | 0,004632 | | 0,009 | | -0,001711 | | 0,005306 | | -0,001049 | | 0,000706 | | 0,001818 | |
| TNBC_031 | 2 | GPL96 | GSM782553 | 1 | 0 | 57 | 1 | 3 | 45 | 0 | YES | -0,001147 | | 0,005407 | | 0,004955 | | 0,010092 | | -0,001816 | | 0,006299 | | 0,008985 | | 0,000314 | | 0,005251 | |
| TNBC_032 | 3 | GPL96 & GPL97 | GSM79115 | 1 | 0 | 40 | 1 | 3 | 120 | 0 | NO | -0,011548 | | 0,005412 | | 0,008804 | | 0,007584 | | -0,00282 | | 0,008651 | | 0,006517 | | 0,00404 | | 0,006822 | |
| TNBC_033 | 3 | GPL96 & GPL97 | GSM79117 | 1 | 1 | 52 | 2 | 3 | 120 | 0 | | -0,011003 | | 0,00467 | | 0,004631 | | 0,008628 | | -0,00756 | | 0,003087 | | 0,012266 | | -0,00095 | | 0,003048 | |
| TNBC_034 | 3 | GPL96 & GPL97 | GSM79122 | 1 | 0 | 74 | 1 | 12 | 120 | 0 | NO | -0,013772 | | 0,007121 | | 0,008747 | | 0,008386 | | -0,000831 | | 0,009217 | | 0,00379 | | 0,003362 | | 0,004012 | |
| TNBC_035 | 3 | GPL96 & GPL97 | GSM79145 | 1 | 0 | 40 | 2 | 12 | 120 | 0 | NO | -0,004879 | | 0,002342 | | 0,000195 | | 0,008314 | | -0,008118 | | 0,000624 | | 0,004364 | | -0,004371 | | 0,00203 | |
| TNBC_036 | 3 | GPL96 & GPL97 | GSM79147 | 1 | 1 | 34 | 2 | 3 | 8 | 0 | | -0,006145 | | 0,004708 | | 0,006217 | | 0,006972 | | -0,003594 | | 0,004995 | | 0,006676 | | 0,000755 | | 0,007079 | |
| TNBC_037 | 3 | GPL96 & GPL97 | GSM79165 | 1 | 0 | 62 | 2 | 12 | 120 | 0 | NO | -0,002441 | | 0,006255 | | 0,004272 | | 0,010526 | | -0,010984 | | 0,00267 | | 0,003482 | | -0,000267 | | 0,00431 | |
| TNBC_038 | 3 | GPL96 & GPL97 | GSM79191 | 1 | 0 | 58 | 2 | 12 | 120 | 0 | | -0,005865 | | 0,005088 | | 0,00703 | | 0,009244 | | -0,009905 | | 0,002835 | | 0,006083 | | 0 | | 0,004622 | |
| TNBC_039 | 3 | GPL96 & GPL97 | GSM79195 | 1 | 1 | 68 | 1 | 12 | 120 | 0 | | -0,006701 | | 0,004367 | | 0,003765 | | 0,0064 | | -0,013064 | | -0,001054 | | 0,006042 | | -0,000527 | | 0,001882 | |
| TNBC_040 | 3 | GPL96 & GPL97 | GSM79196 | 1 | 1 | 77 | 2 | 3 | 43 | 0 | NO | -0,005028 | | 0,004719 | | 0,002901 | | 0,009206 | | -0,006189 | | 0,003752 | | 0,006453 | | 0,001509 | | 0,00147 | |
| TNBC_041 | 3 | GPL96 & GPL97 | GSM79225 | 1 | 0 | 70 | 1 | 12 | 95 | 0 | NO | -0,016826 | | 0,004308 | | 0,000691 | | 0,007763 | | -0,010364 | | -0,004064 | | 0,008339 | | -0,010242 | | 0,008007 | |
| TNBC_042 | 3 | GPL96 & GPL97 | GSM79231 | 1 | 1 | 77 | 2 | 12 | 6 | 1 | NO | -0,004089 | | 0,006261 | | 0,006623 | | 0,008649 | | -0,007455 | | 0,003402 | | -0,002002 | | -0,001628 | | 0,006948 | |
| TNBC_043 | 3 | GPL96 & GPL97 | GSM79251 | 1 | | 88 | 2 | 3 | 2 | 0 | NO | -0,011043 | | 0,003467 | | 0,006068 | | 0,009799 | | -0,006671 | | 0,004297 | | 0,00628 | | -0,001734 | | 0,003317 | |
| TNBC_044 | 3 | GPL96 & GPL97 | GSM79253 | 1 | 0 | 75 | 1 | 3 | 30 | 1 | NO | -0,005689 | | 0,004389 | | 0,004064 | | 0,008208 | | -0,003454 | | 0,002154 | | 0,003617 | | 0,002519 | | 0,003291 | |
| TNBC_045 | 3 | GPL96 & GPL97 | GSM79255 | 1 | 0 | 77 | 2 | 3 | 37 | 1 | NO | -0,0111 | | 0,005123 | | 0,00361 | | 0,007801 | | -0,011372 | | 0,004813 | | 0,005785 | | 0,001785 | | 0,004735 | |
| TNBC_046 | 3 | GPL96 & GPL97 | GSM79270 | 1 | 0 | 32 | 1 | 12 | 120 | 0 | NO | -0,006358 | | 0,006321 | | 0,008014 | | 0,009331 | | -0,009067 | | 0,009857 | | 0,00605 | | 0,001279 | | 0,003349 | |
| TNBC_047 | 3 | GPL96 & GPL97 | GSM79271 | 1 | 1 | 46 | 2 | 3 | 0 | 0 | NO | -0,00309 | | 0,006026 | | 0,009656 | | 0,009309 | | -0,006875 | | 0,011511 | | 0,005172 | | 0,00224 | | 0,00479 | |
| TNBC_048 | 3 | GPL96 & GPL97 | GSM79280 | 1 | 0 | 50 | 1 | 3 | 120 | 0 | NO | -0,004422 | | 0,005152 | | 0,003935 | | 0,008235 | | -0,002556 | | 0,003448 | | 0,003369 | | 0,001257 | | 0,004787 | |
| TNBC_049 | 3 | GPL96 & GPL97 | GSM79287 | 1 | 1 | 37 | 1 | 3 | 120 | 0 | | -0,015205 | | 0,006182 | | 0,003968 | | 0,006642 | | -0,002632 | | 0,004678 | | 0,007053 | | 0,000585 | | 0,004135 | |
| TNBC_050 | 3 | GPL96 & GPL97 | GSM79292 | 1 | 0 | 84 | 2 | 3 | 11 | 0 | | -0,006641 | | 0,007176 | | 0,006183 | | 0,008626 | | -0,008168 | | 0,004389 | | 0,008377 | | 0,002863 | | 0,005993 | |
| TNBC_051 | 3 | GPL96 & GPL97 | GSM79299 | 1 | 0 | 37 | 2 | 3 | 119 | 0 | NO | -0,015931 | | 0,007027 | | 0,006428 | | 0,0107 | | -0,004751 | | 0,006109 | | 0,007206 | | 0,003114 | | 0,006229 | |
| TNBC_052 | 3 | GPL96 & GPL97 | GSM79303 | 1 | 0 | 44 | 1 | 12 | 118 | 0 | NO | -0,010721 | | 0,007147 | | 0,002449 | | 0,009195 | | -0,002971 | | 0,004417 | | 0,006516 | | -0,000201 | | 0,00269 | |
| TNBC_053 | 3 | GPL96 & GPL97 | GSM79306 | 1 | 0 | 38 | 2 | 3 | 2 | 1 | NO | -0,019967 | | 0,006497 | | 0,006339 | | 0,008399 | | -0,00721 | | 0,00206 | | 0,006706 | | 0,001109 | | 0,004833 | |
| TNBC_054 | 3 | GPL96 & GPL97 | GSM79322 | 1 | 1 | 61 | 2 | 12 | 115 | 1 | NO | -0,01387 | | 0,004041 | | 0,00557 | | 0,007463 | | -0,004223 | | 0,002366 | | 0,006874 | | 0,000728 | | 0,004514 | |
| TNBC_055 | 3 | GPL96 & GPL97 | GSM79329 | 1 | 0 | 68 | 1 | 3 | 7 | 1 | NO | -0,014142 | | 0,005359 | | 0,006476 | | 0,008634 | | -0,003573 | | 0,007927 | | 0,005432 | | 0,001191 | | 0,00428 | |
| TNBC_056 | 3 | GPL96 & GPL97 | GSM79336 | 1 | 0 | 77 | 2 | 3 | 3 | 1 | NO | -0,010975 | | 0,006271 | | 0,006898 | | 0,00682 | | -0,006546 | | 0,006507 | | 0,007755 | | 0,001607 | | 0,005919 | |
| TNBC_057 | 3 | GPL96 & GPL97 | GSM79344 | 1 | | 87 | 1 | 3 | 120 | 0 | NO | -0,008375 | | 0,004694 | | 0,006384 | | 0,008299 | | -0,006797 | | 0,001577 | | 0,00716 | | 0,00154 | | 0,005858 | |
| TNBC_058 | 3 | GPL96 & GPL97 | GSM79356 | 1 | 0 | 61 | 2 | 12 | 120 | 0 | NO | -0,008706 | | 0,004568 | | 0,007379 | | 0,007535 | | 0,00082 | | 0,011712 | | 0,007616 | | 0,003553 | | 0,005036 | |
| TNBC_059 | 4 | GPL96 | GSM782554 | 1 | 1 | 52 | 2 | 3 | 6 | 1 | YES | -0,003684 | | 0,005326 | | 0,00621 | | 0,010836 | | -0,001819 | | 0,007425 | | 0,007541 | | 0,003466 | | 0,003509 | |
| TNBC_060 | 4 | GPL96 | GSM782555 | 1 | 0 | 41 | 2 | 3 | 67 | 0 | YES | -0,000551 | | 0,00241 | | 0,000231 | | 0,008649 | | -0,004159 | | 0,000911 | | 0,002795 | | -0,00111 | | -0,001178 | |
| TNBC_061 | 4 | GPL96 | GSM782556 | 1 | 1 | 41 | 2 | 3 | 11 | 1 | YES | -0,000661 | | 0,002663 | | 0,00138 | | 0,008728 | | -0,005918 | | -0,000157 | | 0,00127 | | -0,001395 | | 0,001142 | |
| TNBC_062 | 4 | GPL96 | GSM782557 | 1 | 0 | 30 | 2 | 3 | 51 | 0 | YES | -0,002217 | | 0,004407 | | 0,003892 | | 0,011167 | | -0,002618 | | 0,005591 | | 0,003285 | | -0,000803 | | 0,001837 | |
| TNBC_063 | 4 | GPL96 | GSM782558 | 1 | 0 | 59 | 1 | 12 | 48 | 0 | YES | 0,000549 | | 0,005462 | | 0,002925 | | 0,008977 | | -0,007677 | | 0,002863 | | 0,003189 | | 0,000008 | | 0,003223 | |
| TNBC_064 | 4 | GPL96 | GSM782559 | 1 | 1 | 71 | 2 | 3 | 58 | 0 | YES | 0,001572 | | 0,002871 | | 0,005044 | | 0,008134 | | -0,002143 | | 0,007754 | | 0,004764 | | 0,001942 | | 0,005073 | |
| TNBC_065 | 4 | GPL96 | GSM782560 | 1 | 0 | 43 | 2 | 3 | 3 | 1 | YES | -0,000952 | | 0,003716 | | 0,003746 | | 0,00752 | | -0,006218 | | 0,004997 | | 0,00281 | | -0,000417 | | 0,003672 | |
| TNBC_066 | 4 | GPL96 | GSM782561 | 1 | 0 | 60 | 1 | 3 | 48 | 0 | YES | 0,000886 | | 0,004013 | | 0,000943 | | 0,009311 | | -0,010963 | | 0,00851 | | 0,004265 | | 0,002905 | | 0,0054 | |
| TNBC_067 | 4 | GPL96 | GSM782562 | 1 | 0 | 31 | 1 | 3 | 10 | 1 | YES | -0,001854 | | 0,004275 | | 0,003679 | | 0,009295 | | -0,005749 | | 0,006235 | | 0,000156 | | 0,000579 | | 0,00244 | |
| TNBC_068 | 4 | GPL96 | GSM782563 | 1 | 0 | 64 | 2 | 3 | 120 | 0 | YES | -0,001913 | | 0,004938 | | 0,001578 | | 0,010361 | | -0,006122 | | 0,004235 | | 0,005606 | | 0,00079 | | 0,00266 | |
| TNBC_069 | 4 | GPL96 | GSM782564 | 1 | 0 | 33 | 1 | 3 | 74 | 1 | YES | -0,002105 | | 0,005312 | | 0,003796 | | 0,010123 | | -0,003942 | | 0,003313 | | 0,00492 | | 0,002092 | | 0,002638 | |
| TNBC_070 | 4 | GPL96 | GSM782565 | 1 | 0 | 57 | 2 | | 104 | 0 | YES | -0,004577 | | 0,004585 | | 0,004376 | | 0,010893 | | -0,006903 | | 0,004924 | | 0,004291 | | 0,00115 | | 0,001299 | |
| TNBC_071 | 4 | GPL96 | GSM782566 | 1 | 0 | 32 | 2 | 3 | 22 | 1 | YES | -0,000443 | | 0,00563 | | 0,006903 | | 0,010099 | | -0,00378 | | 0,006489 | | 0,002874 | | 0,002275 | | 0,006818 | |
| TNBC_072 | 4 | GPL96 | GSM782567 | 1 | 0 | 45 | 1 | 12 | 53 | 0 | YES | 0,00118 | | 0,003688 | | 0,000294 | | 0,008932 | | -0,004699 | | 0,002256 | | 0,00128 | | 0,00009 | | 0,00294 | |
| TNBC_073 | 4 | GPL96 | GSM782568 | 1 | 0 | 56 | 2 | 3 | 37 | 1 | YES | -0,001876 | | 0,005008 | | 0,002204 | | 0,008878 | | -0,002722 | | 0,00349 | | 0,004487 | | -0,000161 | | 0,003964 | |
| TNBC_074 | 6 | GPL96 | GSM65839 | 1 | 0 | 73 | 2 | 3 | 8 | 1 | NO | -0,001833 | | 0,006043 | | 0,004431 | | 0,008587 | | -0,00588 | | 0,003477 | | 0,000244 | | 0,00046 | | 0,006995 | |
| TNBC_075 | 6 | GPL96 | GSM65841 | 1 | 0 | 64 | 1 | 12 | 120 | 0 | NO | -0,003775 | | 0,003982 | | 0,004583 | | 0,010091 | | -0,002982 | | 0,004897 | | 0,002895 | | -0,000788 | | 0,002808 | |
| TNBC_076 | 6 | GPL96 | GSM65842 | 1 | 0 | 47 | 2 | 3 | 120 | 0 | NO | -0,010358 | | 0,002367 | | 0,005536 | | 0,008351 | | -0,01069 | | 0,006408 | | 0,006144 | | 0,00121 | | 0,003767 | |
| TNBC_077 | 6 | GPL96 | GSM65843 | 1 | 0 | 39 | 2 | 3 | 84 | 1 | NO | -0,002436 | | 0,003985 | | 0,005392 | | 0,007713 | | -0,004678 | | 0,004669 | | 0,003392 | | 0,000295 | | 0,003458 | |
| TNBC_078 | 6 | GPL96 | GSM65845 | 1 | 0 | 32 | 2 | 3 | 120 | 0 | NO | -0,013936 | | 0,004441 | | 0,007172 | | 0,011528 | | -0,002691 | | 0,007191 | | 0,002758 | | 0,001297 | | 0,005304 | |

| | | | | | | | | | | | | | | | | | | | | |
|---|---|---|---|---|---|---|---|---|---|---|---|---|---|---|---|---|---|---|---|---|
| TNBC_079 | 6 | GPL96 | GSM65852 | 1 | 0 | 53 | 2 | | 42 | 1 | NO | -0,00992 | 0,006736 | 0,012103 | 0,010366 | -0,003976 | 0,006856 | 0,00652 | 0,001025 | 0,008377 |
| TNBC_080 | 6 | GPL96 | GSM65861 | 1 | 0 | 71 | 1 | 12 | 32 | 1 | NO | -0,004172 | 0,002338 | 0,003221 | 0,008448 | -0,005856 | 0,001294 | 0,001991 | -0,000487 | 0,004506 |
| TNBC_081 | 6 | GPL96 | GSM65865 | 1 | 0 | 43 | 2 | 3 | 120 | 0 | NO | -0,007031 | 0,002045 | -0,005918 | 0,006841 | -0,007284 | -0,006007 | 0,005259 | -0,007696 | 0,005449 |
| TNBC_082 | 6 | GPL96 | GSM65876 | 1 | 0 | 52 | 1 | 12 | 63 | 0 | NO | -0,008284 | 0,002122 | 0,000571 | 0,007865 | -0,003856 | 0,000201 | 0,004608 | -0,002235 | -0,001998 |
| TNBC_083 | 6 | GPL96 | GSM65878 | 1 | 0 | 38 | 1 | 12 | 9 | 1 | NO | -0,009007 | 0,004214 | -0,000128 | 0,009513 | -0,001339 | 0,001978 | 0,006161 | -0,001548 | 0,004342 |
| TNBC_084 | 6 | GPL96 | GSM65879 | 1 | 0 | 47 | 1 | | 42 | 0 | NO | -0,013014 | 0,00483 | 0,000715 | 0,008548 | -0,00325 | -0,001215 | 0,007566 | -0,004362 | 0,003664 |
| TNBC_085 | 6 | GPL96 | GSM150795 | 1 | 0 | 51 | 1 | | | | NO | -0,006167 | 0,003953 | -0,001564 | 0,008535 | -0,00532 | -0,007194 | 0,003436 | -0,004532 | 0,003766 |
| TNBC_086 | 6 | GPL96 | GSM150797 | 1 | 0 | 29 | 2 | 3 | | | NO | -0,008454 | 0,002065 | 0,000494 | 0,009653 | -0,003045 | -0,001833 | 0,006989 | -0,005022 | 0,001803 |
| TNBC_087 | 7 | GPL96 & GPL97 | GSM107076 | 1 | | | | 3 | 15 | 1 | NO | -0,005358 | 0,005688 | 0,006883 | 0,007789 | 0,000041 | 0 | 0,004357 | 0,00239 | 0,0068 |
| TNBC_088 | 7 | GPL96 & GPL97 | GSM107084 | 1 | | | | 3 | 13 | 1 | | 0,003102 | 0,004616 | 0,008323 | 0,008134 | -0,006507 | 0,003859 | 0,003027 | 0,001173 | 0,006318 |
| TNBC_089 | 7 | GPL96 & GPL97 | GSM107094 | 1 | | | | | 49 | 1 | NO | -0,001882 | 0,005442 | 0,003887 | 0,008429 | 0,000777 | 0,002005 | 0,004364 | 0,000532 | 0,002946 |
| TNBC_090 | 7 | GPL96 & GPL97 | GSM107114 | 1 | | | | 3 | 11 | 1 | NO | -0,002671 | 0,005227 | 0,004921 | 0,007745 | -0,004006 | -0,001297 | 0,005601 | -0,000801 | 0,007287 |
| TNBC_091 | 7 | GPL96 & GPL97 | GSM107117 | 1 | | | | 3 | 76 | 0 | NO | -0,006871 | 0,005282 | 0,007129 | 0,007387 | -0,005282 | 0,001976 | 0,005865 | 0,002061 | 0,006399 |
| TNBC_092 | 7 | GPL96 & GPL97 | GSM107120 | 1 | | | | 3 | 90 | 0 | NO | -0,002546 | 0,004065 | 0,005583 | 0,011211 | -0,005003 | 0,003127 | 0,002718 | 0,002769 | 0,001876 |
| TNBC_093 | 7 | GPL96 & GPL97 | GSM107124 | 1 | | | | 3 | 98 | 0 | NO | -0,008804 | 0,003815 | 0,003563 | 0,009181 | -0,0013 | -0,006037 | 0,005213 | -0,001593 | 0,003731 |
| TNBC_094 | 7 | GPL96 & GPL97 | GSM107128 | 1 | | | | 12 | 92 | 0 | NO | -0,011896 | 0,004064 | 0,011526 | 0,010972 | 0,002069 | 0,005541 | 0,005228 | 0,000665 | 0,007019 |
| TNBC_095 | 7 | GPL96 & GPL97 | GSM107131 | 1 | | | | 3 | 76 | 0 | NO | -0,002147 | 0,001923 | -0,002594 | 0,007289 | -0,005456 | -0,012029 | 0,00213 | -0,001655 | 0,005545 |
| TNBC_096 | 7 | GPL96 & GPL97 | GSM107143 | 1 | | | | 12 | 89 | 0 | NO | -0,014151 | 0,003508 | 0,004838 | 0,008144 | -0,00383 | 0,000887 | 0,003714 | -0,000323 | 0,000847 |
| TNBC_097 | 7 | GPL96 & GPL97 | GSM107144 | 1 | | | | 3 | 79 | 0 | NO | -0,005839 | 0,005319 | 0,008318 | 0,009678 | -0,002479 | 0,004359 | 0,003901 | 0,005439 | 0,003159 |
| TNBC_098 | 7 | GPL96 & GPL97 | GSM107148 | 1 | | | | 12 | 73 | 0 | NO | -0,009663 | 0,002617 | -0,001651 | 0,008978 | -0,004751 | -0,001812 | 0,004332 | -0,003221 | 0,002496 |
| TNBC_099 | 7 | GPL96 & GPL97 | GSM107150 | 1 | | | | 12 | 75 | 1 | NO | -0,010101 | 0,010506 | 0,007139 | 0,009857 | -0,000771 | 0,001704 | 0,004402 | 0,001055 | 0,003123 |
| TNBC_100 | 7 | GPL96 & GPL97 | GSM107153 | 1 | | | | 3 | 87 | 0 | NO | -0,013718 | 0,005313 | 0,008207 | 0,00896 | -0,005114 | 0,00004 | 0,007071 | 0,000159 | 0,004202 |
| TNBC_101 | 7 | GPL96 & GPL97 | GSM107157 | 1 | | | | 12 | 100 | 0 | NO | -0,00548 | 0,00452 | -0,0014 | 0,00744 | -0,00388 | -0,00616 | 0,000426 | -0,00228 | 0,00476 |
| TNBC_102 | 7 | GPL96 & GPL97 | GSM107167 | 1 | | | | 3 | 98 | 0 | NO | -0,00453 | 0,005865 | 0,004247 | 0,010435 | -0,009424 | 0,000688 | 0,006582 | 0,000243 | 0,002386 |
| TNBC_103 | 7 | GPL96 & GPL97 | GSM107181 | 1 | | | | 3 | 13 | 1 | | -0,006794 | 0,005862 | 0,00691 | 0,011724 | -0,001398 | 0,005513 | 0,007134 | 0,004076 | 0,005047 |
| TNBC_104 | 7 | GPL96 & GPL97 | GSM107201 | 1 | | | | 12 | 95 | 0 | NO | -0,002982 | 0,003421 | -0,003333 | 0,007938 | -0,008377 | -0,007149 | 0,004428 | -0,003684 | 0,001886 |
| TNBC_105 | 7 | GPL96 & GPL97 | GSM107204 | 1 | | | | 3 | 82 | 0 | NO | -0,006922 | 0,005444 | 0,002195 | 0,008779 | -0,001182 | 0,000802 | 0,008293 | 0,001984 | 0,003461 |
| TNBC_106 | 7 | GPL96 & GPL97 | GSM107206 | 1 | | | | 3 | 17 | 1 | NO | -0,007609 | 0,005315 | 0,006959 | 0,007839 | -0,010592 | 0,001835 | 0,00375 | 0,00195 | 0,003556 |
| TNBC_107 | 7 | GPL96 & GPL97 | GSM107210 | 1 | | | | 3 | 77 | 1 | NO | 0,00094 | 0,00481 | 0,007103 | 0,009095 | -0,007779 | 0,008042 | 0,005111 | 0,002631 | 0,003119 |
| TNBC_108 | 7 | GPL96 & GPL97 | GSM107211 | 1 | | | | 3 | 86 | 0 | | -0,004934 | 0,006368 | 0,008266 | 0,010838 | -0,004133 | 0,006284 | 0,005679 | 0,002488 | 0,007338 |
| TNBC_109 | 7 | GPL96 & GPL97 | GSM107217 | 1 | | | | | 18 | 1 | NO | -0,006763 | 0,004457 | 0,007193 | 0,011611 | -0,012002 | 0,002307 | 0,003023 | 0,00215 | 0,006177 |
| TNBC_110 | 7 | GPL96 & GPL97 | GSM107224 | 1 | | | | 3 | 72 | 0 | NO | -0,003917 | 0,004406 | 0,002893 | 0,009569 | -0,002492 | 0,003605 | 0,003052 | 0,004273 | 0,004095 |
| TNBC_111 | 7 | GPL96 & GPL97 | GSM107225 | 1 | | | | 3 | 19 | 1 | NO | -0,004238 | 0,003714 | 0,005156 | 0,007996 | -0,005505 | 0,004326 | 0,002694 | 0,003714 | 0,001966 |
| TNBC_112 | 9 | GPL570 | GSM38092 | 1 | | 55 | | | | | | 0,003142 | 0,005399 | 0,008667 | 0,009933 | 0,000612 | 0,001371 | 0,00273 | 0,001687 | 0,006116 |
| TNBC_113 | 9 | GPL570 | GSM46933 | 1 | | 65 | | | | | | 0,001155 | 0,007498 | 0,008674 | 0,010501 | 0,001092 | 0,008191 | 0,003536 | 0,005608 | 0,008779 |
| TNBC_114 | 9 | GPL570 | GSM46938 | 1 | | 65 | | | | | | 0,001622 | 0,005624 | 0,008868 | 0,009993 | 0,000671 | 0,004737 | 0,00388 | 0,00292 | 0,006662 |
| TNBC_115 | 9 | GPL570 | GSM46952 | 1 | | 45 | | | | | | -0,001716 | 0,005983 | 0,010229 | 0,010734 | -0,003322 | 0,008403 | 0,003868 | 0,004773 | 0,008887 |
| TNBC_116 | 9 | GPL570 | GSM46968 | 1 | 0 | 45 | 2 | 3 | | | | 0,001264 | 0,007803 | 0,011416 | 0,012103 | 0,000466 | 0,00951 | 0,002733 | 0,004611 | 0,008778 |
| TNBC_117 | 9 | GPL570 | GSM53031 | 1 | | 75 | | | | | | -0,005877 | 0,004122 | 0,006408 | 0,00835 | -0,000931 | -0,001808 | 0,002523 | 0,001569 | 0,005292 |
| TNBC_118 | 9 | GPL570 | GSM53033 | 1 | | 65 | | | | | | 0,002442 | 0,006966 | 0,009923 | 0,010662 | 0,00569 | 0,007437 | 0,006854 | 0,004502 | 0,009139 |
| TNBC_119 | 9 | GPL570 | GSM53172 | 1 | 1 | 55 | 2 | 3 | | | | -0,003082 | 0,004428 | -0,000658 | 0,007121 | 0,001436 | -0,011669 | 0,000532 | 0,000957 | 0,005535 |
| TNBC_120 | 9 | GPL570 | GSM53187 | 1 | | 65 | | | | | | 0,003892 | 0,006127 | 0,011827 | 0,016874 | 0,000778 | 0,005097 | 0,000677 | 0,00236 | 0,006654 |
| TNBC_121 | 9 | GPL570 | GSM89023 | 1 | 1 | 45 | 2 | 3 | | | | 0,003518 | 0,005277 | 0,01 | 0,012819 | 0,00106 | 0,00588 | 0,002662 | 0,006843 | 0,006096 |
| TNBC_122 | 9 | GPL570 | GSM102506 | 1 | 1 | 35 | 1 | 3 | | | | -0,000953 | 0,003245 | 0,012235 | 0,009453 | 0,002241 | 0,010251 | 0,004101 | 0,002009 | 0,00613 |
| TNBC_123 | 9 | GPL570 | GSM102525 | 1 | 0 | 55 | 2 | 3 | | | | -0,000647 | 0,004992 | 0,014977 | 0,011744 | 0,001785 | 0,007424 | 0,005322 | 0,003673 | 0,011175 |
| TNBC_124 | 9 | GPL570 | GSM102526 | 1 | 0 | 45 | 1 | 12 | | | | 0,000046 | 0,003587 | 0,009187 | 0,009928 | 0,000023 | 0,002708 | 0,005011 | 0,002268 | 0,005809 |
| TNBC_125 | 9 | GPL570 | GSM102545 | 1 | 1 | 65 | 1 | 12 | | | | -0,001079 | 0,003622 | 0,002183 | 0,007911 | -0,001259 | -0,010429 | 0,004013 | -0,008271 | 0,005112 |
| TNBC_126 | 9 | GPL570 | GSM102564 | 1 | 0 | 45 | 2 | 3 | | | | 0,000648 | 0,005737 | 0,010897 | 0,009697 | -0,000072 | 0,0048 | 0,004017 | 0,002328 | 0,007729 |
| TNBC_127 | 9 | GPL570 | GSM117581 | 1 | 1 | 45 | 1 | 3 | | | | 0,000529 | 0,005773 | 0,011748 | 0,011269 | 0,002925 | 0,006908 | 0,003239 | 0,005824 | 0,009253 |
| TNBC_128 | 9 | GPL570 | GSM117661 | 1 | 1 | 55 | 2 | 3 | | | | -0,00035 | 0,005271 | 0,005809 | 0,011591 | -0,007315 | 0,002555 | 0,001943 | 0,000995 | 0,007692 |
| TNBC_129 | 9 | GPL570 | GSM117692 | 1 | 0 | 65 | 2 | 3 | | | | -0,002825 | 0,005107 | 0,013229 | 0,011355 | 0,002227 | 0,005677 | 0,003499 | 0,004862 | 0,007715 |
| TNBC_130 | 9 | GPL570 | GSM138011 | 1 | | 85 | | | | | | 0,0003 | 0,008912 | 0,012214 | 0,012237 | 0,00374 | 0,009143 | 0,003382 | 0,005102 | 0,00912 |
| TNBC_131 | 9 | GPL570 | GSM152698 | 1 | 1 | 75 | 2 | 3 | | | | -0,007471 | 0,004678 | 0,013247 | 0,011791 | -0,001719 | 0,006755 | 0,006303 | 0,005561 | 0,007853 |
| TNBC_132 | 9 | GPL570 | GSM179854 | 1 | | | | | | | | 0,002974 | 0,006064 | 0,008875 | 0,01013 | -0,000767 | 0,002463 | 0,001922 | 0,004763 | 0,007853 |
| TNBC_133 | 9 | GPL570 | GSM179885 | 1 | | | | | | | | 0,001527 | 0,005438 | 0,00997 | 0,009803 | 0,008563 | 0,006822 | 0,00148 | 0,002886 | 0,007036 |
| TNBC_134 | 12 | GPL96 | GSM50035 | 1 | 1 | 61 | 2 | | 22 | 1 | YES | -0,002653 | 0,006286 | 0,006343 | 0,010957 | -0,012168 | 0,004613 | 0,003857 | 0,00346 | 0,004267 |
| TNBC_135 | 12 | GPL96 | GSM50040 | 1 | 0 | 57 | 2 | | 91 | 0 | YES | -0,00166 | 0,006722 | 0,005809 | 0,011785 | -0,001494 | 0,006335 | 0,004388 | 0,004316 | 0,00675 |
| TNBC_136 | 12 | GPL96 | GSM50041 | 1 | 0 | 73 | 2 | | | | | -0,002765 | 0,004147 | 0,005558 | 0,008956 | -0,007027 | 0,004665 | 0,004885 | 0,003024 | 0,005817 |
| TNBC_137 | 12 | GPL96 | GSM50045 | 1 | 0 | 58 | 2 | | 120 | 0 | YES | -0,001914 | 0,005563 | 0,003619 | 0,009331 | -0,004636 | 0,005264 | 0,005143 | 0,003469 | 0,003499 |
| TNBC_138 | 12 | GPL96 | GSM50046 | 1 | 0 | 32 | 1 | | 96 | 0 | YES | -0,001614 | 0,004103 | 0,005581 | 0,008453 | -0,008234 | 0,007796 | 0,0052 | 0,002818 | 0,006565 |
| TNBC_139 | 12 | GPL96 | GSM50048 | 1 | 0 | 49 | 1 | | 73 | 0 | YES | 0,000028 | 0,003264 | 0,004559 | 0,007344 | -0,00484 | 0,005093 | 0,010439 | 0,001942 | 0,00363 |
| TNBC_140 | 12 | GPL96 | GSM50050 | 1 | 0 | 63 | 1 | | | | | -0,002687 | 0,004044 | 0,001413 | 0,009723 | -0,003379 | 0,004349 | 0,003673 | 0,003878 | 0,005097 |
| TNBC_141 | 12 | GPL96 | GSM50052 | 1 | 0 | 47 | 2 | | 55 | 0 | YES | -0,002669 | 0,004065 | 0,002265 | 0,008968 | -0,007603 | 0,006703 | -0,006951 | 0,001831 | 0,00031 |
| TNBC_142 | 12 | GPL96 | GSM50055 | 1 | 0 | 33 | 2 | | | | | -0,004152 | 0,003344 | 0,004069 | 0,008165 | -0,005629 | 0,002452 | 0,002184 | 0,001589 | 0,003233 |
| TNBC_143 | 12 | GPL96 | GSM50057 | 1 | 0 | 46 | 2 | | | | | 0,000501 | 0,004255 | 0,006452 | 0,007203 | -0,008622 | 0,004033 | 0,000079 | 0,000083 | 0,006592 |
| TNBC_144 | 12 | GPL96 | GSM50058 | 1 | 0 | 51 | 2 | | | | | -0,003596 | 0,002564 | -0,000118 | 0,009639 | -0,009815 | 0,00283 | 0,00081 | 0,001209 | 0,000884 |
| TNBC_145 | 12 | GPL96 | GSM50060 | 1 | 0 | 52 | 2 | | 42 | 1 | YES | 0,001366 | 0,003635 | 0,007351 | 0,009455 | -0,00235 | 0,001175 | -0,000474 | 0,00123 | 0,003225 |
| TNBC_146 | 12 | GPL96 | GSM50061 | 1 | 0 | 61 | 2 | | 48 | 1 | YES | -0,003908 | 0,003362 | 0,000259 | 0,008592 | -0,011523 | 0,000661 | -0,000016 | 0,00069 | -0,000316 |
| TNBC_147 | 12 | GPL96 | GSM50063 | 1 | 0 | 49 | 2 | | 37 | 1 | YES | -0,005149 | 0,00323 | 0,001127 | 0,008135 | -0,007831 | 0,000884 | 0,003816 | 0,001554 | 0,003169 |
| TNBC_148 | 12 | GPL96 | GSM50064 | 1 | 0 | 70 | 2 | | 11 | 1 | YES | -0,004504 | 0,003493 | 0,000375 | 0,00791 | -0,005918 | -0,001992 | 0,004398 | -0,001501 | 0,005687 |
| TNBC_149 | 12 | GPL96 | GSM50066 | 1 | 0 | 80 | 2 | | 15 | 1 | YES | -0,008191 | 0,003249 | 0,00261 | 0,008247 | -0,015383 | 0,000916 | 0,003511 | 0,000472 | 0,000139 |
| TNBC_150 | 12 | GPL96 | GSM50067 | 1 | 0 | 78 | 2 | | 9 | 1 | | -0,006233 | 0,004998 | 0,007359 | 0,008595 | -0,007469 | 0,010243 | 0,005152 | 0,004723 | 0,007195 |
| TNBC_151 | 12 | GPL96 | GSM50069 | 1 | 0 | 73 | 2 | | 46 | 1 | YES | -0,002722 | 0,004114 | 0,002359 | 0,00974 | -0,008348 | 0,002178 | 0,001442 | 0,000302 | 0,00239 |
| TNBC_152 | 12 | GPL96 | GSM50070 | 1 | 1 | 58 | 2 | | 40 | 1 | YES | -0,00683 | 0,005498 | 0,006037 | 0,006632 | -0,001332 | 0,003146 | 0,005276 | 0,001332 | 0,007284 |
| TNBC_153 | 12 | GPL96 | GSM50071 | 1 | 1 | 78 | 2 | | 70 | 0 | NO | -0,003435 | 0,002438 | 0,002687 | 0,006538 | -0,007868 | -0,001884 | 0,002913 | -0,002382 | 0,003075 |
| TNBC_154 | 12 | GPL96 | GSM50073 | 1 | 1 | 50 | 2 | | 87 | 0 | YES | -0,007245 | 0,006985 | 0,00508 | 0,009121 | -0,003897 | 0,005282 | 0,003027 | 0,004301 | 0,006639 |
| TNBC_155 | 12 | GPL96 | GSM50075 | 1 | 1 | 49 | 2 | | 88 | 0 | YES | -0,007641 | 0,002556 | -0,000312 | 0,008124 | -0,010197 | 0,000227 | 0,003366 | -0,000483 | -0,000312 |
| TNBC_156 | 12 | GPL96 | GSM50079 | 1 | 1 | 40 | 2 | | 78 | 0 | YES | -0,000482 | 0,002269 | 0,006523 | 0,00848 | -0,004453 | 0,004963 | 0,003928 | 0,003063 | 0,003148 |
| TNBC_157 | 12 | GPL96 | GSM50089 | 1 | 1 | 30 | 2 | | 105 | 0 | | -0,008396 | 0,003559 | -0,004502 | 0,007848 | -0,009035 | -0,012959 | 0,005106 | -0,001886 | 0,002555 |
| TNBC_158 | 12 | GPL96 | GSM50092 | 1 | 1 | 50 | 2 | | 77 | 0 | YES | -0,007707 | 0,00423 | 0,005187 | 0,007418 | -0,003709 | 0,005071 | 0,003621 | 0,002289 | 0,0031 |

| | | | | | | | | | | | | | | | | | | | | |
|---|---|---|---|---|---|---|---|---|---|---|---|---|---|---|---|---|---|---|---|---|
| TNBC_159 | 12 | GPL96 | GSM50093 | 1 | 1 | 64 | 2 | | 94 | 0 | YES | -0,000233 | 0,004076 | 0,008151 | 0,006754 | -0,004658 | 0,007452 | 0,00248 | 0,002882 | 0,00524 |
| TNBC_160 | 12 | GPL96 | GSM50094 | 1 | 1 | 57 | 1 | | 39 | 1 | YES | -0,006716 | 0,004587 | 0,008272 | 0,007508 | -0,007098 | 0,00273 | 0,005325 | 0,003112 | 0,006307 |
| TNBC_161 | 12 | GPL96 | GSM50096 | 1 | 1 | 32 | 2 | | 17 | 1 | YES | -0,011093 | 0,0043 | 0,008886 | 0,008083 | -0,005819 | 0,005245 | 0,005811 | 0,003325 | 0,005589 |
| TNBC_162 | 12 | GPL96 | GSM50102 | 1 | 1 | 48 | 2 | | 8 | 1 | YES | -0,008367 | 0,004683 | 0,00594 | 0,008453 | -0,00594 | 0,007139 | 0,005229 | 0,00277 | 0,00494 |
| TNBC_163 | 12 | GPL96 | GSM50106 | 1 | 1 | 60 | 2 | | 19 | 1 | YES | -0,005255 | 0,004922 | 0,008008 | 0,01799 | -0,006896 | 0,001474 | 0,004604 | 0,004838 | 0,002363 |
| TNBC_164 | 12 | GPL96 | GSM50107 | 1 | 1 | 52 | 2 | | 75 | 0 | YES | -0,003992 | 0,00474 | 0,006598 | 0,009592 | -0,013001 | 0,006515 | 0,006012 | 0,004131 | 0,003936 |
| TNBC_165 | 12 | GPL96 | GSM50112 | 1 | 1 | 43 | 2 | | 14 | 1 | YES | -0,002857 | 0,006934 | 0,00713 | 0,008253 | -0,011134 | 0,004517 | 0,003171 | 0,00127 | 0,005152 |
| TNBC_166 | 12 | GPL96 | GSM50119 | 1 | 1 | 54 | 2 | | 50 | 0 | YES | -0,002475 | 0,004684 | 0,005402 | 0,008595 | -0,00181 | 0,004737 | 0,002096 | 0,004151 | 0,001517 |
| TNBC_167 | 12 | GPL96 | GSM50123 | 1 | 1 | 59 | 2 | | 47 | 0 | YES | -0,00269 | 0,004559 | 0,004474 | 0,00671 | -0,01008 | 0,005635 | 0,006169 | 0,003709 | 0,001699 |
| TNBC_168 | 12 | GPL96 | GSM50125 | 1 | 1 | 59 | 2 | | | | | -0,002038 | 0,003857 | 0,008292 | 0,009201 | -0,003581 | 0,004352 | -0,00076 | 0,002782 | 0,006942 |
| TNBC_169 | 14 | GPL570 | GSM151295 | 1 | 1 | 53 | 2 | | 6 | 1 | NO | -0,003487 | 0,003267 | 0,010537 | 0,010466 | -0,010939 | 0,003226 | 0,003346 | 0,000617 | 0,006857 |
| TNBC_170 | 14 | GPL570 | GSM151309 | 1 | 1 | 50 | 2 | 3 | 120 | 0 | NO | -0,004555 | 0,00282 | 0,00883 | 0,007752 | -0,004942 | 0,001411 | 0,00495 | 0,000294 | 0,00814 |
| TNBC_171 | 17 | GPL96 | GSM36788 | 1 | 0 | 40 | 2 | 12 | 79 | 0 | NO | -0,015837 | 0,004593 | 0,008679 | 0,008706 | -0,003792 | 0,005501 | 0,003473 | 0,002777 | 0,002644 |
| TNBC_172 | 17 | GPL96 | GSM36793 | 1 | 0 | 36 | 2 | 12 | 101 | 0 | NO | -0,008815 | 0,00362 | 0,000221 | 0,006494 | -0,004394 | 0,001686 | 0,006412 | -0,00945 | 0,004034 |
| TNBC_173 | 17 | GPL96 | GSM36795 | 1 | 0 | 71 | 2 | 12 | 88 | 0 | NO | -0,011708 | 0,004245 | 0,004216 | 0,008888 | -0,012221 | 0,001795 | 0,005224 | 0,000684 | 0,003789 |
| TNBC_174 | 17 | GPL96 | GSM36797 | 1 | 0 | 44 | 1 | 3 | 9 | 1 | NO | -0,017369 | 0,006143 | 0,007281 | 0,008949 | -0,006302 | 0,006169 | 0,009055 | 0,000927 | 0,006831 |
| TNBC_175 | 17 | GPL96 | GSM36798 | 1 | 0 | 41 | 1 | 3 | 106 | 0 | NO | -0,019436 | 0,004388 | 0,005761 | 0,009664 | -0,005842 | 0,003849 | 0,000583 | -0,002342 | 0,003365 |
| TNBC_176 | 17 | GPL96 | GSM36809 | 1 | 0 | 69 | 1 | 3 | 56 | 0 | NO | -0,012939 | 0,004647 | 0,00615 | 0,007207 | -0,001948 | 0,001419 | 0,009039 | -0,000723 | 0,004007 |
| TNBC_177 | 17 | GPL96 | GSM36822 | 1 | 0 | 73 | 2 | 3 | 50 | 0 | NO | -0,004783 | 0,004438 | 0,007626 | 0,010151 | -0,001461 | 0,002896 | 0,0062 | 0,000213 | 0,00186 |
| TNBC_178 | 17 | GPL96 | GSM36824 | 1 | 0 | 70 | 2 | 3 | 86 | 0 | NO | -0,010026 | 0,005756 | 0,007161 | 0,00818 | -0,00829 | 0,003691 | 0,005432 | 0,002038 | 0,003223 |
| TNBC_179 | 17 | GPL96 | GSM36828 | 1 | 0 | 51 | 2 | 3 | 76 | 0 | NO | -0,010403 | 0,003643 | 0,005916 | 0,00711 | -0,002914 | 0,005449 | 0,002338 | 0,00102 | 0,005129 |
| TNBC_180 | 17 | GPL96 | GSM36835 | 1 | 0 | 68 | 2 | 3 | 14 | 1 | NO | -0,017444 | 0,003794 | 0,006497 | 0,005269 | 0,001583 | 0,002839 | 0,004768 | -0,003904 | 0,00273 |
| TNBC_181 | 17 | GPL96 | GSM36846 | 1 | 0 | 36 | 1 | 3 | 103 | 0 | NO | -0,012885 | 0,003792 | 0,00459 | 0,00821 | -0,000342 | 0,003421 | 0,004614 | -0,000399 | -0,000314 |
| TNBC_182 | 17 | GPL96 | GSM36855 | 1 | 0 | 34 | 2 | 3 | 120 | 0 | NO | -0,013588 | 0,004625 | 0,007003 | 0,00682 | -0,003841 | 0,006219 | 0,004739 | 0,000287 | 0,002012 |
| TNBC_183 | 17 | GPL96 | GSM36862 | 1 | 0 | 42 | 2 | 3 | 25 | 1 | NO | -0,016633 | 0,006705 | 0,003915 | 0,009033 | -0,006909 | 0,00673 | 0,003052 | 0,00023 | 0,003327 |
| TNBC_184 | 17 | GPL96 | GSM36876 | 1 | 0 | 42 | 2 | 3 | 120 | 0 | NO | -0,015571 | 0,004273 | 0,003258 | 0,008547 | -0,010123 | 0,008333 | -0,000725 | 0,001709 | 0,004086 |
| TNBC_185 | 17 | GPL96 | GSM36883 | 1 | 0 | 73 | 2 | 12 | 88 | 0 | NO | -0,004985 | 0,002281 | 0,002338 | 0,006901 | -0,009126 | -0,000113 | 0,007212 | -0,003464 | 0,005718 |
| TNBC_186 | 17 | GPL96 | GSM36889 | 1 | 0 | 44 | 2 | 3 | 108 | 0 | NO | -0,011478 | 0,002947 | 0,003227 | 0,008644 | -0,007886 | 0,000449 | 0,005044 | 0,000477 | 0,002835 |
| TNBC_187 | 17 | GPL96 | GSM36890 | 1 | 0 | 64 | 2 | 12 | 116 | 0 | NO | -0,012861 | 0,002611 | 0,006348 | 0,008354 | -0,004947 | -0,000577 | 0,006651 | -0,002061 | 0,004699 |
| TNBC_188 | 17 | GPL96 | GSM36891 | 1 | 0 | 65 | 1 | 3 | 87 | 0 | NO | -0,010956 | 0,004372 | 0,005683 | 0,007869 | -0,000738 | 0,003388 | 0,006561 | 0,00071 | 0,00071 |
| TNBC_189 | 17 | GPL96 | GSM36901 | 1 | 0 | 29 | 2 | 3 | 101 | 0 | NO | -0,010558 | 0,005041 | 0,007173 | 0,008753 | -0,001605 | 0,009455 | 0,002448 | 0,003662 | 0,003787 |
| TNBC_190 | 17 | GPL96 | GSM36905 | 1 | 0 | 47 | 1 | 12 | 17 | 1 | NO | -0,012097 | 0,004268 | 0,002645 | 0,006467 | -0,007829 | 0,001074 | 0,003123 | 0,000393 | 0,002356 |
| TNBC_191 | 17 | GPL96 | GSM36906 | 1 | 0 | 47 | 1 | 3 | 101 | 0 | NO | -0,00986 | 0,004517 | 0,005833 | 0,008724 | -0,00795 | 0,003691 | 0,003142 | 0,001652 | 0,00271 |
| TNBC_192 | 17 | GPL96 | GSM36909 | 1 | 0 | 78 | 2 | 12 | 95 | 0 | NO | -0,015275 | 0,003314 | -0,000699 | 0,009528 | -0,005437 | -0,000311 | 0,003392 | -0,001786 | 0,002511 |
| TNBC_193 | 17 | GPL96 | GSM36912 | 1 | 0 | 53 | 2 | 3 | 97 | 0 | NO | -0,004529 | 0,004794 | 0,006516 | 0,011177 | -0,00498 | 0,006013 | 0,005358 | 0,002357 | 0,003099 |
| TNBC_194 | 17 | GPL96 | GSM36923 | 1 | 0 | 46 | 2 | 3 | 23 | 1 | NO | -0,007694 | 0,006587 | 0,005696 | 0,010231 | -0,007343 | 0,00305 | 0,004337 | 0,000837 | 0,008126 |
| TNBC_195 | 17 | GPL96 | GSM36927 | 1 | 0 | 50 | 2 | 3 | 8 | 1 | NO | -0,013111 | 0,005853 | 0,004447 | 0,010172 | -0,007284 | 0,011143 | 0,001861 | -0,001022 | 0,004447 |
| TNBC_196 | 17 | GPL96 | GSM36931 | 1 | 0 | 49 | 2 | 12 | 37 | 1 | NO | -0,01382 | 0,00383 | 0,005732 | 0,00758 | -0,006348 | 0,001152 | 0,008044 | 0,000455 | 0,002464 |
| TNBC_197 | 17 | GPL96 | GSM36935 | 1 | 0 | 50 | 1 | 3 | 114 | 0 | NO | -0,012642 | 0,00349 | 0,003243 | 0,009042 | -0,008602 | 0,001924 | 0,004335 | 0,001676 | 0,003023 |
| TNBC_198 | 17 | GPL96 | GSM36949 | 1 | 0 | 53 | 2 | 3 | 6 | 1 | NO | -0,011119 | 0,003799 | 0,006038 | 0,00956 | -0,00244 | 0,006314 | 0,004759 | 0,001233 | 0,004327 |
| TNBC_199 | 17 | GPL96 | GSM36952 | 1 | 0 | 46 | 1 | 3 | 13 | 1 | NO | -0,009405 | 0,005294 | 0,0029 | 0,0078 | -0,007969 | 0,001267 | 0,001604 | 0,000056 | 0,001436 |
| TNBC_200 | 17 | GPL96 | GSM36959 | 1 | 0 | 75 | 2 | 3 | 120 | 0 | NO | -0,014366 | 0,003527 | 0,00599 | 0,009284 | -0,004772 | 0,006846 | 0,002018 | 0,001556 | 0,004746 |
| TNBC_201 | 17 | GPL96 | GSM36960 | 1 | 0 | 48 | 1 | 12 | 51 | 1 | NO | -0,013055 | 0,003984 | 0,005276 | 0,009206 | -0,001642 | 0,008425 | 0,005119 | 0,001157 | -0,00035 |
| TNBC_202 | 17 | GPL96 | GSM36961 | 1 | 0 | 68 | 1 | 12 | 120 | 0 | NO | -0,010552 | 0,004232 | 0,004616 | 0,008326 | -0,007639 | 0,001759 | 0,006792 | 0,00272 | 0,002336 |
| TNBC_203 | 17 | GPL96 | GSM36966 | 1 | 0 | 70 | 2 | 3 | 90 | 0 | NO | -0,011739 | 0,004304 | 0,006334 | 0,008975 | -0,003375 | 0,005845 | 0,001857 | 0,001149 | 0,005625 |
| TNBC_204 | 17 | GPL96 | GSM36969 | 1 | 0 | 41 | 1 | 3 | 32 | 1 | NO | -0,008581 | 0,005704 | 0,006764 | 0,008682 | -0,002625 | 0,006259 | 0,005205 | 0,000404 | 0,006234 |
| TNBC_205 | 17 | GPL96 | GSM36977 | 1 | 0 | 72 | 2 | 3 | 120 | 0 | NO | -0,011463 | 0,003632 | 0,005812 | 0,006243 | -0,000726 | 0,007238 | 0,001634 | 0,000673 | 0,003121 |
| TNBC_206 | 17 | GPL96 | GSM36981 | 1 | 0 | 60 | 2 | 3 | 87 | 0 | NO | -0,011333 | 0,005107 | 0,008337 | 0,008467 | -0,010578 | 0,007764 | 0,002986 | -0,000156 | 0,004299 |
| TNBC_207 | 17 | GPL96 | GSM36991 | 1 | 0 | 66 | 1 | 3 | 114 | 0 | NO | -0,011745 | 0,005314 | 0,006655 | 0,008939 | -0,007474 | 0,006779 | 0,004143 | 0,001887 | 0,00519 |
| TNBC_208 | 17 | GPL96 | GSM37002 | 1 | 0 | 53 | 2 | 3 | 16 | 1 | NO | -0,012127 | 0,00578 | 0,007565 | 0,009152 | -0,00034 | 0,004448 | 0,007052 | 0,002323 | 0,004873 |
| TNBC_209 | 17 | GPL96 | GSM37017 | 1 | 0 | 41 | 2 | 3 | 108 | 0 | NO | -0,014779 | 0,003786 | 0,008199 | 0,007755 | -0,007285 | 0,005353 | 0,005907 | 0,000966 | 0,004987 |
| TNBC_210 | 17 | GPL96 | GSM37021 | 1 | 0 | 58 | 2 | 12 | 109 | 0 | NO | -0,012832 | 0,006634 | 0,008093 | 0,0115 | -0,013267 | 0,006531 | 0,001102 | 0,002254 | 0,006147 |
| TNBC_211 | 17 | GPL96 | GSM37022 | 1 | 0 | 58 | 1 | 12 | 30 | 1 | NO | -0,013942 | 0,004108 | 0,006265 | 0,009166 | -0,008473 | 0,00362 | 0,004527 | -0,000745 | 0,007908 |
| TNBC_212 | 17 | GPL96 | GSM37044 | 1 | 0 | 74 | 2 | 3 | 112 | 0 | NO | -0,011153 | 0,004105 | 0,003567 | 0,007954 | -0,003453 | -0,00133 | 0,006765 | -0,002717 | 0,007983 |
| TNBC_213 | 17 | GPL96 | GSM37045 | 1 | 0 | 43 | 1 | 3 | 120 | 0 | NO | -0,013772 | 0,004371 | 0,009532 | 0,010243 | -0,005451 | 0,006689 | 0,004424 | 0,002949 | 0,00603 |
| TNBC_214 | 17 | GPL96 | GSM37047 | 1 | 0 | 44 | 1 | 3 | 108 | 0 | NO | -0,013741 | 0,005257 | 0,006481 | 0,007928 | 0,000056 | 0,008206 | 0,001311 | 0,003004 | 0,003922 |
| TNBC_215 | 17 | GPL96 | GSM37048 | 1 | 0 | 43 | 1 | 3 | 87 | 0 | NO | -0,010162 | 0,002953 | -0,000058 | 0,007383 | -0,00275 | 0,001274 | 0,004075 | 0,000087 | 0,004198 |
| TNBC_216 | 17 | GPL96 | GSM37050 | 1 | 0 | 69 | 1 | 12 | 18 | 1 | NO | -0,013619 | 0,002779 | 0,002311 | 0,007016 | -0,002229 | 0,000303 | 0,003933 | -0,00033 | 0,000138 |
| TNBC_217 | 17 | GPL96 | GSM37051 | 1 | 0 | 43 | 2 | 3 | 18 | 1 | NO | -0,013319 | 0,001736 | 0,004422 | 0,008138 | -0,01001 | 0,001736 | 0,003365 | 0,000543 | 0,002794 |
| TNBC_218 | 17 | GPL96 | GSM120649 | 1 | 0 | 53 | 1 | 12 | 120 | 0 | NO | -0,009927 | 0,005295 | 0,004662 | 0,009496 | -0,001093 | 0,003424 | 0,00727 | 0,000317 | 0,001755 |
| TNBC_219 | 17 | GPL96 | GSM120651 | 1 | 0 | 34 | 2 | 3 | 88 | 0 | NO | -0,012538 | 0,004632 | 0,004219 | 0,008821 | -0,007257 | 0,004042 | 0,006426 | 0,001269 | 0,005664 |
| TNBC_220 | 17 | GPL96 | GSM120655 | 1 | 0 | 43 | 2 | 3 | 10 | 1 | NO | -0,010987 | 0,003368 | 0,007205 | 0,009331 | -0,004196 | 0,008034 | 0,005769 | 0,002071 | 0,005383 |
| TNBC_221 | 17 | GPL96 | GSM120656 | 1 | 0 | 38 | 2 | 3 | 22 | 1 | NO | -0,014511 | 0,004381 | 0,008652 | 0,007748 | -0,006708 | 0,008104 | 0,00473 | 0,000301 | 0,003943 |
| TNBC_222 | 17 | GPL96 | GSM120657 | 1 | 0 | 69 | 2 | 3 | 61 | 0 | NO | -0,011201 | 0,004194 | 0,008415 | 0,007808 | -0,004414 | 0,006566 | 0,004087 | 0,001076 | 0,005187 |
| TNBC_223 | 17 | GPL96 | GSM120658 | 1 | 0 | 42 | 1 | 3 | 120 | 0 | NO | -0,010049 | 0,004664 | 0,006477 | 0,009036 | -0,009702 | 0,00725 | 0,004927 | 0,002772 | 0,002399 |
| TNBC_224 | 17 | GPL96 | GSM120660 | 1 | 0 | 38 | 1 | 3 | 116 | 0 | NO | -0,010935 | 0,00601 | 0,008292 | 0,008737 | -0,002838 | 0,007206 | 0,003048 | 0,00256 | 0,004869 |
| TNBC_225 | 17 | GPL96 | GSM120663 | 1 | 0 | 49 | 2 | 3 | 29 | 1 | NO | -0,009382 | 0,003918 | 0,000276 | 0,007975 | -0,008885 | 0,003587 | -0,003267 | 0,001297 | 0,001932 |
| TNBC_226 | 17 | GPL96 | GSM120666 | 1 | 0 | 34 | 2 | 12 | 87 | 0 | NO | -0,010697 | 0,003443 | 0,003895 | 0,006887 | -0,00525 | 0,004008 | 0,002708 | -0,001637 | 0,002258 |
| TNBC_227 | 17 | GPL96 | GSM120668 | 1 | 0 | 35 | 1 | 3 | 7 | 1 | NO | -0,008653 | 0,004021 | 0,00771 | 0,009762 | -0,001608 | 0,008708 | 0,004354 | 0,002024 | 0,003245 |
| TNBC_228 | 17 | GPL96 | GSM120671 | 1 | 0 | 32 | 2 | 12 | 24 | 1 | NO | -0,009566 | 0,00462 | 0,005136 | 0,010136 | -0,004783 | 0,004457 | 0,00367 | 0,001848 | 0,001984 |
| TNBC_229 | 17 | GPL96 | GSM120672 | 1 | 0 | 47 | 2 | 12 | 120 | 0 | NO | -0,013445 | 0,003998 | 0,002382 | 0,007941 | -0,003478 | 0,003259 | 0,003831 | 0,000822 | 0,001506 |
| TNBC_230 | 17 | GPL96 | GSM120674 | 1 | 0 | 63 | 2 | 3 | 86 | 0 | NO | -0,009532 | 0,005001 | 0,001437 | 0,008068 | -0,011024 | 0,006134 | 0,001953 | 0,001243 | 0,006327 |
| TNBC_231 | 17 | GPL96 | GSM120675 | 1 | 0 | 44 | 1 | 3 | 119 | 0 | NO | -0,009999 | 0,005673 | 0,008398 | 0,00924 | -0,003258 | 0,008314 | 0,00503 | 0,002415 | 0,00396 |
| TNBC_232 | 17 | GPL96 | GSM120677 | 1 | 0 | 82 | 2 | 3 | 68 | 0 | NO | -0,013321 | 0,004886 | 0,008025 | 0,008735 | -0,004095 | 0,006661 | 0,002701 | 0,002184 | 0,005651 |
| TNBC_233 | 17 | GPL96 | GSM120679 | 1 | 0 | 74 | 2 | 3 | 42 | 1 | NO | -0,010236 | 0,005238 | 0,006575 | 0,009595 | -0,002967 | 0,003341 | 0,003598 | 0,003234 | 0,005532 |
| TNBC_234 | 17 | GPL96 | GSM120682 | 1 | 0 | 58 | 1 | 3 | 76 | 0 | NO | -0,010938 | 0,005004 | 0,006399 | 0,006809 | -0,003008 | 0,005551 | 0,001809 | -0,000984 | 0,002379 |
| TNBC_235 | 17 | GPL96 | GSM120683 | 1 | 0 | 63 | 1 | 3 | 120 | 0 | NO | -0,00555 | 0,004445 | 0,009547 | 0,008522 | -0,007811 | 0,002367 | 0,00298 | 0,001078 | 0,006733 |
| TNBC_236 | 17 | GPL96 | GSM120684 | 1 | 0 | 44 | 2 | 12 | 120 | 0 | NO | -0,010519 | 0,006986 | 0,007092 | 0,011395 | -0,00348 | 0,008314 | 0,001798 | 0,000239 | 0,006827 |
| TNBC_237 | 17 | GPL96 | GSM120685 | 1 | 0 | 57 | 1 | 3 | 88 | 0 | NO | -0,00572 | 0,00478 | 0,007792 | 0,008593 | -0,001161 | 0,009229 | 0,002643 | 0,002597 | 0,001713 |
| TNBC_238 | 17 | GPL96 | GSM120686 | 1 | 0 | 66 | 1 | 3 | 81 | 0 | NO | -0,011654 | 0,00609 | 0,004844 | 0,007059 | -0,001107 | 0,001882 | -0,001505 | -0,000747 | 0,006976 |

| | | | | | | | | | | | | | | | | | | | | |
|---|---|---|---|---|---|---|---|---|---|---|---|---|---|---|---|---|---|---|---|---|
| TNBC_239 | 17 | GPL96 | GSM120687 | 1 | 0 | 41 | 2 | 12 | 9 | 1 | NO | -0,011493 | 0,007451 | 0,008166 | 0,009266 | -0,003107 | 0,008111 | 0,005163 | 0,003877 | 0,006571 |
| TNBC_240 | 17 | GPL96 | GSM120688 | 1 | 0 | 44 | 2 | 3 | 120 | 0 | NO | -0,01384 | 0,00738 | 0,009894 | 0,013165 | -0,002757 | 0,009245 | 0,005295 | 0,002136 | 0,00392 |
| TNBC_241 | 17 | GPL96 | GSM120690 | 1 | 0 | 58 | 2 | 3 | 88 | 0 | NO | -0,010365 | 0,002101 | 0,005197 | 0,007213 | 0,000227 | 0,003976 | 0,001911 | 0,001306 | 0,003067 |
| TNBC_242 | 17 | GPL96 | GSM120691 | 1 | 0 | 61 | 2 | 12 | 103 | 0 | NO | -0,010725 | 0,004581 | 0,004087 | 0,009381 | -0,007434 | 0,008531 | 0,000529 | 0,00384 | -0,000713 |
| TNBC_243 | 17 | GPL96 | GSM120692 | 1 | 0 | 39 | 2 | 3 | 89 | 0 | NO | -0,01228 | 0,005775 | 0,008663 | 0,010429 | -0,002944 | 0,006392 | 0,002917 | 0,002047 | 0,005747 |
| TNBC_244 | 17 | GPL96 | GSM120695 | 1 | 0 | 38 | 2 | 12 | 90 | 0 | NO | -0,013116 | 0,00555 | 0,005798 | 0,009636 | -0,003948 | 0,00809 | 0,003314 | 0,002154 | 0,003507 |
| TNBC_245 | 17 | GPL96 | GSM120696 | 1 | 0 | 33 | 2 | 3 | 120 | 0 | NO | -0,01001 | 0,004393 | 0,007035 | 0,011011 | -0,007702 | 0,008731 | 0,004131 | 0,003364 | 0,003726 |
| TNBC_246 | 17 | GPL96 | GSM120698 | 1 | 0 | 57 | 2 | 12 | 61 | 0 | NO | -0,012765 | 0,004801 | 0,004882 | 0,009983 | -0,005537 | 0,012247 | 0,000745 | 0,001909 | 0,002919 |
| TNBC_247 | 17 | GPL96 | GSM120699 | 1 | 0 | 40 | 1 | 3 | 80 | 0 | NO | -0,009933 | 0,003919 | 0,005007 | 0,010178 | -0,007675 | 0,01143 | 0,001511 | 0,004545 | 0,004327 |
| TNBC_248 | 17 | GPL96 | GSM120701 | 1 | 0 | 50 | 2 | 3 | 69 | 0 | NO | -0,0015 | 0,004732 | 0,006233 | 0,009032 | -0,007647 | 0,006262 | 0,005959 | 0,001731 | 0,005713 |
| TNBC_249 | 17 | GPL96 | GSM120702 | 1 | 0 | 39 | 2 | 3 | 4 | 1 | NO | -0,008936 | 0,005395 | 0,007615 | 0,010622 | -0,001433 | 0,008739 | 0,004708 | 0,00326 | 0,002754 |
| TNBC_250 | 17 | GPL96 | GSM120703 | 1 | 0 | 59 | 2 | 12 | 91 | 0 | NO | -0,008527 | 0,005065 | 0,009443 | 0,009243 | -0,007726 | 0,002718 | 0,002452 | 0,002862 | 0,004321 |
| TNBC_251 | 17 | GPL96 | GSM120705 | 1 | 0 | 73 | 2 | 3 | 83 | 0 | NO | -0,009845 | 0,004196 | 0,006911 | 0,012094 | -0,007377 | 0,008803 | 0,004113 | 0,002605 | 0,007569 |
| TNBC_252 | 17 | GPL96 | GSM120706 | 1 | 0 | 46 | 1 | 3 | 102 | 0 | NO | -0,004931 | 0,004536 | 0,003944 | 0,008903 | -0,009692 | 0,006818 | 0,003863 | 0,003409 | 0,003663 |
| TNBC_253 | 20 | GPL96 | GSM782569 | 2 | 0 | 65 | 2 | 12 | 11 | 1 | | -0,001842 | 0,003903 | 0,000165 | 0,008796 | -0,008301 | -0,001044 | -0,003825 | -0,003188 | 0,002556 |
| TNBC_254 | 20 | GPL96 | GSM782570 | 2 | 1 | 32 | 2 | 3 | 39 | 1 | | -0,008208 | 0,006809 | 0,008262 | 0,011115 | -0,004091 | 0,007885 | 0,004442 | 0,002315 | 0,008101 |
| TNBC_255 | 20 | GPL96 | GSM782571 | 2 | 0 | 56 | 2 | 3 | 45 | 0 | | -0,007417 | 0,00619 | 0,007313 | 0,012301 | -0,008409 | 0,005589 | 0,003695 | 0,002037 | 0,008618 |
| TNBC_256 | 20 | GPL96 | GSM782572 | 2 | 0 | 55 | 2 | 12 | 45 | 0 | | -0,00165 | 0,004152 | 0,007338 | 0,010068 | -0,004977 | 0,002873 | 0,006693 | -0,000341 | 0,003527 |
| TNBC_257 | 20 | GPL96 | GSM782573 | 2 | 1 | 44 | 2 | 12 | 23 | 1 | | -0,002231 | 0,004377 | 0,006354 | 0,009008 | -0,004885 | 0,007794 | 0,003647 | 0,003982 | 0,006184 |
| TNBC_258 | 20 | GPL96 | GSM782574 | 2 | 0 | 67 | 2 | 12 | 34 | 1 | | -0,002575 | 0,004011 | 0,00374 | 0,010137 | -0,009838 | 0,001355 | -0,00056 | -0,000271 | 0,00187 |
| TNBC_259 | 20 | GPL96 | GSM782575 | 2 | 1 | 46 | 2 | 3 | 15 | 1 | | -0,006389 | 0,005297 | -0,000289 | 0,011397 | -0,004334 | -0,007448 | 0,00328 | -0,002857 | 0,002279 |
| TNBC_260 | 20 | GPL96 | GSM782576 | 2 | 0 | 60 | 2 | 3 | 51 | 0 | | -0,001939 | 0,003508 | 0,001141 | 0,009696 | -0,001055 | 0,000114 | 0,002103 | -0,00365 | 0,000371 |
| TNBC_261 | 20 | GPL96 | GSM782577 | 2 | 1 | 63 | 2 | 12 | 44 | 0 | | -0,001593 | 0,005649 | 0,003679 | 0,009357 | -0,00055 | 0,000463 | 0,005901 | 0,000009 | 0,003853 |
| TNBC_262 | 20 | GPL96 | GSM782578 | 2 | 1 | 60 | 2 | 3 | 30 | 0 | | -0,003018 | 0,003865 | 0,003468 | 0,008126 | -0,004129 | 0,000556 | -0,001754 | -0,000026 | 0,001429 |
| TNBC_263 | 20 | GPL96 | GSM782579 | 2 | 1 | 44 | 2 | 12 | 27 | 0 | | -0,004094 | 0,002811 | 0,001228 | 0,009033 | -0,009743 | -0,001556 | 0,004389 | -0,001119 | 0,004721 |
| TNBC_264 | 20 | GPL96 | GSM782580 | 2 | 1 | 69 | 2 | 12 | 26 | 0 | | -0,001867 | 0,003544 | 0,004491 | 0,009929 | -0,008928 | 0,006223 | 0,004742 | 0,002895 | 0,001596 |
| TNBC_265 | 20 | GPL96 | GSM782581 | 2 | 1 | 50 | 2 | 3 | 25 | 0 | | 0,000752 | 0,006152 | 0,0054 | 0,010611 | -0,003251 | 0,007226 | 0,004209 | 0,002445 | 0,002257 |
| TNBC_266 | 20 | GPL96 | GSM782582 | 2 | 1 | 44 | 2 | 3 | 10 | 1 | | -0,001051 | 0,005655 | 0,009459 | 0,010585 | -0,00553 | 0,007657 | 0,005988 | 0,00548 | 0,006056 |
| TNBC_267 | 20 | GPL96 | GSM782583 | 2 | | | | | | | | 0,000407 | 0,004987 | 0,007989 | 0,009363 | -0,004071 | 0,005597 | 0,004222 | 0,002442 | 0,008472 |
| TNBC_268 | 20 | GPL96 | GSM782584 | 2 | 0 | 39 | 2 | 3 | 23 | 0 | | -0,001695 | 0,005377 | 0,005881 | 0,009987 | -0,002861 | 0,010278 | 0,004273 | 0,004265 | 0,005324 |
| TNBC_269 | 20 | GPL96 | GSM782585 | 2 | 0 | 41 | 2 | 3 | 23 | 0 | | -0,00238 | 0,006365 | 0,005776 | 0,011446 | -0,005482 | 0,006418 | 0,003321 | 0,002621 | 0,006552 |
| TNBC_270 | 20 | GPL96 | GSM782586 | 2 | 1 | 44 | 2 | 3 | 23 | 0 | | -0,001861 | 0,004262 | 0,006905 | 0,010358 | -0,005611 | 0,004046 | 0,004197 | 0,003291 | 0,00348 |
| TNBC_271 | 20 | GPL96 | GSM782587 | 2 | 1 | 42 | 2 | 12 | 24 | 0 | | -0,001545 | 0,005341 | 0,007409 | 0,009739 | -0,003482 | 0,008377 | 0,006763 | 0,003979 | 0,004267 |
| TNBC_272 | 21 | GPL96 | GSM505351 | 3 | 1 | 50 | 2 | 3 | | | | -0,007294 | 0,004713 | 0,005593 | 0,01061 | -0,002088 | 0,004925 | 0,007875 | 0,003347 | 0,005895 |
| TNBC_273 | 21 | GPL96 | GSM505331 | 3 | 0 | 75 | 2 | 3 | | | | -0,001028 | 0,005163 | 0,006483 | 0,009025 | -0,004718 | 0,002464 | 0,006489 | -0,000215 | 0,004741 |
| TNBC_274 | 21 | GPL96 | GSM505460 | 3 | 0 | 52 | 2 | 3 | | | | -0,000194 | 0,00295 | 0,006699 | 0,00949 | 0,000147 | 0,002665 | 0,008029 | 0,002367 | 0,003157 |
| TNBC_275 | 21 | GPL96 | GSM505345 | 3 | 0 | 65 | 1 | 3 | | | | -0,006452 | 0,005555 | 0,002732 | 0,006389 | -0,007295 | 0,001818 | 0,005469 | 0,001235 | -0,000097 |
| TNBC_276 | 21 | GPL96 | GSM505369 | 3 | 0 | 59 | 2 | 3 | | | | -0,004468 | 0,004151 | 0,010117 | 0,007364 | -0,001006 | 0,003561 | 0,009 | 0,002313 | 0,003716 |
| TNBC_277 | 21 | GPL96 | GSM505343 | 3 | 1 | 66 | 2 | 3 | | | | -0,011771 | 0,004412 | 0,004325 | 0,008249 | -0,001752 | 0,00245 | 0,00435 | 0,001907 | 0,00442 |
| TNBC_278 | 21 | GPL96 | GSM505372 | 3 | 0 | 38 | 2 | 3 | | | | -0,003666 | 0,004688 | 0,008471 | 0,007897 | -0,00301 | 0,006401 | 0,00101 | -0,000689 | 0,005333 |
| TNBC_279 | 21 | GPL96 | GSM505358 | 3 | 0 | 38 | 1 | 3 | | | | -0,005349 | 0,004853 | 0,011177 | 0,007368 | -0,001648 | 0,006279 | 0,004856 | 0,003804 | 0,005699 |
| TNBC_280 | 21 | GPL96 | GSM505336 | 3 | 1 | 61 | 1 | 3 | | | | -0,001809 | 0,005461 | 0,008121 | 0,010364 | 0,001991 | 0,007311 | 0,008758 | 0,003383 | 0,006312 |
| TNBC_281 | 21 | GPL96 | GSM505333 | 3 | 1 | 29 | 2 | 3 | | | | -0,010363 | 0,006201 | 0,00566 | 0,008962 | -0,005255 | 0,005708 | 0,003684 | 0,002549 | 0,006427 |
| TNBC_282 | 21 | GPL96 | GSM505350 | 3 | 1 | 67 | 2 | 12 | | | | -0,000935 | 0,005883 | 0,005198 | 0,007258 | -0,004492 | -0,001072 | 0,00368 | -0,002984 | 0,004445 |
| TNBC_283 | 21 | GPL96 | GSM505370 | 3 | 1 | 35 | 1 | 3 | | | | -0,007656 | 0,00321 | 0,00505 | 0,008285 | -0,001527 | 0,003076 | 0,006349 | 0,001771 | 0,005613 |
| TNBC_284 | 21 | GPL96 | GSM505359 | 3 | 0 | 48 | 2 | 3 | | | | -0,003354 | 0,005021 | 0,006808 | 0,011976 | -0,008787 | 0,001767 | 0,009015 | 0,001918 | 0,00534 |
| TNBC_285 | 21 | GPL96 | GSM505360 | 3 | 1 | 52 | 2 | 3 | | | | -0,000979 | 0,005091 | 0,005144 | 0,008384 | 0,000533 | 0,000956 | 0,009301 | -0,001124 | 0,004047 |
| TNBC_286 | 21 | GPL96 | GSM505357 | 3 | 1 | 44 | 2 | 3 | | | | -0,003411 | 0,006398 | 0,007 | 0,010154 | -0,002975 | 0,005504 | 0,007212 | 0,001225 | 0,005812 |
| TNBC_287 | 21 | GPL96 | GSM505421 | 3 | 1 | 60 | 2 | 3 | | | | -0,011004 | 0,003714 | 0,003996 | 0,00889 | 0,007102 | 0,007585 | 0,004131 | 0,004545 | 0,00391 |
| TNBC_288 | 21 | GPL96 | GSM505417 | 3 | 1 | 51 | 2 | 3 | | | | -0,007123 | 0,005757 | 0,007016 | 0,010938 | -0,001922 | 0,008003 | 0,000675 | 0,002665 | 0,004138 |
| TNBC_289 | 21 | GPL96 | GSM505423 | 3 | 1 | 48 | 2 | 3 | | | | -0,013755 | 0,005967 | 0,004084 | 0,008565 | -0,00766 | 0,009815 | 0,00189 | 0,007342 | 0,00712 |
| TNBC_290 | 21 | GPL96 | GSM505411 | 3 | 1 | 46 | 2 | 3 | | | | -0,003869 | 0,006785 | 0,002841 | 0,011854 | -0,013335 | 0,006835 | 0,007191 | 0,002387 | 0,001531 |
| TNBC_291 | 21 | GPL96 | GSM505413 | 3 | 1 | 51 | 2 | 12 | | | | -0,010095 | 0,004027 | -0,003073 | 0,00708 | -0,006083 | 0,004343 | 0,00421 | 0,002385 | 0,003404 |
| TNBC_292 | 21 | GPL96 | GSM505435 | 3 | 1 | 71 | 1 | 12 | | | | -0,007605 | 0,002044 | 0,001943 | 0,007312 | -0,010539 | -0,002745 | 0,005699 | 0,002101 | -0,000081 |
| TNBC_293 | 21 | GPL96 | GSM505487 | 3 | 1 | 63 | 1 | 3 | | | | -0,003228 | 0,002335 | 0,00499 | 0,008233 | -0,00069 | 0,00157 | 0,006457 | 0,002755 | 0,005789 |
| TNBC_294 | 21 | GPL96 | GSM505459 | 3 | 1 | 62 | 2 | 3 | | | | 0,000345 | 0,005177 | 0,010285 | 0,011012 | -0,005515 | 0,003644 | -0,004649 | 0,00283 | 0,003125 |
| TNBC_295 | 21 | GPL96 | GSM505467 | 3 | 1 | 46 | 2 | 12 | | | | -0,00283 | 0,001936 | 0,000892 | 0,005542 | 0,00584 | 0,003396 | 0,006961 | 0,002148 | 0,003726 |
| TNBC_296 | 21 | GPL96 | GSM505428 | 3 | 1 | 38 | 2 | 3 | | | | -0,010471 | 0,005553 | 0,008046 | 0,007225 | -0,005394 | 0,002252 | 0,004909 | 0,002927 | 0,003894 |
| TNBC_297 | 21 | GPL96 | GSM505449 | 3 | 1 | 41 | 2 | 3 | | | | 0,001458 | 0,002937 | 0,004213 | 0,007966 | -0,006897 | 0,002577 | 0,009101 | 0,002028 | 0,003859 |
| TNBC_298 | 21 | GPL96 | GSM505473 | 3 | 1 | 62 | 2 | 3 | | | | 0,001351 | 0,00426 | 0,008241 | 0,007925 | -0,001742 | 0,000419 | 0,006266 | 0,003484 | 0,005044 |
| TNBC_299 | 21 | GPL96 | GSM505462 | 3 | 1 | 50 | 2 | 12 | | | | -0,010694 | 0,002278 | 0,005926 | 0,009807 | -0,012186 | 0,008551 | 0,004964 | 0,00687 | 0,003117 |
| TNBC_300 | 21 | GPL96 | GSM505450 | 3 | 1 | 58 | 2 | 3 | | | | -0,006406 | 0,003932 | 0,004132 | 0,009724 | -0,012189 | 0,003871 | 0,003792 | 0,001509 | 0,001653 |
| TNBC_301 | 23 | GPL570 | GSM232199 | 1 | 0 | 73 | 1 | 3 | 120 | 0 | NO | -0,002239 | 0,003882 | 0,006686 | 0,010121 | -0,008681 | 0,001936 | 0,008964 | 0,000148 | 0,008316 |
| TNBC_302 | 23 | GPL570 | GSM232204 | 1 | 0 | 82 | 1 | | 120 | 0 | NO | -0,001317 | 0,004182 | 0,00647 | 0,011102 | -0,005246 | -0,000183 | 0,009212 | -0,002111 | 0,004207 |
| TNBC_303 | 24 | GPL96 | GSM26871 | 2 | 0 | | 2 | 3 | | | | -0,012641 | 0,004781 | 0,007847 | 0,009226 | -0,00148 | 0,005497 | 0,00403 | 0,000788 | 0,002767 |
| TNBC_304 | 24 | GPL96 | GSM26878 | 2 | 1 | | 2 | 12 | | | | -0,012089 | 0,003723 | 0,007163 | 0,008346 | -0,00507 | 0,001913 | 0,006057 | -0,003501 | 0,00529 |
| TNBC_305 | 24 | GPL96 | GSM26880 | 2 | 0 | | 2 | 3 | | | | -0,015734 | 0,005135 | 0,009016 | 0,010254 | -0,001803 | 0,001394 | 0,006813 | -0,001601 | 0,005794 |
| TNBC_306 | 24 | GPL96 | GSM26882 | 2 | 1 | | 2 | 3 | | | | -0,016004 | 0,004879 | 0,008635 | 0,008612 | -0,003981 | 0,005443 | 0,006907 | 0,00118 | 0,004302 |
| TNBC_307 | 24 | GPL96 | GSM26883 | 2 | 0 | | 2 | | | | | -0,009524 | 0,003806 | 0,005344 | 0,01 | -0,002174 | 0,002214 | 0,006771 | -0,00074 | 0,002742 |
| TNBC_308 | 24 | GPL96 | GSM26884 | 2 | 1 | | 2 | 12 | | | | -0,011487 | 0,004735 | 0,008326 | 0,011231 | -0,000527 | 0,005117 | 0,005775 | -0,000168 | 0,002461 |
| TNBC_309 | 24 | GPL96 | GSM26886 | 2 | 1 | | 2 | 12 | | | | -0,011261 | 0,003996 | 0,006264 | 0,009254 | -0,002144 | 0,001393 | 0,00654 | -0,002757 | 0,006159 |
| TNBC_310 | 24 | GPL96 | GSM26887 | 2 | 0 | | 2 | 12 | | | | -0,011985 | 0,00488 | 0,011039 | 0,010532 | -0,004165 | 0,003845 | 0,006426 | 0,001473 | 0,00736 |
| TNBC_311 | 24 | GPL96 | GSM26888 | 2 | 0 | | | 3 | | | | -0,014421 | 0,003963 | 0,008037 | 0,010099 | -0,003064 | 0,005483 | 0,005071 | -0,000611 | 0,000632 |
| TNBC_312 | 24 | GPL96 | GSM26889 | 2 | 1 | | 2 | 12 | | | | -0,012665 | 0,005345 | 0,009918 | 0,008846 | -0,001634 | 0,005488 | 0,008671 | 0,000073 | 0,004461 |
| TNBC_313 | 24 | GPL96 | GSM26892 | 2 | 1 | | 2 | 3 | | | | -0,01358 | 0,003215 | 0,009601 | 0,009101 | -0,006752 | 0,008006 | 0,005803 | 0,002173 | 0,004044 |
| TNBC_314 | 24 | GPL96 | GSM26893 | 2 | 1 | | 2 | 3 | | | | -0,011149 | 0,005248 | 0,00736 | 0,010801 | -0,004876 | 0,006842 | 0,006052 | 0,001965 | 0,006916 |
| TNBC_315 | 24 | GPL96 | GSM26895 | 2 | 0 | | 2 | 3 | | | | -0,010807 | 0,006101 | 0,011169 | 0,012083 | -0,004188 | 0,00707 | 0,006149 | 0,000995 | 0,006521 |
| TNBC_316 | 24 | GPL96 | GSM26898 | 2 | 1 | | 2 | 3 | | | | -0,016339 | 0,006024 | 0,011299 | 0,012179 | -0,004361 | 0,005374 | 0,004556 | 0,002095 | 0,007283 |
| TNBC_317 | 24 | GPL96 | GSM26902 | 2 | 1 | | 2 | 3 | | | | -0,013313 | 0,004341 | 0,008927 | 0,009985 | -0,002241 | 0,007114 | 0,002392 | 0,001959 | 0,006218 |
| TNBC_318 | 24 | GPL96 | GSM26903 | 2 | 0 | | 2 | | | | | -0,010458 | 0,004022 | 0,007483 | 0,008309 | -0,000814 | 0,00077 | 0,005514 | -0,001564 | 0,005991 |

| | | | | | | | | | | | | | | | | | | | | |
|---|---|---|---|---|---|---|---|---|---|---|---|---|---|---|---|---|---|---|---|---|
| TNBC_319 | 24 | GPL96 | GSM26905 | 2 | 1 | | 2 | 12 | | | | -0,013214 | 0,004481 | 0,008533 | 0,009623 | -0,007186 | 0,007315 | 0,006008 | 0,00241 | 0,004919 |
| TNBC_320 | 24 | GPL96 | GSM26906 | 2 | 1 | | 2 | 3 | | | | -0,009495 | 0,003668 | 0,0089 | 0,008308 | -0,000603 | 0,004073 | 0,005217 | -0,000621 | 0,002915 |
| TNBC_321 | 24 | GPL96 | GSM26908 | 2 | 0 | | 2 | 3 | | | | -0,012002 | 0,004826 | 0,009633 | 0,010405 | -0,003748 | 0,005184 | 0,004976 | 0,002755 | 0,005145 |
| TNBC_322 | 24 | GPL96 | GSM26910 | 2 | 1 | | 2 | 3 | | | | -0,01183 | 0,00508 | 0,007035 | 0,006506 | -0,005908 | 0,002784 | 0,004154 | -0,005058 | 0,007626 |
| TNBC_323 | 24 | GPL96 | GSM26912 | 2 | 0 | | 2 | 3 | | | | -0,014198 | 0,004529 | 0,007633 | 0,009732 | -0,007952 | 0,005655 | 0,002458 | 0,001542 | 0,004329 |
| TNBC_324 | 25 | GPL96 | GSM282385 | 1 | 0 | 57 | 2 | 12 | 89 | 0 | NO | -0,002659 | 0,004686 | 0,004096 | 0,013892 | -0,002772 | 0,005196 | 0,006277 | 0,005496 | 0,006738 |
| TNBC_325 | 25 | GPL96 | GSM282398 | 1 | 0 | 50 | 2 | 3 | 15 | 1 | NO | -0,009787 | 0,003575 | 0,011766 | 0,010865 | -0,009165 | 0,007203 | -0,00017 | 0,001252 | 0,010868 |
| TNBC_326 | 25 | GPL96 | GSM282413 | 1 | 0 | 68 | 1 | 3 | 17 | 1 | NO | -0,010753 | 0,004426 | 0,007724 | 0,010353 | 0,000827 | 0,005282 | 0,0048 | -0,001529 | 0,005398 |
| TNBC_327 | 25 | GPL96 | GSM282427 | 1 | 0 | 56 | 2 | 3 | 120 | 0 | NO | -0,003192 | 0,005851 | 0,005314 | 0,010209 | -0,005153 | 0,004769 | 0,005878 | 0,003597 | 0,003162 |
| TNBC_328 | 25 | GPL96 | GSM282435 | 1 | 0 | 40 | 2 | 3 | 115 | 0 | NO | -0,005725 | 0,005213 | 0,009653 | 0,010603 | -0,003713 | 0,004968 | 0,004513 | 0,002826 | 0,00388 |
| TNBC_329 | 25 | GPL96 | GSM282440 | 1 | 0 | 52 | 2 | 3 | 106 | 0 | NO | -0,001959 | 0,004688 | 0,00683 | 0,010026 | -0,002495 | 0,006808 | 0,008229 | 0,005089 | 0,00493 |
| TNBC_330 | 25 | GPL96 | GSM282446 | 1 | 0 | 67 | 1 | 12 | 114 | 0 | NO | -0,004214 | 0,004424 | 0,003861 | 0,01034 | 0,000029 | 0,0031 | 0,003662 | -0,00021 | 0,005595 |
| TNBC_331 | 25 | GPL96 | GSM282454 | 1 | 0 | 43 | 1 | 3 | 8 | 1 | NO | -0,012923 | 0,004722 | 0,007517 | 0,012154 | -0,002782 | 0,008363 | 0,004315 | 0,002694 | 0,005352 |
| TNBC_332 | 25 | GPL96 | GSM282457 | 1 | 0 | 54 | 1 | 3 | 120 | 0 | NO | -0,003647 | 0,003521 | 0,003493 | 0,008677 | 0,005682 | 0,001496 | 0,005007 | 0,000184 | 0,006814 |
| TNBC_333 | 25 | GPL96 | GSM282464 | 1 | 0 | 49 | 2 | 3 | 120 | 0 | NO | -0,010668 | 0,004926 | 0,004372 | 0,011296 | -0,002228 | 0,005289 | 0,004341 | 0,000748 | 0,005816 |
| TNBC_334 | 25 | GPL96 | GSM282465 | 1 | 0 | 55 | 2 | 3 | 90 | 0 | NO | -0,004828 | 0,002773 | 0,004057 | 0,008286 | -0,005834 | 0,005189 | 0,006163 | -0,001479 | 0,005459 |
| TNBC_335 | 25 | GPL96 | GSM282474 | 1 | 0 | 45 | 2 | 3 | 120 | 0 | NO | -0,004966 | 0,003871 | 0,00709 | 0,011986 | -0,005471 | 0,006596 | 0,006148 | 0,002901 | 0,002599 |
| TNBC_336 | 25 | GPL96 | GSM282482 | 1 | 0 | 58 | 2 | 12 | 116 | 1 | NO | -0,003187 | 0,003052 | 0,003214 | 0,008036 | -0,001207 | 0,001465 | 0,004348 | -0,000213 | 0,00278 |
| TNBC_337 | 25 | GPL96 | GSM282493 | 1 | 0 | 64 | 2 | 12 | 120 | 0 | NO | -0,003258 | 0,004142 | 0,005068 | 0,009667 | -0,006953 | -0,000439 | -0,000837 | -0,000199 | 0,007721 |
| TNBC_338 | 25 | GPL96 | GSM282497 | 1 | 0 | 65 | 2 | 3 | 93 | 0 | NO | -0,004617 | 0,005038 | 0,00434 | 0,009687 | -0,000617 | 0,005627 | 0,003174 | 0,001641 | 0,005708 |
| TNBC_339 | 25 | GPL96 | GSM282511 | 1 | 0 | 58 | 2 | 3 | 120 | 0 | NO | -0,00595 | 0,004199 | 0,005472 | 0,008348 | 0,000044 | 0,009881 | 0,005003 | -0,00183 | 0,006127 |
| TNBC_340 | 25 | GPL96 | GSM282528 | 1 | 0 | 69 | 1 | 12 | 120 | 0 | NO | -0,004914 | 0,005005 | 0,004398 | 0,009803 | -0,00155 | 0,006166 | 0,004047 | 0,000418 | 0,005917 |
| TNBC_341 | 25 | GPL96 | GSM282535 | 1 | 0 | 34 | 1 | 3 | 37 | 1 | NO | -0,005184 | 0,003446 | -0,001837 | 0,007069 | -0,001889 | -0,00226 | 0,004814 | -0,00195 | 0,002409 |
| TNBC_342 | 25 | GPL96 | GSM282551 | 1 | 0 | 58 | 1 | 12 | 8 | 1 | NO | -0,013272 | 0,006514 | 0,005898 | 0,010864 | -0,002749 | 0,001833 | 0,005645 | 0,002429 | 0,004129 |
| TNBC_343 | 25 | GPL96 | GSM282565 | 1 | 0 | 55 | 2 | 3 | 78 | 1 | NO | -0,01289 | 0,00489 | 0,005624 | 0,012033 | -0,001266 | 0,00336 | 0,005985 | 0,002574 | 0,006378 |
| TNBC_344 | 25 | GPL96 | GSM282569 | 1 | 0 | 50 | 2 | 3 | 10 | 1 | NO | -0,010953 | 0,004656 | 0,006778 | 0,009507 | 0,001666 | 0,007217 | 0,002029 | 0,000756 | 0,006992 |
| TNBC_345 | 29 | GPL96 | ArrayExpress | 1 | 1 | 72 | 2 | 12 | 104 | 0 | YES | 0,000607 | 0,003166 | 0,003991 | 0,007354 | -0,003929 | 0,004081 | 0,004897 | 0,000974 | 0,009128 |
| TNBC_346 | 29 | GPL96 | ArrayExpress | 1 | 0 | 57 | 2 | 12 | 120 | 0 | YES | -0,001341 | 0,004115 | 0,00663 | 0,008698 | -0,002754 | 0,005368 | 0,005262 | 0,002058 | 0,005036 |
| TNBC_347 | 29 | GPL96 | ArrayExpress | 1 | 0 | 55 | 1 | 12 | 81 | 0 | NO | -0,001658 | 0,003826 | 0,004243 | 0,008417 | -0,013307 | -0,001843 | 0,004847 | -0,000723 | 0,006368 |
| TNBC_348 | 29 | GPL96 | ArrayExpress | 1 | 1 | 51 | 1 | 12 | 18 | 0 | NO | -0,004148 | 0,003843 | 0,005553 | 0,008308 | -0,011704 | 0,001446 | 0,003445 | -0,000031 | 0,004039 |
| TNBC_349 | 29 | GPL96 | ArrayExpress | 1 | 0 | 50 | 1 | 12 | 43 | 1 | YES | -0,002119 | 0,003307 | 0,007854 | 0,010223 | -0,012405 | 0,004604 | 0,00301 | 0,002401 | 0,007845 |
| TNBC_350 | 29 | GPL96 | ArrayExpress | 1 | 0 | 45 | 1 | 12 | 10 | 1 | NO | -0,002343 | 0,004327 | 0,009592 | 0,009188 | -0,002538 | 0,002449 | 0,003254 | -0,003269 | 0,004997 |
| TNBC_351 | 29 | GPL96 | ArrayExpress | 1 | 0 | 47 | 2 | 3 | 101 | 0 | YES | -0,001642 | 0,00536 | 0,008342 | 0,010395 | -0,004647 | 0,003013 | -0,000059 | 0,001996 | 0,005164 |
| TNBC_352 | 29 | GPL96 | ArrayExpress | 1 | 0 | 60 | 2 | 3 | 37 | 1 | YES | -0,001765 | 0,00373 | 0,007074 | 0,008159 | -0,002046 | 0,002961 | 0,006039 | -0,002091 | 0,005277 |
| TNBC_353 | 29 | GPL96 | ArrayExpress | 1 | 0 | 44 | 2 | | 120 | 0 | YES | -0,004861 | 0,003098 | 0,008005 | 0,009173 | -0,00564 | 0,005766 | 0,009052 | 0,000009 | 0,00626 |
| TNBC_354 | 29 | GPL96 | ArrayExpress | 1 | 1 | 49 | 2 | 3 | 120 | 0 | YES | -0,00224 | 0,004309 | 0,005975 | 0,007873 | -0,005306 | 0,004427 | 0,000086 | 0,004445 | 0,00575 |
| TNBC_355 | 29 | GPL96 | ArrayExpress | 1 | 1 | 43 | 2 | 12 | 53 | 0 | YES | -0,000236 | 0,003682 | 0,008585 | 0,009438 | -0,003815 | 0,000617 | 0,003017 | 0,000203 | 0,006679 |
| TNBC_356 | 29 | GPL96 | ArrayExpress | 1 | 0 | 69 | 1 | 12 | 83 | 0 | NO | -0,003319 | 0,001951 | 0,005468 | 0,006187 | -0,007271 | 0,003028 | -0,001819 | 0,00185 | 0,004143 |
| TNBC_357 | 29 | GPL96 | ArrayExpress | 1 | 0 | 44 | 2 | 3 | 120 | 0 | YES | 0,001055 | 0,003223 | 0,005992 | 0,007094 | -0,002169 | 0,002242 | 0,000056 | 0,000918 | 0,008004 |
| TNBC_358 | 29 | GPL96 | ArrayExpress | 1 | 1 | 54 | 2 | 3 | 16 | 1 | YES | -0,00123 | 0,003344 | 0,00542 | 0,007845 | -0,005651 | 0,000682 | 0,004311 | -0,001914 | 0,003927 |
| TNBC_359 | 29 | GPL96 | ArrayExpress | 1 | 1 | 40 | 2 | 12 | 117 | 0 | YES | -0,001178 | 0,004571 | 0,008519 | 0,010676 | -0,00377 | 0,005896 | 0,003478 | 0,001991 | 0,005492 |
| TNBC_360 | 29 | GPL96 | ArrayExpress | 1 | 1 | 39 | 1 | 3 | 4 | 1 | YES | -0,004036 | 0,002938 | 0,004934 | 0,007651 | -0,005265 | 0,001555 | 0,000397 | -0,00086 | 0,005938 |
| TNBC_361 | 29 | GPL96 | ArrayExpress | 1 | 1 | 50 | 2 | 3 | 103 | 0 | YES | 0,000395 | 0,002951 | 0,006559 | 0,007594 | -0,00124 | 0,004898 | 0,000554 | 0,001122 | 0,006151 |
| TNBC_362 | 29 | GPL96 | ArrayExpress | 1 | 0 | 36 | 1 | 12 | 120 | 0 | NO | -0,002773 | 0,003195 | 0,007607 | 0,008101 | -0,005226 | 0,003463 | 0,003435 | 0,001362 | 0,00523 |
| TNBC_363 | 29 | GPL96 | ArrayExpress | 1 | 1 | 74 | 1 | 3 | 98 | 0 | YES | -0,002255 | 0,00341 | 0,006158 | 0,008997 | -0,004468 | 0,002688 | 0,001335 | 0,001399 | 0,005315 |
| TNBC_364 | 29 | GPL96 | ArrayExpress | 1 | 1 | 50 | 2 | 12 | 119 | 0 | YES | -0,002722 | 0,003915 | 0,004387 | 0,006621 | 0,00028 | -0,001609 | 0,003432 | -0,007597 | 0,006129 |
| TNBC_365 | 29 | GPL96 | ArrayExpress | 1 | 0 | 63 | 2 | | 25 | 0 | YES | 0,000148 | 0,003989 | 0,007785 | 0,008537 | -0,001697 | 0,007991 | -0,001977 | 0,004346 | 0,007755 |
| TNBC_366 | 29 | GPL96 | ArrayExpress | 1 | 0 | 51 | 2 | 3 | 94 | 0 | NO | -0,003047 | 0,005722 | 0,009297 | 0,010057 | -0,005443 | 0,00917 | 0,004549 | 0,001077 | 0,009063 |
| TNBC_367 | 29 | GPL96 | ArrayExpress | 1 | 1 | 72 | 2 | | 69 | 0 | NO | -0,003484 | 0,00358 | 0,00758 | 0,008878 | -0,006053 | 0,005895 | 0,003843 | 0,00294 | 0,004424 |
| TNBC_368 | 30 | GPL96 | GSM125130 | 2 | | | | | | | NO | -0,009358 | 0,003496 | 0,002541 | 0,011586 | 0,00062 | -0,00324 | 0,003938 | -0,001187 | 0,004554 |
| TNBC_369 | 31 | GPL96 | GSM505513 | 3 | 1 | 42 | 2 | 3 | | | | -0,006853 | 0,005351 | 0,007706 | 0,008984 | -0,007659 | 0,010569 | 0,00371 | 0,004718 | 0,005794 |
| TNBC_370 | 31 | GPL96 | GSM505514 | 3 | 1 | 44 | 2 | 3 | | | | -0,014359 | 0,006155 | 0,007957 | 0,01095 | -0,007083 | 0,002699 | 0,00584 | 0,000396 | 0,003708 |
| TNBC_371 | 31 | GPL96 | GSM505516 | 3 | 1 | 39 | 2 | 3 | | | | -0,005584 | 0,005709 | 0,0095 | 0,009654 | -0,000315 | 0,007974 | 0,005026 | 0,003016 | 0,004509 |
| TNBC_372 | 31 | GPL96 | GSM505517 | 3 | 1 | 59 | 2 | 12 | | | | -0,009527 | 0,003289 | -0,002003 | 0,006927 | -0,002535 | -0,007349 | 0,000804 | -0,003067 | -0,000196 |
| TNBC_373 | 31 | GPL96 | GSM505526 | 3 | 1 | 68 | 2 | 12 | | | | -0,003536 | 0,004993 | 0,007391 | 0,007856 | -0,003634 | 0,003678 | 0,004314 | 0,003722 | 0,007309 |
| TNBC_374 | 31 | GPL96 | GSM505527 | 3 | 1 | 60 | 2 | 12 | | | | -0,003477 | 0,005877 | 0,004248 | 0,007689 | -0,006349 | 0,003895 | 0,00129 | 0,001248 | 0,006345 |
| TNBC_375 | 31 | GPL96 | GSM505528 | 3 | 1 | 52 | 2 | 3 | | | | -0,000282 | 0,002986 | 0,004632 | 0,008696 | -0,004732 | 0,001074 | 0,002399 | 0,000826 | 0,004665 |
| TNBC_376 | 31 | GPL96 | GSM505531 | 3 | 1 | 46 | 2 | 3 | | | | -0,006402 | 0,002258 | 0,002812 | 0,006488 | -0,00618 | -0,00282 | 0,00455 | 0,001713 | 0,002017 |
| TNBC_377 | 31 | GPL96 | GSM505533 | 3 | 0 | 50 | 2 | 12 | | | | -0,020329 | 0,005053 | -0,004048 | 0,006465 | -0,000566 | -0,007402 | 0,003046 | -0,004977 | 0,000604 |
| TNBC_378 | 31 | GPL96 | GSM505534 | 3 | 0 | 66 | 2 | 12 | | | | -0,006187 | 0,007927 | 0,007905 | 0,011395 | -0,003893 | 0,008448 | 0,006156 | 0,003107 | 0,004954 |
| TNBC_379 | 31 | GPL96 | GSM505541 | 3 | 1 | 62 | 2 | 3 | | | | -0,011948 | 0,005303 | 0,003661 | 0,009008 | -0,005547 | 0,002823 | 0,005066 | 0,000218 | 0,004742 |
| TNBC_380 | 31 | GPL96 | GSM505543 | 3 | 1 | 68 | 2 | 3 | | | | -0,003786 | 0,005601 | 0,012238 | 0,012171 | -0,005278 | 0,005619 | 0,003031 | 0,001328 | 0,003419 |
| TNBC_381 | 31 | GPL96 | GSM505545 | 3 | 1 | 32 | 2 | 3 | | | | -0,009891 | 0,006006 | 0,006523 | 0,00732 | -0,000924 | 0,005125 | 0,007447 | 0,003887 | 0,00456 |
| TNBC_382 | 31 | GPL96 | GSM505548 | 3 | 1 | 51 | 2 | 12 | | | | -0,015395 | 0,004028 | -0,00205 | 0,006604 | 0,000904 | -0,001646 | 0,00632 | -0,002532 | 0,001858 |
| TNBC_383 | 31 | GPL96 | GSM505552 | 3 | 1 | 40 | 2 | 3 | | | | -0,010686 | 0,003684 | 0,003258 | 0,008943 | -0,003185 | -0,001423 | 0,003042 | 0,000482 | 0,005078 |
| TNBC_384 | 31 | GPL96 | GSM505553 | 3 | 1 | 50 | 2 | 3 | | | | -0,001586 | 0,005492 | 0,006318 | 0,010128 | -0,010247 | 0,00566 | 0,001746 | 0,003317 | 0,005625 |
| TNBC_385 | 31 | GPL96 | GSM505554 | 3 | 1 | 52 | 2 | 3 | | | | -0,011798 | 0,003199 | 0,004392 | 0,008197 | -0,007157 | 0,002006 | 0,008244 | 0,000634 | 0,001035 |
| TNBC_386 | 31 | GPL96 | GSM505561 | 3 | 0 | 48 | 1 | 12 | | | | -0,009859 | 0,005342 | 0,005026 | 0,010261 | -0,005094 | 0,003683 | 0,005296 | 0,001304 | 0,002548 |
| TNBC_387 | 31 | GPL96 | GSM505562 | 3 | 1 | 61 | 2 | 3 | | | | -0,009539 | 0,007281 | 0,009575 | 0,012511 | -0,007139 | 0,006689 | 0,006952 | 0,003632 | 0,006823 |
| TNBC_388 | 31 | GPL96 | GSM505563 | 3 | 1 | 51 | 2 | 3 | | | | -0,004176 | 0,004107 | 0,003455 | 0,006524 | -0,00707 | 0,002675 | 0,004908 | -0,000631 | 0,002619 |
| TNBC_389 | 31 | GPL96 | GSM505564 | 3 | 1 | 49 | 2 | 3 | | | | -0,006379 | 0,005388 | 0,006293 | 0,011272 | -0,005676 | 0,007431 | 0,004953 | 0,003595 | 0,00608 |
| TNBC_390 | 31 | GPL96 | GSM505566 | 3 | 1 | 57 | 2 | 3 | | | | 0,001184 | 0,00453 | 0,005518 | 0,008998 | -0,00181 | 0,003977 | 0,005402 | 0,004209 | 0,000272 |
| TNBC_391 | 31 | GPL96 | GSM505568 | 3 | 1 | 50 | 2 | 3 | | | | -0,001651 | 0,004619 | 0,000325 | 0,008841 | -0,007155 | -0,000136 | 0,002577 | -0,001558 | 0,003314 |
| TNBC_392 | 31 | GPL96 | GSM505570 | 3 | 1 | 43 | 2 | 12 | | | | -0,011749 | 0,003661 | 0,006614 | 0,009544 | -0,000913 | 0,002019 | 0,006289 | -0,004646 | 0,0016 |
| TNBC_393 | 31 | GPL96 | GSM505572 | 3 | 1 | 58 | 2 | 3 | | | | -0,000946 | 0,004473 | 0,000495 | 0,008265 | -0,010839 | -0,005947 | 0,004986 | -0,001225 | 0,00174 |
| TNBC_394 | 31 | GPL96 | GSM505574 | 3 | 0 | 72 | 2 | 3 | | | | -0,005374 | 0,0047 | 0,008468 | 0,00846 | -0,001598 | 0,004289 | 0,005346 | 0,001852 | 0,002134 |
| TNBC_395 | 31 | GPL96 | GSM505575 | 3 | 0 | 33 | 2 | 3 | | | | -0,005589 | 0,007039 | 0,007981 | 0,009768 | -0,004626 | 0,001753 | 0,005467 | 0,003814 | 0,005292 |
| TNBC_396 | 31 | GPL96 | GSM505580 | 3 | 1 | 46 | 2 | 12 | | | | -0,009221 | 0,003941 | 0,00151 | 0,007976 | -0,000208 | 0,000999 | 0,005687 | -0,000723 | 0,004554 |
| TNBC_397 | 31 | GPL96 | GSM505583 | 3 | 1 | 53 | 2 | 3 | | | | -0,009498 | 0,006911 | 0,009128 | 0,009865 | -0,002429 | 0,002595 | 0,000907 | 0,003216 | 0,006497 |
| TNBC_398 | 31 | GPL96 | GSM505584 | 3 | 1 | 51 | 2 | 3 | | | | -0,002536 | 0,005775 | 0,002877 | 0,009681 | -0,00633 | 0,001679 | 0,005063 | 0,000487 | 0,002779 |

| | | | | | | | | | | | | | | | | | | | | |
|---|---|---|---|---|---|---|---|---|---|---|---|---|---|---|---|---|---|---|---|---|
| TNBC_399 | 31 | GPL96 | GSM505586 | 3 | 1 | 64 | 2 | 3 | | | | -0,010139 | 0,005308 | 0,000697 | 0,009581 | -0,004627 | -0,004682 | 0,001345 | -0,001417 | 0,005077 |
| TNBC_400 | 31 | GPL96 | GSM505588 | 3 | 0 | 36 | 2 | 3 | | | | -0,010872 | 0,004557 | 0,007015 | 0,00546 | -0,001548 | 0,003694 | 0,007135 | -0,00334 | 0,004603 |
| TNBC_401 | 31 | GPL96 | GSM505592 | 3 | 1 | 50 | 2 | 3 | | | | -0,01001 | 0,004833 | 0,010031 | 0,008828 | -0,004358 | 0,00476 | 0,000471 | 0,00355 | 0,007311 |
| TNBC_402 | 32 | GPL570 | GSM85474 | 1 | | | | 3 | | | | -0,010896 | 0,005632 | 0,011748 | 0,012112 | -0,001599 | 0,006335 | 0,006864 | 0,002627 | 0,002078 |
| TNBC_403 | 32 | GPL570 | GSM85476 | 1 | | | | 3 | | | | -0,004493 | 0,001562 | 0,004916 | 0,007756 | -0,003754 | 0,002844 | 0,003774 | -0,000308 | 0,005166 |
| TNBC_404 | 32 | GPL570 | GSM85477 | 1 | | | | 3 | | | | -0,000416 | 0,003492 | 0,006812 | 0,010118 | -0,002242 | 0,001886 | 0,005127 | 0,002567 | 0,004176 |
| TNBC_405 | 32 | GPL570 | GSM85478 | 1 | | | | 3 | | | | 0,000805 | 0,006234 | 0,006743 | 0,009405 | -0,0016 | 0,008071 | 0,006249 | 0,002589 | 0,006216 |
| TNBC_406 | 32 | GPL570 | GSM85479 | 1 | | | | 3 | | | | 0,002555 | 0,005197 | 0,008225 | 0,009698 | -0,000518 | 0,005311 | 0,00152 | 0,00436 | 0,005563 |
| TNBC_407 | 32 | GPL570 | GSM85480 | 1 | | | | 3 | | | | -0,001887 | 0,000768 | 0,003284 | 0,00804 | 0,002801 | -0,000031 | 0,002991 | -0,002063 | 0,00007 |
| TNBC_408 | 32 | GPL570 | GSM85481 | 1 | | | | 3 | | | | 0,001117 | 0,00676 | 0,007967 | 0,012113 | -0,003305 | 0,006109 | 0,006566 | 0,003648 | 0,008579 |
| TNBC_409 | 32 | GPL570 | GSM85482 | 1 | | | | 3 | | | | -0,010145 | 0,004099 | 0,002506 | 0,008949 | -0,000798 | 0,004781 | 0,005264 | -0,000313 | 0,005665 |
| TNBC_410 | 32 | GPL570 | GSM85483 | 1 | | | | 3 | | | | -0,0037 | 0,003679 | 0,0069 | 0,0084 | 0,002542 | 0,003534 | 0,003392 | 0,002174 | 0,004143 |
| TNBC_411 | 32 | GPL570 | GSM85484 | 1 | | | | 3 | | | | -0,011184 | 0,004003 | 0,006263 | 0,01854 | -0,012382 | 0,006464 | 0,008349 | 0,001565 | 0,005189 |
| TNBC_412 | 32 | GPL570 | GSM85485 | 1 | | | | 3 | | | | 0,000102 | 0,004409 | 0,009694 | 0,009383 | -0,007503 | 0,003157 | 0,006011 | 0,003133 | 0,004957 |
| TNBC_413 | 32 | GPL570 | GSM85486 | 1 | | | | 3 | | | | -0,001055 | 0,003845 | 0,005994 | 0,008524 | -0,010934 | 0,008942 | 0,00558 | 0,000993 | 0,006422 |
| TNBC_414 | 32 | GPL570 | GSM85487 | 1 | | | | 3 | | | | -0,00152 | 0,006906 | 0,0119 | 0,009829 | -0,00002 | 0,006507 | 0,004076 | 0,004249 | 0,0061 |
| TNBC_415 | 32 | GPL570 | GSM85489 | 1 | | | | 3 | | | | 0,000891 | 0,002542 | 0,004694 | 0,009439 | -0,001036 | 0,007444 | 0,005945 | 0,002679 | 0,002984 |
| TNBC_416 | 32 | GPL570 | GSM85490 | 1 | | | | 3 | | | | 0,00836 | 0,00246 | 0,006975 | 0,009734 | -0,002056 | 0,003801 | 0,006906 | 0,002223 | 0,005185 |
| TNBC_417 | 32 | GPL570 | GSM85492 | 1 | | | | 3 | | | | -0,003517 | 0,002242 | 0,006585 | 0,006373 | -0,002722 | 0,002105 | 0,004533 | -0,002066 | 0,006819 |
| TNBC_418 | 33 | GPL96 | GSM782588 | 1 | 0 | 59 | 1 | 12 | 56 | 0 | NO | -0,008405 | 0,001877 | -0,004131 | 0,007916 | 0,001222 | -0,00886 | 0,003803 | -0,00261 | 0,006017 |
| TNBC_419 | 33 | GPL96 | GSM782589 | 1 | 1 | 62 | 2 | 3 | 66 | 1 | NO | -0,004401 | 0,002636 | 0,002582 | 0,008455 | -0,00518 | 0,000247 | 0,004338 | 0,000646 | 0,004137 |
| TNBC_420 | 34 | GPL96 | GSM305166 | 1 | 0 | | | | 101 | 0 | NO | -0,006139 | 0,004316 | 0,006315 | 0,010626 | -0,002847 | 0,001339 | 0,006308 | 0,000195 | 0,003807 |
| TNBC_421 | 35 | GPL96 | GSM121673 | 1 | | | | | | | | -0,002802 | 0,006544 | 0,007454 | 0,008464 | -0,002967 | 0,006042 | 0,005604 | 0,002787 | 0,007862 |
| TNBC_422 | 35 | GPL96 | GSM121675 | 1 | | | | | | | | -0,012883 | 0,005755 | 0,006155 | 0,008843 | -0,00211 | 0,002506 | 0,007441 | 0,002117 | 0,007499 |
| TNBC_423 | 35 | GPL96 | GSM121678 | 1 | | | | | | | | -0,004554 | 0,004487 | 0,002169 | 0,007901 | -0,000831 | 0,004759 | 0,005991 | -0,0022 | 0,004224 |
| TNBC_424 | 35 | GPL96 | GSM121684 | 1 | | | | | | | | -0,004263 | 0,005854 | 0,006619 | 0,007467 | -0,001764 | 0,002636 | 0,00649 | -0,000222 | 0,006012 |
| TNBC_425 | 35 | GPL96 | GSM121685 | 1 | | | | | | | | -0,012583 | 0,006466 | 0,00388 | 0,006579 | -0,001662 | 0,001289 | 0,00718 | -0,001014 | 0,006146 |
| TNBC_426 | 35 | GPL96 | GSM121690 | 1 | | | | | | | | -0,002376 | 0,004218 | 0,004001 | 0,006777 | -0,008434 | -0,000007 | -0,001465 | -0,000122 | 0,004143 |
| TNBC_427 | 35 | GPL96 | GSM121692 | 1 | | | | | | | | -0,007909 | 0,004994 | 0,003576 | 0,008089 | -0,000224 | 0,001085 | 0,005694 | 0,00045 | 0,004335 |
| TNBC_428 | 35 | GPL96 | GSM121693 | 1 | | | | | | | | -0,004763 | 0,004747 | 0,005338 | 0,008222 | -0,001514 | 0,002867 | 0,00719 | -0,00155 | 0,003214 |
| TNBC_429 | 35 | GPL96 | GSM121697 | 1 | | | | | | | | -0,002554 | 0,004632 | 0,001333 | 0,007773 | -0,004226 | 0,001341 | 0,001347 | -0,003796 | 0,008535 |
| TNBC_430 | 35 | GPL96 | GSM121700 | 1 | | | | | | | | -0,004023 | 0,005184 | 0,002796 | 0,009276 | -0,002341 | 0,005432 | 0,002501 | -0,001184 | 0,002975 |
| TNBC_431 | 35 | GPL96 | GSM121703 | 1 | | | | | | | | -0,013726 | 0,004797 | 0,002679 | 0,006868 | -0,001939 | -0,000071 | 0,004192 | -0,001536 | 0,005961 |
| TNBC_432 | 35 | GPL96 | GSM121708 | 1 | | | | | | | | -0,005145 | 0,005298 | 0,002719 | 0,00818 | 0,00449 | 0,004181 | 0,005387 | 0,002438 | 0,003196 |
| TNBC_433 | 35 | GPL96 | GSM121709 | 1 | | | | | | | | -0,009095 | 0,004419 | 0,002128 | 0,009292 | -0,001491 | 0,005798 | 0,006989 | 0,003821 | 0,005331 |
| TNBC_434 | 35 | GPL96 | GSM121721 | 1 | | | | | | | | -0,00502 | 0,005885 | 0,003076 | 0,00666 | -0,00083 | 0,002367 | 0,004014 | -0,003045 | 0,004163 |
| TNBC_435 | 35 | GPL96 | GSM121724 | 1 | | | | | | | | -0,015329 | 0,004693 | 0,000979 | 0,006432 | -0,002614 | 0,003726 | 0,0064 | -0,00011 | 0,004997 |
| TNBC_436 | 35 | GPL96 | GSM121758 | 1 | | | | | | | | -0,010633 | 0,005007 | 0,001148 | 0,007451 | -0,004959 | -0,001013 | 0,002593 | -0,004656 | 0,004532 |
| TNBC_437 | 35 | GPL96 | GSM121763 | 1 | | | | | | | | -0,003611 | 0,003878 | 0,002745 | 0,007713 | -0,001119 | 0,006107 | 0,003661 | -0,000002 | 0,002299 |
| TNBC_438 | 35 | GPL96 | GSM121764 | 1 | | | | | | | | -0,006629 | 0,005937 | 0,004676 | 0,010372 | 0,001213 | 0,0025 | 0,00724 | 0,001958 | 0,002653 |
| TNBC_439 | 35 | GPL96 | GSM121765 | 1 | | | | | | | | -0,001823 | 0,00507 | -0,001699 | 0,009202 | -0,001868 | -0,000151 | 0,004564 | -0,005572 | 0,007399 |
| TNBC_440 | 35 | GPL96 | GSM121768 | 1 | | | | | | | | -0,004792 | 0,00577 | 0,003101 | 0,008942 | -0,001964 | 0,002406 | 0,005557 | -0,000555 | 0,005171 |
| TNBC_441 | 35 | GPL96 | GSM121772 | 1 | | | | | | | | -0,003657 | 0,004603 | 0,005285 | 0,009129 | -0,007935 | 0,007049 | 0,002802 | 0,002046 | 0,007087 |
| TNBC_442 | 35 | GPL96 | GSM121777 | 1 | | | | | | | | -0,003131 | 0,005041 | 0,005463 | 0,008723 | -0,011383 | 0,002967 | 0,004233 | -0,000057 | 0,004531 |
| TNBC_443 | 35 | GPL96 | GSM121781 | 1 | | | | | | | | -0,001115 | 0,005198 | 0,004238 | 0,009211 | -0,005339 | 0,004235 | -0,001087 | -0,000159 | 0,008428 |
| TNBC_444 | 35 | GPL96 | GSM121800 | 1 | | | | | | | | -0,008197 | 0,003918 | 0,004193 | 0,00822 | -0,007554 | 0,003874 | 0,004303 | 0,003176 | 0,004727 |
| TNBC_445 | 35 | GPL96 | GSM121807 | 1 | | | | | | | | -0,005118 | 0,006203 | 0,003201 | 0,006314 | -0,007464 | 0,003459 | 0,00808 | 0,000605 | 0,002002 |
| TNBC_446 | 35 | GPL96 | GSM121812 | 1 | | | | | | | | -0,004232 | 0,004995 | 0,002326 | 0,007516 | -0,002992 | 0,004623 | 0,007125 | 0,002161 | 0,003438 |
| TNBC_447 | 35 | GPL96 | GSM121818 | 1 | | | | | | | | -0,008949 | 0,005446 | 0,001736 | 0,007191 | -0,002662 | 0,002445 | -0,000137 | -0,000093 | 0,005706 |
| TNBC_448 | 35 | GPL96 | GSM121822 | 1 | | | | | | | | -0,013579 | 0,007571 | 0,010503 | 0,008171 | -0,003591 | 0,006793 | 0,004227 | 0,005877 | 0,005319 |
| TNBC_449 | 35 | GPL96 | GSM121824 | 1 | | | | | | | | -0,004191 | 0,007726 | 0,004476 | 0,008027 | -0,008316 | 0,004126 | 0,004157 | 0,002318 | 0,004074 |
| TNBC_450 | 35 | GPL96 | GSM121825 | 1 | | | | | | | | -0,009072 | 0,006408 | 0,004062 | 0,010716 | -0,003739 | 0,002511 | 0,003218 | 0,001486 | 0,010131 |
| TNBC_451 | 35 | GPL96 | GSM121833 | 1 | | | | | | | | -0,007093 | 0,004704 | 0,006655 | 0,006583 | -0,004897 | 0,004281 | 0,005578 | 0,00206 | 0,004818 |
| TNBC_452 | 35 | GPL96 | GSM121834 | 1 | | | | | | | | -0,009825 | 0,005762 | 0,000192 | 0,009128 | -0,001573 | 0,004929 | 0,002678 | -0,00135 | 0,003753 |
| TNBC_453 | 35 | GPL96 | GSM121836 | 1 | | | | | | | | -0,002181 | 0,007768 | 0,006675 | 0,009395 | -0,009879 | 0,005951 | 0,004524 | 0,002513 | 0,001721 |
| TNBC_454 | 35 | GPL96 | GSM121848 | 1 | | | | | | | | -0,00991 | 0,005766 | 0,004841 | 0,007539 | -0,000331 | 0,004797 | 0,007737 | 0,001303 | 0,003307 |
| TNBC_455 | 35 | GPL96 | GSM121851 | 1 | | | | | | | | -0,003653 | 0,00468 | 0,001814 | 0,007596 | -0,004545 | -0,00098 | 0,003294 | -0,002774 | 0,006346 |
| TNBC_456 | 35 | GPL96 | GSM121852 | 1 | | | | | | | | -0,006984 | 0,004271 | 0,001078 | 0,004262 | -0,008067 | -0,000205 | 0,007468 | -0,002418 | 0,003505 |
| TNBC_457 | 36 | GPL570 | GSM320216 | 1 | | | | | | | | 0,000901 | 0,004162 | 0,0092 | 0,006462 | -0,003127 | 0,004934 | 0,003463 | 0,001974 | 0,008139 |
| TNBC_458 | 36 | GPL570 | GSM320228 | 1 | | | | | | | | -0,000872 | 0,003076 | 0,006473 | 0,008823 | -0,000414 | 0,009805 | 0,006239 | 0,002569 | 0,005129 |
| TNBC_459 | 36 | GPL570 | GSM320230 | 1 | | | | | | | | -0,000914 | 0,003878 | 0,006607 | 0,009528 | -0,002749 | 0,007237 | 0,001577 | 0,000282 | 0,003699 |
| TNBC_460 | 36 | GPL570 | GSM320234 | 1 | | | | | | | | -0,001799 | 0,004942 | 0,009193 | 0,009233 | -0,001863 | 0,003898 | 0,006509 | 0,001418 | 0,005129 |
| TNBC_461 | 36 | GPL570 | GSM320237 | 1 | | | | | | | | 0,000173 | 0,005492 | 0,008343 | 0,009081 | -0,004007 | 0,004117 | 0,004711 | 0,002945 | 0,007039 |
| TNBC_462 | 37 | GPL96 | GSM152339 | 1 | | 56 | 1 | 12 | | | | -0,012466 | 0,004334 | 0,006634 | 0,009272 | -0,007999 | 0,003251 | 0,005579 | -0,001021 | 0,004851 |
| TNBC_463 | 37 | GPL96 | GSM152343 | 1 | | 85 | 2 | 3 | | | | -0,013908 | 0,006028 | 0,008928 | 0,009503 | -0,0104 | 0,003592 | 0,005437 | 0,000061 | 0,004176 |
| TNBC_464 | 37 | GPL96 | GSM152344 | 1 | | 29 | 2 | 12 | | | | -0,009875 | 0,00695 | 0,011318 | 0,010872 | -0,000808 | 0,002573 | 0,009854 | 0,001523 | 0,006268 |
| TNBC_465 | 37 | GPL96 | GSM152349 | 1 | | 65 | 2 | 3 | | | | -0,011063 | 0,005754 | 0,007993 | 0,009022 | -0,004825 | 0,007416 | 0,005085 | 0,005884 | 0,007335 |
| TNBC_466 | 37 | GPL96 | GSM152352 | 1 | | 65 | 2 | 3 | | | | -0,011535 | 0,003917 | 0,008728 | 0,006258 | -0,005031 | 0,005459 | 0,003971 | 0,001617 | 0,005815 |
| TNBC_467 | 37 | GPL96 | GSM152357 | 1 | | 41 | 2 | 3 | | | | -0,015677 | 0,007834 | 0,008941 | 0,008696 | 0,001083 | 0,00325 | 0,004241 | 0,000012 | 0,004166 |
| TNBC_468 | 37 | GPL96 | GSM152361 | 1 | | 51 | 1 | 3 | | | | -0,007559 | 0,007291 | 0,009481 | 0,008688 | -0,005162 | 0,00009 | 0,003041 | 0,00021 | 0,004465 |
| TNBC_469 | 38 | GPL570 | GSM346882 | 1 | | | | 3 | | | | -0,002692 | 0,003964 | 0,009988 | 0,008576 | -0,002207 | 0,007637 | 0,004094 | 0,001464 | 0,007733 |
| TNBC_470 | 38 | GPL570 | GSM346883 | 1 | | | | 3 | | | | -0,000225 | 0,001344 | 0,007493 | 0,007898 | -0,001315 | 0,004227 | 0,003546 | 0,000483 | 0,008288 |
| TNBC_471 | 38 | GPL570 | GSM346884 | 1 | | | | 3 | | | | -0,001878 | 0,004236 | 0,010258 | 0,010338 | 0,000544 | 0,00925 | 0,00631 | 0,002445 | 0,007072 |
| TNBC_472 | 38 | GPL570 | GSM346885 | 1 | | | | 3 | | | | -0,001322 | -0,000252 | 0,00713 | 0,006412 | 0,000315 | 0,003476 | 0,005973 | 0,001784 | 0,00519 |
| TNBC_473 | 38 | GPL570 | GSM346886 | 1 | | | | 3 | | | | -0,000574 | 0,002663 | 0,00555 | 0,008686 | -0,003309 | 0,004677 | 0,00409 | 0,002136 | 0,004654 |
| TNBC_474 | 38 | GPL570 | GSM346889 | 1 | | | | 3 | | | | -0,001025 | 0,004186 | 0,007441 | 0,008205 | -0,002901 | 0,003733 | 0,00514 | 0,00108 | 0,007746 |
| TNBC_475 | 38 | GPL570 | GSM346890 | 1 | | | | 3 | | | | -0,001376 | 0,00269 | 0,007676 | 0,008088 | 0,000478 | 0,00638 | 0,005015 | 0,000715 | 0,007403 |
| TNBC_476 | 38 | GPL570 | GSM346892 | 1 | | | | 3 | | | | -0,000844 | 0,002815 | 0,005518 | 0,006195 | -0,000553 | 0,00295 | 0,005248 | -0,000524 | 0,005821 |
| TNBC_477 | 38 | GPL570 | GSM346900 | 1 | | | | 3 | | | | -0,002024 | 0,004252 | 0,007773 | 0,007726 | -0,002761 | 0,003249 | 0,006992 | 0,001161 | 0,007842 |
| TNBC_478 | 38 | GPL570 | GSM346904 | 1 | | | | 3 | | | | -0,004837 | 0,003007 | 0,009784 | 0,008232 | -0,001898 | 0,007495 | 0,00337 | 0,002986 | 0,006715 |

| | | | | | | | | | | | | | | | | | | | | |
|---|---|---|---|---|---|---|---|---|---|---|---|---|---|---|---|---|---|---|---|---|
| TNBC_479 | 39 | GPL570 | GSM272166 | 4 | | | | | | | | -0,000841 | 0,004666 | 0,005912 | 0,009633 | -0,000467 | 0,004457 | 0,004082 | -0,001074 | 0,008837 |
| TNBC_480 | 39 | GPL570 | GSM272167 | 4 | | | | | | | | -0,001285 | 0,004199 | 0,005528 | 0,009852 | -0,000782 | 0,002798 | -0,001897 | 0,00068 | 0,008185 |
| TNBC_481 | 39 | GPL570 | GSM272221 | 4 | | | | | | | | -0,000911 | 0,005629 | 0,010301 | 0,010406 | -0,004734 | 0,005201 | 0,006741 | 0,003935 | 0,005026 |
| TNBC_482 | 39 | GPL570 | GSM272242 | 4 | | | | | | | | -0,00464 | 0,005223 | 0,004565 | 0,007674 | -0,009002 | 0,002129 | 0,005015 | -0,009644 | 0,007416 |
| TNBC_483 | 39 | GPL570 | GSM272287 | 4 | | | | | | | | 0,000345 | 0,003057 | 0,001889 | 0,008804 | -0,003547 | -0,008929 | 0,004757 | -0,005365 | 0,003493 |
| TNBC_484 | 40 | GPL96 | GSM177925 | 1 | 0 | 52 | 2 | 3 | 120 | 0 | NO | -0,001899 | 0,003218 | 0,009205 | 0,008493 | -0,000501 | 0,003561 | -0,000174 | 0,003719 | 0,003956 |
| TNBC_485 | 40 | GPL96 | GSM177935 | 1 | 0 | 37 | 2 | 3 | 14 | 1 | NO | 0,000714 | 0,00232 | 0,008618 | 0,008745 | 0,001377 | 0,005889 | 0,00022 | 0,00283 | 0,006654 |
| TNBC_486 | 40 | GPL96 | GSM177937 | 1 | 0 | 53 | 2 | 3 | 103 | 1 | NO | -0,001085 | 0,003074 | 0,006795 | 0,00819 | -0,004289 | 0,002868 | 0,002577 | 0,003462 | 0,003953 |
| TNBC_487 | 40 | GPL96 | GSM177943 | 1 | 0 | 39 | 1 | 12 | 108 | 0 | NO | -0,000787 | 0,003453 | 0,008733 | 0,010078 | -0,001041 | 0,004544 | 0,004144 | 0,000457 | 0,002996 |
| TNBC_488 | 40 | GPL96 | GSM177944 | 1 | 0 | 52 | 2 | 12 | 18 | 1 | NO | -0,002145 | 0,004881 | 0,004363 | 0,009614 | -0,000271 | 0,005769 | 0,004779 | 0,001208 | 0,003155 |
| TNBC_489 | 40 | GPL96 | GSM177954 | 1 | 0 | 37 | 2 | 12 | 26 | 1 | NO | -0,00068 | 0,001779 | 0,008947 | 0,01036 | -0,002119 | 0,006252 | 0,005356 | 0,003061 | 0,006435 |
| TNBC_490 | 40 | GPL96 | GSM177956 | 1 | 0 | 40 | 2 | 3 | 17 | 1 | NO | -0,002908 | 0,002934 | 0,006935 | 0,009496 | -0,002908 | 0,001867 | 0,005941 | -0,00008 | 0,004455 |
| TNBC_491 | 40 | GPL96 | GSM177963 | 1 | 0 | 45 | 2 | 3 | 57 | 0 | NO | -0,001263 | 0,003192 | 0,012125 | 0,010815 | -0,001715 | 0,000929 | 0,005821 | 0,002239 | 0,005169 |
| TNBC_492 | 40 | GPL96 | GSM177966 | 1 | 0 | 40 | 2 | 3 | 51 | 0 | NO | 0,000256 | 0,002894 | 0,007889 | 0,008273 | -0,001767 | 0,004867 | 0,004265 | 0,001178 | 0,00794 |
| TNBC_493 | 40 | GPL96 | GSM177970 | 1 | 0 | 47 | 1 | 3 | 108 | 0 | NO | -0,000368 | 0,003078 | 0,006787 | 0,010655 | -0,006156 | 0,005682 | 0,005064 | 0,003025 | 0,006472 |
| TNBC_494 | 40 | GPL96 | GSM177885 | 1 | 0 | 57 | 2 | 3 | 24 | 1 | NO | -0,001977 | 0,004084 | 0,005411 | 0,009521 | -0,002445 | -0,000572 | 0,004746 | -0,000156 | 0,008116 |
| TNBC_495 | 40 | GPL96 | GSM177897 | 1 | 0 | 46 | 2 | 3 | 120 | 0 | NO | -0,002904 | 0,002114 | 0,005272 | 0,00866 | -0,000968 | 0,00433 | 0,00508 | 0,003235 | 0,006597 |
| TNBC_496 | 40 | GPL96 | GSM177898 | 1 | 0 | 57 | 2 | 12 | 93 | 1 | NO | 0,000304 | 0,002054 | 0,002156 | 0,008928 | -0,002409 | -0,001471 | 0,005269 | -0,000659 | 0,006468 |
| TNBC_497 | 40 | GPL96 | GSM177899 | 1 | 0 | 33 | 2 | 3 | 15 | 1 | NO | -0,001241 | 0,004185 | 0,012069 | 0,008176 | -0,000852 | 0,002628 | 0,004973 | 0,001703 | 0,006594 |
| TNBC_498 | 40 | GPL96 | GSM177903 | 1 | 0 | 57 | 2 | 12 | 60 | 1 | NO | -0,00046 | 0,004654 | 0,004245 | 0,010127 | -0,001611 | 0,000332 | 0,003341 | 0,001841 | 0,007135 |
| TNBC_499 | 40 | GPL96 | GSM177909 | 1 | 0 | 58 | 2 | 3 | 4 | 1 | NO | -0,009463 | 0,002751 | 0,007419 | 0,010246 | 0,000429 | 0,001539 | 0,004812 | 0,00217 | 0,005173 |
| TNBC_500 | 40 | GPL96 | GSM177913 | 1 | 0 | 47 | 2 | 3 | 120 | 0 | NO | -0,00278 | 0,004004 | 0,004641 | 0,009078 | -0,0038 | 0,004488 | 0,007515 | 0,001505 | 0,008849 |
| TNBC_501 | 40 | GPL96 | GSM177988 | 1 | 0 | 43 | 2 | 3 | 88 | 0 | NO | -0,003509 | 0,00423 | 0,005972 | 0,007465 | -0,004778 | -0,000423 | 0,003848 | 0,000995 | 0,006096 |
| TNBC_502 | 40 | GPL96 | GSM177999 | 1 | 0 | 43 | 2 | 12 | 13 | 1 | NO | -0,001505 | 0,003476 | 0,005577 | 0,008924 | -0,002594 | 0,00192 | 0,007944 | 0,00096 | 0,008275 |
| TNBC_503 | 40 | GPL96 | GSM178003 | 1 | 0 | 43 | 2 | 3 | 95 | 1 | NO | -0,003925 | 0,00165 | 0,006174 | 0,007799 | -0,000975 | 0,0034 | 0,004062 | 0,00085 | 0,00355 |
| TNBC_504 | 40 | GPL96 | GSM178009 | 1 | 0 | 53 | 2 | 3 | 9 | 1 | NO | -0,002771 | 0,004593 | 0,000499 | 0,011733 | -0,003021 | 0,003245 | 0,005715 | 0,000649 | 0,005293 |
| TNBC_505 | 40 | GPL96 | GSM178012 | 1 | 0 | 38 | 2 | 3 | 120 | 0 | NO | -0,005447 | 0,003065 | 0,009804 | 0,009424 | -0,003116 | 0,00722 | 0,002607 | 0,001951 | 0,005573 |
| TNBC_506 | 40 | GPL96 | GSM178021 | 1 | 0 | 33 | 2 | 3 | 120 | 0 | NO | -0,011573 | 0,004835 | 0,008075 | 0,008898 | -0,002417 | 0,008512 | 0,003149 | 0,0009 | 0,006506 |
| TNBC_507 | 40 | GPL96 | GSM178023 | 1 | 0 | 42 | 2 | 3 | 120 | 0 | NO | -0,002229 | 0,003671 | 0,008155 | 0,005979 | -0,005612 | 0,007185 | 0,007233 | 0,001573 | 0,005717 |
| TNBC_508 | 40 | GPL96 | GSM178058 | 1 | 0 | 46 | 2 | 3 | 120 | 0 | NO | 0,000892 | 0,005527 | 0,009766 | 0,009171 | -0,008477 | 0,003098 | 0,00255 | 0,001264 | 0,007014 |
| TNBC_509 | 40 | GPL96 | GSM178065 | 1 | 0 | 47 | 2 | 3 | 120 | 0 | NO | -0,001428 | 0,002698 | 0,008068 | 0,007936 | -0,005449 | 0,002566 | 0,006659 | 0,001746 | 0,007221 |
| TNBC_510 | 40 | GPL96 | GSM178075 | 1 | 0 | 48 | 2 | 3 | 95 | 0 | NO | -0,008118 | 0,002908 | 0,00354 | 0,010495 | -0,002908 | 0,002883 | 0,005175 | 0,001644 | 0,003869 |
| TNBC_511 | 40 | GPL96 | GSM178078 | 1 | 0 | 39 | 2 | 3 | 120 | 0 | NO | 0,001089 | 0,004677 | 0,006582 | 0,01168 | -0,001287 | 0,005221 | 0,002076 | 0,003019 | 0,00636 |
| TNBC_512 | 40 | GPL96 | GSM178079 | 1 | 0 | 46 | 2 | 3 | 13 | 1 | NO | -0,000712 | 0,006436 | 0,010219 | 0,010956 | -0,002579 | 0,005183 | 0,006214 | 0,002825 | 0,008524 |
| TNBC_513 | 40 | GPL96 | GSM178082 | 1 | 0 | 39 | 2 | 3 | 44 | 1 | NO | -0,00265 | 0,003368 | 0,004086 | 0,010128 | -0,004061 | 0,001634 | 0,005974 | 0,00057 | 0,008221 |
| TNBC_514 | 40 | GPL96 | GSM177978 | 1 | 0 | 47 | 2 | 3 | 120 | 0 | NO | -0,007929 | 0,003027 | 0,008669 | 0,007388 | -0,006029 | 0,00522 | -0,002118 | 0,003254 | 0,005841 |
| TNBC_515 | 40 | GPL96 | GSM177979 | 1 | 0 | 39 | 2 | 3 | 84 | 1 | NO | -0,007881 | 0,003311 | 0,005988 | 0,007753 | -0,008484 | 0,004631 | 0,007288 | 0,000765 | 0,007333 |
| TNBC_516 | 40 | GPL96 | GSM177980 | 1 | 0 | 32 | 2 | 3 | 120 | 0 | NO | -0,011799 | 0,004297 | 0,008047 | 0,010424 | -0,004283 | 0,007797 | 0,007105 | 0,001761 | 0,007856 |
| TNBC_517 | 40 | GPL96 | GSM177985 | 1 | 0 | 45 | 2 | 3 | 120 | 0 | NO | -0,001154 | 0,002497 | 0,005299 | 0,008348 | -0,002216 | 0,002025 | 0,005765 | 0,001473 | 0,00539 |
| TNBC_518 | 40 | GPL96 | GSM177993 | 1 | 0 | 38 | 1 | 12 | 9 | 1 | NO | -0,003805 | 0,002546 | 0,004076 | 0,008621 | 0,003859 | -0,00104 | 0,00513 | -0,001224 | 0,006873 |
| TNBC_519 | 40 | GPL96 | GSM177994 | 1 | 0 | 47 | 1 | 12 | 42 | 0 | NO | -0,011318 | 0,003849 | 0,001937 | 0,008508 | -0,001952 | 0,00047 | 0,005967 | 0,001054 | 0,005043 |
| TNBC_520 | 40 | GPL96 | GSM178013 | 1 | 0 | 40 | 1 | 3 | 120 | 0 | NO | -0,004578 | 0,003131 | 0,010223 | 0,007319 | -0,000069 | 0,00857 | 0,009869 | 0,00408 | 0,009489 |
| TNBC_521 | 40 | GPL96 | GSM178016 | 1 | 0 | 40 | 2 | 12 | 120 | 0 | NO | -0,002087 | 0,001624 | 0,002933 | 0,008477 | -0,004342 | -0,00054 | 0,005074 | -0,002393 | 0,003403 |
| TNBC_522 | 40 | GPL96 | GSM178034 | 1 | 0 | 32 | 1 | 12 | 120 | 0 | NO | -0,00525 | 0,005698 | 0,009656 | 0,00895 | -0,00753 | 0,008101 | 0,006396 | 0,001804 | 0,004934 |
| TNBC_523 | 40 | GPL96 | GSM178035 | 1 | 0 | 50 | 1 | 3 | 120 | 0 | NO | -0,002681 | 0,0028 | 0,006097 | 0,009177 | -0,004681 | 0,004464 | 0,005439 | 0,00172 | 0,005734 |
| TNBC_524 | 42 | GPL570 | GSM308259 | 1 | | 61 | 2 | 3 | 13 | 1 | NO | -0,00331 | 0,003636 | 0,007364 | 0,009839 | -0,005793 | 0,002177 | 0,005867 | 0,001161 | 0,005716 |
| TNBC_525 | 42 | GPL570 | GSM308261 | 1 | | 55 | 2 | 3 | 18 | 1 | NO | -0,004347 | 0,003278 | 0,006517 | 0,008636 | -0,003394 | 0,00446 | 0,004037 | 0,001883 | 0,006346 |
| TNBC_526 | 42 | GPL570 | GSM308271 | 1 | | 50 | 1 | 12 | 29 | 1 | NO | -0,002267 | 0,001736 | 0,007052 | 0,009125 | -0,004881 | 0,006547 | 0,006722 | 0,002546 | 0,006886 |
| TNBC_527 | 42 | GPL570 | GSM308272 | 1 | | 56 | 2 | 3 | 14 | 1 | NO | -0,006303 | 0,004093 | 0,007949 | 0,007375 | -0,004165 | 0,00245 | 0,00539 | -0,000693 | 0,007313 |
| TNBC_528 | 42 | GPL570 | GSM308273 | 1 | | 55 | 1 | 12 | 19 | 1 | NO | -0,005555 | 0,003557 | 0,012033 | 0,008034 | -0,003753 | 0,007076 | 0,005215 | 0,004854 | 0,008739 |
| TNBC_529 | 42 | GPL570 | GSM308275 | 1 | | 39 | 2 | 3 | 7 | 1 | NO | -0,004887 | 0,002493 | 0,004489 | 0,009111 | -0,00456 | 0,005965 | 0,004956 | 0,004006 | 0,007419 |
| TNBC_530 | 42 | GPL570 | GSM308280 | 1 | | 51 | 2 | 12 | 16 | 1 | NO | -0,010601 | 0,00366 | 0,007425 | 0,00784 | -0,005407 | 0,00334 | 0,004874 | 0,001444 | 0,006651 |
| TNBC_531 | 42 | GPL570 | GSM308281 | 1 | | 66 | 1 | 3 | 5 | 1 | NO | -0,00594 | 0,004217 | 0,010073 | 0,008815 | -0,003992 | 0,004024 | 0,006648 | 0,000351 | 0,005825 |
| TNBC_532 | 42 | GPL570 | GSM308283 | 1 | | 55 | 1 | 3 | 13 | 1 | NO | -0,004455 | 0,002911 | 0,008189 | 0,008955 | -0,004467 | 0,0077 | 0,003809 | 0,003834 | 0,007712 |
| TNBC_533 | 42 | GPL570 | GSM308285 | 1 | | 50 | 2 | 3 | 7 | 1 | NO | -0,010873 | 0,005043 | 0,010354 | 0,010524 | -0,000495 | 0,002407 | 0,006023 | 0,001183 | 0,010368 |
| TNBC_534 | 42 | GPL570 | GSM308290 | 1 | | 62 | 2 | 3 | 6 | 1 | NO | -0,00535 | 0,003759 | 0,005831 | 0,008835 | -0,005853 | 0,002889 | 0,003386 | 0,001042 | 0,006989 |
| TNBC_535 | 42 | GPL570 | GSM308295 | 1 | | 63 | 2 | 3 | 11 | 1 | NO | -0,00628 | 0,003328 | 0,007286 | 0,009447 | 0,000401 | 0,001135 | 0,004687 | 0,003445 | 0,00503 |
| TNBC_536 | 42 | GPL570 | GSM308301 | 1 | | 42 | 1 | 12 | 15 | 1 | YES | -0,001226 | 0,003245 | 0,007116 | 0,009342 | 0,000165 | 0,00229 | 0,004356 | 0,001859 | 0,006547 |
| TNBC_537 | 42 | GPL570 | GSM308307 | 1 | | 36 | 2 | 3 | 29 | 1 | NO | -0,003107 | 0,003375 | 0,008129 | 0,008535 | -0,001539 | 0,004781 | 0,004435 | 0,004798 | 0,007968 |
| TNBC_538 | 42 | GPL570 | GSM308309 | 1 | | 42 | 2 | 12 | 20 | 1 | NO | -0,002421 | 0,002759 | 0,007977 | 0,008697 | 0,001573 | 0,00545 | 0,005485 | 0,001266 | 0,006843 |
| TNBC_539 | 42 | GPL570 | GSM308311 | 1 | | 57 | 2 | 12 | 36 | 1 | NO | -0,001688 | 0,004815 | 0,006773 | 0,011571 | 0,001777 | 0,006282 | 0,008142 | 0,004202 | 0,007889 |
| TNBC_540 | 42 | GPL570 | GSM308312 | 1 | | 44 | 2 | 12 | 115 | 1 | NO | -0,001834 | 0,004107 | 0,00998 | 0,009668 | 0,000084 | 0,007801 | 0,00088 | 0,004085 | 0,008911 |
| TNBC_541 | 42 | GPL570 | GSM308313 | 1 | | | | | 4 | 1 | | -0,003119 | 0,004435 | 0,009216 | 0,010017 | -0,000645 | 0,005744 | 0,006647 | 0,002786 | 0,006789 |
| TNBC_542 | 42 | GPL570 | GSM308314 | 1 | | 58 | 2 | 3 | 8 | 1 | NO | -0,002097 | 0,004611 | 0,006456 | 0,010579 | -0,009505 | 0,006842 | 0,004544 | 0,003098 | 0,005246 |
| TNBC_543 | 42 | GPL570 | GSM308316 | 1 | | 43 | 1 | 3 | 0 | 1 | NO | -0,001955 | 0,000909 | 0,003968 | 0,007428 | -0,000175 | 0,00076 | 0,001138 | 0,000442 | 0,008684 |
| TNBC_544 | 42 | GPL570 | GSM308324 | 1 | | 57 | 2 | 3 | 14 | 1 | NO | 0,000754 | 0,005145 | 0,007278 | 0,009401 | -0,000232 | 0,002858 | 0,005422 | 0,00054 | 0,009312 |
| TNBC_545 | 42 | GPL570 | GSM308328 | 1 | | 52 | 2 | 3 | 16 | 1 | NO | -0,000353 | 0,005851 | 0,007959 | 0,009008 | 0,002148 | 0,002842 | 0,000136 | 0,001824 | 0,010708 |
| TNBC_546 | 42 | GPL570 | GSM308329 | 1 | | 40 | 2 | 3 | 1 | 1 | | 0,000284 | 0,002066 | 0,006304 | 0,009432 | 0,000539 | 0,005789 | -0,004751 | 0,001984 | 0,005682 |
| TNBC_547 | 42 | GPL570 | GSM308330 | 1 | | 31 | 2 | 3 | 0 | 1 | | -0,000544 | 0,002854 | 0,009229 | 0,008479 | 0,004824 | 0,00529 | 0,000513 | 0,001499 | 0,006377 |
| TNBC_548 | 42 | GPL570 | GSM308333 | 1 | | 30 | 1 | 3 | 18 | 1 | NO | -0,002064 | 0,004909 | 0,009268 | 0,009278 | -0,002882 | 0,006176 | 0,001622 | 0,004488 | 0,006912 |
| TNBC_549 | 42 | GPL570 | GSM308335 | 1 | | 38 | 2 | 12 | 50 | 1 | NO | -0,00117 | 0,004916 | 0,008673 | 0,008408 | -0,00058 | 0,006543 | 0,002144 | 0,002063 | 0,006448 |
| TNBC_550 | 42 | GPL570 | GSM308336 | 1 | | 28 | 2 | 12 | 8 | 1 | NO | -0,003603 | 0,001947 | 0,007113 | 0,008092 | -0,002966 | 0,004823 | 0,000136 | 0,000239 | 0,006598 |
| TNBC_551 | 42 | GPL570 | GSM308338 | 1 | | | | | 10 | 1 | | -0,002623 | 0,004132 | 0,008455 | 0,008987 | 0,000456 | 0,005862 | 0,002678 | 0,002878 | 0,008145 |
| TNBC_552 | 42 | GPL570 | GSM308339 | 1 | | 49 | 2 | 3 | 3 | 1 | NO | -0,003811 | 0,004461 | 0,01134 | 0,009147 | -0,003794 | 0,00428 | 0,004943 | 0,004013 | 0,006803 |
| TNBC_553 | 42 | GPL570 | GSM308340 | 1 | | 46 | 2 | 12 | 60 | 1 | NO | 0,003317 | 0,004164 | 0,009772 | 0,0105 | -0,00534 | 0,00217 | 0,005881 | 0,00057 | 0,007011 |
| TNBC_554 | 42 | GPL570 | GSM308341 | 1 | | 57 | 2 | 12 | 17 | 1 | NO | -0,002263 | 0,004923 | 0,010327 | 0,011711 | -0,004709 | 0,005455 | 0,004452 | 0,001608 | 0,005514 |
| TNBC_555 | 42 | GPL570 | GSM308344 | 1 | | 79 | 2 | 12 | 6 | 1 | NO | 0,000648 | 0,002217 | 0,008159 | 0,00897 | -0,005327 | 0,001975 | 0,004821 | -0,001172 | 0,007907 |
| TNBC_556 | 42 | GPL570 | GSM308346 | 1 | | 69 | 2 | 3 | 13 | 1 | NO | 0,000476 | 0,004259 | 0,006297 | 0,008416 | -0,001377 | 0,001586 | 0,006739 | 0,00113 | 0,00517 |
| TNBC_557 | 42 | GPL570 | GSM308348 | 1 | | 39 | 2 | 3 | 25 | 1 | YES | 0,001916 | 0,003122 | 0,010187 | 0,008686 | -0,003159 | 0,001368 | 0,004801 | 0,000733 | 0,00616 |
| TNBC_558 | 42 | GPL570 | GSM308349 | 1 | | 51 | 2 | 3 | 4 | 1 | NO | 0,001036 | 0,003916 | 0,007285 | 0,00889 | -0,002584 | 0,000716 | 0,004424 | 0,000351 | 0,006355 |

| | | | | | | | | | | | | | | | | | | | |
|---|---|---|---|---|---|---|---|---|---|---|---|---|---|---|---|---|---|---|---|
| TNBC_559 | 42 | GPL570 | GSM308350 | 1 | 65 | | 3 | 15 | 1 | NO | -0,000335 | 0,002474 | 0,007547 | 0,00909 | -0,000338 | 0,006032 | 0,006071 | 0,001101 | 0,00475 |
| TNBC_560 | 42 | GPL570 | GSM308352 | 1 | 44 | 2 | 3 | 0 | 1 | | -0,000997 | 0,003283 | 0,009932 | 0,008552 | -0,001903 | 0,001621 | 0,003682 | 0,001569 | 0,006184 |
| TNBC_561 | 42 | GPL570 | GSM308354 | 1 | 35 | 1 | 3 | 21 | 1 | NO | 0,001114 | 0,005176 | 0,007145 | 0,008887 | -0,002717 | 0,004242 | 0,002521 | 0,002643 | 0,007462 |
| TNBC_562 | 42 | GPL570 | GSM308356 | 1 | 39 | 2 | 3 | 18 | 1 | YES | -0,001154 | 0,004555 | 0,009391 | 0,008012 | -0,003945 | 0,002755 | 0,005793 | 0,000617 | 0,006918 |
| TNBC_563 | 42 | GPL570 | GSM308364 | 1 | 44 | 1 | 12 | 29 | 1 | NO | -0,001456 | -0,000488 | -0,000863 | 0,006767 | -0,004723 | -0,006312 | 0,005206 | -0,005004 | 0,003512 |
| TNBC_564 | 42 | GPL570 | GSM308374 | 1 | 51 | 2 | 3 | 6 | 1 | NO | -0,000032 | 0,004122 | 0,009087 | 0,008325 | -0,004453 | 0,002315 | 0,001574 | 0,003145 | 0,006635 |
| TNBC_565 | 42 | GPL570 | GSM308376 | 1 | 56 | 2 | 3 | 6 | 1 | NO | -0,001001 | 0,006223 | 0,011021 | 0,01023 | 0,001685 | 0,008861 | 0,003387 | 0,004449 | 0,007978 |
| TNBC_566 | 42 | GPL570 | GSM308377 | 1 | 28 | 2 | 3 | 19 | 1 | YES | 0,000176 | 0,004159 | 0,00752 | 0,0098 | -0,001513 | 0,006448 | 0,003893 | 0,002301 | 0,00734 |
| TNBC_567 | 42 | GPL570 | GSM308378 | 1 | 41 | 2 | 3 | 15 | 1 | NO | 0,00014 | 0,004471 | 0,011297 | 0,008932 | -0,000886 | 0,00353 | 0,006363 | 0,002917 | 0,006646 |
| TNBC_568 | 42 | GPL570 | GSM308380 | 1 | 56 | 2 | 3 | 7 | 1 | NO | -0,000956 | 0,00357 | 0,007984 | 0,008324 | -0,003419 | 0,00209 | 0,005881 | 0,001523 | 0,006413 |
| TNBC_569 | 42 | GPL570 | GSM308388 | 1 | 37 | 1 | 3 | 17 | 1 | YES | -0,006146 | 0,002572 | 0,006537 | 0,007906 | -0,003601 | 0,002194 | 0,003822 | 0,001194 | 0,008439 |
| TNBC_570 | 42 | GPL570 | GSM308391 | 1 | | | | 7 | 1 | | -0,002155 | 0,00396 | 0,008052 | 0,009415 | 0,001006 | 0,007005 | 0,002725 | 0,002439 | 0,005998 |
| TNBC_571 | 42 | GPL570 | GSM308392 | 1 | | | | 6 | 1 | | -0,006765 | 0,004951 | 0,007804 | 0,00826 | -0,005008 | 0,002836 | 0,00287 | 0,000986 | 0,006019 |
| TNBC_572 | 42 | GPL570 | GSM308393 | 1 | 40 | 2 | 3 | 33 | 1 | NO | -0,001325 | 0,002772 | 0,010265 | 0,008129 | -0,002231 | 0,005392 | 0,003354 | 0,001187 | 0,005927 |
| TNBC_573 | 42 | GPL570 | GSM308394 | 1 | 34 | 2 | 3 | 22 | 1 | YES | -0,002091 | 0,003522 | 0,006101 | 0,009426 | -0,000479 | 0,007844 | 0,002962 | 0,002854 | 0,006832 |
| TNBC_574 | 42 | GPL570 | GSM308404 | 1 | 50 | 2 | 3 | 8 | 1 | NO | -0,002243 | 0,005815 | 0,010546 | 0,01072 | -0,00112 | 0,004694 | 0,00196 | 0,003196 | 0,009181 |
| TNBC_575 | 42 | GPL570 | GSM308405 | 1 | | | | 24 | 1 | | 0,000625 | 0,003861 | 0,007813 | 0,008404 | -0,004274 | 0,001309 | 0,003911 | -0,002127 | 0,007829 |
| TNBC_576 | 42 | GPL570 | GSM308407 | 1 | 40 | 1 | 3 | 94 | 1 | NO | 0,000117 | 0,001742 | 0,008371 | 0,009054 | -0,000089 | 0,004359 | 0,004414 | 0,0012 | 0,008448 |
| TNBC_577 | 42 | GPL570 | GSM308408 | 1 | 45 | 2 | 12 | 4 | 1 | NO | -0,004677 | 0,003609 | 0,008629 | 0,008288 | -0,003084 | 0,005265 | 0,002818 | 0,002257 | 0,006913 |
| TNBC_578 | 42 | GPL570 | GSM308438 | 1 | 63 | 2 | 12 | 35 | 1 | NO | 0,001685 | 0,001995 | 0,006656 | 0,008754 | -0,002001 | 0,001488 | 0,003728 | -0,001475 | 0,006057 |
| TNBC_579 | 42 | GPL570 | GSM308449 | 1 | 56 | 2 | 3 | 47 | 1 | NO | 0,001285 | 0,002068 | 0,005704 | 0,008222 | -0,001783 | 0,002755 | 0,008892 | 0,000743 | 0,005174 |

High levels of expression

Low levels of expression

**Table S3. WikiPathways_2015.**

| Term | Overlap | P-value | Adjusted P-va | Old P | Old A | Odds Ratio | Combined Sc | Genes |
|---|---|---|---|---|---|---|---|---|
| RB in Cancer(I | 26/87 | 3,12E-13 | 1,26E-10 | 0 | 0 | 5,56518761 | 160,252847 | TOP2A;CDKN1A;PCNA;PRIM1;SMC2;CCNB2;CCNB1;CDC45;SKP2;RFC5;PLK4;BARD1;RRM1;RFC4;RRM2;H2AFZ;CDC25B;CCNA2;ANLN;CCNE1;CDK4;POLE2;RBP1;MCM3;CDK1;MCM4 |
| Cell Cycle(Hor | 25/104 | 1,69E-10 | 3,42E-08 | 0 | 0 | 4,47643604 | 100,712464 | CDKN1A;PCNA;BUB1B;MCM10;CDC14A;CDC20;CCNB2;CCNB1;CDC45;SKP2;BUB1;CDKN2A;PLK1;CDC6;CDC25C;CDC25B;CCNA2;ESPL1;CCNE1;CDK4;CDK1;MCM3;MCM4;MCM2;MAD2L1 |
| Gastric cancer | 13/43 | 2,33E-07 | 3,13E-05 | 0 | 0 | 5,6298991 | 85,9913803 | TOP2A;UBE2C;AURKA;TPX2;ESM1;CENPF;HIST4H4;RUVBL1;MCM4;MYBL2;HIST1H4I;ECT2;KIF20B |
| DNA Replicati | 12/41 | 1,00E-06 | 1,01E-04 | 0 | 0 | 5,45033383 | 75,2861341 | RFC5;PRIM2;PCNA;CDC45;RFC4;PRIM1;POLE2;MCM3;MCM4;MCM10;CDC6;MCM2 |
| Cori Cycle(Ho | 6/15 | 7,78E-05 | 0,00224534 | 0 | 0 | 7,44878957 | 70,4749077 | TPI1;PGAM1;SLC2A1;PGK1;GAPDH;PFKP |
| ATM Signaling | 11/40 | 5,57E-06 | 3,22E-04 | 0 | 0 | 5,12104283 | 61,9517677 | CDKN1A;CCNB1;RAD51;CRADD;FANCD2;CCNE1;H2AFX;CDK1;BRCA1;BID;CDC25C |
| DNA Replicati | 11/41 | 7,25E-06 | 3,66E-04 | 0 | 0 | 4,99613935 | 59,1271582 | RFC5;PCNA;CDC45;RFC4;PRIM1;POLE2;MCM3;MCM4;MCM10;CDC6;MCM2 |
| Matrix Metall | 8/29 | 1,05E-04 | 0,00283088 | 0 | 0 | 5,13709626 | 47,0585361 | MMP12;MMP14;MMP7;MMP13;MMP16;BSG;TIMP1;MMP10 |
| Gastric cancer | 9/35 | 7,14E-05 | 0,00221791 | 0 | 0 | 4,78850758 | 45,7190274 | TOP2A;FANCI;RFC4;UBE2C;DSCC1;ATAD2;S100A6;COL9A1;COL9A3 |
| G1 to S cell cy | 13/60 | 1,40E-05 | 6,30E-04 | 0 | 0 | 4,03476102 | 45,0853765 | PRIM2;CDKN1A;PCNA;CDKN2A;PRIM1;CDC45;CCNE1;CDK4;POLE2;MCM3;CDK1;MCM4;MCM2 |
| Spinal Cord In | 18/96 | 3,05E-06 | 2,06E-04 | 0 | 0 | 3,49162011 | 44,3395274 | EGR1;NOS2;ARG1;CXCL3;CXCL2;KLK8;GFAP;MMP12;CXCL10;NR4A1;GJA1;VCAN;COL2A1;CDK4;CASP3;IL1B;SLIT1;CDK1 |
| Matrix Metall | 8/30 | 1,37E-04 | 0,00344977 | 0 | 0 | 4,96585971 | 44,1875771 | MMP12;MMP14;MMP7;MMP13;MMP16;BSG;TIMP1;MMP10 |
| DNA Damage | 14/69 | 1,47E-05 | 5,95E-04 | 0 | 0 | 3,77837152 | 42,0356611 | CDKN1A;H2AFX;BRCA1;CDC25C;CCNB2;CCNB1;RAD51;FANCD2;CCNE1;CDK4;CASP3;CDK1;PMAIP1;BID |
| Spinal Cord In | 20/114 | 2,60E-06 | 2,10E-04 | 0 | 0 | 3,26701297 | 42,0199766 | EGR1;NOS2;ARG1;MIF;KLK8;CXCL2;GFAP;MMP12;LGALS3;CXCL10;NR4A1;GJA1;VCAN;COL2A1;CDK4;CASP3;IL1B;CDK1;SLIT1;CCL2 |
| miRNA regula | 13/63 | 2,45E-05 | 8,99E-04 | 0 | 0 | 3,84262954 | 40,8003304 | CDKN1A;H2AFX;BRCA1;CDC25C;CCNB2;CCNB1;RAD51;FANCD2;CCNE1;CDK4;CASP3;CDK1;BID |
| G1 to S cell cy | 13/67 | 4,86E-05 | 0,00163621 | 0 | 0 | 3,61321882 | 35,8860627 | CDKN1A;PCNA;CDKN2A;PRIM1;CCNB1;CDC45;CCNE1;CDK4;POLE2;MCM3;CDK1;MCM4;MCM2 |
| Glycolysis anc | 10/47 | 1,61E-04 | 0,00381679 | 0 | 0 | 3,96212211 | 34,6152701 | TPI1;PKM;PGAM1;SLC2A1;PGK1;ALDOC;ENO1;GAPDH;HK2;PFKP |
| Integrated Ca | 8/35 | 4,33E-04 | 0,00873893 | 0 | 0 | 4,25645118 | 32,9689838 | BARD1;BLM;CDK4;CASP3;PLK1;CDK1;BRCA1;CDC25B |
| Glycolysis anc | 10/49 | 2,31E-04 | 0,00518936 | 0 | 0 | 3,80040284 | 31,8176899 | TPI1;PKM;PGAM1;SLC2A1;PGK1;ALDOC;ENO1;GAPDH;HK2;PFKP |
| IL1 and megal | 6/24 | 0,00138024 | 0,0242442 | 0 | 0 | 4,65549348 | 30,6587456 | IL1B;F2R;CCL2;TIMP1;PLA2G7;HBEGF |
| Urea cycle an | 5/20 | 0,00348687 | 0,05031054 | 0 | 0 | 4,65549348 | 26,3442785 | GATM;PYCRL;ACY1;ARG1;SMS |
| Leptin Insulin | 4/15 | 0,00700997 | 0,08581903 | 0 | 0 | 4,96585971 | 24,6327587 | SOCS2;SOCS3;SOCS1;PIK3R3 |
| Retinol metak | 7/37 | 0,00315297 | 0,04717777 | 0 | 0 | 3,52307615 | 20,2908414 | SULT2B1;CYP26A1;CYP26B1;RDH10;ALDH1A2;RBP1;RARA |
| Senescence a | 15/108 | 6,50E-04 | 0,01250637 | 0 | 0 | 2,58638527 | 18,9799544 | CDKN1A;CEBPB;SERPINB2;PCNA;CDKN2A;IGFBP3;SERPINE1;IL24;FN1;INHBA;CDC25B;MMP14;IL1B;CCL3;CD44 |
| Integrated Pa | 23/195 | 3,34E-04 | 0,00710303 | 0 | 0 | 2,19643795 | 17,5807516 | TOP2A;EGR1;CDKN1A;BLM;RRM1;PCNA;IGFBP3;PLK1;BUB1B;BRCA1;INHBA;PTGS2;CDC25C;CCNA2;GPRC5A;RAD51;WT1;CCNE1;CDK4;CASP3;RARA;BID;JUNB |
| Oncostatin M | 10/65 | 0,00231757 | 0,03601144 | 0 | 0 | 2,86491907 | 17,3821426 | SOCS3;EGR1;CEBPB;MMP13;CASP3;SERPINE1;OSM;CCL2;JUNB;CYR61 |
| Vitamin A anc | 7/40 | 0,00496547 | 0,06917419 | 0 | 0 | 3,25884544 | 17,2889782 | SULT2B1;CYP26A1;CYP26B1;RDH10;ALDH1A2;RBP1;RARA |
| miRNA Regula | 14/105 | 0,00145391 | 0,02447409 | 0 | 0 | 2,48292986 | 16,2222259 | CDKN1A;H2AFX;BRCA1;CDC25C;CCNB2;CCNB1;RAD51;FANCD2;CCNE1;CDK4;CASP3;CDK1;PMAIP1;BID |
| Complement | 8/51 | 0,00548236 | 0,07382917 | 0 | 0 | 2,92109395 | 15,2078545 | PROS1;SERPINE1;F2R;C3AR1;PLAUR;A2M;F3;KNG1 |
| Type II interfe | 7/44 | 0,00847812 | 0,10073999 | 0 | 0 | 2,96258676 | 14,1323294 | SOCS3;CXCL10;SOCS1;NOS2;HIST4H4;IL1B;HIST1H4I |
| Cardiac Proge | 8/53 | 0,00694574 | 0,08768991 | 0 | 0 | 2,81086399 | 13,9689469 | SOX2;TBX20;CXCR4;PAX6;GATA4;NCAM1;SCN5A;INHBA |
| Kennedy path | 3/14 | 0,03604482 | 0,29124218 | 0 | 0 | 3,99042299 | 13,2601436 | PCYT1B;ETNK2;ETNK1 |
| TP53 Network | 4/21 | 0,02380625 | 0,21858466 | 0 | 0 | 3,54704265 | 13,2581614 | CDKN1A;CDKN2A;PMAIP1;BID |
| Purine metab | 18/158 | 0,00209782 | 0,03390074 | 0 | 0 | 2,1214907 | 13,0829309 | GUCY2C;PRIM2;RRM1;RRM2;PRIM1;AK4;NME5;ADCY8;AK7;APRT;NT5E;ATIC;PKM;POLE2;POLR2D;ENPP1;POLR2F;PDE9A |
| Alpha6-Beta4 | 9/64 | 0,00691587 | 0,09012939 | 0 | 0 | 2,61871508 | 13,0253229 | DSP;LAMA5;CDKN1A;RTKN;MMP7;CASP3;PIK3R3;MET;EIF4E |
| Complement | 8/55 | 0,00868617 | 0,10026318 | 0 | 0 | 2,70865075 | 12,8553203 | PROS1;SERPINE1;F2R;C3AR1;PLAUR;A2M;F3;KNG1 |
| PodNet: prote | 30/306 | 0,0011106 | 0,02039471 | 0 | 0 | 1,82568372 | 12,4198563 | COL18A1;LAMA5;CDKN1A;KHDRBS3;SH3KBP1;ARPC1B;SEMA3F;PTGS2;CXXC5;CYR61;CTNNA1;NCK2;KIRREL3;EGLN3;CDKN2A;IGFBP3;PLAUR;KRT7;F3;SMAD7;CYP26A1;CXCL10;VANGL2;F |
| Prostaglandin | 5/30 | 0,02058146 | 0,20280266 | 0 | 0 | 3,10366232 | 12,0526528 | ANXA2;ANXA3;S100A6;PTGS2;S100A10 |
| Prostaglandin | 5/31 | 0,02348575 | 0,22065681 | 0 | 0 | 3,00354418 | 11,2673798 | ANXA2;ANXA3;S100A6;PTGS2;S100A10 |
| Type II interfe | 7/50 | 0,01675586 | 0,18295592 | 0 | 0 | 2,60707635 | 10,6603534 | SOCS3;CXCL10;SOCS1;NOS2;HIST4H4;IL1B;HIST1H4I |
| Monoamine T | 5/32 | 0,0266418 | 0,23398451 | 0 | 0 | 2,90968343 | 10,5483993 | IL1B;TNFRSF11B;CDC25C;SLC6A2;STX1A |
| Apoptosis-rel | 7/52 | 0,02047331 | 0,21208249 | 0 | 0 | 2,50680418 | 9,74804203 | SOCS3;CDKN1A;F2R;CTNNA1;BIRC5;IER3;SMAD7 |
| TGF Beta Sign | 7/52 | 0,02047331 | 0,20678043 | 0 | 0 | 2,50680418 | 9,74804203 | TGIF1;BAMBI;SERPINE1;LEF1;INHBA;SKIL;SMAD7 |
| Fluoropyrimic | 5/35 | 0,03767616 | 0,29845431 | 0 | 0 | 2,66028199 | 8,72234013 | RRM1;RRM2;TK1;UPP1;UMPS |
| TGF Beta Sign | 7/55 | 0,02706861 | 0,23267487 | 0 | 0 | 2,37006941 | 8,55448231 | TGIF1;BAMBI;SERPINE1;LEF1;INHBA;SKIL;SMAD7 |
| Adipogenesis( | 14/134 | 0,01298877 | 0,14576288 | 0 | 0 | 1,94557937 | 8,45095476 | EGR2;CDKN1A;CEBPB;SERPINE1;OSM;GATA4;AHR;MIF;SOCS3;CYP26A1;CYP26B1;SOCS1;RARA;TRIB3 |
| AGE/RAGE pa | 8/66 | 0,02452038 | 0,22013851 | 0 | 0 | 2,25720896 | 8,37029682 | LGALS3;MSR1;MMP14;MMP7;MMP13;NOS2;CASP3;EZR |
| Wnt Signaling | 11/100 | 0,01831332 | 0,1946995 | 0 | 0 | 2,04841713 | 8,19392797 | SOX2;FOSL1;MMP7;RACGAP1;WNT7B;TCF7;LEF1;FZD9;WNT7A;CD44;NKD2 |
| Nucleotide M | 3/19 | 0,07879036 | 0,50525881 | 0 | 0 | 2,94031167 | 7,47122796 | RRM1;RRM2;MTHFD2 |
| One Carbon M | 4/28 | 0,06071227 | 0,45421773 | 0 | 0 | 2,66028199 | 7,45307114 | AHCY;ATIC;MTHFD1L;MTHFD2 |
| Serotonin Tra | 2/11 | 0,11491923 | 0,64482456 | 0 | 0 | 3,38581344 | 7,32529461 | IL1B;STX1A |

| | | | | | | | | |
|---|---|---|---|---|---|---|---|---|
| Endochondral | 7/59 | 0,03791907 | 0,294602 | 0 | 0 | 2,20938674 | 7,22977879 | ADAMTS4;MMP13;COL2A1;MGP;ENPP1;SLC38A2;SOX5 |
| Adipogenesis | 13/129 | 0,02137056 | 0,20556445 | 0 | 0 | 1,87663303 | 7,21704449 | EGR2;CEBPB;SERPINE1;OSM;GATA4;AHR;MIF;SOCS3;CYP26A1;CYP26B1;SOCS1;RARA;TRIB3 |
| Wnt Signaling | 10/93 | 0,02740598 | 0,23066703 | 0 | 0 | 2,00236279 | 7,20248674 | SOX2;FOSL1;MMP7;RACGAP1;WNT7B;LEF1;FZD9;WNT7A;CD44;NKD2 |
| Hypertrophy I | 3/20 | 0,08914654 | 0,56273751 | 0 | 0 | 2,79329609 | 6,75272007 | EIF4E;CYR61;HBEGF |
| Nucleotide M | 3/20 | 0,08914654 | 0,55408001 | 0 | 0 | 2,79329609 | 6,75272007 | RRM1;RRM2;MTHFD2 |
| Nuclear recep | 4/30 | 0,07478599 | 0,5120939 | 0 | 0 | 2,48292986 | 6,43854678 | CYP26A1;NR1I2;RARA;ABCG1 |
| One Carbon M | 4/30 | 0,07478599 | 0,503559 | 0 | 0 | 2,48292986 | 6,43854678 | AHCY;ATIC;MTHFD1L;MTHFD2 |
| Hypertrophy I | 3/21 | 0,10003659 | 0,5857215 | 0 | 0 | 2,66028199 | 6,12455237 | EIF4E;CYR61;HBEGF |
| Sphingolipid M | 3/21 | 0,10003659 | 0,57735405 | 0 | 0 | 2,66028199 | 6,12455237 | CERS3;SPHK1;SGMS2 |
| Integrated Br | 14/151 | 0,03278089 | 0,27027507 | 0 | 0 | 1,72654063 | 5,90115986 | BARD1;BLM;DCAKD;PLK1;AHR;BRCA1;HMGCR;CDC25B;SMAD7;AURKA;RAD51;CDK4;CASP3;BID |
| IL-4 Signaling | 6/53 | 0,06356837 | 0,46693856 | 0 | 0 | 2,10814799 | 5,80929543 | SOCS3;CEBPB;SOCS1;NFIL3;BIRC5;SOCS5 |
| Oxidative Dan | 4/32 | 0,0903495 | 0,55304844 | 0 | 0 | 2,32774674 | 5,59606568 | CDKN1A;PCNA;CASP3;C3AR1 |
| Trans-sulfurat | 4/32 | 0,0903495 | 0,54479399 | 0 | 0 | 2,32774674 | 5,59606568 | AHCY;MTHFD1L;MTHFD2;PSAT1 |
| ErbB Signaling | 6/54 | 0,06842329 | 0,47660361 | 0 | 0 | 2,06910821 | 5,54943517 | CDKN1A;BCL2L11;TGFA;NRG2;EREG;HBEGF |
| Histone modi | 1/5 | 0,24119161 | 0,95530795 | 0 | 0 | 3,72439479 | 5,29669869 | HAT1 |
| Endochondral | 7/67 | 0,06730077 | 0,47700894 | 0 | 0 | 1,94557937 | 5,25030868 | ADAMTS4;MMP13;COL2A1;MGP;ENPP1;SLC38A2;SOX5 |
| Apoptosis Mo | 8/80 | 0,06491417 | 0,46830939 | 0 | 0 | 1,86219739 | 5,09253129 | BCL2L11;CRADD;CDKN2A;CASP3;BIRC5;PMAIP1;TNFRSF11B;BID |
| Cytokines and | 3/23 | 0,12328628 | 0,65536392 | 0 | 0 | 2,42895312 | 5,08439672 | CSF3;IL1B;CXCL2 |
| Pathogenic Es | 6/56 | 0,07875909 | 0,51320438 | 0 | 0 | 1,99521149 | 5,07055389 | TUBB6;TUBB3;ARPC1B;NCK2;ARHGEF2;EZR |
| EGFR1 Signali | 15/172 | 0,04415672 | 0,33659085 | 0 | 0 | 1,62400935 | 5,06692567 | USP6NL;TGIF1;ERRFI1;CEBPB;SH3KBP1;KRT7;PIK3R3;EPS8;SOCS3;GJA1;SOCS1;ELF3;HAT1;NCK2;SPRY2 |
| ErbB signaling | 5/45 | 0,09211386 | 0,5472647 | 0 | 0 | 2,06910821 | 4,93426415 | CDKN1A;TGFA;NRG2;EREG;HBEGF |
| Signaling Path | 8/83 | 0,0770227 | 0,51011756 | 0 | 0 | 1,79488905 | 4,60147646 | ERRFI1;CDKN1A;CCNE1;CDKN2A;CDK4;SPRY2;BRCA1;MET |
| Nuclear Recep | 4/35 | 0,1163105 | 0,63499247 | 0 | 0 | 2,12822559 | 4,57886018 | CYP26A1;NR1I2;RARA;ABCG1 |
| Cytokines and | 3/25 | 0,14826802 | 0,7582314 | 0 | 0 | 2,23463687 | 4,26532667 | CSF3;IL1B;CXCL2 |
| Cholesterol Bi | 2/15 | 0,19107748 | 0,89761977 | 0 | 0 | 2,48292986 | 4,10943836 | HMGCR;DHCR7 |
| Cholesterol Bi | 2/15 | 0,19107748 | 0,8873023 | 0 | 0 | 2,48292986 | 4,10943836 | HMGCR;DHCR7 |
| Nuclear Recep | 4/37 | 0,13521729 | 0,70945178 | 0 | 0 | 2,01318637 | 4,02812866 | NR4A2;NR4A1;NR1I2;RARA |
| Arrhythmoge | 7/74 | 0,1017732 | 0,57910383 | 0 | 0 | 1,76153807 | 4,02512948 | DSP;GJA1;LEF1;TCF7;CTNNA1;DSC2;ITGA9 |
| Regulation of | 1/6 | 0,28194975 | 1 | 0 | 0 | 3,10366232 | 3,92931845 | GATA4 |
| Nuclear Recep | 4/38 | 0,14510881 | 0,75158924 | 0 | 0 | 1,96020778 | 3,78373298 | NR4A2;NR4A1;NR1I2;RARA |
| Proteasome D | 6/63 | 0,12133294 | 0,65358011 | 0 | 0 | 1,77352133 | 3,74074122 | HIST1H2AH;H2AFZ;HIST1H2AK;H2AFX;HIST1H2AE;HIST1H2AC |
| Apoptosis Mo | 2/16 | 0,21105567 | 0,897542 | 0 | 0 | 2,32774674 | 3,62112046 | CASP3;BID |
| Neurotransmi | 2/16 | 0,21105567 | 0,88819261 | 0 | 0 | 2,32774674 | 3,62112046 | SLC38A2;STX1A |
| Wnt Signaling | 9/106 | 0,11617019 | 0,64291449 | 0 | 0 | 1,58111099 | 3,40365603 | VANGL2;RUVBL1;LEF1;FZD9;CDK1;FHL2;WNT7A;CDC25C;MAPK8IP1 |
| Extracellular | 3/28 | 0,18838487 | 0,89538219 | 0 | 0 | 1,99521149 | 3,3305432 | TGFA;PROM1;MET |
| AMPK Signalin | 6/69 | 0,16512348 | 0,83387356 | 0 | 0 | 1,61930208 | 2,91646303 | CCNA2;CDKN1A;CCNB1;PIK3R3;HMGCR;CAMKK2 |
| Eicosanoid Sy | 2/18 | 0,2515005 | 0,95854906 | 0 | 0 | 2,06910821 | 2,85601141 | PTGES2;PTGS2 |
| Aryl Hydrocar | 4/43 | 0,1983363 | 0,90031307 | 0 | 0 | 1,73227665 | 2,80246196 | CDKN1A;PSRC1;NCOA7;AHR |
| Hair Follicle D | 4/43 | 0,1983363 | 0,89030959 | 0 | 0 | 1,73227665 | 2,80246196 | VCAN;LEF1;NCAM1;INHBA |
| IL-6 signaling | 4/43 | 0,1983363 | 0,88052597 | 0 | 0 | 1,73227665 | 2,80246196 | SOCS3;TIMP1;A2M;JUNB |
| Folate Metab | 6/70 | 0,17300322 | 0,85235734 | 0 | 0 | 1,59616919 | 2,80039115 | GPX2;AHCY;MTHFD2;IL1B;SERPINE1;CCL2 |
| Primary Focal | 6/70 | 0,17300322 | 0,84208798 | 0 | 0 | 1,59616919 | 2,80039115 | KIRREL3;LAMA5;CDKN1A;PCNA;WT1;PLAUR |
| One carbon m | 4/44 | 0,2096243 | 0,91062601 | 0 | 0 | 1,69290672 | 2,64506244 | PCYT1B;ETNK2;GAD1;ETNK1 |
| Regulation of | 4/44 | 0,2096243 | 0,9009385 | 0 | 0 | 1,69290672 | 2,64506244 | SPRED1;CDK1;KIF2C;AURKB |
| Ovarian Infert | 3/31 | 0,23086659 | 0,95173573 | 0 | 0 | 1,80212651 | 2,64176478 | EGR1;CEBPB;CDK4 |
| Apoptosis(Ho | 7/86 | 0,17880568 | 0,85997016 | 0 | 0 | 1,51574206 | 2,60928277 | BCL2L11;CRADD;CDKN2A;CASP3;BIRC5;PMAIP1;BID |
| Blood Clotting | 2/19 | 0,27181934 | 1 | 0 | 0 | 1,96020778 | 2,55340122 | SERPINB2;SERPINE1 |
| Ovarian Infert | 3/32 | 0,24540284 | 0,96255095 | 0 | 0 | 1,74581006 | 2,45260854 | EGR1;CEBPB;CDK4 |
| ACE Inhibitor | 1/8 | 0,35702133 | 1 | 0 | 0 | 2,32774674 | 2,39748545 | KNG1 |
| Folate-Alcoho | 1/8 | 0,35702133 | 1 | 0 | 0 | 2,32774674 | 2,39748545 | CEBPB |
| Androgen rec | 7/90 | 0,20879954 | 0,91690233 | 0 | 0 | 1,44837575 | 2,26870772 | PSMC3IP;CDKN1A;CCNE1;SNORD96A;FHL2;BRCA1;ETV5 |
| Interferon typ | 5/61 | 0,22904597 | 0,95396464 | 0 | 0 | 1,52639131 | 2,24964522 | SOCS3;SOCS1;PRMT1;SNORD96A;EIF4E |
| miR-targeted | 24/362 | 0,16859139 | 0,84087555 | 0 | 0 | 1,23460601 | 2,19794109 | GALNT7;SLC38A1;CEBPB;SERPINE2;ANXA2;MYO10;CDKN2A;PLK1;CYR61;AURKB;CORO1C;MIR24-2;GJA1;NT5E;PKM;MTHFD2;P4HA2;PSAT1;RDH10;MIR140;TRIP13;SLC38A2;MET;SLC25A |

| | | | | | | | | |
|---|---|---|---|---|---|---|---|---|
| Semaphorin i | 5/62 | 0,23907963 | 0,95631854 | 0 | 0 | 1,50177209 | 2,14897367 | SEMA5A;SEMA7A;CD72;TREM2;MET |
| Apoptosis Mo | 2/21 | 0,31233523 | 1 | 0 | 0 | 1,77352133 | 2,06380812 | CASP3;BID |
| Eicosanoid Sy | 2/21 | 0,31233523 | 1 | 0 | 0 | 1,77352133 | 2,06380812 | PTGES2;PTGS2 |
| Globo Sphing | 2/21 | 0,31233523 | 1 | 0 | 0 | 1,77352133 | 2,06380812 | B3GALT5;ST6GALNAC4 |
| PluriNetWork | 19/284 | 0,19148787 | 0,87910341 | 0 | 0 | 1,24583628 | 2,05928117 | CDKN1A;TFAP2C;KDM3A;CDKN2A;IGFBP3;LEF1;WWP2;BRCA1;UTF1;ETV5;SMAD7;SOX2;SOCS2;P4HA1;CASP3;MYBL2;SGK1;RCOR2;CD44 |
| DNA Damage | 7/93 | 0,23243861 | 0,94853735 | 0 | 0 | 1,40165395 | 2,04519413 | FOSL1;CDKN1A;BCL2L11;CDKN2A;WNT7B;PMAIP1;WNT7A |
| Apoptosis(Mu | 6/79 | 0,24998662 | 0,96185329 | 0 | 0 | 1,41432713 | 1,96074941 | BCL2L11;CRADD;CDKN2A;CASP3;BIRC5;BID |
| Effects of Nitr | 1/9 | 0,39156304 | 1 | 0 | 0 | 2,06910821 | 1,94001395 | NOS2 |
| Macrophage | 1/9 | 0,39156304 | 1 | 0 | 0 | 2,06910821 | 1,94001395 | F3 |
| Mismatch rep | 1/9 | 0,39156304 | 1 | 0 | 0 | 2,06910821 | 1,94001395 | PCNA |
| Mismatch rep | 1/9 | 0,39156304 | 1 | 0 | 0 | 2,06910821 | 1,94001395 | PCNA |
| BDNF signalin | 10/142 | 0,23297933 | 0,9412365 | 0 | 0 | 1,31140662 | 1,91046442 | SHC4;GABRB3;EGR1;EGR2;BCL2L11;CASP3;SNORD96A;NCK2;NCAM1;EIF4E |
| Hedgehog Sig | 2/22 | 0,33242598 | 1 | 0 | 0 | 1,69290672 | 1,86446262 | CCNB1;CDK1 |
| Signal Transd | 2/22 | 0,33242598 | 1 | 0 | 0 | 1,69290672 | 1,86446262 | RACGAP1;SPHK1 |
| IL-1 Signaling | 3/36 | 0,30464208 | 1 | 0 | 0 | 1,55183116 | 1,84453399 | IL1RN;IL1B;IL1RAP |
| Peptide GPCR | 5/66 | 0,28028459 | 1 | 0 | 0 | 1,4107556 | 1,7944103 | CCR1;CXCR2;C3AR1;CXCR4;GCAT |
| SIDS Susceptil | 11/161 | 0,24795514 | 0,96321035 | 0 | 0 | 1,27230878 | 1,77424406 | SOX2;EGR1;IL1RN;GJA1;CEBPB;KCNJ8;CASP3;IL1B;SCN5A;FOXM1;GAPDH |
| Amyotrophic | 3/37 | 0,31959132 | 1 | 0 | 0 | 1,50988978 | 1,72234971 | CASP3;SLC1A2;BID |
| Chemokine si | 11/165 | 0,27326676 | 1 | 0 | 0 | 1,24146493 | 1,61056092 | SHC4;CCR1;CXCL10;CXCR2;ELMO1;PIK3R3;CXCR4;CCL2;CXCL3;ADCY8;CXCL2 |
| Osteoblast(M | 1/10 | 0,42425076 | 1 | 0 | 0 | 1,86219739 | 1,596705 | TNFRSF11B |
| SREBF and mi | 1/10 | 0,42425076 | 1 | 0 | 0 | 1,86219739 | 1,596705 | HMGCR |
| Trans-sulfurat | 1/10 | 0,42425076 | 1 | 0 | 0 | 1,86219739 | 1,596705 | AHCY |
| Vitamin D Me | 1/10 | 0,42425076 | 1 | 0 | 0 | 1,86219739 | 1,596705 | DHCR7 |
| Neural Crest | 7/101 | 0,29920591 | 1 | 0 | 0 | 1,29063186 | 1,55730646 | CDH6;COL2A1;MIA;SNAI1;HOXA1;SOX5;GFAP |
| Blood Clotting | 2/24 | 0,37205005 | 1 | 0 | 0 | 1,55183116 | 1,5343372 | SERPINB2;SERPINE1 |
| Signal Transd | 2/24 | 0,37205005 | 1 | 0 | 0 | 1,55183116 | 1,5343372 | RACGAP1;SPHK1 |
| NOD pathway | 3/39 | 0,3494865 | 1 | 0 | 0 | 1,43245953 | 1,50593088 | NLRP10;IL1B;CARD9 |
| Oxidation by | 3/39 | 0,3494865 | 1 | 0 | 0 | 1,43245953 | 1,50593088 | CYB5R2;CYP26A1;CYP26B1 |
| Focal Adhesio | 12/186 | 0,29805626 | 1 | 0 | 0 | 1,20141767 | 1,45428369 | COMP;LAMA5;COL2A1;TNN;COL11A1;STYK1;TNC;FN1;MET;PGF;THBS4;ITGA9 |
| Insulin Signali | 10/153 | 0,30728311 | 1 | 0 | 0 | 1,21712248 | 1,43618721 | SOCS3;EGR1;RPS6KA6;SOCS1;SLC2A1;ENPP1;PIK3R3;TRIB3;SGK1;EIF4E |
| Parkinsons Di | 3/40 | 0,36438867 | 1 | 0 | 0 | 1,39664805 | 1,40996397 | GP1BB;CCNE1;CASP3 |
| Myometrial R | 10/155 | 0,32139969 | 1 | 0 | 0 | 1,20141767 | 1,36369291 | CNN2;GJA1;RGS1;CALD1;IGFBP3;IL1B;MAFF;RGS20;ADM;ADCY8 |
| Ganglio Sphin | 1/11 | 0,4551839 | 1 | 0 | 0 | 1,69290672 | 1,3324086 | ST3GAL5 |
| Corticotropin | 6/90 | 0,35380933 | 1 | 0 | 0 | 1,24146493 | 1,28987851 | NR4A2;FOSL1;NR4A1;GJA1;CASP3;JUNB |
| Wnt Signaling | 4/58 | 0,37910624 | 1 | 0 | 0 | 1,28427406 | 1,24566725 | FOSL1;WNT7B;FZD9;WNT7A |
| EGF/EGFR Sig | 10/161 | 0,36446578 | 1 | 0 | 0 | 1,15664434 | 1,16742728 | USP6NL;EPS8;ERRFI1;GJA1;PCNA;SH3KBP1;NCK2;SPRY2;MYBL2;AURKA |
| Physiological | 2/27 | 0,42957049 | 1 | 0 | 0 | 1,37940548 | 1,16555546 | MYEF2;GATA4 |
| IL-7 Signaling | 3/43 | 0,40867706 | 1 | 0 | 0 | 1,29920748 | 1,16256986 | CCNA2;BCL2L11;CDK4 |
| Tryptophan m | 3/43 | 0,40867706 | 1 | 0 | 0 | 1,29920748 | 1,16256986 | PRMT1;ALDH1A2;DHCR24 |
| Wnt Signaling | 4/60 | 0,40375874 | 1 | 0 | 0 | 1,24146493 | 1,12593142 | FOSL1;WNT7B;FZD9;WNT7A |
| Alanine and a | 1/12 | 0,48445657 | 1 | 0 | 0 | 1,55183116 | 1,12465472 | GAD1 |
| Alanine and a | 1/12 | 0,48445657 | 1 | 0 | 0 | 1,55183116 | 1,12465472 | GAD1 |
| Insulin Signali | 10/163 | 0,37898869 | 1 | 0 | 0 | 1,14245239 | 1,10846318 | SOCS3;EGR1;RPS6KA6;SOCS1;SLC2A1;ENPP1;PIK3R3;TRIB3;SGK1;EIF4E |
| Heart Develo | 3/44 | 0,42324189 | 1 | 0 | 0 | 1,26968004 | 1,0916854 | FOXC1;TBX20;GATA4 |
| Metapathway | 8/129 | 0,39004925 | 1 | 0 | 0 | 1,1548511 | 1,08727184 | SULT2B1;CYP26A1;CYP26B1;MGST2;HS6ST2;CHST2;HS3ST1;HS2ST1 |
| Leptin signali | 4/61 | 0,41601928 | 1 | 0 | 0 | 1,22111305 | 1,07094504 | SOCS2;SOCS3;IL1RN;IL1B |
| Focal Adhesio | 11/183 | 0,39456658 | 1 | 0 | 0 | 1,11935362 | 1,04096238 | LAMA5;COL2A1;TNN;COL11A1;STYK1;TNC;FN1;MET;PGF;THBS4;ITGA9 |
| miR-targeted | 10/166 | 0,40085764 | 1 | 0 | 0 | 1,12180566 | 1,02549744 | MIR24-2;CEBPB;PKM;SERPINE2;ANXA2;MTHFD2;RDH10;PTRH1;KCNN4;CORO1C |
| IL-3 Signaling | 6/97 | 0,42168711 | 1 | 0 | 0 | 1,15187468 | 0,9946342 | SOCS2;SOCS3;BCL2L11;SLC2A1;RARA;BIRC5 |
| Arylhydrocar | 2/29 | 0,46633039 | 1 | 0 | 0 | 1,28427406 | 0,97972247 | CDKN1A;AHR |
| Dopminergic | 2/29 | 0,46633039 | 1 | 0 | 0 | 1,28427406 | 0,97972247 | SOX2;NR4A2 |
| Iron uptake a | 7/115 | 0,42266618 | 1 | 0 | 0 | 1,13351146 | 0,97614899 | GABRB3;CYBRD1;ATP6V0A4;ATP1A1;ATP1B1;SGK1;ATP6V1C2 |
| Tryptophan m | 3/46 | 0,45198148 | 1 | 0 | 0 | 1,21447656 | 0,96443292 | PRMT1;ALDH1A2;DHCR24 |

| | | | | | | | | |
|---|---|---|---|---|---|---|---|---|
| Homologous r | 1/13 | 0,51215781 | 1 | 0 | 0 | 1,43245953 | 0,95849088 | RAD51 |
| Homologous r | 1/13 | 0,51215781 | 1 | 0 | 0 | 1,43245953 | 0,95849088 | RAD51 |
| Osteoclast(M | 1/13 | 0,51215781 | 1 | 0 | 0 | 1,43245953 | 0,95849088 | TNFRSF11B |
| Quercetin and | 1/13 | 0,51215781 | 1 | 0 | 0 | 1,43245953 | 0,95849088 | ACOX2 |
| Serotonin and | 1/13 | 0,51215781 | 1 | 0 | 0 | 1,43245953 | 0,95849088 | ARC |
| Myometrial R | 9/151 | 0,42303915 | 1 | 0 | 0 | 1,10991898 | 0,9548528 | CNN2;GJA1;RGS1;IGFBP3;IL1B;MAFF;RGS20;ADM;ADCY8 |
| Differentiatio | 3/47 | 0,46612915 | 1 | 0 | 0 | 1,18863663 | 0,90727748 | NT5E;WNT7B;INHBA |
| Heart Develop | 3/47 | 0,46612915 | 1 | 0 | 0 | 1,18863663 | 0,90727748 | FOXC1;TBX20;GATA4 |
| Dopminergic I | 2/30 | 0,48417559 | 1 | 0 | 0 | 1,24146493 | 0,90044401 | SOX2;NR4A2 |
| Inflammatory | 2/30 | 0,48417559 | 1 | 0 | 0 | 1,24146493 | 0,90044401 | LAMA5;FN1 |
| Diurnally Regu | 3/48 | 0,48011246 | 1 | 0 | 0 | 1,16387337 | 0,85397453 | CEBPB;TUBB3;STBD1 |
| Inflammatory | 2/31 | 0,50164517 | 1 | 0 | 0 | 1,20141767 | 0,8288127 | LAMA5;FN1 |
| Biogenic Amir | 1/14 | 0,53837191 | 1 | 0 | 0 | 1,330141 | 0,82363085 | GAD1 |
| Keap1-Nrf2(M | 1/14 | 0,53837191 | 1 | 0 | 0 | 1,330141 | 0,82363085 | CEBPB |
| Osteoblast Sig | 1/14 | 0,53837191 | 1 | 0 | 0 | 1,330141 | 0,82363085 | TNFRSF11B |
| Kit Receptor S | 4/67 | 0,48803977 | 1 | 0 | 0 | 1,11175964 | 0,7975301 | SPRED1;SOCS1;SH3KBP1;SOCS5 |
| TSH signaling | 4/67 | 0,48803977 | 1 | 0 | 0 | 1,11175964 | 0,7975301 | EGR1;CCNE1;CDK4;TSHB |
| Diurnally Regu | 3/50 | 0,50754333 | 1 | 0 | 0 | 1,11731844 | 0,75773542 | CEBPB;TUBB3;STBD1 |
| Id Signaling Pa | 3/51 | 0,52097197 | 1 | 0 | 0 | 1,09541023 | 0,71427213 | TGIF1;CCNA2;CCNE1 |
| Biogenic Amir | 1/15 | 0,56317864 | 1 | 0 | 0 | 1,24146493 | 0,71279752 | GAD1 |
| GPCRs, Class C | 1/15 | 0,56317864 | 1 | 0 | 0 | 1,24146493 | 0,71279752 | GPRC5A |
| TFs Regulate r | 1/15 | 0,56317864 | 1 | 0 | 0 | 1,24146493 | 0,71279752 | MYEF2 |
| Selenium Mic | 5/88 | 0,51408798 | 1 | 0 | 0 | 1,0580667 | 0,70399617 | GPX2;IL1B;SERPINE1;CCL2;PTGS2 |
| Metapathway | 10/181 | 0,50949511 | 1 | 0 | 0 | 1,02883834 | 0,69378173 | SULT2B1;CYP26A1;CYP26B1;GPX2;MGST2;CHST5;HS6ST2;CHST2;HS3ST1;HS2ST1 |
| Exercise-indu | 3/52 | 0,53419836 | 1 | 0 | 0 | 1,07434465 | 0,67360125 | CEBPB;STBD1;HIST1H2BL |
| TOR Signaling | 2/34 | 0,55169991 | 1 | 0 | 0 | 1,09541023 | 0,65149636 | PRR5;DDIT4 |
| Deregulation | 1/16 | 0,58665349 | 1 | 0 | 0 | 1,16387337 | 0,62071804 | RAB27B |
| GPCRs, Class C | 1/16 | 0,58665349 | 1 | 0 | 0 | 1,16387337 | 0,62071804 | GPRC5A |
| ID signaling pa | 1/16 | 0,58665349 | 1 | 0 | 0 | 1,16387337 | 0,62071804 | CCNE1 |
| Osteoclast Sig | 1/16 | 0,58665349 | 1 | 0 | 0 | 1,16387337 | 0,62071804 | TNFRSF11B |
| Transcriptiona | 1/16 | 0,58665349 | 1 | 0 | 0 | 1,16387337 | 0,62071804 | CEBPB |
| G13 Signaling | 2/35 | 0,56757729 | 1 | 0 | 0 | 1,0641128 | 0,60269045 | RTKN;CIT |
| Vitamin B12 M | 3/54 | 0,56001635 | 1 | 0 | 0 | 1,03455411 | 0,5998234 | IL1B;SERPINE1;CCL2 |
| Peptide GPCR | 4/73 | 0,55615841 | 1 | 0 | 0 | 1,02038213 | 0,59866035 | CCR1;CXCR2;C3AR1;CXCR4 |
| Amino Acid m | 5/92 | 0,5539783 | 1 | 0 | 0 | 1,0120638 | 0,597755 | PKM;ARG1;P4HA2;SMS;FARSB |
| Hematopoieti | 3/55 | 0,57259598 | 1 | 0 | 0 | 1,01574403 | 0,56635339 | CSF3;IL1B;CXCR4 |
| FAS pathway a | 2/36 | 0,58304405 | 1 | 0 | 0 | 1,03455411 | 0,55813422 | CASP3;LMNB1 |
| GPCRs, Other | 5/94 | 0,57330999 | 1 | 0 | 0 | 0,99053053 | 0,55106057 | CHRM3;PROKR2;F2R;CXCR2;CELSR1 |
| ACE Inhibitor | 1/17 | 0,6088679 | 1 | 0 | 0 | 1,09541023 | 0,54349211 | KNG1 |
| Drug Inductio | 1/17 | 0,6088679 | 1 | 0 | 0 | 1,09541023 | 0,54349211 | NR1I2 |
| Mitochondria | 1/17 | 0,6088679 | 1 | 0 | 0 | 1,09541023 | 0,54349211 | MYEF2 |
| Serotonin Rec | 1/17 | 0,6088679 | 1 | 0 | 0 | 1,09541023 | 0,54349211 | GATA4 |
| Serotonin and | 1/17 | 0,6088679 | 1 | 0 | 0 | 1,09541023 | 0,54349211 | ARC |
| Statin Pathwa | 1/17 | 0,6088679 | 1 | 0 | 0 | 1,09541023 | 0,54349211 | HMGCR |
| Parkin-Ubiqui | 4/76 | 0,58833635 | 1 | 0 | 0 | 0,98010389 | 0,51990246 | TUBB6;TUBB3;GP1BB;CCNE1 |
| IL-6 signaling | 5/97 | 0,60146092 | 1 | 0 | 0 | 0,95989556 | 0,48800488 | SOCS3;CEBPB;CASP3;SGK1;EIF4E |
| FAS pathway a | 2/38 | 0,61274022 | 1 | 0 | 0 | 0,98010389 | 0,48006882 | CASP3;LMNB1 |
| SREBF and mi | 1/18 | 0,6298895 | 1 | 0 | 0 | 1,03455411 | 0,47818216 | HMGCR |
| Sulfation Biot | 1/18 | 0,6298895 | 1 | 0 | 0 | 1,03455411 | 0,47818216 | SULT2B1 |
| TarBasePathw | 1/18 | 0,6298895 | 1 | 0 | 0 | 1,03455411 | 0,47818216 | GJA1 |
| IL-1 signaling | 3/58 | 0,60896129 | 1 | 0 | 0 | 0,96320555 | 0,47775051 | IL1B;CCL2;IL1RAP |
| Alzheimers Di | 4/79 | 0,61910659 | 1 | 0 | 0 | 0,94288476 | 0,45209233 | CASP3;IL1B;BID;GAPDH |
| Kit receptor si | 3/59 | 0,62061363 | 1 | 0 | 0 | 0,94688003 | 0,45170586 | SOCS1;SNAI1;JUNB |

| | | | | | | | | |
|---|---|---|---|---|---|---|---|---|
| MicroRNAs in | 4/80 | 0,62903647 | 1 | 0 | 0 | 0,9310987 | 0,43162574 | VMP1;MYEF2;PIK3R3;GATA4 |
| Mitochondria | 1/19 | 0,64978227 | 1 | 0 | 0 | 0,98010389 | 0,42254038 | MYEF2 |
| Serotonin Rec | 1/19 | 0,64978227 | 1 | 0 | 0 | 0,98010389 | 0,42254038 | EGR1 |
| Interleukin-11 | 2/40 | 0,64079021 | 1 | 0 | 0 | 0,9310987 | 0,41438842 | SOCS3;BIRC5 |
| Oxidation by ( | 3/61 | 0,64319984 | 1 | 0 | 0 | 0,91583478 | 0,40415771 | CYB5R2;CYP26A1;CYP26B1 |
| Calcium Regu | 7/143 | 0,65279262 | 1 | 0 | 0 | 0,91156516 | 0,38877869 | CHRM3;GJA1;GJB4;RGS1;RGS20;ATP1B1;ADCY8 |
| G13 Signaling | 2/41 | 0,65420336 | 1 | 0 | 0 | 0,90838897 | 0,38546307 | RTKN;CIT |
| Structural Pat | 2/41 | 0,65420336 | 1 | 0 | 0 | 0,90838897 | 0,38546307 | IL1RAP;EIF4E |
| TWEAK Signal | 2/42 | 0,66721341 | 1 | 0 | 0 | 0,88676066 | 0,35882356 | CASP3;CCL2 |
| Regulation of | 7/146 | 0,67403544 | 1 | 0 | 0 | 0,89283437 | 0,35219868 | CXCL10;SOCS1;IL1B;PLK1;CCL3;PIK3R3;TREM1 |
| Glycerophosp | 1/21 | 0,68642033 | 1 | 0 | 0 | 0,88676066 | 0,3336571 | AGPS |
| Triacylglyceric | 1/21 | 0,68642033 | 1 | 0 | 0 | 0,88676066 | 0,3336571 | AGPS |
| Calcium Regu | 7/149 | 0,69444914 | 1 | 0 | 0 | 0,87485784 | 0,31900497 | CHRM3;GJA1;GJB4;RGS1;RGS20;ATP1B1;ADCY8 |
| Proteasome D | 3/66 | 0,69543165 | 1 | 0 | 0 | 0,84645336 | 0,30745095 | H2AFZ;H2AFX;HIST1H2AE |
| Glutathione m | 1/22 | 0,70327719 | 1 | 0 | 0 | 0,84645336 | 0,29795512 | GPX2 |
| Oxidative Stre | 1/22 | 0,70327719 | 1 | 0 | 0 | 0,84645336 | 0,29795512 | JUNB |
| miR-targeted | 8/172 | 0,71097297 | 1 | 0 | 0 | 0,86613832 | 0,29545786 | GALNT7;SLC38A1;PKM;ANXA2;P4HA2;HOXA7;SLC38A2;MET |
| Non-odorant | 12/256 | 0,72638446 | 1 | 0 | 0 | 0,87290503 | 0,27904666 | CCR1;CHRM3;PROKR2;GPRC5A;CXCR2;F2R;C3AR1;FZD9;LPAR2;CXCR4;LPAR3;CELSR1 |
| MAPK signalir | 7/155 | 0,73272089 | 1 | 0 | 0 | 0,84099237 | 0,26154057 | NR4A1;DUSP5;TMEM37;CASP3;IL1B;DUSP6;CDC25B |
| Histone Modi | 3/70 | 0,73288889 | 1 | 0 | 0 | 0,7980846 | 0,24801371 | HIST4H4;HIST1H3G;HIST1H4I |
| IL-9 Signaling | 1/24 | 0,73432338 | 1 | 0 | 0 | 0,77591558 | 0,23960722 | SOCS3 |
| Triacylglyceric | 1/24 | 0,73432338 | 1 | 0 | 0 | 0,77591558 | 0,23960722 | AGPS |
| TNF-alpha NF | 8/179 | 0,75124111 | 1 | 0 | 0 | 0,83226699 | 0,23805218 | UNC5CL;CRADD;PEG3;FANCD2;CASP3;PFDN2;MCC;TRAIP |
| miRNAs involv | 3/71 | 0,74166445 | 1 | 0 | 0 | 0,78684397 | 0,2351549 | CDKN1A;CCNE1;H2AFX |
| Alzheimers Di | 3/73 | 0,75852556 | 1 | 0 | 0 | 0,7652866 | 0,21150898 | CASP3;IL1B;BID |
| EPO Receptor | 1/26 | 0,76212388 | 1 | 0 | 0 | 0,71622977 | 0,19456107 | SOCS1 |
| EPO Receptor | 1/26 | 0,76212388 | 1 | 0 | 0 | 0,71622977 | 0,19456107 | SOCS1 |
| Glutathione a | 1/26 | 0,76212388 | 1 | 0 | 0 | 0,71622977 | 0,19456107 | AHCY |
| Synaptic Vesic | 2/51 | 0,76710384 | 1 | 0 | 0 | 0,73027349 | 0,19361968 | SLC38A1;STX1A |
| Wnt Signaling | 2/51 | 0,76710384 | 1 | 0 | 0 | 0,73027349 | 0,19361968 | GJA1;LEF1 |
| FSH signaling | 1/27 | 0,77491438 | 1 | 0 | 0 | 0,68970274 | 0,17587608 | SGK1 |
| GPCRs, Other | 3/77 | 0,78956997 | 1 | 0 | 0 | 0,72553145 | 0,17141901 | GPR39;FZD9;CELSR1 |
| Nanoparticle- | 1/28 | 0,78701774 | 1 | 0 | 0 | 0,6650705 | 0,15928737 | FN1 |
| RANKL/RANK | 2/54 | 0,79407736 | 1 | 0 | 0 | 0,68970274 | 0,15902779 | FHL2;TNFRSF11B |
| GPCRs, Class | 7/170 | 0,81336292 | 1 | 0 | 0 | 0,76678716 | 0,15840126 | CCR1;CHRM3;F2R;CXCR2;C3AR1;CXCR4;GCAT |
| Cytoplasmic R | 5/127 | 0,8184163 | 1 | 0 | 0 | 0,73314858 | 0,14691135 | RPS6KA6;SNORD32A;RPL34;SNORD83B;RPL37 |
| Toll-like recep | 4/104 | 0,81617008 | 1 | 0 | 0 | 0,71622977 | 0,14548956 | CXCL10;IL1B;CCL3;PIK3R3 |
| MAPK Signalir | 7/173 | 0,82698442 | 1 | 0 | 0 | 0,75349028 | 0,14314012 | NR4A1;DUSP5;TMEM37;CASP3;IL1B;DUSP6;CDC25B |
| Delta-Notch S | 3/81 | 0,81722371 | 1 | 0 | 0 | 0,68970274 | 0,13921126 | NOV;LEF1;SKP2 |
| Hypothetical I | 1/30 | 0,80930858 | 1 | 0 | 0 | 0,62073246 | 0,13133147 | ADCY8 |
| IL17 signaling | 1/30 | 0,80930858 | 1 | 0 | 0 | 0,62073246 | 0,13133147 | CEBPB |
| TCA Cycle(Mu | 1/30 | 0,80930858 | 1 | 0 | 0 | 0,62073246 | 0,13133147 | PDK3 |
| MicroRNAs in | 4/109 | 0,84343345 | 1 | 0 | 0 | 0,68337519 | 0,11636122 | MYEF2;MIR140;PIK3R3;GATA4 |
| SIDS Susceptil | 2/59 | 0,83292229 | 1 | 0 | 0 | 0,63125335 | 0,11540254 | SCN5A;FOXM1 |
| Hypothetical I | 1/32 | 0,82926838 | 1 | 0 | 0 | 0,58193669 | 0,1089452 | ADCY8 |
| Monoamine G | 1/32 | 0,82926838 | 1 | 0 | 0 | 0,58193669 | 0,1089452 | CHRM3 |
| Oxidative Stre | 1/32 | 0,82926838 | 1 | 0 | 0 | 0,58193669 | 0,1089452 | JUNB |
| Statin Pathwa | 1/32 | 0,82926838 | 1 | 0 | 0 | 0,58193669 | 0,1089452 | HMGCR |
| TNF alpha Sig | 3/86 | 0,84738942 | 1 | 0 | 0 | 0,64960374 | 0,10757108 | CASP3;PLK1;BID |
| Alpha 6 Beta 4 | 1/33 | 0,83845127 | 1 | 0 | 0 | 0,56430224 | 0,09942939 | LAMA5 |
| Fatty Acid Bet | 1/33 | 0,83845127 | 1 | 0 | 0 | 0,56430224 | 0,09942939 | TPI1 |
| Monoamine G | 1/33 | 0,83845127 | 1 | 0 | 0 | 0,56430224 | 0,09942939 | CHRM3 |
| Toll Like Rece | 1/33 | 0,83845127 | 1 | 0 | 0 | 0,56430224 | 0,09942939 | CASP3 |

| | | | | | | | | |
|---|---|---|---|---|---|---|---|---|
| Fatty Acid Bet | 1/34 | 0,84714069 | 1 | 0 | 0 | 0,54770512 | 0,09085798 | TPI1 |
| Signaling of H | 1/34 | 0,84714069 | 1 | 0 | 0 | 0,54770512 | 0,09085798 | MET |
| ESC Pluripote | 4/116 | 0,87581627 | 1 | 0 | 0 | 0,64213703 | 0,0851467 | WNT7B;FZD9;WNT7A;SMAD7 |
| Endothelin Pa | 1/35 | 0,85536312 | 1 | 0 | 0 | 0,5320564 | 0,08312275 | GUCY1B2 |
| Regulation of | 5/144 | 0,89149567 | 1 | 0 | 0 | 0,64659632 | 0,07426463 | CHRM3;F2R;FN1;PIK3R3;EZR |
| PPAR signalin | 2/68 | 0,88652282 | 1 | 0 | 0 | 0,54770512 | 0,06597021 | ACOX2;OLR1 |
| Regulation of | 5/148 | 0,90445635 | 1 | 0 | 0 | 0,62912074 | 0,06317708 | CHRM3;F2R;FN1;PIK3R3;EZR |
| IL-5 Signaling | 1/40 | 0,89030255 | 1 | 0 | 0 | 0,46554935 | 0,05409401 | SPRED1 |
| miR-targeted | 1/41 | 0,89620501 | 1 | 0 | 0 | 0,45419449 | 0,0497734 | CEBPB |
| Cytoplasmic R | 2/74 | 0,912882 | 1 | 0 | 0 | 0,50329659 | 0,04587481 | RPL34;RPL37 |
| Estrogen sign | 2/74 | 0,912882 | 1 | 0 | 0 | 0,50329659 | 0,04587481 | POLR2F;BRCA1 |
| Eukaryotic Tra | 1/42 | 0,90179015 | 1 | 0 | 0 | 0,44338033 | 0,04583375 | POLR2F |
| IL-2 Signaling | 1/42 | 0,90179015 | 1 | 0 | 0 | 0,44338033 | 0,04583375 | SOCS3 |
| Splicing facto | 1/42 | 0,90179015 | 1 | 0 | 0 | 0,44338033 | 0,04583375 | CHL1 |
| mRNA Proces | 4/137 | 0,94061238 | 1 | 0 | 0 | 0,54370727 | 0,03328802 | SNRPD1;PRMT1;SNRPG;HNRNPA1 |
| Translation Fa | 1/46 | 0,92128499 | 1 | 0 | 0 | 0,40482552 | 0,03318997 | EIF4E |
| Allograft Reje | 2/82 | 0,9391711 | 1 | 0 | 0 | 0,45419449 | 0,02850416 | CASP3;IL1B |
| Selenium Met | 1/48 | 0,92953042 | 1 | 0 | 0 | 0,38795779 | 0,0283503 | GPX2 |
| TCR Signaling | 2/89 | 0,95581955 | 1 | 0 | 0 | 0,41847132 | 0,0189091 | PSTPIP1;IL1B |
| Translation Fa | 1/60 | 0,9637284 | 1 | 0 | 0 | 0,31036623 | 0,01146672 | EIF4E |
| Notch Signali | 1/61 | 0,96568192 | 1 | 0 | 0 | 0,30527826 | 0,01066055 | CDKN1A |
| SREBP signalli | 1/64 | 0,97093404 | 1 | 0 | 0 | 0,29096834 | 0,00858262 | HMGCR |
| IL-5 Signaling | 1/68 | 0,97670919 | 1 | 0 | 0 | 0,27385256 | 0,0064537 | SOCS1 |
| GPCRs, Class | 7/259 | 0,98727531 | 1 | 0 | 0 | 0,50329659 | 0,00644539 | CCR1;GPR39;CHRM3;CXCR2;F2R;C3AR1;CXCR4 |
| mRNA proces | 11/398 | 0,99603849 | 1 | 0 | 0 | 0,51467767 | 0,00204295 | BARD1;LSM8;NPM1;MYEF2;SNRPD1;PCBP4;PRMT1;RPL37;BRCA1;EIF4E;WDR55 |
| G Protein Sig | 1/88 | 0,99230978 | 1 | 0 | 0 | 0,21161334 | 0,00163364 | ADCY8 |
| B Cell Recept | 1/93 | 0,99417128 | 1 | 0 | 0 | 0,20023628 | 0,00117054 | ILF2 |
| G Protein Sig | 1/96 | 0,99506432 | 1 | 0 | 0 | 0,1939789 | 9,60E-04 | ADCY8 |
| Integrin-medi | 1/97 | 0,9953305 | 1 | 0 | 0 | 0,19197911 | 8,99E-04 | ITGA9 |
| Integrin-medi | 1/100 | 0,99604598 | 1 | 0 | 0 | 0,18621974 | 7,38E-04 | ITGA9 |
| Odorant GPC | 1/122 | 0,99883208 | 1 | 0 | 0 | 0,15263913 | 1,78E-04 | GPR142 |

**Table S4.** ***MMTV-R26*** $^{Met}$ **tumour versus controls.**

| index | gene name | pvalue | log2FC |
|---|---|---|---|
| 702 | Foxm1 | 0.0236012658 | 1.41555447003448 |
| 459 | Aurka | 0.0030585754 | 1.81423953455477 |
| 544 | Aurkb | 0.0189702518 | 1.65732268945046 |
| 465 | Cdk1 | 0.0119652018 | 1.81069099796158 |
| 2243 | Cdk2 | 0.0068316319 | 0.657367566261536 |
| 1178 | Cdk4 | 0.0057604442 | 1.0252517662403 |
| 4394 | Cdk6 | 0.3377133336 | 0.350047272557211 |
| 7210 | Cdk5 | 0.6642639085 | -0.195680364197381 |
| 7604 | Cdk3-ps | 0.7824790071 | -0.162854424239089 |
| 2630 | Wee1 | 0.1340048910 | 0.584243867281207 |
| 2791 | Ccna1 | 0.1175335340 | -0.869644269247914 |
| 717 | Ccna2 | 0.0385074205 | 1.39925211687813 |
| 581 | Ccnb1 | 0.0302622659 | 1.57829056186044 |
| 634 | Ccnb2 | 0.0222839844 | 1.4964121428095 |
| 1092 | Ccnd1 | 0.0763555189 | 1.07948033011888 |
| 1454 | Ccnd2 | 0.0009653828 | -1.7032664178437 |
| 6145 | Ccnd3 | 0.3000047990 | -0.293298972372172 |
| 671 | Ccne1 | 0.0145447076 | 1.44663741701661 |
| 3771 | Ccne2 | 0.3245121465 | 0.420617509929978 |
| 7726 | Ccng1 | 0.6626222740 | -0.154629679856764 |
| 7486 | Ccng2 | 0.8279623388 | 0.0840532742418599 |

**Table S5A. A11_vs_No.**

| ID | logFC | AveExpr | t | P.Value | adj.P.Val | B |
|---|---|---|---|---|---|---|
| B7-H3 | 0,322884 | -0,02672 | 2,05581 | 0,045767 | 0,999887 | -3,97403 |
| VHL-EPPK1 | -0,3044 | -0,01284 | -2,76152 | 0,008363 | 0,999887 | -3,31209 |
| EMA | -0,24786 | 0,022802 | -2,33677 | 0,02407 | 0,999887 | -3,72549 |
| HER3_pY12 | -0,15254 | 0,046123 | -2,23746 | 0,030372 | 0,999887 | -3,81583 |

cut-off : P.Value<0.05

**Table S5B. R547_vs No.**

| ID | logFC | AveExpr | t | P.Value | adj.P.Val | B |
|---|---|---|---|---|---|---|
| Rb_pS807_ | -2,26559 | -0,3406 | -10,164 | 4,06E-13 | 8,75E-11 | 19,76045 |
| c-Met_pY1 | -1,63916 | -0,11708 | -3,31643 | 0,001835 | 0,032951 | -1,91629 |
| Mcl-1 | -1,47577 | -0,13578 | -11,4027 | 1,01E-14 | 4,35E-12 | 23,37069 |
| NDRG1_pT | -1,35527 | 0,115251 | -2,61771 | 0,012093 | 0,115828 | -3,65773 |
| Connexin-4 | -1,28807 | -0,0405 | -4,16411 | 0,000143 | 0,0051 | 0,504974 |
| 53BP1 | -1,18178 | 0,044267 | -3,111 | 0,00327 | 0,050334 | -2,45599 |
| Akt_pS473 | -1,1062 | 0,043018 | -2,08676 | 0,04274 | 0,241278 | -4,77765 |
| ARID1A | -1,05097 | 0,008024 | -3,55905 | 0,000907 | 0,018618 | -1,25293 |
| Pdcd4 | -1,02392 | -0,38581 | -3,47551 | 0,001159 | 0,02172 | -1,48433 |
| Akt2_pS474 | -1,0155 | -0,16412 | -2,53926 | 0,014721 | 0,132182 | -3,83529 |
| Cdc6 | -1,00313 | 0,051176 | -6,1805 | 1,84E-07 | 2,64E-05 | 6,97937 |
| HER2_pY12 | -0,99549 | -0,03211 | -3,77354 | 0,000478 | 0,012771 | -0,64537 |
| Rad51 | -0,96045 | 0,041184 | -4,19764 | 0,000129 | 0,005091 | 0,606083 |
| Shc_pY317 | -0,94145 | -0,16483 | -3,02665 | 0,004123 | 0,057328 | -2,67128 |
| Hif-1-alpha | -0,8917 | -0,06664 | -5,43683 | 2,26E-06 | 0,000162 | 4,526962 |
| Cyclin-D3 | -0,77841 | -0,15553 | -3,7562 | 0,000504 | 0,012771 | -0,69516 |
| FN14 | -0,75776 | -0,03562 | -5,84998 | 5,61E-07 | 4,84E-05 | 5,884974 |
| EPHA2 | 0,750958 | 0,195028 | 3,173865 | 0,002745 | 0,045508 | -2,2931 |
| Src_pY416 | -0,7413 | -0,01775 | -2,46144 | 0,017831 | 0,153707 | -4,00744 |
| HES1 | -0,70096 | -0,08295 | -2,22051 | 0,031584 | 0,205169 | -4,51425 |
| p90RSK_pT | -0,65159 | 0,103213 | -4,63795 | 3,16E-05 | 0,001945 | 1,962754 |
| Pyk2_pY40 | -0,64559 | -0,01398 | -2,65085 | 0,011119 | 0,108912 | -3,58154 |
| HER2 | -0,63882 | -0,10338 | -3,96435 | 0,000267 | 0,008218 | -0,08997 |
| p38-MAPK | -0,62517 | -0,1338 | -3,06915 | 0,00367 | 0,052852 | -2,56329 |
| PHLPP | -0,61105 | 0,013041 | -2,85531 | 0,006538 | 0,074151 | -3,09655 |
| Coup-TFII | -0,60824 | -0,00264 | -2,18078 | 0,034592 | 0,207074 | -4,59387 |
| GCN5L2 | -0,59346 | -0,23753 | -2,12037 | 0,039652 | 0,227866 | -4,7127 |
| PLC-gamma | -0,58438 | -0,08715 | -2,49914 | 0,016257 | 0,142994 | -3,92454 |
| JNK_pT183 | -0,57219 | -0,13916 | -2,02355 | 0,04912 | 0,251844 | -4,89748 |
| ULK1_pS75 | -0,54843 | -0,1893 | -2,95175 | 0,005052 | 0,063966 | -2,85921 |
| Snail | -0,5303 | 0,250236 | -2,04292 | 0,047082 | 0,247468 | -4,86109 |
| NF-kB-p65_ | -0,52334 | -0,21469 | -2,44942 | 0,018362 | 0,155176 | -4,03368 |
| C-Raf_pS33 | -0,52271 | 0,00189 | -2,70278 | 0,009735 | 0,102341 | -3,46078 |
| Chk1 | -0,51207 | 0,162593 | -2,67078 | 0,010568 | 0,105925 | -3,5354 |
| Mnk1 | -0,49885 | -0,06276 | -2,06787 | 0,044566 | 0,243139 | -4,81378 |
| Smad1 | -0,49774 | -0,19949 | -2,28733 | 0,027045 | 0,194271 | -4,37778 |
| ERRalpha | -0,49309 | -0,0124 | -4,19553 | 0,00013 | 0,005091 | 0,599714 |
| PDK1_pS24 | -0,48156 | -0,08182 | -2,78229 | 0,007922 | 0,085362 | -3,27266 |
| Aurora-ABC | -0,47789 | 0,075737 | -3,7764 | 0,000474 | 0,012771 | -0,63713 |
| Histone-H3 | -0,47253 | 0,093741 | -2,67731 | 0,010393 | 0,105925 | -3,52021 |
| MMP14 | -0,45283 | 0,114206 | -2,24761 | 0,029667 | 0,205169 | -4,45927 |
| RBM15 | -0,41724 | 0,077333 | -2,86792 | 0,006323 | 0,073649 | -3,06581 |
| IRS1 | -0,39491 | -0,16165 | -2,02407 | 0,049064 | 0,251844 | -4,89652 |
| PI3K-p85 | -0,39342 | -0,11462 | -2,08293 | 0,043105 | 0,241278 | -4,78501 |
| DNMT1 | 0,374448 | 0,187822 | 3,149138 | 0,002941 | 0,046953 | -2,35741 |
| IR-b | -0,3613 | 0,194304 | -3,56096 | 0,000902 | 0,018618 | -1,24759 |
| RPA32_pS4 | -0,33419 | -0,05857 | -3,61927 | 0,000759 | 0,017218 | -1,08425 |
| 4E-BP1_pS | -0,33365 | -0,09775 | -2,80358 | 0,007493 | 0,082804 | -3,22163 |
| SOD1 | 0,33142 | 0,07373 | 4,141726 | 0,000154 | 0,0051 | 0,437667 |

| | | | | | | |
|---|---|---|---|---|---|---|
| MITF | -0,32289 | -0,12981 | -3,73257 | 0,000541 | 0,012952 | -0,76284 |
| PKC-b-II_p! | -0,31958 | -0,02681 | -2,59914 | 0,012673 | 0,118745 | -3,70013 |
| SHP2 | -0,29991 | -0,06484 | -5,99037 | 3,49E-07 | 3,76E-05 | 6,349228 |
| HER3_pY12 | -0,29196 | 0,046123 | -4,28241 | 9,88E-05 | 0,004939 | 0,863208 |
| S6 | 0,2893 | 0,054635 | 2,355064 | 0,023047 | 0,182137 | -4,2362 |
| MSI2 | 0,286749 | -0,04978 | 3,494525 | 0,001096 | 0,02148 | -1,43193 |
| Wee1_pS6 | -0,24814 | 0,100067 | -2,91346 | 0,0056 | 0,067042 | -2,95408 |
| Akt2 | -0,24708 | -0,17563 | -2,0576 | 0,045587 | 0,24326 | -4,83331 |
| Atg4B | -0,23819 | -0,10914 | -3,22048 | 0,002409 | 0,041526 | -2,17102 |
| GSK-3a-b | -0,23404 | 0,00883 | -2,35012 | 0,023319 | 0,182137 | -4,24663 |
| CD45 | 0,225285 | 0,008777 | 3,068295 | 0,003679 | 0,052852 | -2,56547 |
| Bcl2 | 0,217647 | 0,084614 | 2,125147 | 0,03923 | 0,227866 | -4,70341 |
| CD49b | 0,179144 | 0,083165 | 2,941456 | 0,005194 | 0,063966 | -2,8848 |
| Bcl-xL | -0,17639 | -0,0211 | -2,07496 | 0,043873 | 0,242425 | -4,80025 |
| HSP27 | 0,169276 | 0,015585 | 2,993791 | 0,004509 | 0,060053 | -2,75411 |
| Annexin-VI | 0,165943 | 0,064325 | 2,433534 | 0,019085 | 0,158184 | -4,0682 |
| Notch1-cle | 0,165721 | 0,058174 | 2,343927 | 0,023665 | 0,182137 | -4,25969 |
| PAI-1 | 0,165641 | 0,03994 | 4,268752 | 0,000103 | 0,004939 | 0,821628 |
| SF2 | 0,161461 | 0,101404 | 2,29015 | 0,026866 | 0,194271 | -4,37194 |
| Rheb | 0,154196 | 0,01574 | 2,98661 | 0,004598 | 0,060053 | -2,77213 |
| ATP5A | 0,153526 | 0,072204 | 2,198071 | 0,033253 | 0,205169 | -4,55936 |
| 14-3-3-eps | 0,151653 | 0,027058 | 2,584417 | 0,013151 | 0,120599 | -3,73357 |
| CD29 | 0,150243 | 0,064657 | 2,311589 | 0,025546 | 0,189836 | -4,32744 |
| MEK2 | 0,145272 | -0,0442 | 2,182739 | 0,034438 | 0,207074 | -4,58997 |
| Gli1 | 0,14334 | -0,00908 | 2,197165 | 0,033322 | 0,205169 | -4,56117 |
| MelanA | 0,141021 | -0,03267 | 2,170769 | 0,035389 | 0,208942 | -4,61375 |
| Bid | 0,139832 | 0,076547 | 2,233226 | 0,030671 | 0,205169 | -4,48852 |
| Tau | 0,139304 | 0,086972 | 2,056308 | 0,045717 | 0,24326 | -4,83577 |
| Rb | 0,135846 | 0,097094 | 2,40044 | 0,020674 | 0,168122 | -4,13957 |
| UQCRC2 | 0,134148 | 0,034731 | 2,201867 | 0,032965 | 0,205169 | -4,55175 |
| Collagen-V | 0,13084 | 0,014401 | 2,231359 | 0,030803 | 0,205169 | -4,49231 |
| BiP-GRP78 | 0,127991 | 0,042868 | 2,24745 | 0,029678 | 0,205169 | -4,45961 |
| CD31 | 0,125889 | 0,029025 | 2,209031 | 0,032428 | 0,205169 | -4,53737 |
| Syk | 0,123404 | 0,060814 | 2,331125 | 0,024395 | 0,184457 | -4,2866 |
| TIGAR | 0,122938 | -0,03684 | 2,208326 | 0,032481 | 0,205169 | -4,53878 |
| ADAR1 | 0,122292 | 0,069972 | 2,018462 | 0,049668 | 0,251844 | -4,907 |

cut-off : P.Value<0.05

**Table S5C. Adav_vs_No.**

| ID | logFC | AveExpr | t | P.Value | adj.P.Val | B |
|---|---|---|---|---|---|---|
| p53 | 0,893274 | 0,016506 | 2,733352 | 0,008997 | 0,271392 | -3,02026 |
| cdc2_pY15 | -0,88263 | 0,010496 | -9,99755 | 6,77E-13 | 2,92E-10 | 19,00459 |
| Pyk2_pY40 | -0,85188 | -0,01398 | -3,49786 | 0,001086 | 0,066855 | -1,08732 |
| Chk1 | -0,8297 | 0,162593 | -4,32739 | 8,56E-05 | 0,009228 | 1,290798 |
| p38-MAPK- | 0,787398 | -0,03056 | 2,894573 | 0,00589 | 0,271392 | -2,63827 |
| CDK1_pT14 | 0,780187 | -0,00726 | 2,727352 | 0,009138 | 0,271392 | -3,03418 |
| Snail | -0,68821 | 0,250236 | -2,65124 | 0,011108 | 0,281613 | -3,20892 |
| Rad51 | -0,63621 | 0,041184 | -2,78055 | 0,007958 | 0,271392 | -2,91 |
| JNK2 | 0,631696 | -0,12274 | 2,102989 | 0,041224 | 0,668733 | -4,35644 |
| Hif-1-alpha | -0,61398 | -0,06664 | -3,74354 | 0,000523 | 0,045111 | -0,40843 |
| MMP14 | -0,49777 | 0,114206 | -2,47066 | 0,017434 | 0,385051 | -3,60906 |
| EMA | -0,46177 | 0,022802 | -4,35352 | 7,88E-05 | 0,009228 | 1,369238 |
| Mcl-1 | -0,4581 | -0,13578 | -3,53955 | 0,000961 | 0,066855 | -0,97383 |
| RPA32 | -0,36247 | -0,0536 | -2,85045 | 0,006622 | 0,271392 | -2,74429 |
| RBM15 | 0,353431 | 0,077333 | 2,429356 | 0,019279 | 0,395682 | -3,69765 |
| ERRalpha | -0,3129 | -0,0124 | -2,66236 | 0,010797 | 0,281613 | -3,18361 |
| VHL-EPPK1 | -0,28053 | -0,01284 | -2,54495 | 0,014514 | 0,347533 | -3,44696 |
| PARP | -0,26728 | -0,03808 | -2,46061 | 0,017868 | 0,385051 | -3,63073 |
| SHP2 | -0,24647 | -0,06484 | -4,92299 | 1,24E-05 | 0,002681 | 3,117465 |
| DDB-1 | -0,23711 | 0,062349 | -2,09577 | 0,041893 | 0,668733 | -4,37016 |
| c-Abl_pY41 | -0,22358 | 0,002527 | -2,71453 | 0,009445 | 0,271392 | -3,06385 |
| SOD1 | 0,218735 | 0,07373 | 2,733509 | 0,008993 | 0,271392 | -3,0199 |
| HER3_pY12 | -0,2131 | 0,046123 | -3,12582 | 0,003138 | 0,169078 | -2,06519 |
| MITF | -0,20391 | -0,12981 | -2,35725 | 0,022927 | 0,449157 | -3,84954 |
| Raptor | -0,14889 | -0,08896 | -2,18487 | 0,034271 | 0,596674 | -4,19821 |
| C-Raf | -0,14113 | 0,036654 | -2,04081 | 0,0473 | 0,728084 | -4,47329 |
| MIG6 | 0,125608 | 0,101079 | 2,180557 | 0,03461 | 0,596674 | -4,20667 |
| PAI-1 | 0,085934 | 0,03994 | 2,214627 | 0,032014 | 0,596674 | -4,13952 |

cut-off : P.Value<0.05

**Table S5D. A11+R547_vs_No.**

| ID | logFC | AveExpr | t | P.Value | adj.P.Val | B |
|---|---|---|---|---|---|---|
| Rb_pS807_ | -2,29183 | -0,3406 | -10,2817 | 2,83E-13 | 1,22E-10 | 20,1024 |
| c-Met_pY1 | -2,10188 | -0,11708 | -4,25263 | 0,000109 | 0,001732 | 0,821176 |
| Connexin-4 | -1,72293 | -0,0405 | -5,56994 | 1,44E-06 | 7,77E-05 | 5,002576 |
| NDRG1_pT | -1,6529 | 0,115251 | -3,19258 | 0,002605 | 0,016512 | -2,19027 |
| Histone-H3 | 1,598981 | 0,365476 | 4,665457 | 2,89E-05 | 0,000595 | 2,095144 |
| Akt_pS473 | -1,52954 | 0,043018 | -2,88535 | 0,006036 | 0,030131 | -2,96795 |
| Akt2_pS47 | -1,435 | -0,16412 | -3,58822 | 0,000832 | 0,008152 | -1,11931 |
| HER2_pY12 | -1,33878 | -0,03211 | -5,07481 | 7,54E-06 | 0,000271 | 3,395449 |
| 53BP1 | -1,24197 | 0,044267 | -3,26945 | 0,002097 | 0,013907 | -1,98797 |
| Stat3 | -1,17506 | -0,32293 | -2,81043 | 0,007359 | 0,034105 | -3,14962 |
| SHP-2_pY5 | -1,14238 | -0,1488 | -2,83113 | 0,006969 | 0,033108 | -3,09976 |
| Rad51 | -1,14162 | 0,041184 | -4,98944 | 1E-05 | 0,000308 | 3,121753 |
| Mcl-1 | -1,10592 | -0,13578 | -8,545 | 6,78E-11 | 1,46E-08 | 14,75228 |
| Akt1_pS47 | -1,09595 | -0,06283 | -2,87423 | 0,006217 | 0,030317 | -2,99511 |
| ARID1A | -1,08243 | 0,008024 | -3,6656 | 0,000661 | 0,006627 | -0,90172 |
| Rictor | -1,06039 | -0,12713 | -3,07822 | 0,00358 | 0,020301 | -2,48566 |
| Stat5a | -1,05491 | -0,3732 | -2,13957 | 0,037978 | 0,09966 | -4,61741 |
| Shc_pY317 | -1,05035 | -0,16483 | -3,37677 | 0,001543 | 0,01143 | -1,70071 |
| Hif-1-alpha | -1,04831 | -0,06664 | -6,39173 | 8,97E-08 | 9,67E-06 | 7,713684 |
| Src_pY416 | -1,00952 | -0,01775 | -3,35204 | 0,001657 | 0,011708 | -1,76739 |
| Pdcd4 | -0,99856 | -0,38581 | -3,38943 | 0,001488 | 0,011384 | -1,66646 |
| Cdc6 | -0,98756 | 0,051176 | -6,08459 | 2,54E-07 | 2,19E-05 | 6,696983 |
| NF-kB-p65_ | -0,95184 | -0,21469 | -4,455 | 5,7E-05 | 0,000961 | 1,440272 |
| FAK_pY397 | -0,95056 | -0,14486 | -2,90697 | 0,005698 | 0,028891 | -2,91493 |
| Caspase-3- | 0,94917 | 0,182884 | 3,392577 | 0,001475 | 0,011384 | -1,65794 |
| HES1 | -0,93768 | -0,08295 | -2,9704 | 0,004804 | 0,025252 | -2,75781 |
| PTEN | -0,93694 | -0,31196 | -2,43659 | 0,018944 | 0,062306 | -4,0047 |
| Cyclin-D3 | -0,92211 | -0,15553 | -4,44962 | 5,8E-05 | 0,000961 | 1,423672 |
| Pyk2_pY40 | -0,90613 | -0,01398 | -3,72063 | 0,000561 | 0,006041 | -0,74549 |
| PHLPP | -0,89112 | 0,013041 | -4,164 | 0,000143 | 0,002132 | 0,553697 |
| FOXO3 | -0,86716 | -0,14293 | -2,51256 | 0,015728 | 0,055943 | -3,83818 |
| HER2 | -0,86001 | -0,10338 | -5,33703 | 3,15E-06 | 0,000136 | 4,242825 |
| H2AX_pS14 | 0,853326 | 0,173534 | 4,894752 | 1,37E-05 | 0,000368 | 2,819654 |
| Coup-TFII | -0,84421 | -0,00264 | -3,02683 | 0,004121 | 0,022204 | -2,61617 |
| FRS2-a_pY | -0,82822 | -0,13486 | -2,49132 | 0,016573 | 0,056243 | -3,88512 |
| p70-S6K1 | -0,82207 | -0,31847 | -3,15838 | 0,002867 | 0,017401 | -2,27932 |
| IRS2 | -0,81457 | -0,1149 | -2,65633 | 0,010965 | 0,041448 | -3,51276 |
| mTOR | -0,77909 | -0,09916 | -2,76846 | 0,008213 | 0,035399 | -3,24995 |
| JNK_pT183 | -0,76043 | -0,13916 | -2,68926 | 0,010079 | 0,0406 | -3,43638 |
| PRAS40_pT | -0,75448 | -0,16259 | -2,88251 | 0,006082 | 0,030131 | -2,9749 |
| S6 | 0,736784 | 0,054635 | 5,99783 | 3,41E-07 | 2,45E-05 | 6,410296 |
| DVL3 | -0,73359 | -0,25384 | -2,66601 | 0,010697 | 0,041448 | -3,4904 |
| ULK1_pS75 | -0,72225 | -0,1893 | -3,88731 | 0,000338 | 0,00394 | -0,26522 |
| WIPI1 | -0,71426 | -0,2543 | -2,32003 | 0,025043 | 0,07253 | -4,25255 |
| GCN5L2 | -0,6997 | -0,23753 | -2,49994 | 0,016225 | 0,055943 | -3,8661 |
| FN14 | -0,69583 | -0,03562 | -5,37186 | 2,8E-06 | 0,000134 | 4,356063 |
| Caspase-7- | 0,690076 | 0,19787 | 5,007771 | 9,41E-06 | 0,000308 | 3,180425 |
| p38-MAPK | -0,68958 | -0,1338 | -3,38535 | 0,001506 | 0,011384 | -1,67749 |
| EPHA2 | 0,66279 | 0,195028 | 2,801229 | 0,007539 | 0,034412 | -3,17171 |

| | | | | | | |
|---|---|---|---|---|---|---|
| Myosin-IIa | -0,6588 | -0,20646 | -2,05618 | 0,04573 | 0,115938 | -4,77794 |
| Smad1 | -0,65278 | -0,19949 | -2,99984 | 0,004436 | 0,023603 | -2,68412 |
| XIAP | -0,64464 | -0,31104 | -2,50512 | 0,016019 | 0,055943 | -3,85467 |
| AMPKa_pT | 0,637662 | 0,086728 | 2,402575 | 0,020568 | 0,064501 | -4,07799 |
| PLC-gamma | -0,62274 | -0,08715 | -2,6632 | 0,010774 | 0,041448 | -3,49688 |
| PDK1 | -0,61755 | -0,14864 | -2,74466 | 0,008737 | 0,036812 | -3,30639 |
| Akt2 | -0,6143 | -0,17563 | -5,11559 | 6,58E-06 | 0,000258 | 3,52657 |
| Akt_pT308 | -0,60106 | 0,080566 | -2,24855 | 0,029602 | 0,082848 | -4,39988 |
| DM-Histon | 0,589655 | 0,167256 | 4,114998 | 0,000167 | 0,002229 | 0,406843 |
| DUSP6 | -0,58146 | 0,009088 | -2,04799 | 0,046561 | 0,116331 | -4,79343 |
| Chk1_pS34 | -0,57914 | -0,24295 | -2,40184 | 0,020604 | 0,064501 | -4,07956 |
| Rad23A | -0,57874 | -0,07387 | -3,16779 | 0,002792 | 0,017192 | -2,25487 |
| Jak2 | -0,57759 | -0,22193 | -2,2394 | 0,030237 | 0,084077 | -4,41848 |
| RRM2 | -0,57091 | -0,05838 | -2,45146 | 0,018271 | 0,060575 | -3,9724 |
| PDK1_pS24 | -0,56093 | -0,08182 | -3,24084 | 0,002274 | 0,01463 | -2,0636 |
| ERRalpha | -0,55114 | -0,0124 | -4,68943 | 2,67E-05 | 0,000595 | 2,170383 |
| mTOR_pS2 | -0,53647 | -0,13298 | -2,05253 | 0,046099 | 0,116191 | -4,78486 |
| Mnk1 | -0,52864 | -0,06276 | -2,19138 | 0,033766 | 0,091854 | -4,51508 |
| PTPN12 | -0,51905 | -0,19282 | -2,0303 | 0,048401 | 0,117209 | -4,82671 |
| UBAC1 | -0,51671 | -0,16793 | -2,14466 | 0,037544 | 0,099476 | -4,60744 |
| SOD1 | 0,514315 | 0,07373 | 6,427349 | 7,95E-08 | 9,67E-06 | 7,831688 |
| 4E-BP1_pT | -0,50097 | -0,04807 | -3,67529 | 0,000642 | 0,006591 | -0,87431 |
| TAZ | -0,4996 | -0,1195 | -3,11465 | 0,003237 | 0,019377 | -2,3923 |
| H2AX_pS13 | 0,486855 | 0,008345 | 2,326162 | 0,024683 | 0,072369 | -4,23976 |
| FoxO3a_pS | -0,48521 | -0,12973 | -2,13753 | 0,038153 | 0,09966 | -4,62141 |
| Rictor_pT1 | -0,47208 | -0,05303 | -3,10129 | 0,003359 | 0,019563 | -2,4266 |
| Chk1 | -0,45662 | 0,162593 | -2,38155 | 0,021634 | 0,065204 | -4,12291 |
| eEF2K | -0,4502 | -0,05665 | -2,09476 | 0,041987 | 0,107079 | -4,70433 |
| PI3K-p85 | -0,44397 | -0,11462 | -2,35055 | 0,023296 | 0,069245 | -4,18857 |
| PAK4 | -0,43814 | -0,11223 | -3,57622 | 0,000862 | 0,008259 | -1,1528 |
| DNMT1 | 0,418827 | 0,187822 | 3,522371 | 0,001011 | 0,008853 | -1,30247 |
| IRS1 | -0,41406 | -0,16165 | -2,12223 | 0,039487 | 0,10191 | -4,65122 |
| MMP14 | -0,41074 | 0,114206 | -2,03872 | 0,047517 | 0,117029 | -4,8109 |
| EMA | -0,40236 | 0,022802 | -3,79339 | 0,00045 | 0,004975 | -0,53714 |
| KAP1 | 0,376733 | 0,102915 | 2,143759 | 0,037621 | 0,099476 | -4,60921 |
| Caspase-8 | 0,365651 | 0,089702 | 4,624539 | 3,3E-05 | 0,000646 | 1,96701 |
| cdc2_pY15 | 0,365367 | 0,010496 | 4,138511 | 0,000155 | 0,002229 | 0,477218 |
| Erk5 | -0,34933 | -0,12646 | -3,95072 | 0,000278 | 0,003333 | -0,07982 |
| Lasu1 | -0,34757 | -0,10668 | -3,51686 | 0,001027 | 0,008853 | -1,31771 |
| Bcl2 | 0,344742 | 0,084614 | 3,366136 | 0,001591 | 0,01143 | -1,72941 |
| TRAP1 | 0,34178 | 0,101822 | 2,046688 | 0,046694 | 0,116331 | -4,79589 |
| MSI2 | 0,336344 | -0,04978 | 4,098924 | 0,000176 | 0,002229 | 0,358832 |
| SF2 | 0,328865 | 0,101404 | 4,664607 | 2,9E-05 | 0,000595 | 2,092478 |
| 4E-BP1_pS | -0,31499 | -0,09775 | -2,64677 | 0,011235 | 0,041448 | -3,53481 |
| RPA32_pS4 | -0,3135 | -0,05857 | -3,39528 | 0,001463 | 0,011384 | -1,65062 |
| Aurora-ABC | -0,30407 | 0,075737 | -2,40284 | 0,020555 | 0,064501 | -4,07742 |
| Akt1 | -0,29883 | -0,05668 | -2,50279 | 0,016111 | 0,055943 | -3,85981 |
| MITF | -0,29732 | -0,12981 | -3,43703 | 0,001296 | 0,010746 | -1,53703 |
| MLH1 | 0,294071 | 0,049452 | 2,812933 | 0,007311 | 0,034105 | -3,14361 |
| ATP5A | 0,293366 | 0,072204 | 4,200206 | 0,000128 | 0,001971 | 0,662682 |
| Cyclin-B1 | 0,291068 | 0,036954 | 2,663442 | 0,010768 | 0,041448 | -3,49633 |

| | | | | | | |
|---|---|---|---|---|---|---|
| SHP2 | -0,288 | -0,06484 | -5,7525 | 7,8E-07 | 4,8E-05 | 5,60165 |
| CD49b | 0,280556 | 0,083165 | 4,606587 | 3,5E-05 | 0,000655 | 1,910916 |
| IRF-1 | 0,280384 | 0,061071 | 3,683751 | 0,000626 | 0,006583 | -0,85033 |
| Caspase-8- | 0,27515 | 0,072456 | 2,747617 | 0,008671 | 0,036812 | -3,29939 |
| ADAR1 | 0,271711 | 0,069972 | 4,484658 | 5,18E-05 | 0,00093 | 1,531909 |
| Annexin-VI | 0,271659 | 0,064325 | 3,983839 | 0,000251 | 0,003096 | 0,017558 |
| GRB7 | 0,270155 | 0,054688 | 2,794121 | 0,007681 | 0,034412 | -3,18874 |
| Annexin-I | 0,269303 | 0,056642 | 2,359733 | 0,022792 | 0,068217 | -4,16918 |
| XPA | 0,265541 | -0,00526 | 2,654045 | 0,011029 | 0,041448 | -3,51804 |
| SCD | 0,265092 | 0,107627 | 3,2823 | 0,002022 | 0,013618 | -1,95385 |
| Cdc42 | 0,264264 | 0,085081 | 2,962677 | 0,004906 | 0,025474 | -2,77706 |
| Tau | 0,261725 | 0,086972 | 3,8634 | 0,000364 | 0,004127 | -0,33475 |
| Atg4B | -0,26122 | -0,10914 | -3,53186 | 0,000983 | 0,008853 | -1,27617 |
| Src | 0,256932 | 0,012489 | 2,319502 | 0,025074 | 0,07253 | -4,25366 |
| Rheb | 0,255832 | 0,01574 | 4,955205 | 1,12E-05 | 0,000322 | 3,012348 |
| Syk | 0,254503 | 0,060814 | 4,807616 | 1,82E-05 | 0,000461 | 2,543092 |
| Porin | 0,252939 | 0,069191 | 3,040147 | 0,003974 | 0,021681 | -2,58248 |
| NAPSIN-A | 0,250254 | -0,04255 | 3,311146 | 0,001863 | 0,012949 | -1,87701 |
| Collagen-V | 0,240595 | 0,014401 | 4,103134 | 0,000174 | 0,002229 | 0,371399 |
| HER3_pY12 | -0,23531 | 0,046123 | -3,45149 | 0,001243 | 0,010506 | -1,49753 |
| cdc25C | 0,235098 | 0,079258 | 2,165246 | 0,035836 | 0,095933 | -4,56694 |
| Rb | 0,23336 | 0,097094 | 4,123541 | 0,000163 | 0,002229 | 0,432391 |
| AMPK-a2_ | 0,229382 | -0,05915 | 2,646325 | 0,011247 | 0,041448 | -3,53584 |
| p21 | 0,226293 | 0,021595 | 2,271983 | 0,028032 | 0,079486 | -4,35198 |
| Notch1-cle | 0,224098 | 0,058174 | 3,169607 | 0,002778 | 0,017192 | -2,25014 |
| 14-3-3-bet | 0,220424 | 0,061595 | 3,087975 | 0,003485 | 0,020026 | -2,46073 |
| CD29 | 0,21904 | 0,064657 | 3,370087 | 0,001573 | 0,01143 | -1,71875 |
| N-Ras | 0,209379 | 0,051875 | 3,062862 | 0,003734 | 0,020633 | -2,52481 |
| D-a-Tubuli | 0,203756 | 0,028271 | 3,395323 | 0,001463 | 0,011384 | -1,6505 |
| CD31 | 0,20223 | 0,029025 | 3,548618 | 0,000935 | 0,008765 | -1,22968 |
| Glutamate- | 0,201878 | -0,02475 | 2,184328 | 0,034314 | 0,092432 | -4,52912 |
| XBP-1 | 0,201576 | 0,02769 | 3,250042 | 0,002216 | 0,01447 | -2,03931 |
| BiP-GRP78 | 0,200515 | 0,042868 | 3,520935 | 0,001015 | 0,008853 | -1,30644 |
| DNA-Ligase | 0,198414 | 0,085852 | 2,033771 | 0,048035 | 0,117209 | -4,8202 |
| Patched | 0,1962 | -0,01103 | 2,210008 | 0,032355 | 0,088823 | -4,4778 |
| GATA6 | 0,192752 | 0,04835 | 2,580979 | 0,013265 | 0,048451 | -3,68498 |
| Gli1 | 0,191127 | -0,00908 | 2,92966 | 0,005362 | 0,027511 | -2,85898 |
| RIP3 | 0,187771 | 0,039092 | 2,340856 | 0,023838 | 0,070372 | -4,20896 |
| HSP27 | 0,185949 | 0,015585 | 3,288678 | 0,001986 | 0,013586 | -1,9369 |
| DM-K9-His | 0,184723 | -0,00753 | 2,030252 | 0,048406 | 0,117209 | -4,8268 |
| PAI-1 | 0,183096 | 0,03994 | 4,718609 | 2,43E-05 | 0,000582 | 2,262134 |
| DAPK2 | 0,181077 | 0,015904 | 2,495172 | 0,016417 | 0,056155 | -3,87664 |
| Slfn11 | 0,180745 | -0,01853 | 2,189827 | 0,033886 | 0,091854 | -4,51817 |
| MRAP | 0,180454 | 0,053994 | 2,871651 | 0,00626 | 0,030317 | -3,00141 |
| Chk1_pS29 | 0,179698 | 0,044032 | 2,66379 | 0,010758 | 0,041448 | -3,49552 |
| PR | 0,177004 | 0,048848 | 2,792653 | 0,00771 | 0,034412 | -3,19225 |
| YAP | 0,174133 | -0,02923 | 2,408514 | 0,020276 | 0,064501 | -4,06525 |
| FOXM1 | 0,174033 | -0,00113 | 2,229208 | 0,030957 | 0,085528 | -4,43912 |
| PD-1 | 0,173112 | 0,011749 | 2,294632 | 0,026585 | 0,076389 | -4,30532 |
| TIGAR | 0,173096 | -0,03684 | 3,109319 | 0,003285 | 0,019396 | -2,406 |
| Smad4 | 0,172685 | 0,042306 | 3,070192 | 0,00366 | 0,020485 | -2,50614 |

| | | | | | | |
|---|---|---|---|---|---|---|
| PCNA | 0,171705 | 0,071675 | 2,124914 | 0,03925 | 0,101909 | -4,64601 |
| UQCRC2 | 0,170038 | 0,034731 | 2,79095 | 0,007745 | 0,034412 | -3,19632 |
| CD134 | 0,169887 | 0,019186 | 2,694285 | 0,00995 | 0,040458 | -3,42468 |
| Ambra1_pS | 0,169089 | 0,121705 | 2,734925 | 0,008961 | 0,037135 | -3,32936 |
| Notch3 | 0,166996 | 0,090122 | 2,769574 | 0,008189 | 0,035399 | -3,24731 |
| PDH | 0,165939 | 0,028879 | 2,483139 | 0,016909 | 0,056936 | -3,90313 |
| Cyclin-D1 | 0,163783 | 0,037598 | 2,776165 | 0,00805 | 0,035399 | -3,23162 |
| MIF | 0,163741 | -0,0104 | 2,668715 | 0,010624 | 0,041448 | -3,48412 |
| LAD1 | 0,162616 | -0,0278 | 2,252256 | 0,029349 | 0,082676 | -4,39233 |
| MelanA | 0,162572 | -0,03267 | 2,502521 | 0,016122 | 0,055943 | -3,86041 |
| MEK2 | 0,161968 | -0,0442 | 2,43359 | 0,019082 | 0,062306 | -4,01119 |
| N-Cadherir | 0,160043 | 0,03129 | 2,381597 | 0,021631 | 0,065204 | -4,1228 |
| 14-3-3-eps | 0,1599 | 0,027058 | 2,724949 | 0,009195 | 0,037742 | -3,35285 |
| Bid | 0,158638 | 0,076547 | 2,533567 | 0,014931 | 0,054076 | -3,79147 |
| PD-L1 | 0,158394 | 0,052266 | 2,829978 | 0,00699 | 0,033108 | -3,10254 |
| MIG6 | 0,157951 | 0,101079 | 2,742023 | 0,008797 | 0,036812 | -3,31261 |
| SFRP1 | 0,157398 | -0,00456 | 2,399884 | 0,020702 | 0,064501 | -4,08376 |
| Bak | 0,154867 | -0,01284 | 2,479593 | 0,017057 | 0,056989 | -3,91092 |
| PDHK1 | 0,15269 | 0,068548 | 2,646183 | 0,011251 | 0,041448 | -3,53617 |
| PRAS40 | 0,147202 | 0,046683 | 2,524564 | 0,015268 | 0,054836 | -3,81153 |
| Creb | 0,144895 | 0,013834 | 2,408974 | 0,020253 | 0,064501 | -4,06426 |
| E2F1 | 0,14419 | 0,059001 | 2,391116 | 0,021143 | 0,065089 | -4,10251 |
| Bad_pS112 | 0,134345 | 0,068923 | 2,098507 | 0,041638 | 0,106821 | -4,69712 |
| SOX17 | 0,133305 | 0,018711 | 2,026371 | 0,048818 | 0,117546 | -4,83407 |
| BAP1 | 0,129545 | 0,048736 | 2,397876 | 0,020802 | 0,064501 | -4,08806 |
| Ets-1 | 0,123696 | -0,02815 | 2,277757 | 0,027657 | 0,078941 | -4,34012 |
| Hexokinase | 0,122342 | -0,02538 | 2,381739 | 0,021624 | 0,065204 | -4,1225 |
| DAPK1_pS | 0,104625 | 0,050536 | 2,042284 | 0,047148 | 0,116786 | -4,80419 |

cut-off : P.Value<0.05

**Table S5E. A11+Adav_vs_No.**

| ID | logFC | AveExpr | t | P.Value | adj.P.Val | B |
|---|---|---|---|---|---|---|
| Histone-H3 | 1,660532 | 0,365476 | 4,845047 | 1,61E-05 | 0,00099 | 2,782526 |
| PAR | 1,510213 | 0,33572 | 2,973889 | 0,004759 | 0,070328 | -2,58996 |
| Caspase-3- | 1,396942 | 0,182884 | 4,993032 | 9,88E-06 | 0,00071 | 3,250265 |
| Pyk2_pY40 | -0,96877 | -0,01398 | -3,97781 | 0,000256 | 0,008491 | 0,140618 |
| H2AX_pS13 | 0,893769 | 0,008345 | 4,270372 | 0,000103 | 0,00402 | 1,009495 |
| cdc2_pY15 | -0,87577 | 0,010496 | -9,91986 | 8,61E-13 | 3,71E-10 | 18,94938 |
| Hif-1-alpha | -0,85022 | -0,06664 | -5,18395 | 5,25E-06 | 0,000452 | 3,858932 |
| CDK1_pT14 | 0,844253 | -0,00726 | 2,951311 | 0,005058 | 0,070328 | -2,64583 |
| p38-MAPK- | 0,823227 | -0,03056 | 3,026283 | 0,004128 | 0,068422 | -2,45923 |
| H2AX_pS14 | 0,790435 | 0,173534 | 4,534002 | 4,42E-05 | 0,002117 | 1,813286 |
| ER-a_pS11 | 0,790237 | -0,08919 | 2,959113 | 0,004953 | 0,070328 | -2,62656 |
| DM-Histon | 0,75352 | 0,167256 | 5,258556 | 4,09E-06 | 0,000441 | 4,098187 |
| Rad51 | -0,75204 | 0,041184 | -3,2868 | 0,001996 | 0,039113 | -1,7879 |
| DUSP6 | -0,717 | 0,009088 | -2,52538 | 0,015237 | 0,164176 | -3,64407 |
| Chk1 | -0,70422 | 0,162593 | -3,67296 | 0,000647 | 0,014672 | -0,73396 |
| Connexin-4 | -0,69722 | -0,0405 | -2,254 | 0,02923 | 0,237703 | -4,21971 |
| PHLPP | -0,64405 | 0,013041 | -3,00953 | 0,00432 | 0,068966 | -2,50118 |
| Caspase-7- | 0,627079 | 0,19787 | 4,550612 | 4,19E-05 | 0,002117 | 1,86451 |
| Snail | -0,61763 | 0,250236 | -2,37936 | 0,021747 | 0,203763 | -3,96003 |
| MMP14 | -0,5732 | 0,114206 | -2,84508 | 0,006717 | 0,087728 | -2,90488 |
| RBM15 | 0,549739 | 0,077333 | 3,778706 | 0,000471 | 0,012677 | -0,43449 |
| MLH1 | 0,469404 | 0,049452 | 4,490071 | 5,09E-05 | 0,002194 | 1,678118 |
| Akt1 | -0,46457 | -0,05668 | -3,89089 | 0,000335 | 0,010298 | -0,11219 |
| KAP1 | 0,457076 | 0,102915 | 2,60094 | 0,012616 | 0,146961 | -3,47509 |
| SOD1 | 0,449312 | 0,07373 | 5,615004 | 1,24E-06 | 0,000178 | 5,25006 |
| Akt2 | -0,44902 | -0,17563 | -3,73922 | 0,00053 | 0,013442 | -0,54683 |
| Bcl2 | 0,424827 | 0,084614 | 4,148096 | 0,000151 | 0,005416 | 0,643143 |
| Stathmin-1 | 0,403128 | -0,0511 | 2,188235 | 0,034009 | 0,262066 | -4,3515 |
| EMA | -0,39344 | 0,022802 | -3,70931 | 0,00058 | 0,013889 | -0,63151 |
| Annexin-I | 0,357552 | 0,056642 | 3,133002 | 0,003076 | 0,055247 | -2,18841 |
| RPA32 | -0,34467 | -0,0536 | -2,71053 | 0,009543 | 0,117518 | -3,22361 |
| IGF1R_pY1 | -0,33658 | 0,123006 | -2,28208 | 0,027379 | 0,231375 | -4,16249 |
| Mcl-1 | -0,33517 | -0,13578 | -2,58973 | 0,012977 | 0,147185 | -3,50038 |
| Rictor_pT1 | -0,32518 | -0,05303 | -2,13624 | 0,038264 | 0,280358 | -4,4535 |
| PARP | -0,31278 | -0,03808 | -2,87949 | 0,006131 | 0,082578 | -2,82167 |
| SHP2 | -0,29082 | -0,06484 | -5,80895 | 6,45E-07 | 0,000139 | 5,881525 |
| Erk5 | -0,28423 | -0,12646 | -3,21449 | 0,00245 | 0,045903 | -1,97766 |
| S6 | 0,283645 | 0,054635 | 2,309028 | 0,025701 | 0,226063 | -4,10707 |
| 4E-BP1_pT | -0,27786 | -0,04807 | -2,03849 | 0,047541 | 0,315236 | -4,63982 |
| Caspase-8 | 0,269812 | 0,089702 | 3,412426 | 0,001392 | 0,029351 | -1,45229 |
| HER3_pY12 | -0,26268 | 0,046123 | -3,85302 | 0,000376 | 0,01079 | -0,22148 |
| DDB-1 | -0,25911 | 0,062349 | -2,29022 | 0,026862 | 0,231375 | -4,14581 |
| DRP1 | -0,24934 | -0,07825 | -2,41783 | 0,019825 | 0,189875 | -3,87817 |
| ERRalpha | -0,24313 | -0,0124 | -2,06867 | 0,044487 | 0,299593 | -4,58305 |
| PKCa | -0,23768 | -0,16464 | -2,61958 | 0,012037 | 0,144105 | -3,43285 |
| B-Raf_pS44 | -0,2342 | -0,04863 | -2,35566 | 0,023014 | 0,206645 | -4,00996 |
| Raptor | -0,23191 | -0,08896 | -3,40317 | 0,00143 | 0,029351 | -1,47726 |
| Lasu1 | -0,22083 | -0,10668 | -2,23452 | 0,03058 | 0,244071 | -4,25908 |
| XPA | 0,212115 | -0,00526 | 2,120066 | 0,039679 | 0,280358 | -4,48481 |

| | | | | | | |
|---|---|---|---|---|---|---|
| ATP5A | 0,20923 | 0,072204 | 2,995605 | 0,004487 | 0,06907 | -2,53596 |
| Annexin-VI | 0,207886 | 0,064325 | 3,048624 | 0,003883 | 0,066941 | -2,40303 |
| GAPDH | -0,19961 | -0,02123 | -2,17838 | 0,034782 | 0,262999 | -4,37098 |
| Atg4B | -0,18483 | -0,10914 | -2,49903 | 0,016261 | 0,170027 | -3,70213 |
| IRF-1 | 0,179682 | 0,061071 | 2,360705 | 0,022739 | 0,206645 | -3,99936 |
| Bad_pS112 | 0,177552 | 0,068923 | 2,773425 | 0,008108 | 0,102776 | -3,07596 |
| SF2 | 0,17565 | 0,101404 | 2,491419 | 0,016569 | 0,170027 | -3,71882 |
| C-Raf | -0,16826 | 0,036654 | -2,43318 | 0,019101 | 0,187105 | -3,84523 |
| CSK | -0,15892 | -0,05276 | -2,07448 | 0,04392 | 0,299593 | -4,57205 |
| SOD2 | 0,14946 | 0,022023 | 2,089779 | 0,042454 | 0,295126 | -4,54294 |
| TUFM | -0,14601 | -0,08513 | -2,26948 | 0,028196 | 0,233703 | -4,18824 |
| Syk | 0,13419 | 0,060814 | 2,534873 | 0,014882 | 0,164176 | -3,62304 |
| CD49b | 0,129286 | 0,083165 | 2,122811 | 0,039436 | 0,280358 | -4,47951 |
| Collagen-V | 0,12828 | 0,014401 | 2,187705 | 0,03405 | 0,262066 | -4,35255 |
| DAPK1_pS3 | 0,12577 | 0,050536 | 2,455041 | 0,018112 | 0,181543 | -3,79804 |
| Rheb | 0,109917 | 0,01574 | 2,128986 | 0,038893 | 0,280358 | -4,46756 |
| BAP1 | 0,109289 | 0,048736 | 2,022942 | 0,049185 | 0,321193 | -4,6688 |

cut-off : P.Value<0.05

**Table S6:** Antibodies used for RPPA analysis of *MMTV-R26^Met^* treated cells

| # | Official Ab Name | Ab Name Reported on Dataset | Gene Name | Company | Catalog # | Species | RPPA Dilution |
|---|---|---|---|---|---|---|---|
| **1** | 14-3-3 beta | 14-3-3-beta | YWHAB | Santa Cruz | sc-628 | Rabbit | 1:75 |
| **2** | 14-3-3 epsilon | 14-3-3-epsilon | YWHAE | Santa Cruz | SC-23957 | Mouse | 1:50 |
| **3** | 14-3-3 zeta | 14-3-3-zeta | YWHAZ | Santa Cruz | sc-1019 | Rabbit | 1:5000 |
| **4** | 4E-BP1 | 4E-BP1 | EIF4EBP1 | CST | 9452 | Rabbit | 1:100 |
| **5** | 4E-BP1 (phospho S65) | 4E-BP1_pS65 | EIF4EBP1 | CST | 9456 | Rabbit | 1:250 |
| **6** | 4E-BP1 (phospho T37/46) | 4E-BP1-pT37-T46 | EIF4EBP1 | CST | 9459 | Rabbit | 1:2000 |
| **7** | 53BP1 | 53BP1 | TP53BP1 | CST | 4937 | Rabbit | 1:300 |
| **8** | A1Up | UBQLN4 | UBQLN4 | Santa Cruz | sc-136145 | Mouse | 1:125 |
| **9** | Acetyl-CoA-Carboxylase | ACC1 | ACACA, B | Epitomics/ Abcam | 1768-1/ ab45174 | Rabbit | 1:1500 |
| **10** | Acetyl-CoA-Carboxylase (phospho S79) | ACC_pS79 | ACACA, B | CST | 3661 | Rabbit | 1:500 |
| **11** | ACSL1 (D2H5) | ACSL1 | ACSL1 | CST | 9189 | Rabbit | 1:500 |
| **12** | ACVRL1 | ACVRL1 | ACVRL1 | Epitomics/ Abcam | 2940-1/ ab108207 | Rabbit | 1:30 |
| **13** | ADAR1 | ADAR1 | ADAR | Abcam | ab88574 | Mouse | 1:100 |
| **14** | Akt | Akt | AKT1, 2, 3 | CST | 4691 | Rabbit | 1:7500 |
| **15** | Akt (phospho S473) | Akt_pS473 | AKT1, 2, 3 | CST | 9271 | Rabbit | 1:150 |
| **16** | Akt (phospho T308) | Akt_pT308 | AKT1, 2, 3 | CST | 2965 | Rabbit | 1:250 |
| **17** | Akt1 | Akt1 | AKT1 | CST | 2938 | Rabbit | 1:1000 |
| **18** | Akt1 (phospho S473) | Akt1_pS473 | AKT1 | CST | 9018 | Rabbit | 1:1000 |
| **19** | Akt2 | Akt2 | AKT2 | CST | 3063 | Rabbit | 1:3000 |
| **20** | Akt2 (phospho S474) | Akt2_pS474 | AKT2 | CST | 8599 | Rabbit | 1:1000 |
| **21** | Ambra1 (phospho S52) | Ambra1_pS52 | AMBRA1 | Millipore | ABC80 | Rabbit | 1:250 |
| **22** | AMPK alpha 2 (phospho S345) | AMPK-a2_pS345 | PRKAA1, 2 | Abcam | ab129081 | Rabbit | 1:200 |
| **23** | AMPKa | AMPKa | PRKAA1, 2 | CST | 2532 | Rabbit | 1:75 |

| | | | | | | | |
|---|---|---|---|---|---|---|---|
| **24** | AMPKa (phospho T172) | AMPKa_pT172 | PRKAA1, 2 | CST | 2535 | Rabbit | 1:100 |
| **25** | Androgen Receptor (D6F11) | AR | AR | CST | 5153 | Rabbit | 1:250 |
| **26** | Annexin I | Annexin-I | ANXA1 | BD Biosciences | 610066 | Mouse | 1:5000 |
| **27** | Annexin VII | Annexin-VII | ANXA7 | BD Biosciences | 610668 | Mouse | 1:20 |
| **28** | A-Raf | A-Raf | ARAF | CST | 4432 | Rabbit | 1:200 |
| **29** | A-Raf (phospho S299) | A-Raf_pS299 | ARAF | CST | 4431 | Rabbit | 1:25 |
| **30** | ARID1A | ARID1A | ARID1A | Sigma-Aldrich | HPA005456 | Rabbit | 1:1000 |
| **31** | ASNS | ASNS | ASNS | Sigma-Aldrich | HPA029318 | Rabbit | 1:500 |
| **32** | Atg3 | Atg3 | ATG3 | CST | 3415 | Rabbit | 1:72 |
| **33** | Atg4B | Atg4B | ATG4B | CST | 13507 | Rabbit | 1:200 |
| **34** | Atg5 | Atg5 | ATG5 | CST | 12994 | Rabbit | 1:1000 |
| **35** | Atg7 | Atg7 | ATG7 | CST | 8558 | Rabbit | 1:1000 |
| **36** | ATM | ATM | ATM | CST | 2873 | Rabbit | 1:250 |
| **37** | ATM (phospho S1981) | ATM_pS1981 | ATM | CST | 5883 | Rabbit | 1:20 |
| **38** | ATP5A | ATP5A | ATP5A | Abcam | ab14748 | Mouse | 1:500 |
| **39** | ATP5H | ATP5H | ATP5H | Abcam | ab110275 | Mouse | 1:30 |
| **40** | ATR | ATR | ATR | CST | 2790 | Rabbit | 1:30 |
| **41** | ATR (phospho S428) | ATR_pS428 | ATR | Abcam | ab178407 | Rabbit | 1:1000 |
| **42** | ATRX | ATRX | ATRX | Abcam | ab97508 | Rabbit | 1:300 |
| **43** | Aurora B/AIM1 | Aurora-B | AURKB | CST | 3094 | Rabbit | 1:38 |
| **44** | Axl | Axl | AXL | CST | 8661 | Rabbit | 1:500 |
| **45** | B7-H3 | B7-H3 | CD276 | CST | 14058 | Rabbit | 1:200 |
| **46** | B7-H4 | B7-H4 | VTCN1 | CST | 14572 | Rabbit | 1:50 |
| **47** | Bad (phospho S112) | Bad_pS112 | BAD | CST | 9291 | Rabbit | 1:50 |
| **48** | Bak | Bak | BAK1 | Epitomics/ Abcam | 1542-1/ ab32371 | Rabbit | 1:400 |
| **49** | BAP1 | BAP1 | BAP1 | Santa Cruz | sc-28383 | Mouse | 1:200 |
| **50** | Bax | Bax | BAX | CST | 2772 | Rabbit | 1:100 |
| **51** | b-Catenin | b-Catenin | CTNNB1 | CST | 9562 | Rabbit | 1:1500 |
| **52** | Bcl2 | Bcl2 | BCL2 | Dako | M0887 | Mouse | 1:50 |
| **53** | Bcl2A1 | Bcl2A1 | BCL2A1 | Abnova | PAB8528 | Rabbit | 1:250 |
| **54** | Bcl-xL | Bcl-xL | BCL2L1 | CST | 2762 | Rabbit | 1:100 |

| | | | | | | | |
|---|---|---|---|---|---|---|---|
| **55** | Beclin 1 | Beclin | BECN1 | ThermoFisher | PA1-16857 | Rabbit | 1:500 |
| **56** | beta Actin | b-Actin | ACTB | CST | 4970 | Rabbit | 1:50 |
| **57** | beta Catenin (phospho T41/S45) | b-Catenin_pT41_S45 | CTNNB1 | CST | 9565 | Rabbit | 1:30 |
| **58** | Bid | Bid | BID | CST | 2002 | Rabbit | 1:500 |
| **59** | Bim (C34C5) | Bim | BCL2L11 | Epitomics/ Abcam | 1036-1/ ab32158 | Rabbit | 1:400 |
| **60** | BiP/GRP78 | BiP-GRP78 | HSPA5 | BD Biosciences | 610978 | Mouse | 1:150 |
| **61** | BMK1/Erk5 (phospho T218/Y220) | BMK1-Erk5_pT218_Y220 | MAPK7 | Millipore | 07-507 | Rabbit | 1:500 |
| **62** | B-Raf | B-Raf | BRAF | CST | 14814 | Rabbit | 1:500 |
| **63** | B-Raf (phospho S445) | B-Raf_pS445 | BRAF | CST | 2696 | Rabbit | 1:75 |
| **64** | BRD4 | BRD4 | BRD4 | CST | 13440 | Rabbit | 1:1000 |
| **65** | CA9 (CAIX) | CA9 | CA9 | CST | 5649 | Rabbit | 1:200 |
| **66** | c-Abl | c-Abl | ABL1 | CST | 2862 | Rabbit | 1:100 |
| **67** | c-Abl (phospho Y412) | Abl_pY412 | ABL1 | CST | 2865 | Rabbit | 1:200 |
| **68** | Caspase 3 (cleaved asp175) | Caspase-3-cleaved | CASP3 | CST | 9661 | Rabbit | 1:500 |
| **69** | Caspase 7 (cleaved) | Caspase-7-cleaved | CASP7 | CST | 9491 | Rabbit | 1:60 |
| **70** | Caspase 8 | Caspase-8 | CASP8 | CST | 9746 | Mouse | 1:150 |
| **71** | Caspase 8 (cleaved asp391) | Caspase-8-cleaved | CASP8 | CST | 9496 | Rabbit | 1:500 |
| **72** | Caspase-3 | Caspase-3 | CASP3 | Epitomics/ Abcam | 1476-1/ ab32042 | Rabbit | 1:250 |
| **73** | Caveolin 1 | Caveolin-1 | CAV1 | CST | 3238 | Rabbit | 1:3000 |
| **74** | CD134/OX40 | CD134 | TNFRSF | Abcam | ab76000 | Rabbit | 1:100 |
| **75** | CD171 (L1) | CD171 | L1CAM | Biolegend | 826701 | Mouse | 1:1000 |
| **76** | CD20 | CD20 | MS4A1 | Epitomics/ Abcam | 1632-1/ ab78237 | Rabbit | 1:75 |
| **77** | CD26 | CD26 | DPP4 | Abcam | ab28340 | Rabbit | 1:1000 |
| **78** | CD29 | CD29 | ITGB1 | BD Biosciences | 610467 | Mouse | 1:30 |
| **79** | CD31 | CD31 | PECAM1 | Dako/ Fisher | M0823/ MS353S | Mouse | 1:25 |
| **80** | CD38 | CD38 | CD38 | Abcam | ab108403 | Rabbit | 1:250 |
| **81** | CD4 | CD4 | CD4 | Abcam | ab133616 | Rabbit | 1:500 |

| | | | | | | | |
|---|---|---|---|---|---|---|---|
| **82** | CD44 | CD44 | CD44 | CST | 3570 | Mouse | 1:20 |
| **83** | CD45 | CD45 | CD45 | DAKO/ ThermoFisher | M070129-2/ MS355P | Mouse | 1:1000 |
| **84** | CD49b | CD49b | ITGA2 | BD Biosciences | 611016 | Mouse | 1:50 |
| **85** | CD86 | CD86 | CD86 | Abcam | ab53004 | Rabbit | |
| **86** | Cdc2 (phospho Y15) | cdc2_pY15 | CDK | CST | 4539 | Rabbit | 1:38 |
| **87** | cdc25C | cdc25C | CDC25C | CST | 4688 | Rabbit | 1:250 |
| **88** | CDK1/2/3 (phospho T14) | CDK1_pT14 | CDK1, 2, 3 | Abcam | ab32384 | Rabbit | 1:1000 |
| **89** | CDKN2A/p16INK4 a | p16INK4a | CDKN2A | Abcam | ab81278 | Rabbit | 1:500 |
| **90** | Chk1 | Chk1 | CHEK | CST | 2360 | Mouse | 1:100 |
| **91** | Chk1 (phospho S296) | Chk1_pS296 | CHEK1 | Abcam | ab79758 | Rabbit | 1:125 |
| **92** | Chk1 (phospho S345) | Chk1_pS345 | CHEK1 | CST | 2348 | Rabbit | 1:30 |
| **93** | Chk2 | Chk2 | CHEK2 | CST | 3440 | Mouse | 1:50 |
| **94** | Chk2 (phospho T68) | Chk2_pT68 | CHEK2 | CST | 2197 | Rabbit | 1:250 |
| **95** | c-IAP2 | c-IAP2 | BIRC3 | CST | 3130 | Rabbit | 1:50 |
| **96** | CIITA | CIITA | CIITA | CST | 3793 | Rabbit | 1:250 |
| **97** | c-Jun (phospho S73) | c-Jun_pS73 | JUN | CST | 9164 | Rabbit | 1:30 |
| **98** | c-Kit | c-Kit | KIT | Epitomics/ Abcam | 1522-1/ ab32363 | Rabbit | 1:250 |
| **99** | Claudin 7 | Claudin-7 | CLDN7 | Abcam | ab79481 | Rabbit | 1:250 |
| **100** | c-Myc | c-Myc | MYC | Santa Cruz | sc-764 | Rabbit | 1:250 |
| **101** | COG3 | COG3 | COG3 | ProteinTech | 11130-1-AP | Rabbit | 1:750 |
| **102** | Collagen-VI/COL6A1 | Collagen-VI | COL6A1 | Santa Cruz | sc-20649 | Rabbit | 1:6000 |
| **103** | Complex II Subunit | Complex-II-Subunit | SDHB | Life Technologies | 459230 | Mouse | 1:200 |
| **104** | Connexin 43 | Connexin-43 | GJA1 | CST | 3512 | Rabbit | 1:150 |
| **105** | Coup-TFII | Coup-TFII | NR2F2 | CST | 6434 | Rabbit | 1:50 |
| **106** | Cox2 | Cox2 | PTGS2 | CST | 4842 | Rabbit | 1:75 |
| **107** | Cox-IV | Cox-IV | COX4I1 | CST | 4850 | Rabbit | 1:5000 |
| **108** | C-Raf | C-Raf | RAF1 | Millipore | 04-739 | Rabbit | 1:100 |
| **109** | C-Raf (phospho S338) | C-Raf_pS338 | RAF1 | CST | 9427 | Rabbit | 1:200 |

| | | | | | | | |
|---|---|---|---|---|---|---|---|
| **110** | Creb | Creb | CREB1 | CST | 9197 | Rabbit | 1:75 |
| **111** | CSK | CSK | CSK | CST | 4980 | Rabbit | 1:300 |
| **112** | CtIP | CtIP | RBBP8 | CST | 9201 | Rabbit | 1:500 |
| **113** | Cyclin B1 | Cyclin B1 | CCNB1 | Epitomics/ Abcam | 1495-1/ ab32053 | Rabbit | 1:1500 |
| **114** | Cyclin D1 | Cyclin-D1 | CCND1 | Millipore Sigma | SAB4502603 | Rabbit | 1:200 |
| **115** | Cyclin D3 | Cyclin D3 | CCND3 | CST | 2936 | Mouse | 1:1000 |
| **116** | Cyclin E1 | Cyclin E1 | CCNE1 | Santa Cruz | sc-247 | Mouse | 1:25 |
| **117** | Cyclophilin-F | Cyclophilin-F | PPIF | Abcam | MSA04/ ab110324 | Mouse | 1:50000 |
| **118** | Cytokeratin 19 | Cytokeratin-19 | KRT19 | Dako | M0888 | Mouse | 1:50 |
| **119** | DAP Kinase 1 (phospho S308) | DAPK1_pS308 | DAPK1 | GeneTex | GTX10524 | Mouse | 1:200 |
| **120** | DAP Kinase 2 | DAPK2 | DAPK2 | Abcam | ab51601 | Rabbit | 1:250 |
| **121** | DDB-1 | DDB-1 | DDB1 | CST | 6998 | Rabbit | 1:5000 |
| **122** | Detyrosinated alpha-Tubulin | D-a-Tubulin | TUBA4A, TUBA3C | Abcam | ab48389 | Rabbit | 1:1500 |
| **123** | Di-Methyl-Histone H3 (Lys4/C64G9) | DM-Histone-H3 | HIST1H3A | CST | 9725 | Rabbit | 1:100 |
| **124** | Dimethyl-K9 Histone H3 | DM-K9-Histone-H3 | HIST3H3 | Abcam | ab1220 | Mouse | 1:250 |
| **125** | DNA Ligase IV | DNA-Ligase-IV | LIG4 | CST | 14649 | Rabbit | 1:1000 |
| **126** | DNA Polymerase gamma (D1Y6R) | POLG | POLG | CST | 13609 | Rabbit | 1:500 |
| **127** | DNMT1 (D63A6) | DNMT1 | DNMT1 | CST | 5032 | Rabbit | 1:500 |
| **128** | DRP1 (D8H5) | DRP1 | DNM1L | CST | 5391 | Rabbit | 1:1000 |
| **129** | DUSP4/MKP2 | DUSP4 | DUSP4 | CST | 5149 | Rabbit | 1:150 |
| **130** | DUSP6 | DUSP6 | DUSP6 | Abcam | ab76310 | Rabbit | 1:750 |
| **131** | Dvl3 | Dvl3 | DVL3 | CST | 3218 | Rabbit | 1:30 |
| **132** | E2F1 | E2F1 | E2F1 | Santa Cruz | sc-251 | Mouse | 1:20 |
| **133** | E-Cadherin | E-Cadherin | CDH1 | CST | 3195 | Rabbit | 1:150 |
| **134** | eEF2 | eEF2 | EEF2 | CST | 2332 | Rabbit | 1:50 |
| **135** | eEF2K | eEF2K | EEF2K | CST | 3692 | Rabbit | 1:50 |
| **136** | EGFR | EGFR | EGFR | CST | 2232 | Rabbit | 1:75 |
| **137** | EGFR (phospho Y1173) | EGFR_pY1173 | EGFR | Epitomics/ Abcam | 1124-1/ ab32578 | Rabbit | 1:300 |
| **138** | eIF4E | eIF4E | EIF4E | CST | 9742 | Rabbit | 1:75 |
| **139** | eIF4E (phospho S209) | eIF4E_pS209 | EIF4E | Abcam | ab76256 | Rabbit | 1:250 |

| | | | | | | | |
|---|---|---|---|---|---|---|---|
| **140** | eIF4G | eIF4G | EIF4G1 | CST | 2498 | Rabbit | 1:1000 |
| **141** | Elk1 (phospho S383) | Elk1_pS383 | ELK1 | CST | 9181 | Rabbit | 1:50 |
| **142** | Enolase-2 (D20H2) | Enolase-2 | ENO2 | CST | 8171 | Rabbit | 1:250 |
| **143** | ENY2 | ENY2 | ENY2 | GeneTex | GTX629542 | Mouse | 1:500 |
| **144** | Eph Receptor A2 | EPHA2 | EPHA2 | Abcam | ab133501 | Rabbit | 1:1000 |
| **145** | Epithelial Membrane Antigen | EMA | MUC1 | DAKO | M061329-2 | Mouse | 1:750 |
| **146** | ErbB3/HER3 | HER3 | ERBB3 | Santa Cruz | sc-285 | Rabbit | 1:300 |
| **147** | ErbB3/HER3 (phospho Y1289) | HER3_pY1289 | ERBB3 | CST | 4791 | Rabbit | 1:50 |
| **148** | ERCC1 | ERCC1 | ERCC1 | Santa Cruz | sc-17809 | Mouse | 1:38 |
| **149** | Erk5 | Erk5 | MAPK7 | CST | 3552 | Rabbit | 1:500 |
| **150** | ERRalpha (E1G1J) | ERRalpha | ESRRA | CST | 13826 | Rabbit | 1:500 |
| **151** | ERRFI1/MIG6 | MIG6 | ERRFI1 | Sigma-Aldrich | WH0054206 M1 | Mouse | 1:50 |
| **152** | Estrogen Receptor | ER | ESR1 | Lab Vision | RM-9101 | Rabbit | 1:40 |
| **153** | Estrogen Receptor alpha | ER-a | ERSA | CST | 13258 | Rabbit | 1:500 |
| **154** | Estrogen Receptor alpha (phospho S118) | ER-a_pS118 | ESR1 | Epitomics/ Abcam | 1091-1/ ab32396 | Rabbit | 1:500 |
| **155** | Ets-1 | Ets-1 | ETS1 | Bethyl | A303-501A | Rabbit | 1:100 |
| **156** | FAK | FAK | PTK2 | Epitomics/ Abcam | 1700-1/ ab40794 | Rabbit | 1:1000 |
| **157** | FAK (phospho Y397) | FAK_pY397 | PTK2 | CST | 3283 | Rabbit | 1:25 |
| **158** | Fatty Acid Synthase | FASN | FASN | CST | 3180 | Rabbit | 1:1000 |
| **159** | FGF-basic | FGF-basic | FGF2 | VWR | 10775-082 (500-P18) | Rabbit | 1:1000 |
| **160** | Fibronectin | Fibronectin | FN1 | Epitomics | 1574-1 | Rabbit | 1:10000 |
| **161** | FoxM1 | FOXM1 | FOXM1 | CST | 5436 | Rabbit | 1:30 |
| **162** | FoxO3a | FoxO3a | FOXO3 | CST | 2497 | Rabbit | 1:20 |
| **163** | FoxO3a (phospho S318/S321) | FoxO3a_pS318_S3 21 | FOXO3 | CST | 9465 | Rabbit | 1:30 |
| **164** | FRS2-a (phospho Y196) | FRS2-a_pY196 | FRS2 | CST | 3864 | Rabbit | 1:100 |
| **165** | G6PD | G6PD | G6PD | Santa Cruz | sc-373887 | Mouse | 1:75 |

| | | | | | | | |
|---|---|---|---|---|---|---|---|
| **166** | Gab2 | Gab2 | GAB2 | CST | 3239 | Rabbit | 1:300 |
| **167** | GAPDH | GAPDH | GAPDH | Ambion/ Invitrogen | AM4300 | Mouse | 1:75000 |
| **168** | GATA3 | GATA3 | GATA3 | BD Biosciences | 558686 | Mouse | 1:150 |
| **169** | GATA6 | GATA6 | GATA6 | CST | 5851 | Rabbit | 1:200 |
| **170** | GCLC | GCLC | GCLC | Proteintech Group | 12601-1-AP | Rabbit | 1:500 |
| **171** | GCLM | GCLM | GCLM | Abcam | ab124827 | Rabbit | 1:500 |
| **172** | GCN5L2 | GCN5L2 | KAT2A | CST | 3305 | Rabbit | 1:30 |
| **173** | Gli1 | Gli1 | GLI1 | CST | 3538 | Rabbit | 1:3000 |
| **174** | Gli3 | Gli3 | GLI3 | Abcam | ab69838 | Rabbit | 1:1000 |
| **175** | Glucose-6 Phosphate Dehydrogenase | G6PD | G6PD | CST | 8866 | Rabbit | 1:30 |
| **176** | Glutamate Dehydrogenase1/ 2 | Glutamate-D1-2 | GLUD1 | Novus | NBP2-16679 | Rabbit | 1:500 |
| **177** | Glutaminase | Glutaminase | GLS | Abcam | ab156876 | Rabbit | 1:150 |
| **178** | Glycogen Synthase | Gys | GYS1 | CST | 3886 | Rabbit | 1:2000 |
| **179** | Glycogen Synthase (phospho S641) | Gys_pS641 | GYS1 | CST | 3891 | Rabbit | 1:300 |
| **180** | GPBB | GPBB | PYGM | Novus | NBP1-32799 | Rabbit | 1:200 |
| **181** | Granzyme B | Granzyme-B | GZMB | CST | 4275 | Rabbit | 1:500 |
| **182** | GRB7 | GRB7 | GRB7 | Abcam | ab183737 | Rabbit | 1:500 |
| **183** | Grp75 (D13H4) | Grp75 | HSPA9 | CST | 3593 | Rabbit | 1:250 |
| **184** | GSK-3alpha/beta | GSK-3a-b | GSK3A, B | Santa Cruz | sc-7291 | Mouse | 1:750 |
| **185** | GSK-3alpha/beta (phospho S21/S9) | GSK-3a-b_pS21_S9 | GSK3A, B | CST | 9331 | Rabbit | 1:200 |
| **186** | GSK-3B | GSK-3B | GSK3B | CST | 9315 | Rabbit | 1:750 |
| **187** | GSK-3beta (phospho S9) | GSK-3b_pS9 | GSK3B | CST | 5558 | Rabbit | 1:250 |
| **188** | H2AX (phospho S140) | H2AX_pS140 | H2AFX | Pierce Biotechnolog y | MA12022 | Mouse | 1:100 |
| **189** | Hamartin/TSC1 | TSC1 | TSC1 | CST | 4906 | Rabbit | 1:200 |
| **190** | HER2 | HER2 | ERBB2 | Lab Vision | MS-325-P1 | Mouse | 1:300 |
| **191** | HER2 (phospho Y1248) | HER2_pY1248 | ERBB2 | R&D systems | AF1768 | Rabbit | 1:1500 |
| **192** | Heregulin | Heregulin | NRG1 | CST | 2573 | Rabbit | 01:30 |

| | | | | | | | |
|---|---|---|---|---|---|---|---|
| **193** | HES1 | HES1 | HES1 | CST | 11988 | Rabbit | 1:500 |
| **194** | Hexokinase II | Hexokinase II | HK2 | CST | 2106 | Rabbit | 1:100 |
| **195** | Hif-1-alpha | Hif-1-alpha | HIF1A | BD Biosciences | 610958 | Mouse | 1:20 |
| **196** | Histone H3 | Histone H3 | HIST3H3 | Abcam | ab1791 | Rabbit | 1:5000 |
| **197** | HLA-DQA1 | HLA-DQA1 | HLA-DQA1 | Abcam | ab128959 | Rabbit | 1:3000 |
| **198** | HLA-DR/DP/DQ/DX | HLA-DR-DP-DQ-DX | HLA-DRA | Santa Cruz | sc-53302 | Mouse | 1:250 |
| **199** | HMHA1 | HMHA1 | HMHA1 | ProteinTech | 14832-1-AP | Rabbit | 1:3000 |
| **200** | HSP27 | HSP27 | HSBP1 | CST | 2402 | Mouse | 1:75 |
| **201** | HSP27 (phospho S82) | HSP27_pS82 | HSBP1 | CST | 2401 | Rabbit | 1:75 |
| **202** | HSP60 | HSP60 | HSP60 | CST | 12165 | Rabbit | 1:1000 |
| **203** | HSP70 | HSP70 | HSPA1A | CST | 4872 | Rabbit | 1:50 |
| **204** | Hsp75/TRAP1 | TRAP1 | TRAP1 | BD Biosciences | 612344 | Mouse | 1:750 |
| **205** | IDO | IDO | IDO1 | CST | 86630 | Rabbit | 1:200 |
| **206** | IGF1R (phospho Y1135/Y1136 | IGF1R_pY1135_Y1136 | IGF1R, INSR | CST | 3024 | Rabbit | 1:30 |
| **207** | IGF-1Receptor beta | IGF1R-b | IGF1R | CST | 3018 | Rabbit | 1:50 |
| **208** | IGFBP2 | IGFBP2 | IGFBP2 | CST | 3922 | Rabbit | 1:50 |
| **209** | IGFBP3 | IGFBP3 | IGFBP3 | BD Biosciences | 611504 | Mouse | 1:1000 |
| **210** | IGFRb | IGFRb | IGF1R | CST | 3027 | Rabbit | 1:250 |
| **211** | IL-6 | IL-6 | IL6 | CST | 12153 | Rabbit | 1:250 |
| **212** | INPP4b | INPP4b | INPP4B | CST | 4039 | Rabbit | 01:30 |
| **213** | Insulin Receptor beta | IR-b | INSR | CST | 3025 | Rabbit | 1:100 |
| **214** | IRF-1 | IRF-1 | IRF1 | CST | 8478 | Rabbit | 1:250 |
| **215** | IRS1 | IRS1 | IRS1 | Millipore | 06-248 | Rabbit | 1:250 |
| **216** | IRS2 | IRS2 | IRS2 | CST | 4502 | Rabbit | 1:100 |
| **217** | JAB1 | JAB1 | COPS5 | Santa Cruz | sc-13157 | Mouse | 1:30 |
| **218** | Jagged1 | Jagged1 | JAG1 | Abcam | ab109536 | Rabbit | 01:50 |
| **219** | Jak2 | Jak2 | JAK2 | CST | 3230 | Rabbit | 1:750 |
| **220** | JNK (phospho T183/Y185) | JNK_pT183_Y185 | MAPK8 | CST | 4668 | Rabbit | 01:30 |
| **221** | JNK2 | JNK2 | MAPK9 | CST | 4672 | Rabbit | 1:25 |
| **222** | KAP1 | KAP1 | TRIM28 | Abcam | ab10484 | Rabbit | 1:2000 |
| **223** | KMT3A/HYPB/HIF-1 | SETD2 | SETD2 | abcam | ab184190 | Rabbit | 1:1000 |

| | | | | | | | |
|---|---|---|---|---|---|---|---|
| **224** | LAD1 | LAD1 | LAD1 | Atlas | HPA028732 | Rabbit | 1:500 |
| **225** | Lasu1/Ureb1 | Lasu1 | HUWE1 | Bethyl | IHC-00439 | Rabbit | 1:1000 |
| **226** | LC3A/B | LC3A-B | MAP1LC3A, B | CST | 4108 | Rabbit | 1:250 |
| **227** | Lck | Lck | LCK | CST | 2752 | Rabbit | 1:75 |
| **228** | LDHA | LDHA | LDHA | CST | 3582 | Rabbit | 1:250 |
| **229** | LRP6 (phospho S1490) | LRP6_pS1490 | LRP6 | CST | 2568 | Rabbit | 1:250 |
| **230** | MAPK (phospho T202/Y204) | MAPK_pT202/Y204 | MAPK1, 3 | CST | 4377 | Rabbit | 1:25 |
| **231** | Mcl-1 | Mcl-1 | MCL1 | CST | 5453 | Rabbit | 1:100 |
| **232** | MDM2 (phospho S166) | MDM2_pS166 | MDM2 | CST | 3521 | Rabbit | 1:60 |
| **233** | MEK1 | MEK1 | MAP2K1 | Epitomics/ Abcam | 1235-1/ ab32576 | Rabbit | 1:1500 |
| **234** | MEK1 (phospho S217/S221) | MEK1_p_S217/ S221 | MAP2K1, 2 | CST | 9154 | Rabbit | 1:50 |
| **235** | MEK2 | MEK2 | MAP2K2 | CST | 9125 | Rabbit | 1:50 |
| **236** | MelanA | MelanA | MLANA | Abcam | ab51061 | Rabbit | 1:500 |
| **237** | Melanoma gp100 | Melan-gp100 | PMEL | Abcam | ab137078 | Rabbit | 1:500 |
| **238** | MERIT40 | MERIT40 | MERIT40 | CST | 12711 | Rabbit | 1:3000 |
| **239** | MERIT40 (phospho S29) | MERIT40_pS29 | BABAM1 | CST | 12110 | Rabbit | 1:300 |
| **240** | Merlin/NF2 | Merlin | NF2 | Novus | 22710002 | Rabbit | 1:250 |
| **241** | MIF | MIF | MIF | Santa Cruz | sc-130329 | Rabbit | 1:100 |
| **242** | MITF (D5G7V) | MITF | MITF | CST | 12590 | Rabbit | 1:500 |
| **243** | Mitofusin-1 | Mitofusin-1 | MFN1 | CST | 14739 | Rabbit | 1:500 |
| **244** | Mitofusin-2 | Mitofusin-2 | MFN2 | CST | 11925 | Rabbit | 1:1000 |
| **245** | MLH1 (4C9C7) | MLH1 | MLH1 | CST | 3515 | Mouse | 1:500 |
| **246** | MLKL | MLKL | MLKL | CST | 14993 | Rabbit | 1:1000 |
| **247** | MMP2 | MMP2 | MMP2 | CST | 4022 | Rabbit | 1:75 |
| **248** | Mnk1 | Mnk1 | MKNK1 | CST | 2195 | Rabbit | 1:750 |
| **249** | Monocarboxylic Acid Transporter 4 | MCT4 | SLC16A4 | Millipore | AB3314P | Rabbit | 1:500 |
| **250** | MR1 | MR1 | MR1 | Santa Cruz | sc-377312 | Mouse | 1:500 |
| **251** | MRAP | MRAP | MRAP | Abcam | ab103319 | Rabbit | 1:500 |
| **252** | MSH2 (D24B5) | MSH2 | MSH2 | CST | 2017 | Rabbit | 1:750 |
| **253** | MSH6 | MSH6 | MSH6 | Novus | 22030002 | Rabbit | 1:1000 |
| **254** | MSI2 (EP1305Y) | MSI2 | MSI2 | Abcam | ab76148 | Rabbit | 1:1000 |

| | | | | | | | |
|---|---|---|---|---|---|---|---|
| **255** | MTCO1 | MTCO1 | MTCO1 | Abcam | ab14705 | Mouse | 1:500 |
| **256** | mTOR | mTOR | MTOR | CST | 2983 | Rabbit | 1:3000 |
| **257** | mTOR (phospho S2448) | mTOR_pS2448 | MTOR | CST | 2971 | Rabbit | 1:50 |
| **258** | MTSS1 | MTSS1 | MTSS1 | Novus | H00009788-M01A | Mouse | 1:250 |
| **259** | Myosin Heavy Chain 11 | Myosin-11 | MYH11 | Novus | 21370002 | Rabbit | 1:1000 |
| **260** | Myosin IIa | Myosin-IIa | MYH9 | CST | 3403 | Rabbit | 1:1000 |
| **261** | Myosin IIa (phospho S1943) | Myosin-IIa_pS1943 | MYH9 | CST | 5026 | Rabbit | 1:750 |
| **262** | Myt1 | Myt1 | PKMYT1 | CST | 4282 | Rabbit | 1:100 |
| **263** | NAPSIN-A | NAPSIN-A | NAPSA | Epitomics/ Abcam | 5795-1/ ab129189 | Rabbit | 1:150 |
| **264** | N-Cadherin | N-Cadherin | CDH2 | CST | 4061 | Rabbit | 1:25 |
| **265** | NDRG1 (phospho T346) | NDRG1_pT346 | NDRG1 | CST | 3217 | Rabbit | 01:50 |
| **266** | NDUFB4 | NDUFB4 | NDUFB4 | Abcam | ab110243 | Mouse | 1:25 |
| **267** | NF-kB p65 (phospho S536) | NF-kB-p65_pS536 | RELA | CST | 3033 | Rabbit | 1:30 |
| **268** | Notch1 | Notch1 | NOTCH1 | CST | 3268 | Rabbit | 01:30 |
| **299** | Notch1 (Cleaved) | Notch1-cleaved | NOTCH1 | CST | 4147 | Rabbit | 1:100 |
| **270** | Notch3 | Notch3 | NOTCH3 | Novus | H00004854-M01 | Mouse | 1:250 |
| **271** | NQO1 | NQO1 | NQO1 | CST | 3187 | Mouse | 1:15000 |
| **272** | N-Ras | N-Ras | NRAS | Santa Cruz | sc-31 | Mouse | 1:50 |
| **273** | NRF2 | NRF2 | NRF2 | CST | 12721 | Rabbit | 1:500 |
| **274** | Oct-4 | Oct-4 | POU5F1 | CST | 2750 | Rabbit | 1:40 |
| **275** | p16/INK4a | p16-INK4a | CDKN2A | Epitomics/ Abcam | 1712-1/ ab40803 | Rabbit | 1:500 |
| **276** | p21 | p21 | CDKN1A | Santa Cruz | sc-6246 | Rabbit | 1:150 |
| **277** | p27 (phospho T157) | p27_pT157 | CDKN1B | R&D Systems | AF1555 | Rabbit | 1:30 |
| **278** | p27 (phospho T198) | p27_pT198 | CDKN1B | Abcam | ab64949 | Rabbit | 01:50 |
| **279** | p27 KIP 1 | p27-Kip-1 | CDKN1B | Epitomics/ Abcam | 1591-1/ ab32034 | Rabbit | 01:40 |
| **280** | p38 (phospho T180/Y182) | p38_pT180_Y182 | MAPK11, 12, 13, 14 | CST | 9211 | Rabbit | 1:38 |
| **281** | p38 alpha MAPK | p38-a | MAPK1 | CST | 9228 | Mouse | 1:300 |
| **282** | p38 MAPK | p38-MAPK | MAPK11, 12, 14 | CST | 9212 | Rabbit | 1:1500 |

| | | | | | | | |
|---|---|---|---|---|---|---|---|
| **283** | p38/MAPK (phospho T180/Y182) | p38-MAPK_pT180_Y182 | MAPK14 | CST | 9215 | Rabbit | 1:250 |
| **284** | p44/42 MAPK | p44-42-MAPK | MAPK1, 3 | CST | 4695 | Rabbit | 1:2000 |
| **285** | p53 | p53 | TP53 | CST | 9282 | Rabbit | 1:2500 |
| **286** | p70 S6 Kinase (phospho T389) | p70-S6K_pT389 | RPS6KB1 | CST | 9205 | Rabbit | 1:50 |
| **287** | p70/S6K1 | p70-S6K1 | RPS6KB1 | Epitomics/ Abcam | 1494-1/ ab32529 | Rabbit | 1:300 |
| **288** | p90RSK (phospho T573) | p90RSK_pT573 | RPS6K | CST | 9346 | Rabbit | 1:25 |
| **289** | PAI-1 | PAI-1 | SERPINE1 | BD Biosciences | 612024 | Mouse | 1:50 |
| **290** | PAICS | PAICS | PAICS | Sigma-Aldrich | HPA035895 | Rabbit | 1:250 |
| **291** | PAK1 | PAK1 | PAK1 | CST | 2602 | Rabbit | 1:750 |
| **292** | PAK4 | PAK4 | PAK4 | CST | 3242 | Rabbit | 1:300 |
| **293** | PAR | PAR | PAR | Trevigen | 4336-BPC-100 | Rabbit | 1:30000 |
| **294** | PARG | PARG | PARG | CST | 66564 | Rabbit | 1:1000 |
| **295** | PARK7/DJ1 | DJ1 | PARK7 | Abcam | ab76008 | Rabbit | 1:5000 |
| **296** | PARP | PARP | PARP1 | CST | 9532 | Rabbit | 1:1000 |
| **297** | Patched | Patched | PTCH1 | Abcam | ab53715 | Rabbit | 1:1000 |
| **298** | Paxillin | Paxillin | PXN | CST | 2542 | Rabbit | 1:250 |
| **299** | P-Cadherin | P-Cadherin | CDH3 | CST | 2130 | Rabbit | 1:38 |
| **300** | PCNA | PCNA | PCNA | CST | 2586 | Mouse | 1:250 |
| **301** | PD-1 | PD-1 | PDCD1 | CST | 43248 | Mouse | 1:500 |
| **302** | Pdcd4 | Pdcd4 | PDCD4 | Rockland | 600-401-965 | Rabbit | 1:750 |
| **303** | PDGFRB | PDGFR-b | PDGFRB | Invitrogen | MA5-15143 | Rabbit | 1:500 |
| **304** | PDH | PDH | PDH | Abcam | ab110332 | Mouse | 1:100 |
| **305** | PDHK1 | PDHK1 | PDHK1 | CST | 3820 | Rabbit | 1:300 |
| **306** | PDK1 | PDK1 | PDPK1 | CST | 3062 | Rabbit | 01:50 |
| **307** | PDK1 (phospho S241) | PDK1_pS241 | PDPK1 | CST | 3061 | Rabbit | 01:50 |
| **308** | PD-L1 | PD-L1 | CD274 | CST | 13684 | Rabbit | 1:250 |
| **309** | PEA-15 | PEA-15 | PEA15 | CST | 2780S | Rabbit | 1:100 |
| **310** | PED/PEA-15 (phospho S116) | PEA-15_pS116 | PEA15 | Life Technologies | 44836G | Rabbit | 1:100 |
| **311** | PHGDH | PHGDH | PHGDH | CST | 13428 | Rabbit | 1:1000 |
| **312** | PI3 Kinase p110 alpha | PI3K-p110-a | PIK3CA | CST | 4255 | Rabbit | 1:50 |
| **313** | PI3K p110 beta | PI3K-p110-b | PIK3BC | Santa Cruz | sc-376412 | Mouse | 1:40 |

| | | | | | | | |
|---|---|---|---|---|---|---|---|
| **314** | PI3K p85 | PI3K-p85 | PIK3R1 | Millipore | 06-195 | Rabbit | 1:15000 |
| **315** | PKA RI alpha | PKA-a | PRKAR1A | CST | 5675 | Rabbit | 1:250 |
| **316** | PKC alpha/beta II (phospho T638/641) | PKC-a-b-II_pT638_T641 | PRKCA, B | CST | 9375 | Rabbit | 1:1000 |
| **317** | PKC(pan) beta II (phospho S660) | PKC-b-II_pS660 | PRKCA, B, D, E, H, Q | CST | 9371 | Rabbit | 1:200 |
| **318** | PKC delta (phospho S664) | PKC-delta_pS664 | PRKCD | Millipore | 07-875 | Rabbit | 1:75 |
| **319** | PKCalpha | PKCa | PRKCA | CST | 2056 | Rabbit | 1:200 |
| **320** | PKM2 | PKM2 | PKM | CST | 4053 | Rabbit | 1:300 |
| **321** | PLC gamma2 (phospho Y759) | PLC-gamma2_pY759 | PLCG2 | CST | 3874 | Rabbit | 01:25 |
| **322** | PLK1 | PLK1 | PLK1 | CST | 4513 | Rabbit | 1:125 |
| **323** | Met (phospho Y1234/Y1235) | c-Met_pY1234_ Y1235 | MET | CST | 3129 | Rabbit | 1:100 |
| **324** | PMS2 | PMS2 | PMS2 | Novus Biologicals | 22510002 | Rabbit | 1:1500 |
| **325** | PRAS40 | PRAS40 | AKT1S1 | Life Technologies | AHO1031 | Mouse | 1:75 |
| **326** | PRAS40 (phospho T246) | PRAS40_pT246 | AKT1S1 | Life Technologies | 441100G | Rabbit | 1:500 |
| **327** | PREX1 | PREX1 | PREX1 | Abcam | ab102739 | Rabbit | 1:100 |
| **328** | Progesterone Receptor [YR85] | PR | PGR | abcam | 206926 | Rabbit | 1:500 |
| **329** | PTEN | PTEN | PTEN | CST | 9552 | Rabbit | 1:500 |
| **330** | PTPN12 | PTPN12 | PTPN12 | Abcam | ab76942 | Rabbit | 1:500 |
| **331** | Puma | Puma | BBC3 | CST | 4976 | Rabbit | 1:50 |
| **332** | PYGB | PYGB | PYGB | Sigma-Aldrich | SAB2900066 | Rabbit | 1:750 |
| **333** | PYGM | PYGM | PYGM | Novus | H00005837-M10 | Mouse | 1:500 |
| **334** | Pyk2 (phospho Y402) | Pyk2_pY402 | PYK2 | CST | 3291 | Rabbit | 1:500 |
| **335** | Pyruvate Dehydrogenase | PDHA1 | PDHA1 | CST | 3205 | Rabbit | 1:200 |
| **336** | Rab11 | Rab11 | RAB11A, B | CST | 3539 | Rabbit | 1:30 |
| **337** | Rab25 | Rab25 | RAB25 | CST | 4314 | Rabbit | 1:30 |
| **338** | Rac1/Cdc42 | Cdc42 | CDC42 | CST | 4651 | Rabbit | 1:100 |
| **339** | Rad23A | Rad23A | RAD23A | CST | 24555 | Rabbit | 1:1000 |
| **340** | Rad50 | Rad50 | RAD50 | CST | 3427 | Rabbit | 1:250 |
| **341** | Rad51 | Rad51 | RAD51 | Millipore | ABE257 | Rabbit | 1:1000 |

| | | | | | | | |
|---|---|---|---|---|---|---|---|
| **342** | Raptor | Raptor | RPTOR | CST | 2280 | Rabbit | 1:300 |
| **343** | Rb | Rb | RB1 | CST | 9309 | Mouse | 1:150 |
| **344** | Rb (phospho S807/811) | Rb_pS807_S811 | RB1 | CST | 9308 | Rabbit | 1:1000 |
| **345** | RBM15 | RBM15 | RBM15 | Novus | 21390002 | Rabbit | 1:5000 |
| **346** | Rheb | Rheb | RHEB | R&D Systems | MAB3426 | Mouse | 1:75 |
| **347** | Rictor | Rictor | RICTOR | CST | 2114 | Rabbit | 1:100 |
| **348** | Rictor (phospho T1135) | Rictor_pT1135 | RICTOR | CST | 3806 | Rabbit | 1:200 |
| **349** | RIP | RIP | RIP | CST | 4926 | Rabbit | 1:75 |
| **350** | RIP3 | RIP3 | RIP3 | CST | 13526 | Rabbit | 1:500 |
| **351** | RPA32 (phospho S4/S8) | RPA32_pS4/S8 | RPA2 | Bethyl | A300-245A | Rabbit | 1:250 |
| **352** | RPA32/RPA2 | RPA32 | RPA2 | CST | 2208 | Rat | 1:150 |
| **353** | RRM1 | RRM1 | RRM1 | CST | 3388 | Rabbit | 1:100 |
| **354** | RRM2 | RRM2 | RRM2 | Life Technologies | PA527856 | Rabbit | 1:250 |
| **355** | RSK | RSK | RPS6KA1, 2, 3 | CST | 9347 | Rabbit | 1:150 |
| **356** | S100A4 | S100A4 | S100A4 | CST | 13018 | Rabbit | 1:1000 |
| **357** | S6 (phospho S235/236) | S6_pS235_S236 | RPS6 | CST | 2211 | Rabbit | 1:2500 |
| **358** | S6 (phospho S240/244) | S6_pS240_S244 | RPS6 | CST | 2215 | Rabbit | 1:1000 |
| **359** | S6 Ribosomal Protein | S6 | RPS6 | CST | 2317 | Mouse | 1:750 |
| **360** | SCD | SCD | SCD | Santa Cruz | sc-58420 | Mouse | 1:20 |
| **361** | SDHA | SDHA | SDHA | CST | 11998 | Rabbit | 1:250 |
| **362** | SFRP1 | SFRP1 | SFRP1 | CST | 4690 | Rabbit | 1:500 |
| **363** | Shc_pY317 | Shc_pY317 | SHC1 | CST | 2431 | Rabbit | 01:25 |
| **364** | SHP-2 (phospho Y542) | SHP-2_pY542 | PTPN11 | CST | 3751 | Rabbit | 1:75 |
| **365** | SHP2 / PTPN11 | SHP2 | PTPN11 | CST | 3397 | Rabbit | 1:250 |
| **366** | SLC1A5 | SLC1A5 | SLC1A5 | Sigma-Aldrich | HPA035240 | Rabbit | 1:150000 |
| **367** | Slfn11 | Slfn11 | SLFN11 | Santa Cruz | sc-374339 | Mouse | 1:150 |
| **368** | Smac | Smac | DIABLO | CST | 2954 | Mouse | 1:150 |
| **369** | Smad1 | Smad1 | SMAD1 | Epitomics/ Abcam | 1649-1/ ab33902 | Rabbit | 1:500 |
| **370** | Smad3 | Smad3 | SMAD3 | Epitomics/ Abcam | 1735-1/ ab40854 | Rabbit | 1:150 |
| **371** | Smad4 | Smad4 | SMAD4 | Santa Cruz | sc-7966 | Mouse | 1:30 |

| | | | | | | | |
|---|---|---|---|---|---|---|---|
| **372** | Snail | Snail | SNAI1 | CST | 3895 | Mouse | 1:50 |
| **373** | SOD1 | SOD1 | SOD1 | CST | 4266 | Mouse | 1:500 |
| **374** | SOD2 (D9V9C) | SOD2 | SOD2 | CST | 13194 | Rabbit | 1:200 |
| **375** | Sox2 | Sox2 | SOX2 | CST | 2748 | Rabbit | 1:50 |
| **376** | Src | Src | SRC | Millipore | 05-184 | Mouse | 1:50 |
| **377** | Src (phospho Y416) | Src_pY419 | SRC | CST | 2101 | Rabbit | 1:25 |
| **378** | Src (phospho Y527) | Src_pY527 | SRC | CST | 2105 | Rabbit | 1:150 |
| **379** | SRSF1/SF2 | SF2 | SRSF1 | Invitrogen | 324500 | Mouse | 1:75 |
| **380** | Stat3 | Stat3 | STAT3 | CST | 4904 | Rabbit | 1:3000 |
| **381** | Stat3 (phospho Y705) | Stat3_pY705 | STAT3 | CST | 9145 | Rabbit | 1:100 |
| **382** | Stat5a | Stat5a | STAT5A | Epitomics/ Abcam | 1289-1/ ab32043 | Rabbit | 1:300 |
| **383** | Stathmin-1 | Stathmin-1 | STMN1 | Epitomics/ Abcam | 1972-1/ ab52630 | Rabbit | 1:75 |
| **384** | STING | STING | TMEM173 | CST | 13647 | Rabbit | 1:250 |
| **385** | Syk | Syk | SYK | Santa Cruz | sc-1240 | Mouse | 1:500 |
| **386** | Tau | Tau | MAPT | Millipore | 05-348 | Mouse | 1:100 |
| **387** | TAZ | TAZ | WWTR1 | CST | 4883 | Rabbit | 1:300 |
| **388** | TFAM | TFAM | TFAM | CST | 7495 | Rabbit | 1:300 |
| **389** | Transferrin R | TFRC | TFRC | Novus | 22500002 | Rabbit | 1:15000 |
| **390** | TIGAR | TIGAR | TIGAR | Epitomics/ Abcam | S1711/ ab137573 | Rabbit | 1:100 |
| **391** | Transglutaminase | Transglutaminase | TGM2 | Lab Vision | MS-224-P1 | Mouse | 1:150 |
| **392** | TRIM25 | TRIM25 | TRIM25 | Abcam | ab167154 | Rabbit | 1:3000 |
| **393** | TTF1 | TTF1 | NKX2-1 | Epitomics/ Abcam | 2044-1/ ab76013 | Rabbit | 1:150 |
| **394** | Tuberin | Tuberin | TSC2 | Epitomics/ Abcam | 1613-1/ ab32554 | Rabbit | 1:2500 |
| **395** | Tuberin/TSC2 (phospho T1462) | Tuberin_pT1462 | TSC2 | CST | 3617 | Rabbit | 1:38 |
| **396** | TUFM | TUFM | TUFM | Abcam | ab173300 | Rabbit | 1:38 |
| **397** | TWEAK Receptor/FN14 | FN14 | TNFRSF12A | CST | 4403 | Rabbit | 1:1000 |
| **398** | TWIST | TWIST | TWIST1 | Santa Cruz | sc-81417 | Mouse | 1:30 |
| **399** | Tyro3 | Tyro3 | TYRO3 | CST | 5585 | Rabbit | 1:30 |
| **400** | UBAC1 | UBAC1 | UBAC1 | Sigma-Aldrich | HPA005651 | Rabbit | 1:250 |
| **401** | Ubiquityl-Histone H2B | U-Histone-H2B | HIST1H2BB | CST | 5546 | Rabbit | 1:500 |

| | | | | | | | |
|---|---|---|---|---|---|---|---|
| **402** | UGT1A | UGT1A | UGT1A1, 3, 4, 5, 7, 8, 10 | Santa Cruz | sc-271268 | Mouse | 1:75 |
| **403** | ULK1 (phospho S757) | ULK1_pS757 | ULK1 | CST | 6888 | Rabbit | 1:300 |
| **404** | UQCRC2 | UQCRC2 | UQCRC2 | MitoSciences / Abcam | MS304/ ab14745 | Mouse | 01:50 |
| **405** | UVRAG | UVRAG | UVRAG | CST | 13115 | Rabbit | 1:100 |
| **406** | VASP | VASP | VASP | CST | 3112 | Rabbit | 1:100 |
| **407** | Vav1 | Vav1 | VAV1 | CST | 2502 | Rabbit | 1:500 |
| **408** | VDAC1/Porin | Porin | VDAC1 | Abcam | ab14734 | Mouse | 1:100 |
| **409** | VEGF Receptor 2 | VEGFR-2 | KDR | CST | 2479 | Rabbit | 1:3000 |
| **410** | VHL/EPPK1 | VHL-EPPK1 | EPPK1 | BD Biosciences | 556347 | Mouse | 1:1500 |
| **411** | Vimentin | Vimentin | VIM | Dako/Fisher | M0725/ MS-129-P | Mouse | 1:250 |
| **412** | Vinculin | Vinculin | VCL | Sigma-Aldrich | SAB4200080 | Mouse | 1:25000 |
| **413** | Wee1 | Wee1 | WEE1 | CST | 4936 | Rabbit | 1:250 |
| **414** | Wee1 (phospho S642) | Wee1_pS642 | WEE1 | CST | 4910 | Rabbit | 1:50 |
| **415** | WIPI1 | WIPI1 | WIPI1 | CST | 12124 | Rabbit | 1:150 |
| **416** | WIPI2 | WIPI2 | WIPI2 | CST | 8567 | Rabbit | 1:150 |
| **417** | XBP-1 | XBP-1 | XBP1 | Santa Cruz | sc-32136 | Goat | 1:200 |
| **418** | XIAP | XIAP | XIAP | CST | 2042 | Rabbit | 1:100 |
| **419** | XPA | XPA | XPA | Santa Cruz | sc-56813 | Mouse | 1:75 |
| **420** | XPF | XPF | ERCC4 | Abcam | ab73720 | Rabbit | 1:100 |
| **421** | XPG | ERCC5 | ERCC5 | Proteintech Group | 11331-1-AP | Rabbit | 1:250 |
| **422** | XRCC1 | XRCC1 | XRCC1 | CST | 2735 | Rabbit | 1:20 |
| **423** | YAP | YAP | YAP1 | Santa Cruz | sc-376830 | Mouse | 1:300 |
| **424** | YAP (phospho S127) | YAP_pS127 | YAP1 | CST | 4911 | Rabbit | 1:250 |
| **425** | YB1 (phospho S102) | YB1_pS102 | YBX1 | CST | 2900 | Rabbit | 1:50 |
| **426** | ZAP-70 | ZAP-70 | ZAP70 | CST | 2705 | Rabbit | 1:500 |

**Table S7:** Cell cycle distribution of MGT11 treated cells - Statistical analysis was performed by two-way ANOVA followed by Tukey test.

| | | no | A11 | R547 | A11 + R547 |
|---|---|---|---|---|---|
| **Sub G0** | no | | ns | ns | * |
| | A11 | | | ns | ns |
| | R547 | | | | ns |
| | A11 + R547 | | | | |
| **G0** | no | | ns | ns | ns |
| | A11 | | | ns | ns |
| | R547 | | | | ns |
| | A11 + R547 | | | | |
| **G1** | no | | ns | ns | ns |
| | A11 | | | ns | ns |
| | R547 | | | | ns |
| | A11 + R547 | | | | |
| **S** | no | | ns | ns | ns |
| | A11 | | | ns | ns |
| | R547 | | | | * |
| | A11 + R547 | | | | |
| **G2** | no | | ns | ** | ns |
| | A11 | | | ** | ns |
| | R547 | | | | *** |
| | A11 + R547 | | | | |
| **M** | no | | ns | ns | ns |
| | A11 | | | ns | ns |
| | R547 | | | | ns |
| | A11 + R547 | | | | |

**Table S8:** List of antibodies used in the study.

| Antibody | Company | Reference | Dilution | Used for | TritonX-100 (%) for IF |
|---|---|---|---|---|---|
| FOX O3A | CliniSciences | 40937 | 1 :500 | IF | 0.5 % |
| $pS_{139}$H2AX (γH2AX) | Cell Signaling | 9718 | 1:400 | IF | 0.2 % |
| Alexa488-conjugated Phalloidin | ThermoFisher Scientific | A12379 | 1:20 | IF | 0.5% |
| pH3 (S10) | Millipore | 06-570 | 1 :500 | IF | 0.3 % |
| alpha-TUBULIN | Sigma | T5168 | 1 :5000 | IF | 0.3 % |
| ACTIN | Sigma | A3853 | 1:6000 | WB | |
| AKT | Cell Signaling | 9272 | 1:2000 | WB | |
| $pS_{473}$AKT | Cell Signaling | 9271 | 1:2000 | WB | |
| ATM | Cell Signaling | 2873 | 1:1000 | WB | |
| $pS_{1987}$ATM | Invitrogen | PA5-37346 | 1:1000 | WB | |
| ATR | Cell Signaling | 13934 | 1:1000 | WB | |
| $pS_{428}$ATR | Cell Signaling | 2853 | 1:1000 | WB | |
| BCL-XL | Transduction Laboratory | B22620 | 1:500 | WB | |
| BIM | Santa Cruz | sc-11425 | 1:1000 | WB | |
| Cleaved-CASPASE 3 | Cell Signaling | 9661 | 1:1000 | WB | |
| CDC2 (CDK1) | Cell Signaling | 28493 | 1:20000 | WB | |
| $pY_{15}$CDC2(CDK1) | Cell Signaling | 4539 | 1:1000 | WB | |
| $pT_{202}/Y_{204}$ERKs | Cell Signaling | 9106 | 1:10000 | WB | |
| ERKs | Cell Signaling | 9102 | 1:10000 | WB | |
| FOX M1 | CliniSciences | 32671 | 1:1000 | WB | |
| $pT_{600}$FOX M1 | CliniSciences | 13207 | 1:1000 | WB | |
| FOX O3A | CliniSciences | 40937 | 1 :1000 | WB | |
| $pY_{627}$GAB1 | Cell Signaling | 3231 | 1:2000 | WB | |
| GAB1 | Upstate | 6579 | 1:1000 | WB | |
| $pS_{139}$H2AX (γH2AX) | Cell Signaling | 9718 | 1:1000 | WB | |
| MCL1 | Santa Cruz | sc-819 | 1:1000 | WB | |
| $MET^{25H2}$ | Cell Signaling | 3127 | 1:1000 | WB | |
| $pY_{1234/35}$MET | Cell Signaling | 3126 | 1:2000 | WB | |
| PARP | Cell Signaling | 9546S | 1:2000 | WB | |
| P53 | Novocastra | CM5 | 1:1000 | WB | |
| $pS_{15}$P53 | Cell Signaling | 9284 | 1:1000 | WB | |
| RB | Cell Signaling | 9313 | 1:1000 | WB | |

| | | | | |
|---|---|---|---|---|
| $pS_{780}RB$ | Cell Signaling | 9307 | 1:1000 | WB |
| $pS_{795}RB$ | Abcam | Ab47474 | 1:1000 | WB |
| RRM2 | abbexa | Abx004031 | 1 :25000 | WB |
| XIAP | Transduction laboratory | 610716 | 1:3000 | WB |
| Goat anti-rabbit IgG-peroxidase | Jackson Immuno Research | 115-035-144 | 1:4000 | WB |
| Goat anti-mouse IgG-peroxidase | Jackson Immuno Research | 115-035-146 | 1:4000 | WB |
| anti-Ki67-APC (clone SolA15) | eBioscience | 17-5698-82 | 1:200 | FACS |

**Table S9:** Drugs used in this study, with the indicated targets and the concentrations used.

| Drug | Company | Target | Concentration (µM) |
|---|---|---|---|
| A-1155463 | Selleckchem/Targetmol | Bcl-xL | 0.3, 1, 3, 10 |
| Adavosertib | Selleckchem | Wee1 | 1, 3, 10 |
| Adavosertib(MK-1775) | Targetmol | Wee1 | 1, 3, 10 |
| Alisertib | Selleckchem | Aurora A | 1, 3, 10 |
| Barasertib | Selleckchem | Aurora B | 1, 3, 10 |
| FDI-6 | Axon Medchem | FOXM1 | 0.3, 1, 3 |
| R547 | Selleckchem/Sigma | Cdk1/2/4 | 1, 3, 10 |

**Table S10:** Oligonucleotides used for RT-qPCR experiments.

| Oligonucleotide (name) | Sequence (Forward) | Sequence (Reverse) |
|---|---|---|
| *PUMA* | ACCGCTCCACCTGCCGTCAC | ACGGGCGACTCTAAGTGCTGC |
| *Fas-L* | GAAGGAACTGGCAGAACTCCGT | GCCACACTCCTCGGCTCTTTTT |
| *B2M* | ACAGTTCCACCCGCCTCACATT | TAGAAAGACCAGTCCTTGCTGAAG |

### Full unedited gels for Figures 4 and 5

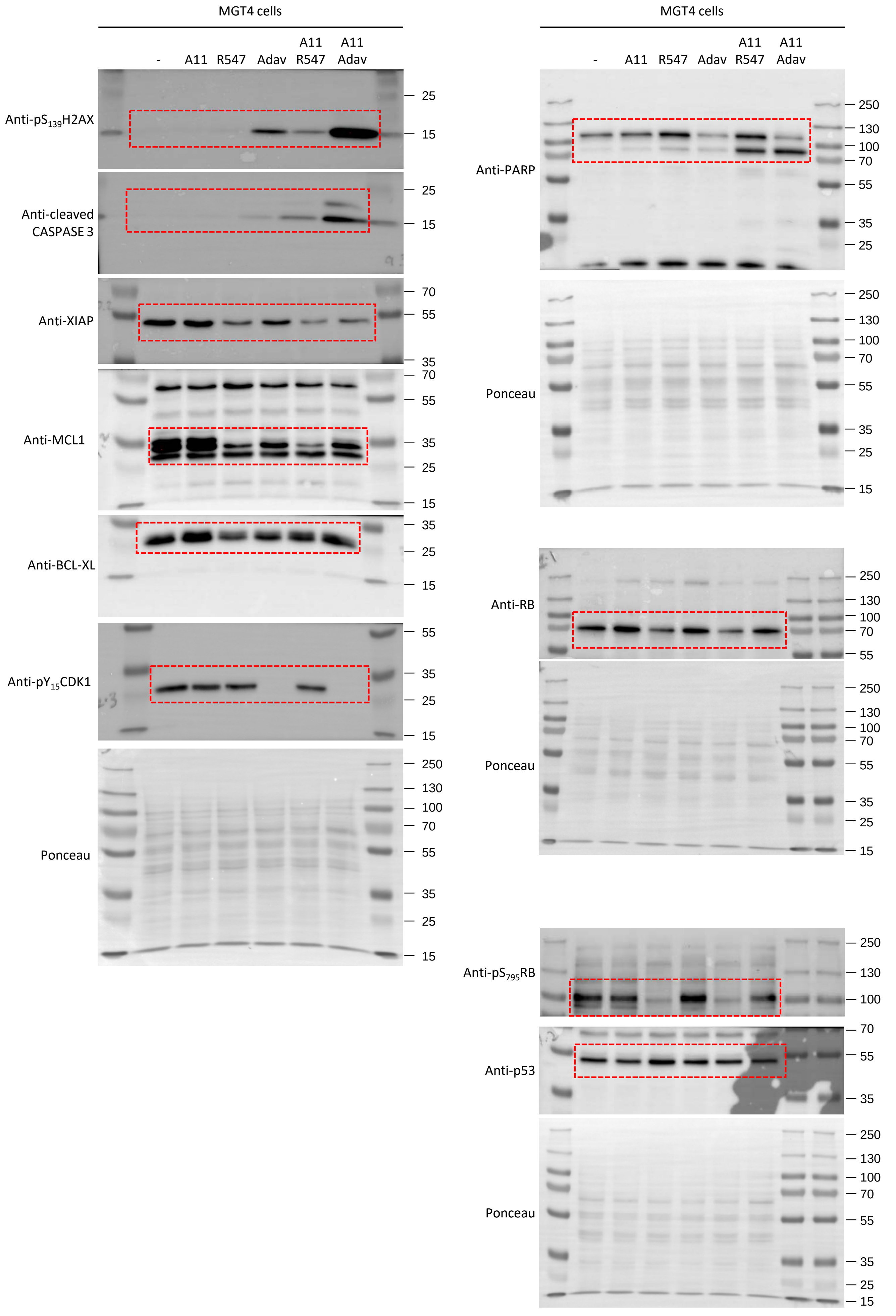

Full unedited gels for Figures 4 and 5

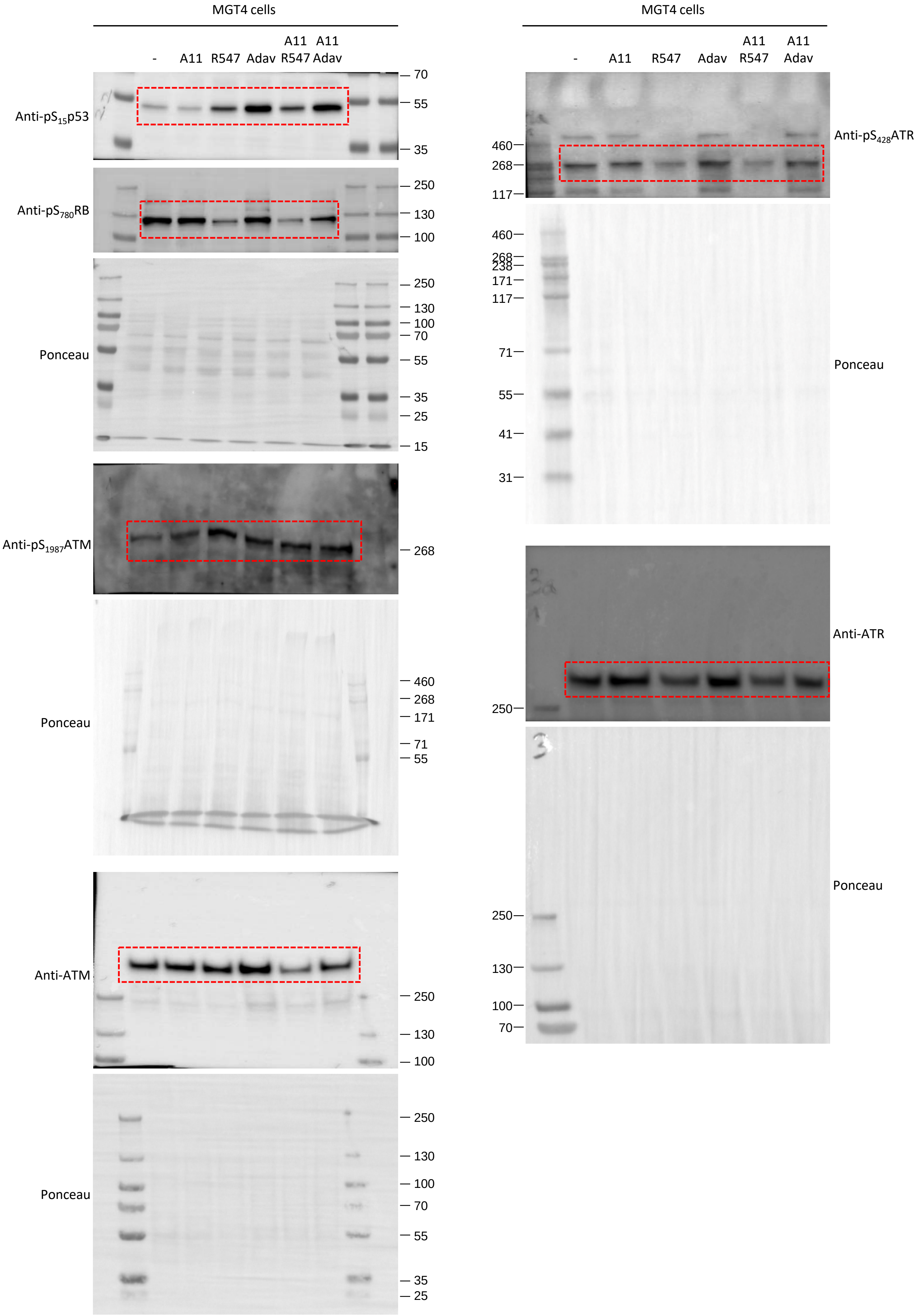

### Full unedited gels for Figures 4 and 5

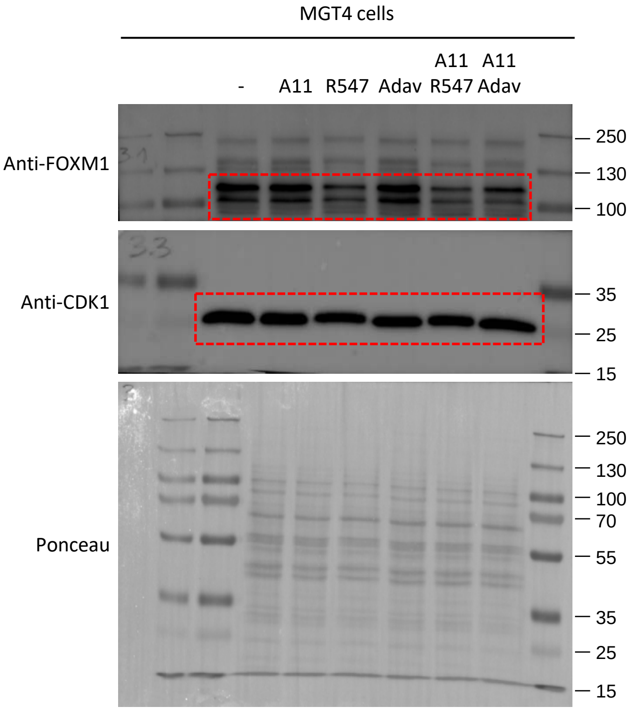

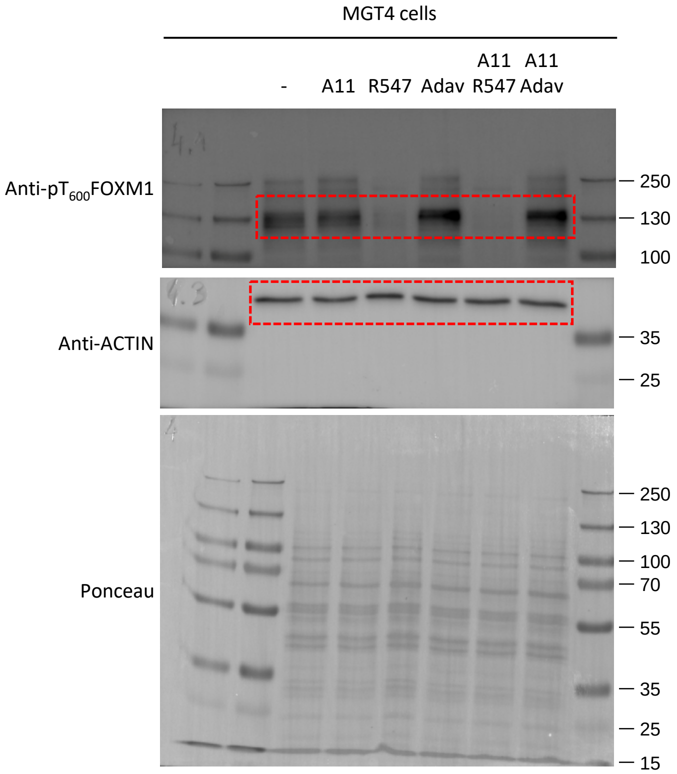

Full unedited gels for Figures 4 and 5

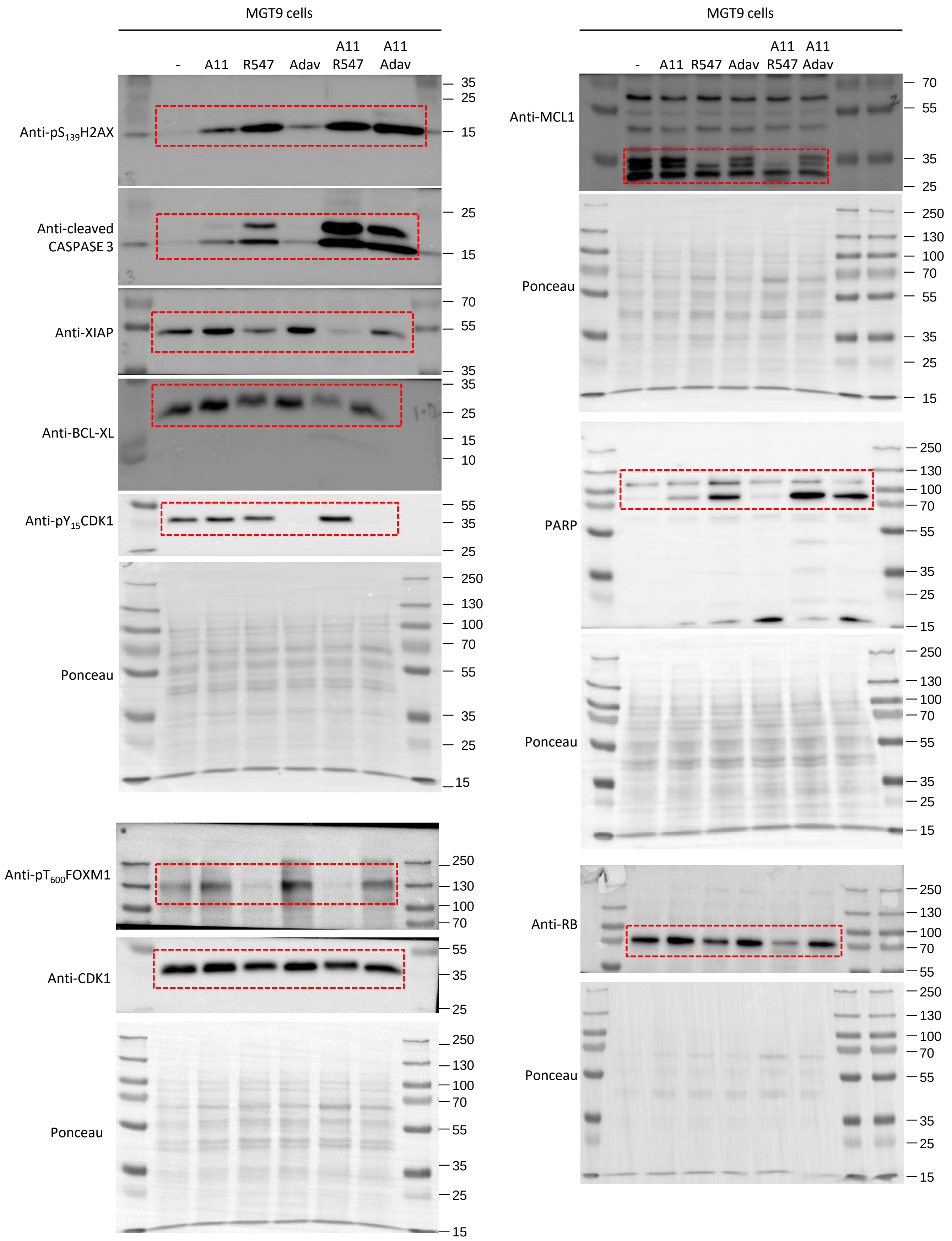

### Full unedited gels for Figures 4 and 5

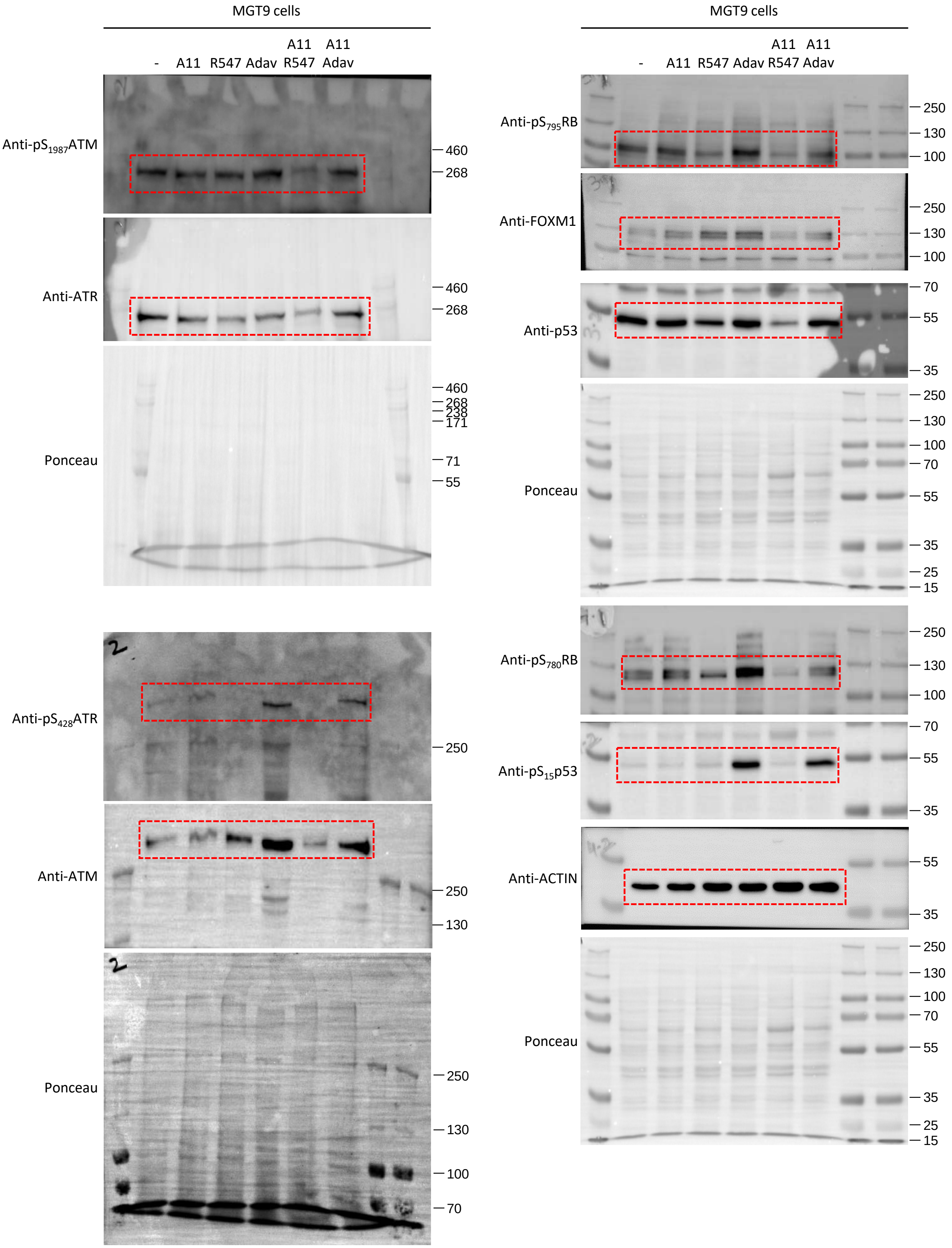

### Full unedited gels for Figures 4 and 5

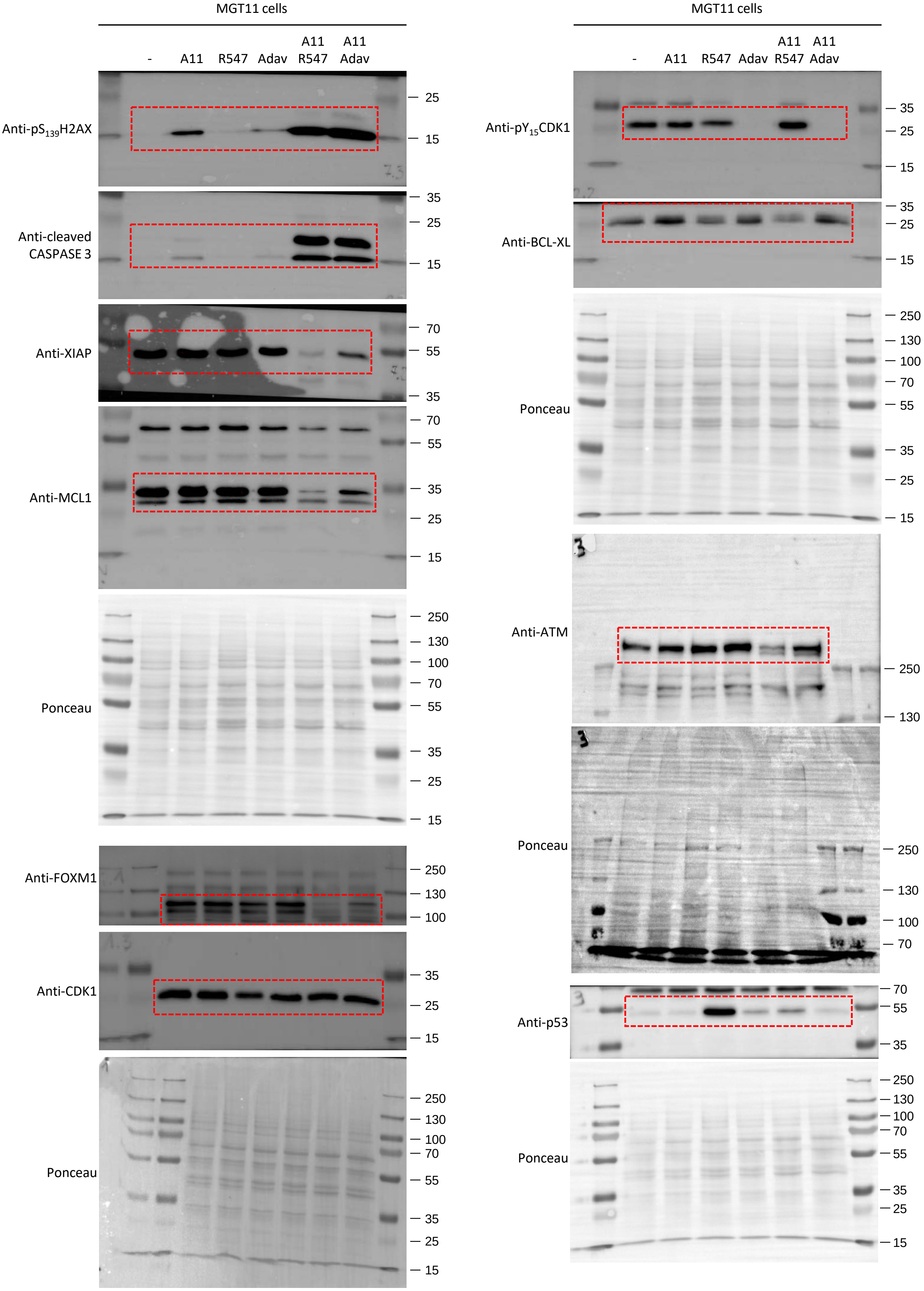

### Full unedited gels for Figures 4 and 5

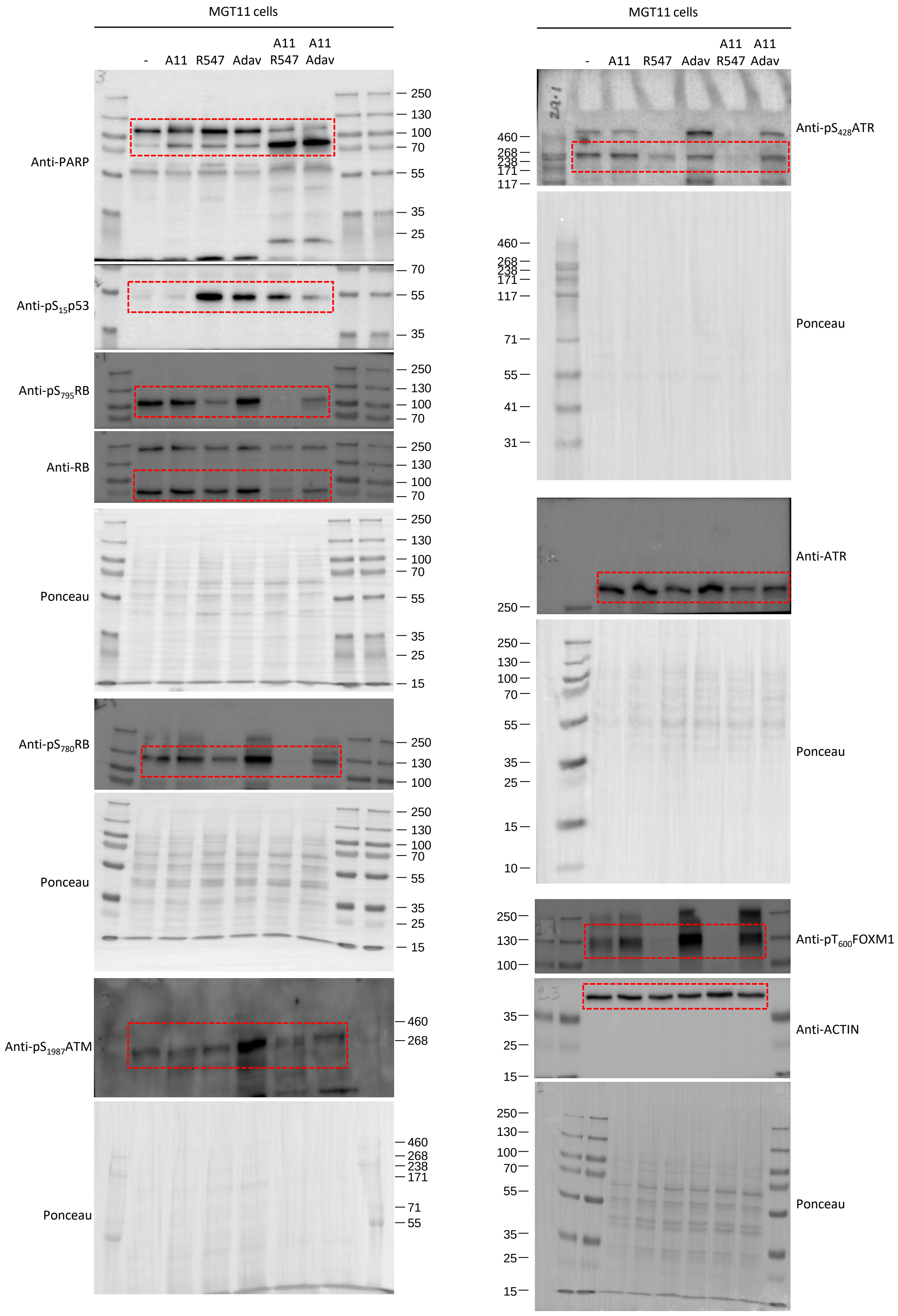

### Full unedited gels for Figure S3

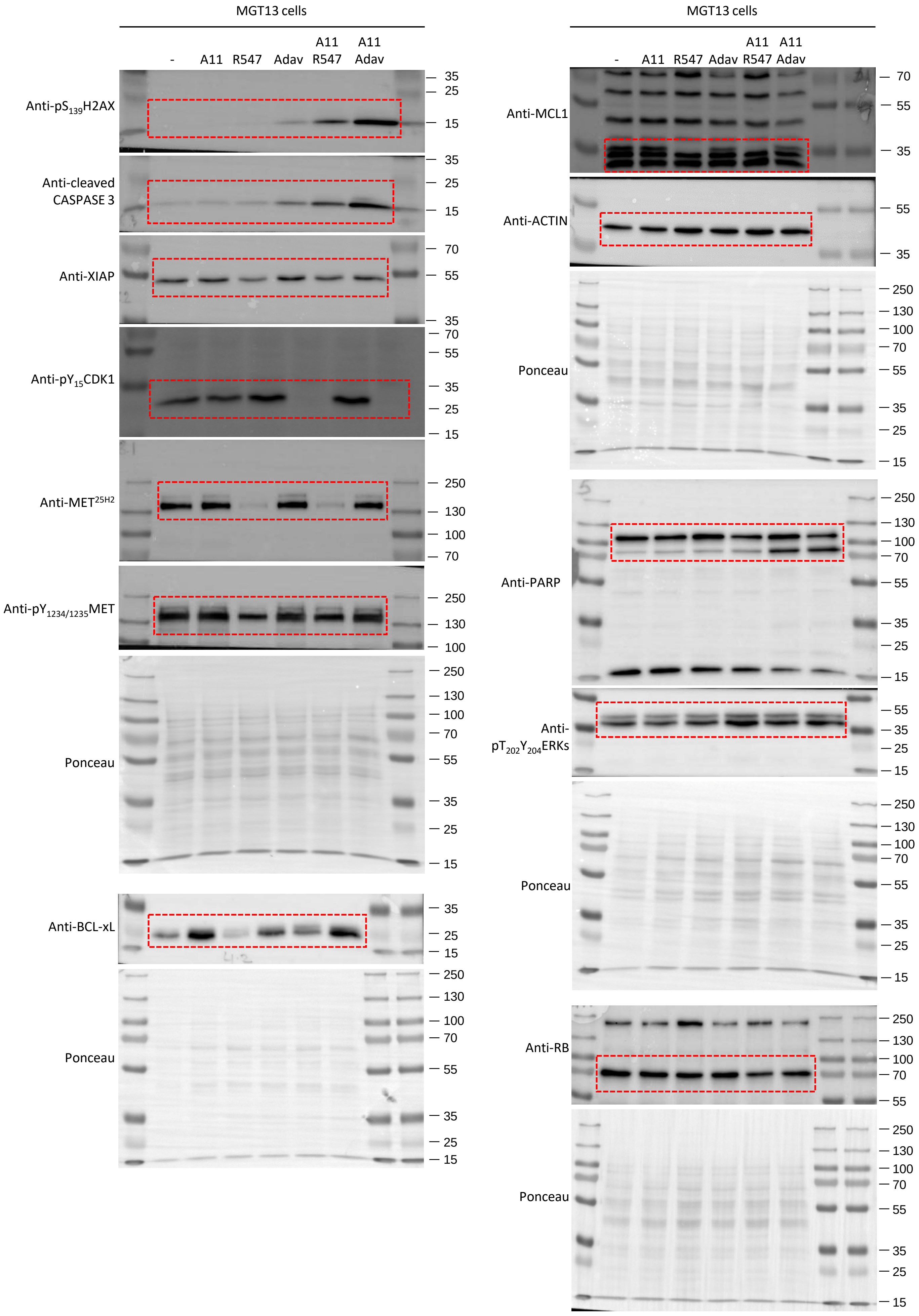

### Full unedited gels for Figure S3

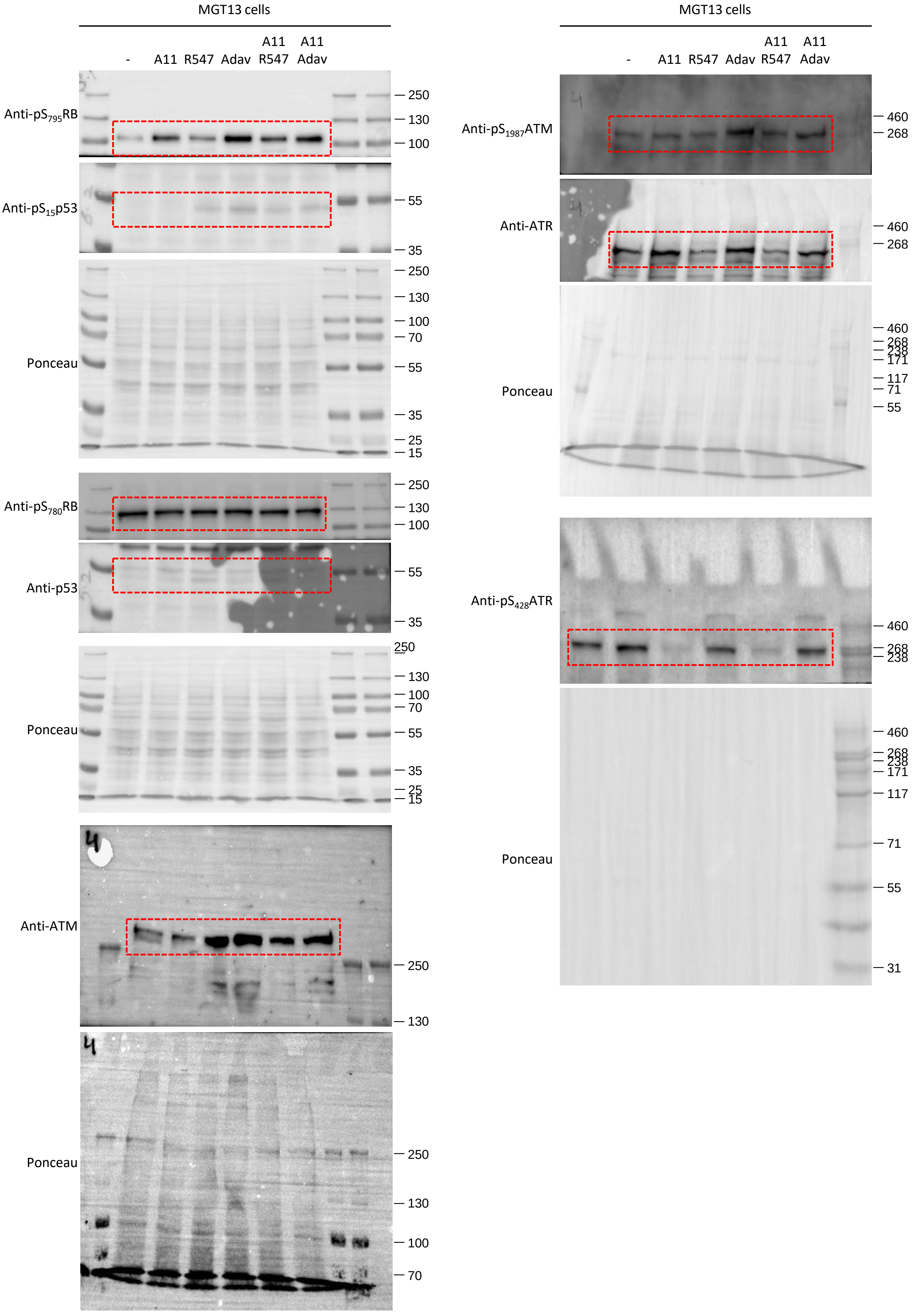

Full unedited gels for Figure 7A

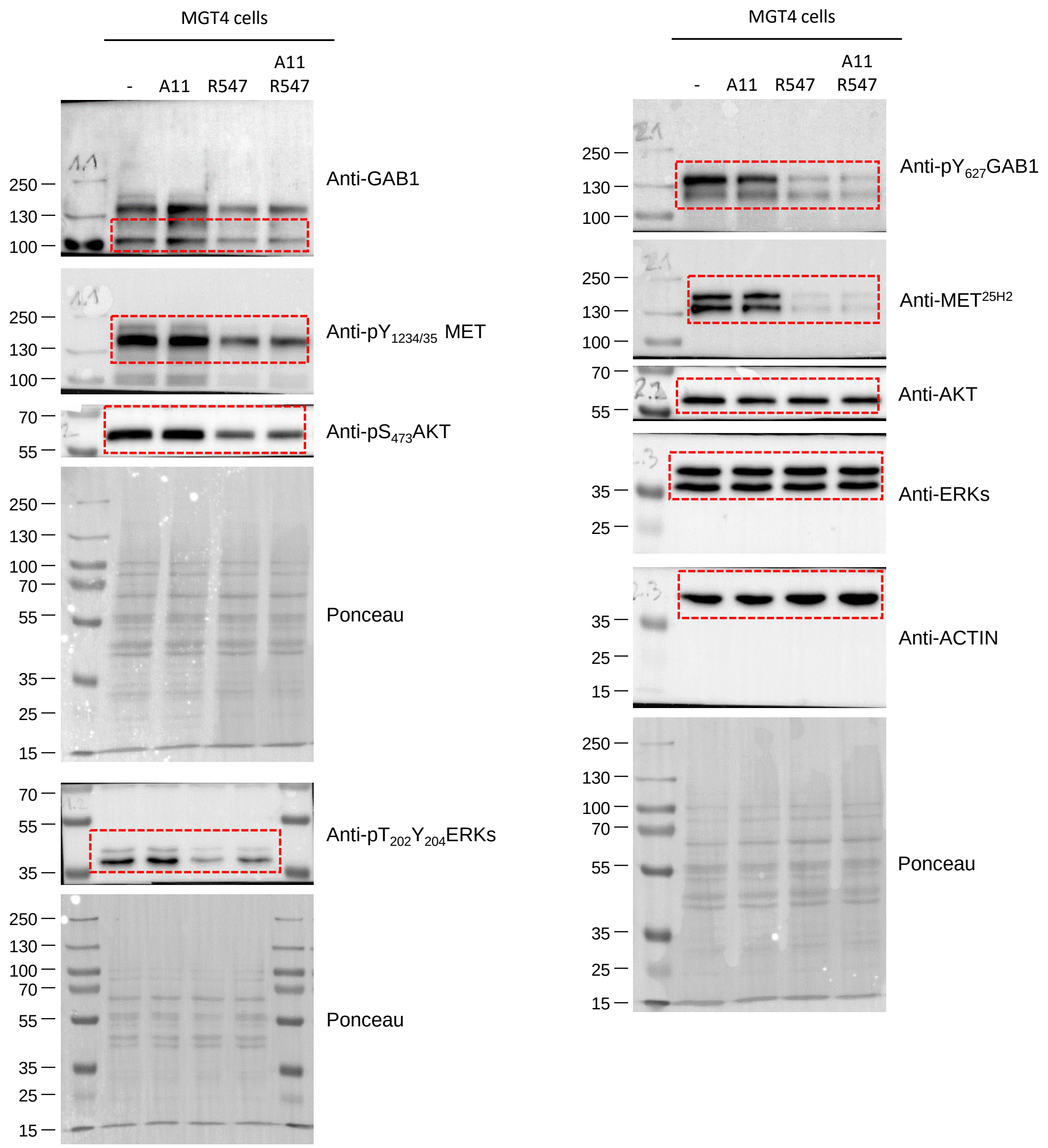

Full unedited gels for Figure 7A

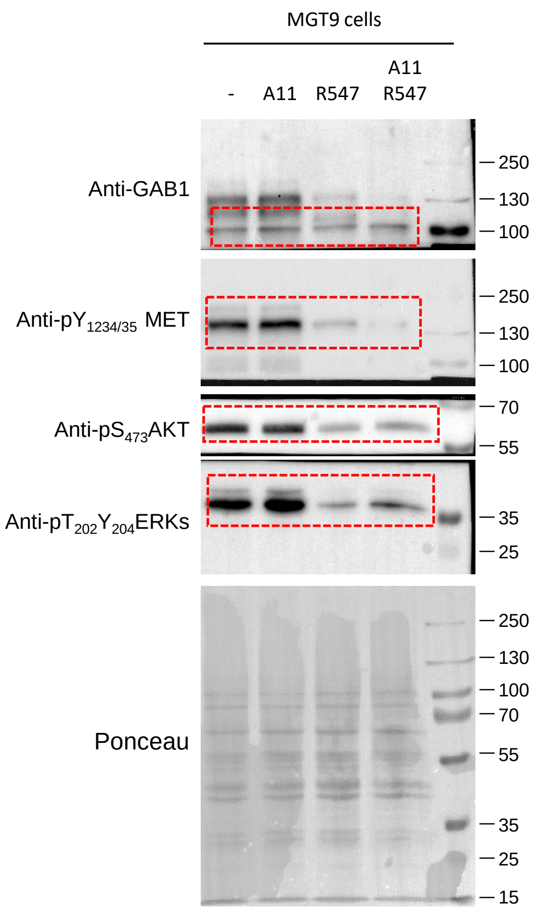

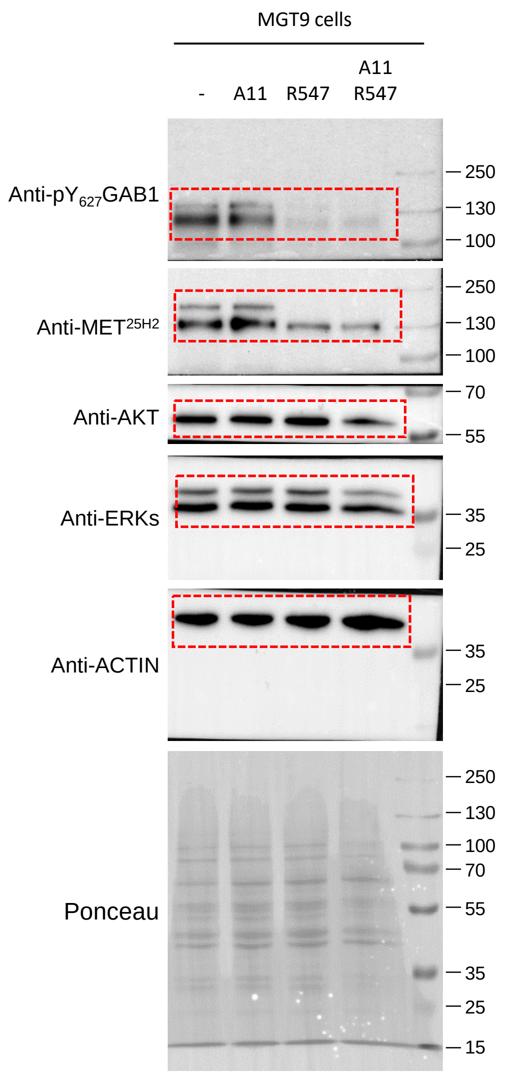

### Full unedited gels for Figure 7A

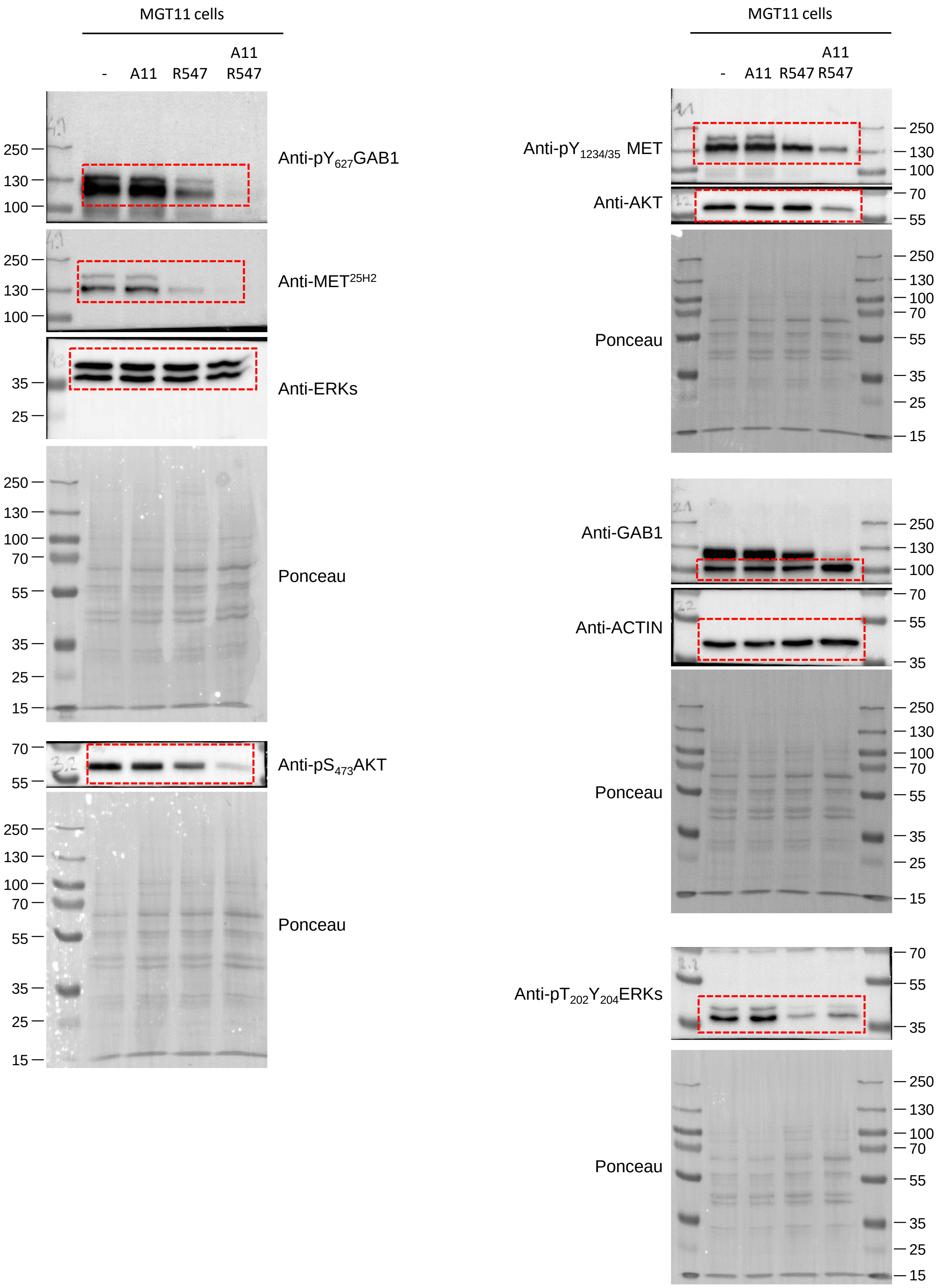

Full unedited gels for Figure 7C

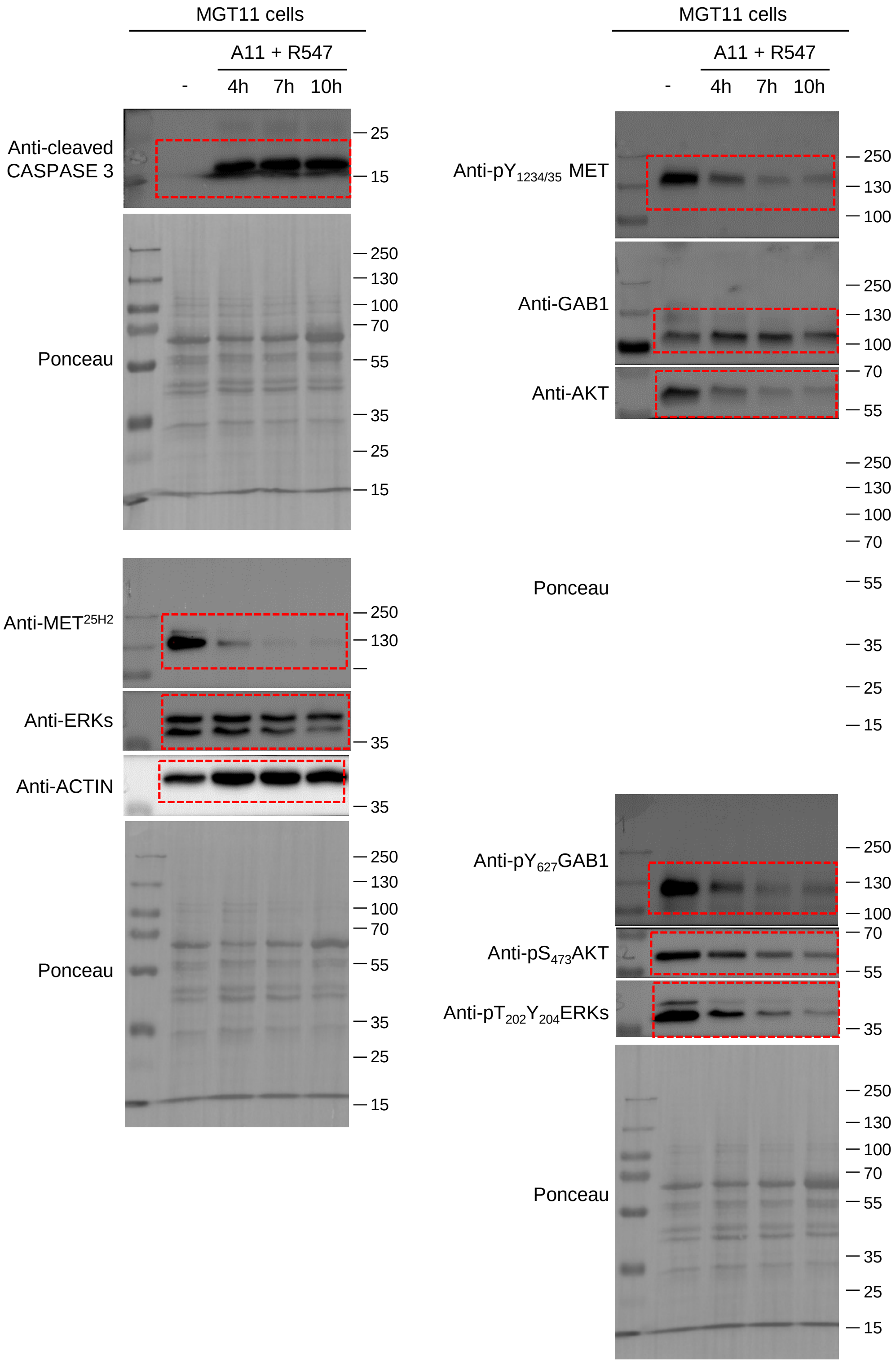
